# Mapping Chemical Diversity: Descriptor-Guided Clustering of Natural Products in the COCONUT Database

**DOI:** 10.64898/2026.06.03.729746

**Authors:** G Shreyasree, Adithya Dileep, Akhileshwar Namani, Prashantha Karunakar

**Affiliations:** Department of Biotechnology, Dayananda Sagar College of Engineering (Affiliated to Visvesvaraya Technological University, Belagavi), Kumaraswamy Layout, Bangalore, Karnataka 560 111, India; Department of Molecular Oncology, Sri Shankara Cancer Hospital and Research Centre, Sri Shankara National Centre for Cancer Prevention and Research, Sri Shankara Cancer Foundation, Bangalore 560004, Karnataka, India

**Keywords:** Natural product chemical space, molecular clustering, descriptor selection, cluster analysis, scaffold diversity, representative analysis, drug discovery

## Abstract

Natural products represent a major source of bioactive compounds for drug discovery, yet their exploration remains challenging due to extensive structural complexity and scaffold diversity. Using the COCONUT database, we developed a cluster-oriented framework to systematically map and characterize the natural product chemical space through feature engineering, molecular clustering, and representative-based analysis. Descriptor selection identified a greedy maximum coverage strategy with a 0.35–0.85 correlation threshold range and 20 descriptors as the optimal feature set, enriched in physicochemical and graph-topological properties. Comparative evaluation of clustering approaches identified UMAP-HDBSCAN as the best-performing pipeline, generating 1,683 clusters with silhouette scores of 0.42 before and 0.24 after noise reassignment. Cluster profiling revealed a highly heterogeneous scaffold landscape, with 67.56% of clusters exhibiting low scaffold dominance and only 15.21% representing highly scaffold-dominated regions, supporting a chemical space composed largely of interconnected transitional clusters. Descriptor analyses showed that natural product clusters were generally enriched in saturated, low-aromaticity chemotypes with moderate lipophilicity and constrained molecular flexibility. Representative-based analyses demonstrated that central representatives (medoid and centroid-closest molecules) closely captured cluster-average properties, whereas diverse representatives better reflected structural breadth, findings further supported through descriptor-based and docking-based validation. Collectively, the results reinforce the natural product chemical space as a continuous yet structured manifold and provide a representative-guided framework for its efficient exploration in drug discovery applications. The complete data can be accessed at: https://github.com/shrek-28/DescriptorClusteringNPSpace

## 1. Introduction

Natural products have historically served as one of the most productive sources of therapeutic agents, due to their extensive structural diversity and biological specificity, which provide valuable scaffolds for low molecular weight drug discovery. Several clinically important drugs, including aspirin, morphine, digitoxin, and quinine, originated from natural sources such as plants, fungi, and marine organisms, and they continue to demonstrate the pharmacological relevance of natural compounds in modern medicine (Dias et al., 2012). A substantial proportion of drugs approved by regulatory agencies such as the FDA and EMA are either natural products or their derivatives. This highlights their sustained contribution to drug development. Consequently, computer-aided drug discovery strategies are increasingly being integrated into natural product research to accelerate lead identification, target screening, and early-stage toxicity assessment (Thomford et al., 2018). The increasing availability of large-scale natural product repositories has further expanded the opportunities for computational exploration of the natural product chemical space. The COCONUT (Collection of Open Natural Products) database represents one of the most comprehensive open-access repositories of natural products, providing extensive structural and chemical information on naturally occurring compounds (Chandrasekhar et al., 2025; Sorokina et al., 2021). The scale and chemical diversity of such datasets have consequently increased the importance of chemoinformatics and computational approaches for systematic analysis and organization of the natural product space.

Feature selection and molecular clustering are necessary components of chemical space analysis in computer-aided drug discovery and virtual screening pipelines (Pérez et al., 2025). Clustering of the chemical space helps in development of predictive models, representative sampling of chemical subspaces and identification of underrepresented spaces. Additionally, within natural product research, clustering allows the study of aggregation patterns within the chemical space, and aids the identification of privileged regions associated with specific biological target classes (Talevi & Bellera, 2024). Feature selection is a crucial preliminary step to clustering, helping reduce large descriptor spaces into small, informative and relevant features that improve clustering quality and interpretability of the model (Huang et al., 2023). Existing approaches for clustering-based feature selection broadly include: filter methods, which rank descriptors independently of clustering algorithms; wrapper methods, which perform iterative clustering to evaluate descriptor subsets; and hybrid methods, which combine descriptor filtering and subsequent cluster-quality optimization (Alelyani et al., 2019). Several studies have proposed specialized descriptor selection strategies for molecule datasets. A study by Böcker et al. utilized an integrated approach with Shannon entropy and unsupervised forward selection (UFS) to ensure that relevant and low-redundancy descriptors were retained in the dataset. Similarly, a graph-based approach to deep clustering of molecules, conducted by Hadipour et al. employed a comprehensive custom feature engineering pipeline, combining atom and bond-level descriptors followed by dimensionality reduction through PCA. This combined local and global molecular features to remove redundant variables. Molecular clustering approaches using physicochemical descriptors have also proved to have practical applications in lead prioritization. A study by Voicu et al. employed a hierarchical clustering approach using the rcdk and cluster packages in R, reducing pyrazole-derived molecules from 23 to 4. This substantially decreased downstream synthesis and screening approaches. In a study involving natural product lead databases (NPLDs), clustered distribution analyses showed that specific compound classes were preferentially associated with privileged binding-site classes such as amine, nucleobase, nucleoside and amino acid phosphate targets. This highlights the utility of clustering for understanding functional organization within the natural product chemical space (Tao et al., 2015a).

Despite feature selection and molecular clustering being extensively used in cheminformatics and virtual screening workflows, several limitations remain. Feature selection in unsupervised learning is largely difficult due to the absence of class labels, which makes it challenging to differentiate between relevant, redundant and misleading features (Huang et al., 2023). Small changes in dataset composition, initialization or hyperparameters can produce different descriptor subsets, which leads to instability and reduced reproducibility in the clustering outcome. Descriptor pre-filtering can remove variables that may be informative when combined with other descriptors, while descriptor subsets selected for one clustering algorithm may not perform equally well with different clustering approaches. Another challenge is the trade-off between interpretability and clustering performance, as highly interpretable descriptors do not always guarantee the generation of clear clusters (Di Martino & Senatore, 2021; Witten & Tibshirani, 2010). Existing clustering approaches also present limitations in terms of practicality. Similarity-based approaches, such as the Taylor–Butina algorithm, scale poorly with increasing dataset size due to their high computational and memory requirements, limiting their applicability to large chemical libraries. Hierarchical clustering methods are constrained by irreversible merging or splitting decisions made during cluster construction, whereas non-hierarchical approaches require the number of clusters and initialization parameters to be specified beforehand, both of which can substantially influence the resulting cluster structure (Talevi & Bellera, 2024). Hence, although several studies have elucidated the clustering of small molecules and natural products using fingerprints, graph representations and mixed descriptor spaces, there remains limited work which specifically focuses on stable and interpretable natural product clustering using descriptors alone.

In this study, we integrate descriptor generation, feature selection, descriptor validation, dimensionality reduction and molecular clustering approaches to investigate the natural product chemical space, derived from the COCONUT database (Chandrasekhar et al., 2025; Sorokina et al., 2021). The primary objective is to construct an interpretable and stable cluster organization of natural products using selected descriptors from various categories. Descriptor subsets were validated to ensure robustness, reduced redundancy and comprehensive chemical space coverage. This integrated computational workflow contributes as an improved characterization of the natural product chemical space and supports a systematic exploration of molecular diversity.

## 2. Materials and Methods

### 2.1 Data Selection and Curation

Data of natural product molecules was sourced from the COCONUT database (Chandrasekhar et al., 2025) in August 2025, from the official webpage. A total of 695,114 molecules were downloaded in both 2D and 3D SDF formats. The 2D structures were used to extract canonical SMILES strings and molecular identifiers, while the 3D structures were used for geometry-based analyses (extractor.py, **see Supplementary data - scripts/data_retrieval**). Molecules that failed structural parsing or sanitization were removed using file-level structural exclusion and the RDKit Chem.SanitizeMol() function. The 3D conformers obtained were geometry-optimized using UFF (Universal Force Field) (Rappe et al., 1992) in Open Babel (O’Boyle et al., 2011) with a maximum of 20,000 steps. UFF was preferred due to its computational efficiency in large-scale processing (energy_min_script.py, **see Supplementary data - scripts/energy_minimization**). Structures that failed optimization were discarded. Post sanitization and optimization-success filtering, a total of 694,534 molecules were obtained, which were separated into chunks of 50000 molecules each and used for subsequent analyses (chunk_creation.py, **see Supplementary data - scripts/data_retrieval**).

Scaffold diversity was assessed using Murcko scaffolds generated from the SMILES representations of all molecules in the dataset (data_preprocessing.py, scaffold_identifier.py; **see Supplementary data - scripts/scaffold_decomposition/analysis**). Scaffold frequency distributions were calculated to characterize the structural composition of the chemical space. Analyses included identification of the 20 most prevalent scaffolds and quantification of scaffolds occurring within different frequency ranges (1–10, 1–100, and 1–1000 occurrences) to evaluate the extent of scaffold redundancy and rarity (histogram_data.py, **see Supplementary data - scripts/scaffold_decomposition/analysis**). The resulting frequency distributions were visualized using bar plots generated in R (top_20_bemis_murcko_scaffolds.R, 10_to_1000.R, 1_to_1000_range.R, 1_to_10_range.R; **see Supplementary data - scripts/scaffold_decomposition/visualization**).

### 2.2 Preparation of the Primary Molecular Descriptor Set

Descriptors were selected from those commonly used in literature (Darlami & Sharma, 2024; A. Kumar & Zhang, 2018; S. Kumar et al., 2020; Lo et al., 2018; Neves et al., 2018; Nicholls et al., 2010; Pirie et al., 2024; Prasanna & Doerksen, 2009; Ramahi et al., 2024; Shearer et al., 2022; Singh et al., 2021; Yukawa & Naven, 2020; Zhou et al., 2024) (**Supplementary Table 1**). A total of 80 descriptor properties were selected, which were then categorized into 7 classes: physicochemical (18 descriptors), three-dimensional (14 descriptors), electrotopological state (3 descriptors), partial charge (3 descriptors), graph and topological state (17 descriptors), pharmacophoric (19 descriptors) and ring and scaffold (6 descriptors) **(Supplementary Table 2)**.

Physicochemical, pharmacophoric and ring descriptors (n=43) encode the global structural features of the molecule derived from a molecule’s 2D representation. Physicochemical descriptors (n=18) capture properties related to size, polarity and structural composition (Vallianatou et al., 2015), pharmacophoric descriptors (n=19) quantify features associated with the interaction of a molecule with a biological target which are responsible for drug effects (Giordano et al., 2022) and ring and scaffold descriptors (n=6) describe the characteristics of the ring systems present within a molecule (Shearer et al., 2022). These were calculated through modules present in RDKit.Chem (physicochemical_descriptors.py, pharmacophoric_descriptors.py, ring_and_scaffold_descriptors.py, **see Supplementary data - scripts/descriptor_generation**).

Three-dimensional properties and graph and topological state properties (n=31) characterize molecular structure in terms of spatial geometry and connectivity patterns. Three dimensional properties (n=14) focus on spatial arrangement and shape of the molecules in 3D space (Meyers et al., 2016), while graph and topological descriptors (n=17) represent the graphical structure of the molecule (Khan et al., 2024) and describe connectivity patterns in the molecular graph. 11 three-dimensional properties were calculated through functions in the RDKit.Chem package, while 3 properties (bounding box volume, conformer count and inertia ratio) were calculated through custom functions incorporating functions present in the RDKit.Chem package. The graph and topological descriptors were calculated using the RDKit.Chem package and the networkx (https://networkx.org/en/) and Mordred (Moriwaki et al., 2018) libraries (3d_descriptors.py, graph_and_topological_descriptors.py, **see Supplementary data - scripts/descriptor_generation**).

Electrotopological state descriptors (n=3) and partial charge descriptors (n=3) capture the electronic environment around the atom to identify important molecular fragments (Roy & Mitra, 2012) and describe electron sharing among atoms (Engler et al., 2019) respectively. Electrotopological state descriptors were generated using RDKit functions, while the partial charge descriptors were generated using qmdesc (Guan et al., 2021) library, which uses quantum mechanical models to estimate partial charges (estate_descriptors.py, partial_charge_descriptors.py, **see Supplementary data - scripts/descriptor_generation**).

### 2.3 Exploratory Correlation Regime Analysis and Descriptor Subset Generation

A correlation matrix was computed to assess pairwise correlations among the 80 descriptors, using the Spearman correlation coefficient, which was selected due to its ability to study both monotonic and linear relationships (Zar, 2014) (correl_matrix_code.py, complete_correlation_matrix.R, **see Supplementary data - scripts/complete_correlation_matrix_generation**). Four correlation bands (0.5 - 0.85, 0.35 - 0.85, 0.25 - 0.85, and 0.35 - 0.75) were used to evaluate redundancy patterns across different regimes, without assuming any band to be optimal prior to analysis. For each band, descriptor pairs with pairwise correlation within the band’s limits were selected and individual descriptors involved were extracted (pairwise_correlation_matrix.py, **see Supplementary data - scripts/complete_correlation_matrix_generation**; band1.py, band2.py, band3.py, band4.py, **see Supplementary data - scripts/flat_file_generation_correlation**).

The extracted descriptors were used to obtain descriptor subsets using the greedy maximum coverage (Hanchuan Peng et al., 2005) and hierarchical clustering techniques (Liu et al., 2011). Both techniques were used with each correlation band to generate subsets of 10, 15, 20 and 25 descriptors each. For hierarchical clustering, subsets were selected based on either maximum or minimum average intra-cluster correlation. The individual descriptor sets were visualized using correlation matrix heatmaps (see **Supplementary data - scripts/greedy_max_coverage_scripts, scripts/hierarchical_clustering_scripts**).

To enable a detailed assessment of the effects of selection strategy, descriptor count, and correlation threshold range on descriptor selection, descriptor sets were generated for all parameter combinations (data_preparation.py, **see Supplementary data - scripts/method_analysis/complete_count**). Descriptor-set overlap was quantified for each variable while keeping the remaining two variables constant, and Jaccard similarity heatmaps were constructed to evaluate descriptor-selection consistency and parameter-specific preferences (comparison_script.py, data_split.py, method_comparison_jaccard, **see Supplementary data - scripts/overlap_computation**). Descriptor and descriptor-type enrichment analyses were performed by quantifying descriptor occurrence globally across all combinations, as well as within individual selection strategies, descriptor counts, and correlation threshold ranges, thereby identifying preferentially selected descriptors and descriptor categories (method_analysis.py, descriptor_presence, desc_presence_plots; desc_type_presence.py, descriptor_type_presence.R; **see Supplementary data - scripts/method_analysis/desc_type_presence, scripts/method_analysis/descriptor_occurrence**). Finally, the descriptor correlation structure was examined by calculating the Maximum Absolute Correlation (MAC) for each combination, with correlation heatmaps used to compare the correlation profiles associated with different parameter settings (correlation_analysis.py, mean_correl_analysis_comparisons.R; **see Supplementary data - scripts/method_analysis/correlation_analysis**).

### 2.4 Pareto Optimization for Descriptor Subset Generation

A multi-objective Pareto optimization framework (Xing et al., 2025) was employed for the selection of optimal correlation band and subset size (n=10, 15, 20, 25) and was applied across all candidate subsets. Each subset was evaluated using two objectives: (i) retained total variance, which is calculated as the trace of the covariance matrix of standardized descriptors normalized by covariance matrix of the full descriptor set (Jolliffe, 2002), and (ii) redundancy, which is calculated as the mean absolute pairwise Spearman correlation among descriptors within the subsets (Tosco & Balle, 2011) (pareto_analysis.py, **see Supplementary data - scripts/pareto_optimization**). This analysis facilitated the selection of Pareto-optimal (non-dominated) solutions that provided the most favorable compromise between descriptor informativeness and redundancy reduction. The final subset size was determined based on the Pareto front structure and identification of a knee point that corresponds to diminishing marginal gains in variance retention relative to increasing redundancy (mean_abs_correl.R, variance_retention_vs_redundancy.R; see **Supplementary data - scripts/pareto_optimization**) (Verhellen, 2022).

Based on the Pareto optimization, all subsequent analyses were performed using the selected correlation threshold range, greedy maximum-coverage selection, and a fixed subset size (n=20 descriptors).

### 2.5 Information Retention and Stability Analysis

Principal Component Analysis (PCA) (Jolliffe, 2002) was used to confirm that the final subset preserves dominant variance structure. PCA was independently applied to the complete dataset (n = 694,534 molecules) using the full set of 80 standardized descriptors and to the reduced subset of 20 descriptors (Giuliani, 2017) (20desc_pca.py, 80desc_pca.py; **see Supplementary data - scripts/pca**). The cumulative explained variance across top 20 PCs were compared to assess the structural firmness of the reduced representation relative to the original descriptor space (cumulative_explained_variance_plot.R; **see Supplementary data - scripts/pca**).

To quantify descriptor stability and overlap, perturbation analysis was conducted (Kalousis et al., 2007). To assess the robustness of descriptor selection, the complete dataset was randomly partitioned into 60:40 training and validation subsets using five random seeds (0, 1, 42, 101, and 123) (Hanchuan Peng et al., 2005). For each split, descriptor matrices were generated using the specified correlation threshold range, followed by greedy maximum coverage feature selection to obtain a final set of 25 descriptors (split_gen.py, flat_file_gen.py, greedy_max_cov.py; **see Supplementary data - scripts/perturbation_analysis.py**). Correlation matrices of the selected descriptors were visualized as heatmaps to evaluate redundancy patterns (visualization_of_matrices.R, **see Supplementary data - scripts/perturbation_analysis.py**). The consistency of descriptor selection across random seeds was assessed by quantifying the overlap between descriptors selected from the 60% and 40% subsets, as well as with the final 25-descriptor set, and the results were visualized using bar plots (perturbation_analysis_presence.py, descriptor_stability_plot.R; **see Supplementary data - scripts/perturbation_analysis**).

### 2.6 Evaluation of Three Clustering Techniques on a Representative Dataset Subset

To identify the optimal clustering strategy, 50 000 molecules were sampled from the complete dataset (sampling.py, **see Supplementary data - scripts/clustering_tries**). Three clustering algorithms were chosen to represent different types of clustering algorithms: Bisecting KMeans (Steinbach et al., 2000), KMeans (Jin & Han, 2011) (kmeans_plusplus.py, kmeans_random.py; **see Supplementary data - scripts/clustering_tries/kmeans**) and HDBSCAN (Hierarchical Density Based Spatial Clustering of Applications with Noise) (Campello et al., 2013), which represent connectivity models, centroid models, and density-based clustering models respectively (Wani, 2024). UMAP (Uniform Manifold Approximation and Projection) (McInnes et al., 2018) was chosen as the dimensionality reduction technique for the HDBSCAN and Bisecting KMeans due to its ability to preserve local and global structure with large-scale and heterogeneous data (umap_hdbscan.py, umap_bkmeans.py; **see Supplementary data - scripts/clustering_tries**). The data was transformed and standardized using the Yeo-Johnson transformer (Yeo, 2000) and StandardScaler (Pedregosa et al., 2012) respectively, to reduce skew. Clustering performance was evaluated using Silhouette score (Rousseeuw, 1987), Davies Bouldin Index (Davies & Bouldin, 1979), Calinski Harabasz Index (Caliński & Harabasz, 1974), cluster purity scores, noise fraction and Dunn index (Dunn, 1973). Each clustering model was trained using multiple hyperparameter combinations (**Supplementary Table 3**). All analyses were carried out using CuML (Raschka et al., 2020) in a Google Colab workspace (Bisong, 2019).

The KMeans technique was evaluated by analysing variations of Silhouette score, Calinski-Harabasz index, inertia and Davies-Bouldin score across different numbers of clusters (k) (kmeans_result_analysis.R; **see Supplementary data - scripts/clustering_try_result_analysis**). The UMAP-HDBSCAN algorithm was assessed using multiple hyperparameter combinations, including analysis of distributions of Silhouette scores across varying UMAP n_neighbours, relationships between cluster purity and noise fraction, and changes in Silhouette score with cluster count (hdbscan_cluster_purity_vs_noise_frac.R, hdbscan_sil_scores_vs_no_of_clusters.R, hdbscan_metric_heatmaps_vs_hyperparams.R, umap_sil_score_vs_neighbours_boxplot.R; **see Supplementary data - scripts/clustering_try_result_analysis**). The UMAP-Bisecting KMeans approach was tested by comparing Silhouette score, Davies-Bouldin index, and Calinski-Harabasz index across trials, along with analyzing the effects of different UMAP and Bisecting KMeans combinations on Silhouette scores and Davies Bouldin indices (bkmeans_metrics_vs_trial.R, bkmeans_silhouette_vs_bkmeans_params.R, bkmeans_silhouette_vs_umap_hyperparams.R; **see Supplementary data - scripts/clustering_try_result_anlaysis**).

### 2.7 Full Scale Clustering of the Dataset using UMAP-HDBSCAN Method

The complete dataset was further clustered using the UMAP-HDBSCAN algorithm, selected due to its ability to preserve local manifold structure and effectively identify clusters in noisy and unevenly distributed datasets. This approach was suited for the current dataset due to its substantial noise and uneven density distributions (umap.py, **see Supplementary data - scripts/complete_umap_hdbscan_clustering**). Prior to complete clustering, the dataset was transformed using the Yeo-Johnson (Power) transform, and scaled using StandardScaler. The manifold was further saved as a .npy file, for reproducibility and subsequent use and prediction data generation was enabled to allow new compounds to be assigned to clusters without requiring the retraining of the entire model **(See Table 1 for complete hyperparameter set)**.

**Table 1:** Final UMAP-HDBSCAN Hyperparameters.

| Hyperparameter | Values |
| --- | --- |
| UMAP |  |
| n_neighbours | 30 |
| min_dist | 0.3 |
| n_components | 12 |
| metric | euclidean |
| low_memory | True |
| dens_map | False |
| n_epochs | 150 |
| verbose | True |
| random_state | 42 |
| HDBSCAN |  |
| min_cluster_size | 100 |
| min_samples | 10 |
| cluster_selection_method | leaf |
| approx_min_span_tree | True |
| metric | euclidean |
| alpha | 0.7 |
| prediction_data | True |
| core_dist_n_jobs | -1 |

The clustering was evaluated using three metrics: Silhouette score, Davies Bouldin Index and Calinski-Harabasz (CH) index. Due to computational constraints, the Silhouette score was calculated on a sample size of 50,000 molecules with a fixed random seed to ensure reproducibility. Molecules labelled as noise by the HDBSCAN algorithm were subsequently reassigned to clusters based on their distance to cluster medoids. Clusters were evaluated both before and after reassignment of noise molecules (hdbscan_noise_assignment.py, **see Supplementary data - scripts/complete_umap_hdbscan_clustering**).

Post-clustering analysis was conducted to study the structural organization of the UMAP-HDBSCAN clusters. Cluster assignments, UMAP embeddings, HDBSCAN membership probabilities and cluster size distributions were extracted for both core and post-noise reassigned clusters. Cluster centroids and medoids were computed within the UMAP manifold space, and molecule membership lists were generated for each cluster before and after noise reassignment (hdbscan_noise_assignment.py, **see Supplementary data - scripts/complete_umap_hdbscan_clustering**). These outputs facilitated downstream analysis of cluster membership and structural diversity.

Representative molecules were selected for each cluster using a centroid proximity and structural diversity criteria. For every cluster, the medoid, the molecule closest to the centroid excluding the medoid, and five structurally diverse molecules were identified, using pairwise Euclidean distances within the UMAP embedding space. The diverse representatives are constrained within the 80th percentile distance radius from the cluster medoid to avoid selection of extreme outlier compounds (hdbscan_noise_assignment.py, **see Supplementary data - scripts/complete_umap_hdbscan_clustering**).

### 2.8 Structural and Chemical Characterization of Identified Clusters

The clusters identified by UMAP-HDBSCAN and noise reassignment were characterized by cluster stability, chemical composition, compactness and diversity. These analyses comprised a combination of density-based statistics, scaffold enrichment analyses, physicochemical descriptor distributions, manifold geometry analyses and molecular similarity metrics.

Cluster stability was assessed using core-assignment statistics derived from HDBSCAN outputs. These include core cluster size, post-noise reassigned cluster size, HDBSCAN membership probability distributions and probability inter-quartile range (IQR). Core fraction was calculated as the ratio of core cluster membership to total post-reassignment cluster membership, as a metric for density stability within a cluster. Clusters were categorized as accepted, borderline and rejected based on core fraction and IQR of membership probability. Clusters with a core fraction > 0.5 were considered “accepted”, while those with core fraction < 0.3 and probability IQR < 0.4 were rejected due to structural instability. The remaining clusters were classified as “borderline”. Thresholds were selected empirically based on the observed distributions of core fraction and membership probability IQR across clusters (core_frac_noise.py, **see Supplementary data - scripts/cluster_characterization**).

The chemical composition of clusters was analyzed using scaffold enrichment analysis and physicochemical descriptor distribution. The dominant scaffold fraction and cluster occupancy fraction were calculated to assess structural enrichment within clusters. For each cluster and its representative molecule set, descriptive statistics including mean, median, standard deviation and interquartile range were calculated for molecular weight, LogP, Total Polar Surface Area (TPSA), hydrogen bond donor count (HBD), hydrogen bond acceptor count (HBA), and number of rotatable bonds. Representative-level profiling was further combined with scaffold profiling for studying local diversity within clusters (chemical_profiling.py, **see Supplementary data - scripts/cluster_characterization**).

Cluster geometry and spatial compactness were studied using the UMAP embedding space. Cluster centrality was evaluated using the distance between centroid and medoid. Intra-cluster spread statistics were computed separately for core molecule subsets and post-noise reassigned clusters, using Euclidean distance distributions. The determinant of the covariance matrix was calculated to evaluate cluster volume and spatial dispersion, while cluster boundary stability was estimated using the density gradient (compactness_and_geometry.py, **see Supplementary data - scripts/cluster_characterization**).

Jaccard-similarity based fingerprints were used to study structural diversity within and between clusters. Within-cluster similarity distribution was calculated as the average pairwise Jaccard similarity between molecules present in individual clusters whereas within-cluster similarity analysis was done by pairwise comparison of cluster medoids. A separation ratio, defined as the ratio of intra-cluster similarity to inter-cluster similarity, was calculated to quantify cluster overlap within the chemical space (diversity_analytics.py, **see Supplementary data - scripts/cluster_characterization**).

Cluster assignment confidence was analyzed using HDBSCAN membership probabilities. The mean, variance and interquartile range of membership probabilities were computed for each cluster, to evaluate assignment confidence and dispersion of membership probabilities across the clustering hierarchy. All data pertaining to cluster characterization is provided in Supplementary data (mean_variance_prob.py, **see Supplementary data - scripts/cluster_characterization**).

Distributional analyses of cluster-level metrics, including cluster size, core fraction, density gradient, separation ratio, scaffold enrichment, and HDBSCAN membership probability statistics, were performed using histogram-based visualizations (histograms.R, **see Supplementary data - scripts/cluster_characterization/visualization**). Pairwise relationships between cluster evaluation metrics were examined through scatterplot-based analyses. The fidelity of representative molecules was assessed by comparing cluster-level descriptor statistics with medoid descriptor values using scatterplots (scatterplot_single_file.R, scatterplot_two_files.R; **see Supplementary data - scripts/cluster_characterization/visualization**). Bland-Altman analyses were additionally conducted to evaluate the agreement between cluster-level descriptor means and corresponding medoid descriptor values, thereby assessing the extent to which medoids accurately represented their respective clusters; the results were visualized using scatterplots (bland_altman_plots.R, **see Supplementary data - scripts/cluster_characterization/visualization**). Summary tables were generated to complement the graphical analyses (rep_difference_analysis.py, bivariate_rep_analysis.py, bland_altman_analysis.py; **see Supplementary data - scripts/cluster_characterization**). All visualizations were produced using the ggplot2 package in R, and statistical summaries were produced using the pandas package in Python.

### 2.9 Representative Suitability Analysis

Clusters were first stratified into small, medium, and large size categories. From each category, six clusters were initially selected, and within each, 25 random molecules were sampled along with the seven predefined representatives to compare representative performance against intra-cluster diversity (stratification.py; **see Supplementary data - scripts/complete_umap_hdbscan_clustering**). Based on molecule availability, five clusters per size stratum were retained for downstream analysis. Docking was then performed to evaluate representative stability for drug discovery applications. The KEAP1 structure (PDB ID: 8XGK) (Berman, 2000; Otake et al., 2024), targeting the Kelch domain active site, was retrieved from the Protein Data Bank and used as the receptor. Docking simulations were carried out using PyRx (Dallakyan & Olson, 2015), with grid parameters set to fully cover the active site to ensure comprehensive ligand sampling.

The suitability of cluster representatives was evaluated by comparing their docking performance against randomly selected molecules from the corresponding clusters. Analyses included the distribution of representative binding affinity scores (docking_score_distribution.R, global_distribution_docking_scores_random.R; **see Supplementary data - scripts/docking_representative_analysis/visualization**), identification of the best-performing representatives through both global and cluster size-stratified assessments (best_rep_analysis.py,, best_size_strata_rep.R, global_best_cluster_performance.R; **see Supplementary data - scripts/docking_representative_analysis**), and characterization of the binding affinity distributions of randomly selected molecules at global and size-stratified levels (). In addition, average binding affinities of cluster representatives and 25 randomly selected molecules per cluster were compared using line-plot visualizations, performed both globally and across cluster size categories (stratified_docking_affinity.R, global_rep_analysis.R, **see Supplementary data - scripts/docking_representative_analysis/visualization**). All analyses were conducted in Python and visualizations were conducted in R.

## 3. Results

### 3.1. Dataset Composition and Structural Landscape

#### 3.1.1. Structural Diversity

The study dataset was derived from molecules retrieved from the COCONUT database. Molecular characterization was performed using the 80 descriptors described in the Methods section, and structural diversity was further examined through Murcko scaffold decomposition. (Bemis & Murcko, 1996).

Following decomposition, 124447 unique scaffolds were detected among 694,534 molecules, corresponding to an average of ∼6 molecules per scaffold (**Supplementary Table 4**). The scaffold distribution is strongly right-skewed. The benzene ring was the most detected scaffold (**Figure 1a, Supplementary Table 5**, n=15322), corresponding to ∼2% of the dataset, reflecting a high prevalence of aromatic structures. Beyond this scaffold, frequencies sharply decrease, with the next common scaffolds occurring between 1146 - 4501 molecules **(Figure 1a, Supplementary Table 4, 5)**. The top 20 scaffolds cumulatively account for 7.6% of the whole dataset (**Figure 1a, b; Supplementary Table 4, 5**).

**Figure 1:**
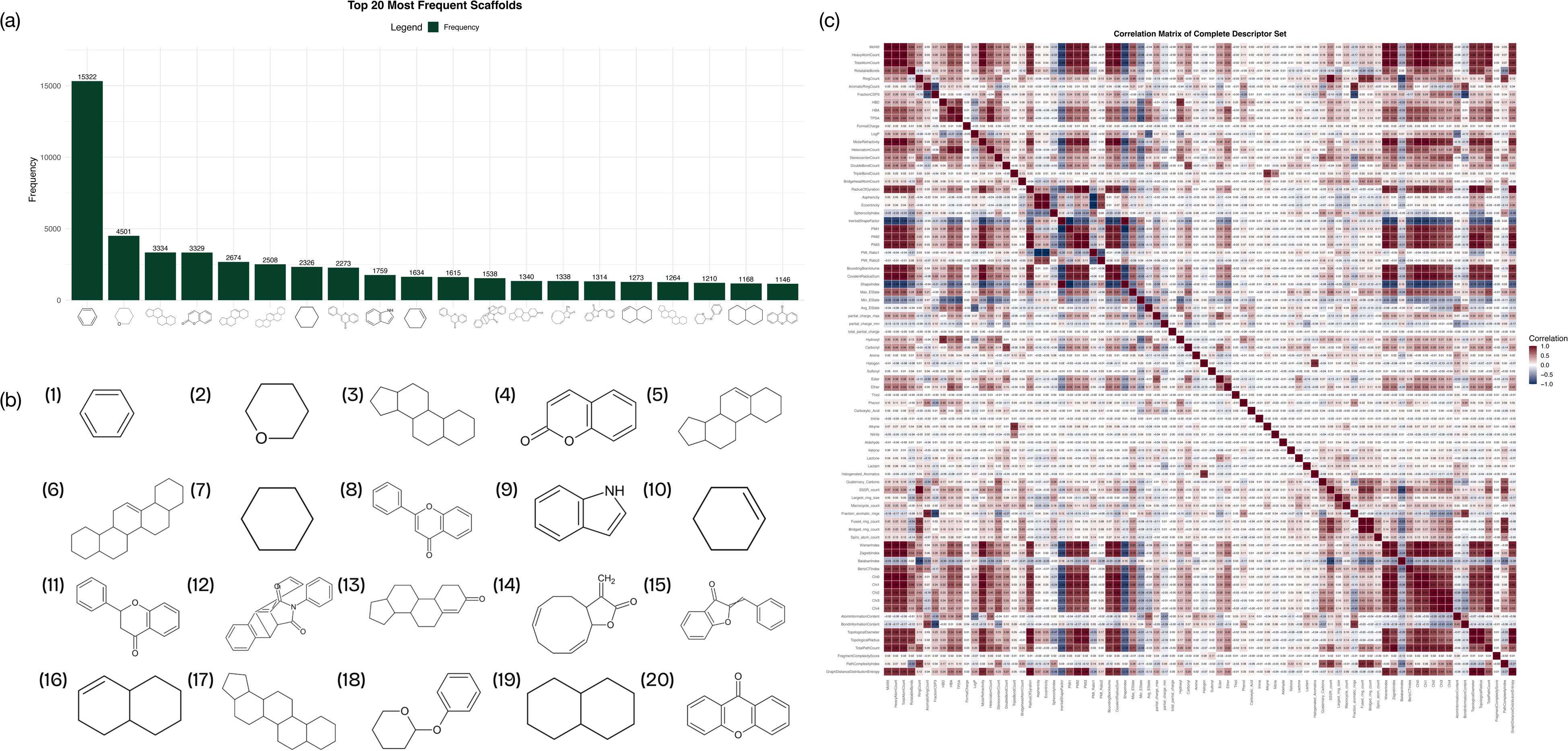
Scaffold and Descriptor Correlation Landscape of the COCONUT Database. **(a) Frequency of the Top 20 Scaffolds in the COCONUT Database:** The bar plot summarizes the frequencies of the 20 most abundant scaffolds identified in the dataset. The x-axis displays the scaffold structures, while the y-axis indicates the number of compounds containing each scaffold. Numerical annotations above the bars represent the corresponding scaffold frequencies. **(b) Top 20 Scaffold Structures:** Representative chemical structures of the 20 most frequently occurring scaffolds in the complete COCONUT dataset. **(c) Complete Descriptor Correlation Landscape:** The heat map illustrates the pairwise Spearman correlation coefficients among the 80 molecular descriptors. Both axes represent the descriptor set, and each cell corresponds to the correlation between a descriptor pair, with coefficient values annotated within the matrix. The color scale ranges from dark blue (−1; strong negative correlation) to dark red (+1; strong positive correlation), with white indicating no correlation (0).

Scaffold diversity varies substantially across the dataset. 17.91% of scaffolds occur in fewer than 1,000 molecules **(Supplementary Figure 1a, Supplementary Table 6a)**. At lower frequency thresholds, 17.84% of scaffolds are observed in fewer than 100 molecules **(Supplementary Figure 1b, Supplementary Table 6b)**, 16.74% scaffolds occur in fewer than 10 molecules, and 8.85% are singletons **(Supplementary Figure 2, Supplementary Table 6c)**. This distribution demonstrates a heavily skewed scaffold landscape characterized by a limited number of recurrent frameworks and a substantial proportion of sparsely represented structures.

#### 3.1.2. Descriptor Redundancy in the Full Space

A pairwise Spearman coefficient correlation matrix was computed using the 80 descriptors. Correlation coefficients ranged from −2 21 × 10^−5^ (minimum) to 1 (maximum), indicating that both strong monotonic as well as weak inverse relationships are present within the correlation space (**Figure 1c, Supplementary Table 7)**. The distribution is dominated by low-magnitude negative correlations ranging between -0.3 to 0 (∼40%), indicating that most descriptor pairs exhibit weak inverse associations. Overall, 60.73% (n=1919) of correlations are positive and 39.27% (n=1241) negative. This indicates that weak inverse correlations are more frequent compared to stronger monotonic relationships. Specifically, 7.44% (n=235) of absolute correlations are above 0.8, among which 6.45% (n=204) are positive and 0.98% (n=31) are negative. These high magnitude associations represent a relatively small yet structurally meaningful component of the dataset (**Supplementary Figure 3, Supplementary Table 8**).

Despite the dominance of weak inverse correlations, localized clusters of strong positive correlation are observed among PMI1, PMI2, PMI3, Chi0, Chi1, Chi2, Chi3, Chi4, topological diameter, topological radius, bounding box volume, covalent radius sum, molecular weight, heavy atom count, and total atom count, with extensive correlation observed in the Chi0-4 descriptor space (**Figure 1c, Supplementary Table 7)**. These patches reflect shared physicochemical and structural information encoded by related descriptor groups which cause localized redundancies within the feature space, with global heterogeneity and local redundancy. This structural pattern justifies the need for systematic descriptor reduction strategies aimed at minimizing localized redundancy without disrupting the broader variance structure of the chemical descriptor space.

### 3.2. Descriptor Subset Generation and Algorithmic Performance Evaluation

The greedy maximum coverage, minimum average correlation hierarchical clustering, and maximum average correlation hierarchical clustering approaches were evaluated across four descriptor counts and four correlation threshold bands using flat files **(Supplementary Table 9)**. Correlation matrix heatmaps were generated for each parameter combination to visualize descriptor relationships and redundancy patterns (**Supplementary Figures 4-27; Supplementary Tables 10-21**). For the hierarchical clustering approaches, dendrograms were additionally constructed to illustrate descriptor grouping structures and facilitate comparison of clustering behavior across descriptor counts and correlation threshold bands **(Supplementary Figures 28-31; Supplementary Table 22)**.

#### 3.2.1. Comparison of Descriptor Prevalence across Descriptor Selection Strategies

Across the 10-descriptor configurations, no descriptors were consistently conserved across all three selection strategies within any of the four bands **(Supplementary Tables 23a, b, c; Supplementary Figures 4, 5, 12, 13, 20, 21)**. Pairwise comparisons showed minimal overlap between the greedy maximum coverage and hierarchical minimum average correlation approaches, whereas the two hierarchical clustering techniques consistently exhibited the highest descriptor overlap. Similar trends were observed for the 15-descriptor configurations, where only a single common descriptor was identified across all three methods in bands 3 and 4, represented by phenol and covalent radius sum, respectively (**Supplementary Table 24; Supplementary Figures 6, 7, 14, 15, 22, 23**). In the 20-descriptor configurations, two descriptors were consistently retained across all three methods in bands 1-3, with asphericity emerging as a recurrent conserved descriptor, while band 4 (0.35-0.75) showed comparatively lower conservation **(Supplementary Table 25; Supplementary Figures 8, 9, 16, 17, 24, 25)**. For the 25-descriptor configurations, 3-5 descriptors were conserved across methods, with descriptor conservation progressively increasing from band 4 to band 1 **(Supplementary Table 26; Supplementary Figures 10, 11, 18, 19, 26, 27)**. Across all descriptor-count configurations, the greedy maximum coverage technique consistently exhibited the highest number of uniquely retained descriptors, whereas the hierarchical maximum average correlation approach generally showed the lowest descriptor uniqueness **(Supplementary Tables 23a, 24a, 25a, 26a; Supplementary Figures 4-11)**.

Across all four correlation bands, the number of descriptors conserved across all three selection strategies increased progressively with increasing descriptor count, ranging from no common descriptors in the 10-descriptor configurations to 4-5 common descriptors in the 25-descriptor configurations **(Supplementary Tables 27-29; Supplementary Figures 4-27)**. In all bands, the greedy maximum coverage approach consistently exhibited the highest number of uniquely retained descriptors **(Supplementary Table 27, Supplementary Figures 4-11)**, whereas the hierarchical maximum average correlation approach generally showed the lowest descriptor uniqueness **(Supplementary Table 28, Supplementary Figures 12-19)**, occasionally shared with the hierarchical minimum average correlation technique in the 25-descriptor configurations **(Supplementary Table 29, Supplementary Figures 20-27)**. Pairwise comparisons further revealed consistent overlap patterns across all bands, where the two hierarchical clustering approaches demonstrated the highest descriptor overlap **(Supplementary Tables 28-29, Supplementary Figures 12-27)**, while comparisons between greedy maximum coverage and hierarchical minimum average correlation consistently exhibited the lowest overlap **(Supplementary Tables 27, 29; Supplementary Figures 4-11, 20-27)**, occasionally shared with the greedy maximum coverage and hierarchical maximum average correlation comparison in higher descriptor-count configurations **(Supplementary Figures 27-28, Supplementary Figures 4-19)**.

#### 3.2.2. Comparison of Descriptor Prevalence across Correlation Threshold Ranges

In the greedy maximum coverage technique, the BertzCT index was consistently retained across all four correlation threshold ranges for the 10-, 15-, 20-, and 25-descriptor configurations, indicating stable descriptor conservation across varying redundancy thresholds **(Supplementary Table 27; Supplementary Figures 4-11)**. Band 4 (0.35-0.75) consistently exhibited the highest number of unique (non-overlapping) descriptors, predominantly comprising Chi-based topological descriptors, suggesting increased descriptor diversification under stricter filtering conditions **(Supplementary Table 27d; Supplementary Figures 5b, 7b, 9b, 11b)**. The 25-descriptor configuration had the highest degree of descriptor overlap across bands, with electrotopological state descriptors, heteroatom count, and BertzCT index frequently retained, whereas lower descriptor counts exhibited comparatively reduced overlap **(Supplementary Table 26; Supplementary Figures 10, 11, 18, 19, 26, 27)**. Across all descriptor counts, the highest multi-band similarity was consistently observed between bands 1, 2, and 3, where descriptors such as PMI2, graph distance distribution entropy, and average electrotopological state were repeatedly conserved **(Supplementary Table 27a, Supplementary Figures 4-11)**. Similarly, pairwise comparisons revealed that band 1-band 2 and band 2-band 3 combinations exhibited the greatest descriptor overlap, indicating relatively stable descriptor retention within intermediate correlation threshold ranges **(Supplementary Table 23, Supplementary Figures 4-5, 12-13, 20-21)**.

For the hierarchical clustering approach using maximum average correlation, PMI Ratio 2 was consistently retained across all four correlation threshold ranges for the 10- and 15-descriptor configurations, while both PMI_Ratio2 and BertzCT index were conserved across all bands in the 20- and 25-descriptor configurations. Among three-band overlaps, the band 1-band 2-band 3 combination consistently exhibited the highest descriptor conservation across all descriptor counts, with the greatest overlap observed for the 20- and 25-descriptor configurations. In contrast, the lowest three-band overlap was generally observed for the band 1-band 3-band 4 combination, where only PMI Ratio 2, along with occasional retention of BertzCT index or Largest ring size in higher descriptor counts, remained conserved. Pairwise comparisons further showed that band 2-band 3 and band 1-band 2 combinations exhibited the highest descriptor overlap across multiple descriptor-count configurations, whereas comparisons involving band 3-band 4 consistently demonstrated the lowest overlap. Analysis of uniquely retained descriptors revealed that band 4 exhibited the highest number of unique descriptors for the 15-, 20-, and 25-descriptor configurations, while band 2 consistently showed the lowest number of unique descriptors across all descriptor counts **(Supplementary Table 28b, d; Supplementary Figures 12-19(b))**.

For the hierarchical clustering approach using minimum average correlation, PMI Ratio 1 was consistently retained across all four correlation threshold ranges for all descriptor-count configurations, while additional descriptors such as bounding box volume, HBD, molar refractivity, Balaban index, and bond information content were conserved in higher descriptor-count settings **(Supplementary Table 29; Supplementary Figures 20-27)**. Among three-band overlaps, the highest descriptor conservation was generally observed for the band 2-band 3-band 4 combination in the 10-, 15-, and 20-descriptor configurations, whereas the 25-descriptor configuration showed maximum overlap for the band 1–band 2–band 3 combination. In contrast, the band 1–band 3–band 4 combination consistently exhibited the lowest three-band overlap across multiple descriptor counts. Pairwise comparisons revealed that band 2–band 4 and band 2–band 3 combinations showed the greatest descriptor overlap across most descriptor-count configurations, while combinations involving band 1–band 3 or band 3–band 4 generally demonstrated lower overlap. Unlike the maximum average correlation approach, no consistent band-specific trend was observed for uniquely retained descriptors, with the highest number of unique descriptors distributed variably across bands 1, 3, and 4 depending on descriptor count **(Supplementary Tables 30-34; Supplementary Figures 20-27)**.

Across the 10-descriptor configurations, no consistent patterns were observed in dual-band overlaps, three-way correlations, or uniquely retained descriptors across the three selection strategies, although a single descriptor remained conserved across all four bands for each method, and both PMI ratios were consistently retained across all bands in the two hierarchical clustering approaches **(Supplementary Table 30; Supplementary Figures 4-5, 12-13, 20-21)**.

Similar trends were observed for the 15-descriptor configurations, where band 4 exhibited the highest number of unique descriptors in the greedy maximum coverage and hierarchical maximum average correlation approaches, while band 1 showed the highest uniqueness in the hierarchical minimum average correlation approach **(Supplementary Table 31; Supplementary Figures 6-7, 14-15, 22-23)**. Additionally, the band 2–band 3 combination demonstrated the highest pairwise overlap in the greedy maximum coverage and hierarchical maximum average correlation techniques, whereas the hierarchical minimum average correlation approach showed maximum overlap for band 2–band 4. In contrast, the 20- and 25-descriptor configurations exhibited clearer and more consistent overlap patterns across all methods, where the band 2–band 3 and band 1–band 2 combinations generally showed the highest descriptor overlap, while combinations involving band 3–band 4 or band 2–band 4 demonstrated lower overlap. Across all three techniques, band 4 consistently exhibited the highest number of uniquely retained descriptors, while the band 1–band 2–band 3 combination demonstrated the greatest three-way descriptor conservation in most higher descriptor-count configurations **(Supplementary Tables 23-26; Supplementary Figures 4-27)**.

#### 3.2.2. Comparison of Descriptor Prevalence across Descriptor Counts

In the greedy maximum coverage technique, 10 descriptors were consistently conserved across all four descriptor-count configurations in bands 2, 3, and 4, while band 1 exhibited comparatively lower conservation with only seven common descriptors across all counts. Among three-way descriptor-count overlaps, the highest descriptor conservation was observed for the 15–20–25 combination across bands 2, 3, and 4, whereas band 1 consistently showed lower overlap. Most three-way descriptor-count comparisons retained only the descriptors already conserved across all four descriptor counts, indicating stable core descriptor retention across configurations. No unique descriptors were observed for the 10- and 15-descriptor configurations across any band, while the 25-descriptor configuration consistently exhibited the highest number of unique descriptors in all bands. Pairwise comparisons further revealed that the 20–25 descriptor combination demonstrated the greatest descriptor overlap across all bands, whereas the 10–15 combination consistently showed the lowest overlap **(Supplementary Table 27, Supplementary Figures 4-11)**.

For the hierarchical clustering approaches, 5–8 descriptors in the maximum average correlation technique and 7–10 descriptors in the minimum average correlation technique were consistently conserved across all four bands, indicating relatively stable descriptor retention across varying descriptor-count configurations. In both approaches, the 15–20–25 descriptor combination consistently exhibited the highest three-way descriptor conservation across all bands, while combinations involving 10–15–25 and 10–20–25 generally demonstrated comparatively lower overlap and frequently lacked uniquely conserved descriptors beyond those already retained across all four descriptor counts. Pairwise comparisons further revealed consistent overlap patterns across both techniques, where the 20–25 descriptor combination demonstrated the greatest descriptor overlap across most bands, whereas the 10–25 combination generally exhibited the lowest overlap. Additionally, the 25-descriptor configuration consistently showed the highest number of uniquely retained descriptors across all bands in both approaches, while lower descriptor counts, particularly the 15-descriptor configuration, frequently exhibited few or no uniquely retained descriptors across multiple bands **(Supplementary Tables 28-29; Supplementary Figures 12-27)**. Although the overall overlap trends were broadly similar between the two hierarchical clustering approaches, the minimum average correlation technique demonstrated comparatively greater descriptor conservation across bands relative to the maximum average correlation approach.

Across all four correlation bands, consistent descriptor conservation patterns were observed across the three selection strategies. In band 1, approximately seven descriptors were conserved across all four descriptor-count configurations for all methods, whereas bands 2 and 3 exhibited comparatively higher conservation with 7–10 common descriptors, and band 4 showed 5–10 conserved descriptors. Among three-way descriptor-count overlaps, the 15–20–25 combination consistently demonstrated the highest descriptor conservation across all methods in all four bands, while combinations involving 10–15–25 and 10–20–25 generally exhibited lower overlap. Pairwise comparisons further revealed that the 20–25 descriptor combination consistently showed the greatest descriptor overlap in bands 2, 3, and 4 across all methods, whereas no consistent pattern was observed for the highest-overlap combinations in band 1. In contrast, the 10–25 descriptor combination generally exhibited the lowest descriptor overlap. Analysis of uniquely retained descriptors showed that the 25-descriptor configuration consistently exhibited the highest number of unique descriptors across all bands, while the 15-descriptor configuration frequently showed no unique descriptors, particularly in bands 2 and 4. Additionally, several three-way descriptor-count combinations, including 10–15–25 and 10–20–25, exhibited no uniquely conserved descriptors across all methods **(Supplementary Tables 27-29; Supplementary Figures 4-27)**, indicating stable retention of a core descriptor subset across multiple parameter configurations.

#### 3.2.4. Descriptor and Descriptor Type Presence across Selection Strategies, Descriptor Counts and Correlation Threshold Ranges

Out of the 80 descriptors evaluated across 48 combinations, 73 appeared at least once, with the BertzCT index being the most frequently occurring descriptor, present in 62.5% of combinations, followed by PMI ratio 2 at 52.1% and PMI ratio 1 at 45.8%. In contrast, heavy atom count, total atom count, PMI3, and total path count each appeared only once. All Chi descriptors were represented, with frequencies ranging from 8.3% for Chi1 to 27.1% for Chi4 (**Supplementary Figure 32, Supplementary Table 34)**. Across descriptor selection strategies, descriptor occurrence ranged from 6.2% to 100%, although no descriptor was universally retained across all strategies. BertzCT index consistently showed strong representation in both greedy maximum coverage and hierarchical clustering based on maximum average correlation approaches **(Supplementary Figure 33, Supplementary Table 35)**. Across descriptor counts, occurrence frequencies ranged from 8.3% to 100%, with BertzCT index remaining the most consistently selected descriptor. Increasing descriptor count to 20 and 25 resulted in higher representation of PMI ratio 1, PMI ratio 2, and asphericity among the top five descriptors, whereas triple bond count was more prominent in the 10 and 15 descriptor subsets. Chi descriptors showed relatively low occurrence in smaller descriptor sets but increased in frequency with larger descriptor counts **(Supplementary Figure 34, Supplementary Table 36)**. Across correlation threshold bands, BertzCT index remained among the top five descriptors in all bands except the strictest threshold band, while Chi descriptors were retained across all four bands. Consistent trends of average EState occurring more frequently than maximum EState, which in turn occurred more frequently than minimum EState, and PMI ratio 2 occurring more frequently than PMI ratio 1, were also observed across all threshold bands **(Supplementary Figure 35, Supplementary Table 37)**.

Across combinations of descriptor selection method, descriptor count, and correlation threshold range, normalized descriptor-type presence revealed clear enrichment patterns. Physicochemical descriptors showed the highest overall presence at 10.82, whereas partial charge descriptors showed the lowest presence at 6.33. Pharmacophoric and graph and topological descriptors also demonstrated consistently high enrichment, closely followed by 3D descriptors, indicating that functional group composition, physicochemical properties, and molecular topology contribute strongly to representation of the molecular landscape **(Supplementary Figure 36, Supplementary Table 38a)**. Descriptor selection strategies displayed distinct preferences, with hierarchical clustering approaches favoring pharmacophoric descriptors, while greedy maximum coverage showed relatively balanced enrichment across physicochemical, 3D, and graph and topological descriptors. The hierarchical clustering approach based on minimum average correlation showed no enrichment of electrotopological state descriptors, whereas partial charge descriptors consistently showed low enrichment across all selection strategies **(Supplementary Figure 37, Supplementary Tables 38b-d)**. Correlation threshold ranges also influenced descriptor-type representation, with wider correlation bands favoring pharmacophoric and 3D descriptors, while narrower correlation ranges showed stronger enrichment of physicochemical, graph and topological, and 3D descriptors. Across all four correlation threshold bands, partial charge descriptors remained minimally represented **(Supplementary Figure 38, Supplementary Tables 38e-h)**. Across descriptor counts, pharmacophoric descriptors consistently showed moderate to high enrichment, whereas EState and partial charge descriptors maintained systematically low presence. Graph and topological descriptors and ring descriptors showed moderate enrichment across all descriptor counts, while physicochemical descriptors demonstrated high enrichment in all cases except the 10 descriptor subsets **(Supplementary Figure 39, Supplementary Tables 38i-l)**.

#### 3.2.5. Mean Absolute Correlation across Selection Strategies, Descriptor Counts and Correlation Threshold Ranges

Average pairwise descriptor correlations differed more strongly between selection methods and correlation bands than between descriptor counts. Across all bands, the greedy maximum coverage approach generally produced higher average correlations than either hierarchical clustering method, while hierarchical minimum-average and maximum-average clustering exhibited highly similar correlation profiles with only minor differences. Increasing the descriptor count from 10 to 25 descriptors resulted in modest increases in correlation for both hierarchical approaches, whereas the greedy maximum coverage method showed relatively stable correlations in Bands 1–3 and a progressive decrease in Band 4. Distinct trends were also observed across correlation bands, with Band 4 consistently yielding the highest average correlations and Band 3 the lowest. The strongest contrast between methods occurred in Band 4, where greedy maximum coverage produced substantially higher correlations (0.466–0.636) than hierarchical maximum-average (0.107–0.209) and hierarchical minimum-average clustering (0.131–0.232). In contrast, Bands 1 and 2 exhibited intermediate correlation values, while Band 3 remained characterized by comparatively low correlations across all methods. Overall, these results indicate that the choice of descriptor selection method exerted a greater influence on the resulting correlation structure than descriptor count, with the two hierarchical clustering approaches producing nearly equivalent outcomes and greedy maximum coverage consistently retaining more highly correlated descriptor subsets **(Figure 2b, Supplementary Table 39)**.

**Figure 2:**
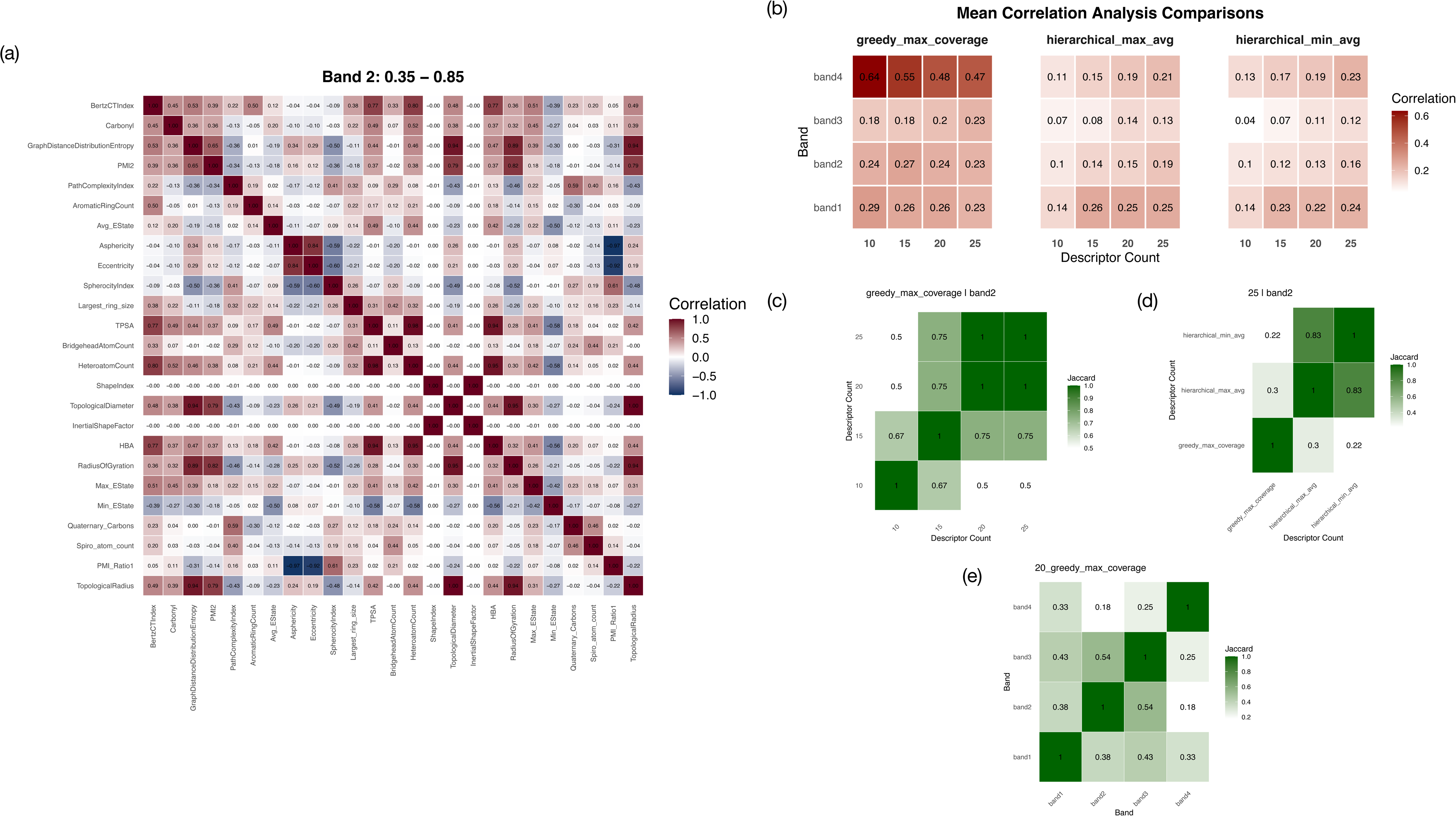
Descriptor Selection Analysis and Descriptor Set Overlap. **(a) Correlation Matrix of the Greedy Maximum Coverage Method (Correlation Range: 0.35–0.85; 25 Descriptors):** The heat map displays the pairwise Spearman correlation coefficients among the 25 descriptors selected by the Greedy Maximum Coverage method. Both axes represent the selected descriptors, and each cell corresponds to the correlation between a descriptor pair, with coefficient values annotated within the matrix. The color scale ranges from dark blue (−1; strong negative correlation) to dark red (+1; strong positive correlation), with white indicating no correlation (0). **(b) Heatmap of Mean Absolute Correlation (MAC) Across Descriptor Selection Strategies, Descriptor Counts, and Correlation Ranges:** The heat map compares the average descriptor redundancy across all experimental conditions. The three facets represent the descriptor selection strategies, the x-axis represents descriptor counts, and the y-axis represents correlation threshold ranges. Each cell is annotated with the corresponding MAC value, with darker shades of red indicating higher average correlation among selected descriptors. **(c) Jaccard Similarity Heat Map of Descriptor Overlap (Greedy Maximum Coverage; Band 2):** The heat map illustrates the overlap between descriptor sets generated by the Greedy Maximum Coverage method across different descriptor counts within Band 2. Both axes represent descriptor counts, and cell annotations indicate the Jaccard similarity coefficient. Darker shades of green correspond to greater descriptor overlap. **(d) Jaccard Similarity Heatmap of Descriptor Overlap (25 descriptors - Band 2):** The heat map illustrates the overlap between descriptor sets generated by the 0.35 - 0.85 correlation threshold range across different selection strategies within Band 2. Both axes represent descriptor selection strategy, and cell annotations indicate the Jaccard similarity coefficient. Darker shades of green correspond to greater descriptor overlap. **(e) Jaccard Similarity Heatmap of Descriptor Overlap (Greedy Maximum Coverage - 20 descriptors):** The heat map illustrates the overlap between descriptor sets generated by the Greedy Maximum Coverage method across different correlation threshold ranges with 20 descriptors. Both axes represent correlation threshold ranges, and cell annotations indicate the Jaccard similarity coefficient. Darker shades of green correspond to greater descriptor overlap.

#### 3.2.6. Descriptor Selection Strategy Optimization

To identify the most effective strategy for descriptor selection, Jaccard similarity-based correlation heatmaps were constructed while maintaining a constant correlation threshold and fixed number of selected descriptors. Comparative analysis across the different threshold-descriptor combinations revealed notable differences in redundancy patterns between the different techniques. Among all the evaluated approaches, the greedy maximum coverage method consistently exhibited the lowest cumulative Jaccard similarity value, indicating reduced overlap and greater diversity. The findings suggest that greedy maximum coverage was more effective in minimizing descriptor redundancy while preserving a more inclusive representation of the feature space **(Fig 2c; Supplementary Figures 40-41)**.

#### 3.2.7. Correlation Threshold and Descriptor Count Optimization

Similar Jaccard-similarity based correlation heatmaps were also constructed for identification of the best correlation threshold and number of descriptors. For correlation threshold optimization, the descriptor selection strategy and number of selected descriptors were kept constant, while for descriptor count optimization, the correlation threshold and descriptor selection strategy were maintained constant. Comparative analysis showed clear differences in descriptor redundancy patterns across both parameter settings. Among the evaluated configurations, band 2 (0.35 - 0.85) **(Fig 2d, Supplementary Figures 42-43)**, and a descriptor count of 20 exhibited lowest cumulative Jaccard similarity values. Since lower cumulative similarity indicates a reduced overlap and descriptor redundancy, these parameter combinations were considered optimal as they are able to provide broader and more diverse representation of the feature space **(Fig 2a, 2e; Supplementary Figures 44-45)**.

### 3.3 Pareto Front and Configuration Selection

Two optimization curves were generated to evaluate descriptor redundancy and information retention across varying correlation threshold ranges and descriptor counts, including mean absolute correlation (MAC) profiles and variance retention versus redundancy curves. Among the evaluated threshold bands, the 0.35–0.85 range (band 2) exhibited the most stable MAC profile across descriptor counts, whereas band 3 (0.25-0.85) demonstrated a consistently increasing MAC trend. In contrast, the 0.5–0.85 band showed a sharp reduction in MAC values from 10 to 15 descriptors followed by comparatively minor fluctuations at higher descriptor counts, while band 4 (0.35-0.75) exhibited a continuous decrease in MAC from 0.63 to 0.47. Collectively, these observations indicate that band 2 maintained the most stable redundancy behavior across descriptor-count configurations relative to the other threshold ranges **(Figure 3a, Supplementary Table 40)**.

**Figure 3:**
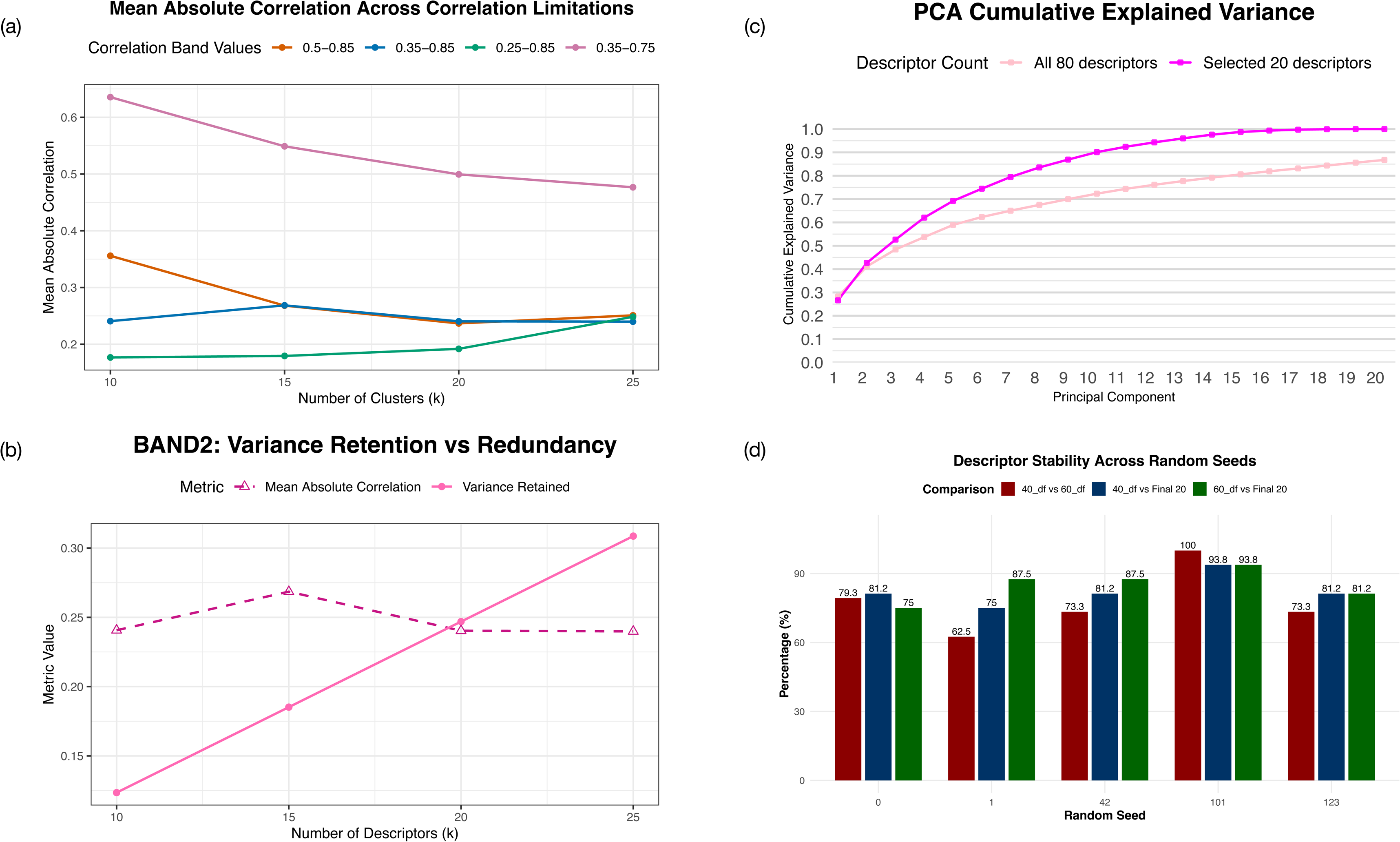
Validation of Selected Correlation Threshold Range, Descriptor Selection Strategy and Descriptor Count. **(a) Pareto Optimization of Mean Absolute Correlation across Correlation Limitations:** The line plot illustrates the mean absolute correlation (MAC) obtained across different descriptor counts and correlation threshold ranges. The x-axis represents the number of selected descriptors (*k*), while the y-axis represents the MAC value. Line colors correspond to the evaluated correlation threshold ranges: orange (0.50–0.85), blue (0.35–0.85), green (0.25–0.85), and pink (0.35–0.75). **(b) Variance Retained v/s Redundancy (0.35 - 0.85 Correlation Threshold Range):** The line plot summarizes the trade-off between retained variance and descriptor redundancy for the selected correlation threshold range. The x-axis represents descriptor count, and the y-axis represents the corresponding metric value. The solid pink line denotes the proportion of variance retained, while the dotted pink line with triangular markers represents the mean absolute correlation. **(c) Cumulative Explained Variance Plot (PCA):** The plot compares cumulative explained variance across the first 20 principal components for PCA performed on the complete descriptor set and the selected descriptor subset. The x-axis represents the number of principal components, and the y-axis represents cumulative explained variance. The dark pink line corresponds to PCA performed on the selected 20-descriptor subset, whereas the light pink line represents PCA performed on the complete set of 80 descriptors. **(d) Descriptor Stability across Perturbed datasets using Different Random Seeds and the complete 20 dataset:** The bar plot evaluates descriptor selection stability using multiple 60/40 dataset splits generated with different random seeds. The x-axis represents the random seeds, and the y-axis represents percentage descriptor overlap. Red bars indicate overlap between descriptor sets selected from the 60% and 40% splits, blue bars indicate overlap between the 40% split descriptor sets and the final 20-descriptor subset, and green bars indicate overlap between the 60% split descriptor sets and the final 20-descriptor subset. Percentage overlap values are annotated above each bar.

Variance retention versus redundancy analysis further demonstrated a steady increase in retained variance from 10 to 25 descriptors across all bands. However, redundancy also increased with descriptor count, with the highest MAC values observed in band 4. Notably, the 20-descriptor configuration within band 2 exhibited the lowest difference between variance retention and MAC values, indicating the most balanced trade-off between descriptor-space representation and redundancy minimization **(Figure 3b, Supplementary Table 40)**. Beyond 20 descriptors, increases in retained variance were comparatively modest relative to the associated increase in redundancy, suggesting diminishing returns at higher descriptor counts. These observations collectively support the selection of the 0.35-0.85 threshold range and the 20-descriptor configuration as the optimal parameter combination for descriptor selection **(Supplementary Figure 46; Supplementary Table 40)**.

### 3.4. Characterization of the final 20 Descriptor Subset

An intermediate subset of 25 descriptors was initially retained following optimization. Redundancy-guided refinement identified multiple descriptors with overlapping correlation behavior and similar physicochemical representation. Iterative removal of descriptors contributing limited additional information resulted in a final 20-descriptor subset with lower internal redundancy while maintaining overall descriptor coverage.

The final 20 selected descriptors included aromatic ring count, bridgehead atom count, heteroatom count, stereocenter count, triple bond count, PMI1, PMI2, spherocity index, inertial shape factor, carbonyl count, largest ring size, atom information content, TPSA, graph distance distribution entropy, path complexity index, shape index, and the average, maximum, and minimum electrotopological states. The final subset was predominantly composed of physicochemical (n=6, 30%) and graph/topological descriptors (n=5, 25%), followed by 3D descriptors (n=4, 20%) and electrotopological descriptors (n=3, 15%), whereas pharmacophoric and ring descriptors showed minimal representation, with only carbonyl count and largest ring size retained from their respective categories. Overall, 6 out of 18 physicochemical descriptors, 5 out of 17 graph/topological descriptors, 4 out of 14 3D descriptors, and all electrotopological descriptors were retained in the final subset, indicating preferential enrichment toward descriptors associated with molecular connectivity, physicochemical characteristics, structural complexity, and atom-level electronic environments **(Figure 2a; Supplementary Tables 2, 13b)**.

### 3.5. Information Retention and Descriptor Stability Analysis

To evaluate the robustness of descriptor selection, PCA was performed on both the final 20 descriptor subset and the complete 80 descriptor dataset. The PCA generated using the 20 selected descriptors achieved 100% cumulative explained variance at PC20 and exceeded 85% explained variance by PC9, indicating that the reduced descriptor set effectively captured the majority of dataset variance within a relatively small number of principal components. In comparison, PCA performed on the complete 80 descriptor set achieved 86.7% cumulative explained variance at PC20, suggesting that the selected descriptor subset retained substantial variance representation while providing a more compact and information-dense feature space **(Fig 3c, Supplementary Table 41)**.

Perturbation-based analysis further demonstrated the stability of descriptor selection across varying random seeds and dataset splits **(Supplementary Figures 47-51; Supplementary Tables 42-46)**. For every random seed tested, a minimum of 20 descriptors were shared between the 40% and 60% subsets. Most random seeds showed 81.2% similarity between the 40% subset and the final 20 descriptor set, with an overall similarity range of 75% to 93.8%. Similarly, the 60% subset showed descriptor similarity values ranging from 75% to 93.8%, with two random seeds demonstrating 87.5% similarity with the final descriptor set **(Fig 3d; Supplementary Table 47)**. These results indicate consistent descriptor retention despite perturbations in dataset composition, supporting the robustness of the descriptor selection framework.

### 3.6. Comparative Analysis of the Three Clustering Techniques

For KMeans clustering, Yeo–Johnson transformation reduced descriptor skewness from 8.05 to 0.46 after scaling. The elbow plot showed a continuous decrease in inertia without a distinct plateau, indicating the absence of well-separated natural clusters and no optimal cluster number **(Supplementary Figure 52c, Supplementary Table 48)**. Silhouette scores were generally higher for KMeans++ initialization at lower cluster counts, but decreased with increasing clusters, whereas random initialization remained relatively stable in the range of 0.12 to 0.123. Between cluster ranges of 420 to 440 and 450 to 460, both initialization methods showed similar silhouette scores, while at 435 and 445 clusters, random initialization slightly outperformed KMeans++. Davies–Bouldin indices were consistently higher for KMeans++ initialization except at k = 335 **(Figure 4a,b; Supplementary Table 48)**. The Calinski–Harabasz index decreased steadily with increasing cluster number for both initialization methods, indicating that higher cluster counts fragmented existing groups rather than improving separation **(Supplementary Figure 52a, Supplementary Table 48)**. Overall, the results suggest limitations of KMeans for molecular chemical space clustering due to the continuous and heterogeneous nature of molecular descriptor space, which does not conform to the spherical cluster assumptions of the algorithm.

**Figure 4:**
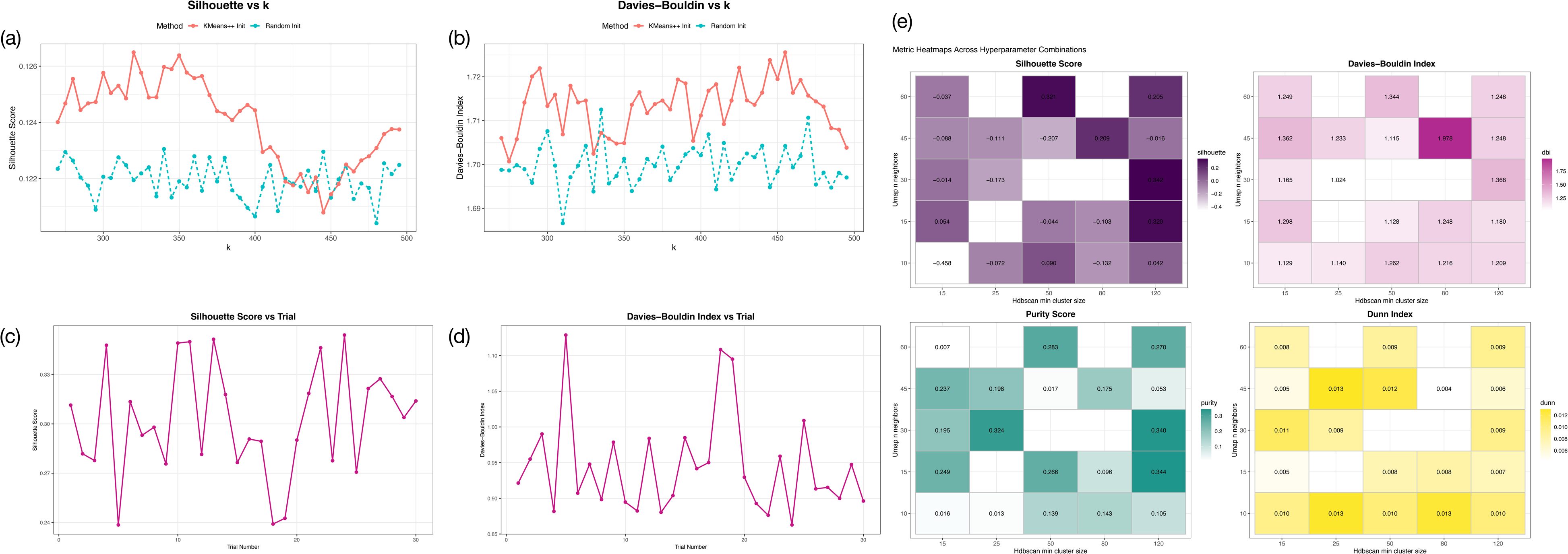
Summary of Performance of different Clustering Techniques on the Sampled Dataset. **(a) Silhouette Scores across different numbers of clusters (KMeans):** The line plot shows silhouette scores obtained across varying numbers of clusters for different K-Means initialization methods. The x-axis represents the number of clusters, and the y-axis represents the silhouette score. The red line corresponds to k-means++ initialization, while the green dotted line represents random initialization. (b) **Davies Bouldin across different numbers of clusters (KMeans):** The line plot presents Davies-Bouldin index values across varying numbers of clusters for different K-Means initialization methods. The x-axis represents the number of clusters, and the y-axis represents the Davies-Bouldin index. The red line corresponds to k-means++ initialization, while the green dotted line represents random initialization. **(c) Silhouette Scores across different trials (UMAP-Bisecting KMeans):** The line plot summarizes silhouette scores obtained across UMAP-Bisecting K-Means trials with different hyperparameter combinations. The x-axis represents the trial number, and the y-axis represents the silhouette score. **(d) Davies Bouldin Indices across different trials (UMAP-Bisecting KMeans):** The line plot summarizes Davies-Bouldin index values obtained across UMAP-Bisecting K-Means trials with different hyperparameter combinations. The x-axis represents the trial number, and the y-axis represents the Davies–Bouldin index. **(e) Summary of Different Clustering Performance Metrics across UMAP-HDBSCAN hyperparameter combinations:** The heat maps summarize clustering performance across different UMAP-HDBSCAN hyperparameter combinations. Purple, pink, yellow, and green heat maps represent silhouette score, Davies-Bouldin index, Dunn index, and cluster purity, respectively. Darker shades indicate higher metric values, and each cell is annotated with the corresponding score. The x-axis represents the HDBSCAN *min_cluster_size* parameter, while the y-axis represents the UMAP *n_neighbors* parameter.

For UMAP–HDBSCAN clustering, silhouette scores largely ranged from -0.2 to 0.5, with cluster counts between 0 and 200 showing higher scores in the range of 0.2 to 0.4, indicating better separation of distinct structural chemotypes **(Supplementary Figure 53b, 54, Supplementary Table 49)**. Silhouette scores declined towards 0 and slightly negative values in the 200 to 400 cluster range, suggesting subdivision of structurally related groups and reduced cluster distinctiveness. Lower quartile silhouette values ranged from -0.231 to 0.047, while maximum silhouette values consistently exceeded 0.3. The highest silhouette scores of 0.32, 0.321, and 0.342 corresponded to low Davies–Bouldin indices of 1.180, 1.344, and 1.248 respectively, indicating dense and well-separated clusters. Dunn indices ranging from 0.007 to 0.009 further reflected the complexity and clustering difficulty of the molecular dataset **(Figure 4e, Supplementary Table 49)**. Cluster purity also showed a consistent decrease with increasing noise fraction, indicating reduced cluster homogeneity due to weaker relationships between data points at higher noise levels **(Supplementary Figure 53a, Supplementary Table 49)**.

For the UMAP-Bisecting KMeans framework, higher silhouette scores were consistently associated with lower minimum distance values and specific neighbour-component combinations **(Supplementary Figure 55, Supplementary Table 50)**. The highest silhouette scores of 0.349 and 0.354 were observed with 10 neighbours and 5 components, and 40 neighbours and 3 components respectively **(Figure 4c)**. Across neighbour settings of 10 to 50, minimum distance values of 0.0 consistently produced silhouette scores above 0.32, while component counts of 2, 3, and 5 also showed consistently high scores above 0.34 **(Supplementary Figure 55, Supplementary Table 50)**. Euclidean and cosine metrics outperformed Manhattan distance, with cosine distance producing the highest silhouette score of 0.351 **(Supplementary Figure 56, Supplementary Table 50)**. Within the Bisecting KMeans stage, higher silhouette scores were observed with cluster counts between 450 and 500 and maximum iteration values between 100 and 300 **(Figure 4c)**. KMeans++ initialization generally produced a greater frequency of silhouette scores above 0.3, although the highest overall silhouette score of 0.324 was obtained using random initialization combined with the biggest inertia bisecting strategy **(Supplementary Figure 56, Supplementary Table 50)**. Overall, the best-performing configuration combined UMAP parameters of 40 neighbours, 5 components, minimum distance of 0.0, and euclidean or cosine metric with Bisecting KMeans settings of 500 clusters, biggest inertia bisecting strategy, and 300 maximum iterations **(Fig 4c, d; Supplementary Figures 55-57 ;Supplementary Table 50)**.

### 3.7. Full-Scale Clustering and Global Chemical Space Organization

UMAP-HDBSCAN was selected as the primary clustering framework due to its improved scalability and clustering performance. Unlike KMeans-based approaches, the framework was seen to be better suited for continuous and non-spherical chemical spaces. Across multiple hyperparameter combinations, UMAP-HDBSCAN consistently produced higher silhouette scores alongside lower and better-balanced Davies-Bouldin indices, indicating improved cluster compactness and separation. The method also showed higher capability in preserving local structural relationships during dimensionality reduction while effectively handling noise. This was particularly important for the present dataset, where substantial molecular diversity limited the effectiveness of centroid-based clustering approaches. Overall, the framework provided a more robust and chemically meaningful representation of the molecular landscape compared to conventional clustering methods.

Complete molecular clustering was performed using UMAP–HDBSCAN, with cluster quality evaluated before and after noise assignment to assess cluster stability and boundary ambiguity. Prior to noise assignment, clustering achieved a silhouette score of 0.42, indicating well-separated core clusters, although 48% of points were classified as noise, reflecting ambiguity at boundary regions. Following incorporation of noise points into clusters, the silhouette score decreased to 0.24, indicating reduced within-cluster cohesion. Clusters showed an average size of 412 molecules, suggesting relatively uniform and moderately large cluster distributions. Similarly, the Davies–Bouldin index increased from 0.85 to 1.15 after noise assignment, while the Calinski–Harabasz index decreased from 111 880.445 to 95 932.22, collectively indicating reduced cluster compactness and separation upon inclusion of uncertain points. Overall, the results suggest the presence of well-defined core chemotype clusters within the UMAP–HDBSCAN embedding, while also highlighting the intrinsic continuity and overlap present within the molecular chemical space.

### 3.8. Structural, Geometric and Chemical Characterization of Clusters

#### 3.8.1. Chemical Characterization

Dominant scaffold fraction (DSF) ranged from 0.0095 to 1.0 (mean = 0.302, median = 0.15), indicating substantial variation in scaffold dominance across clusters. Most clusters (67.56%) exhibited DSF values below 0.25, suggesting structurally diverse scaffold compositions, whereas 15.21% of clusters showed DSF values between 0.95 and 1.0, representing highly scaffold-dominated clusters. The DSF distribution was positively skewed (1.356), indicating enrichment toward lower scaffold dominance **(Supplementary Figure 58, Supplementary Table 51)**.

Descriptor profiling revealed that clusters were generally characterized by low aromaticity and high saturation. Mean aromatic ring count was low (mean = 1.196, median = 1), with 86.87% of clusters containing fewer than three aromatic rings **(Supplementary Figure 59a, Supplementary Table 51)**. In contrast, Fsp3 distributions were shifted toward higher saturation values (mean = 0.624), with 36.54% of clusters exhibiting Fsp3 means greater than 0.75 (**Supplementary Figure 59b, Supplementary Table 51)**. Hydrogen bond acceptor and donor means showed strong right-skewed distributions, with most clusters concentrated in low-to-moderate HBA and HBD ranges (**Supplementary Figure 59c, d; Supplementary Table 51)**. Mean logP values indicated moderate lipophilicity (mean = 4.641) **(Figure 5a; Supplementary Table 51)**, while molecular weight distributions were centered within the 100–600 Da range for most clusters (60.5%) (**Supplementary Figure 60b; Supplementary Table 51)**. Rotatable bond and TPSA distributions similarly showed enrichment toward chemically constrained and moderately polar molecular spaces (**Supplementary Figure 60c, d; Supplementary Table 51)**.

**Figure 5:**
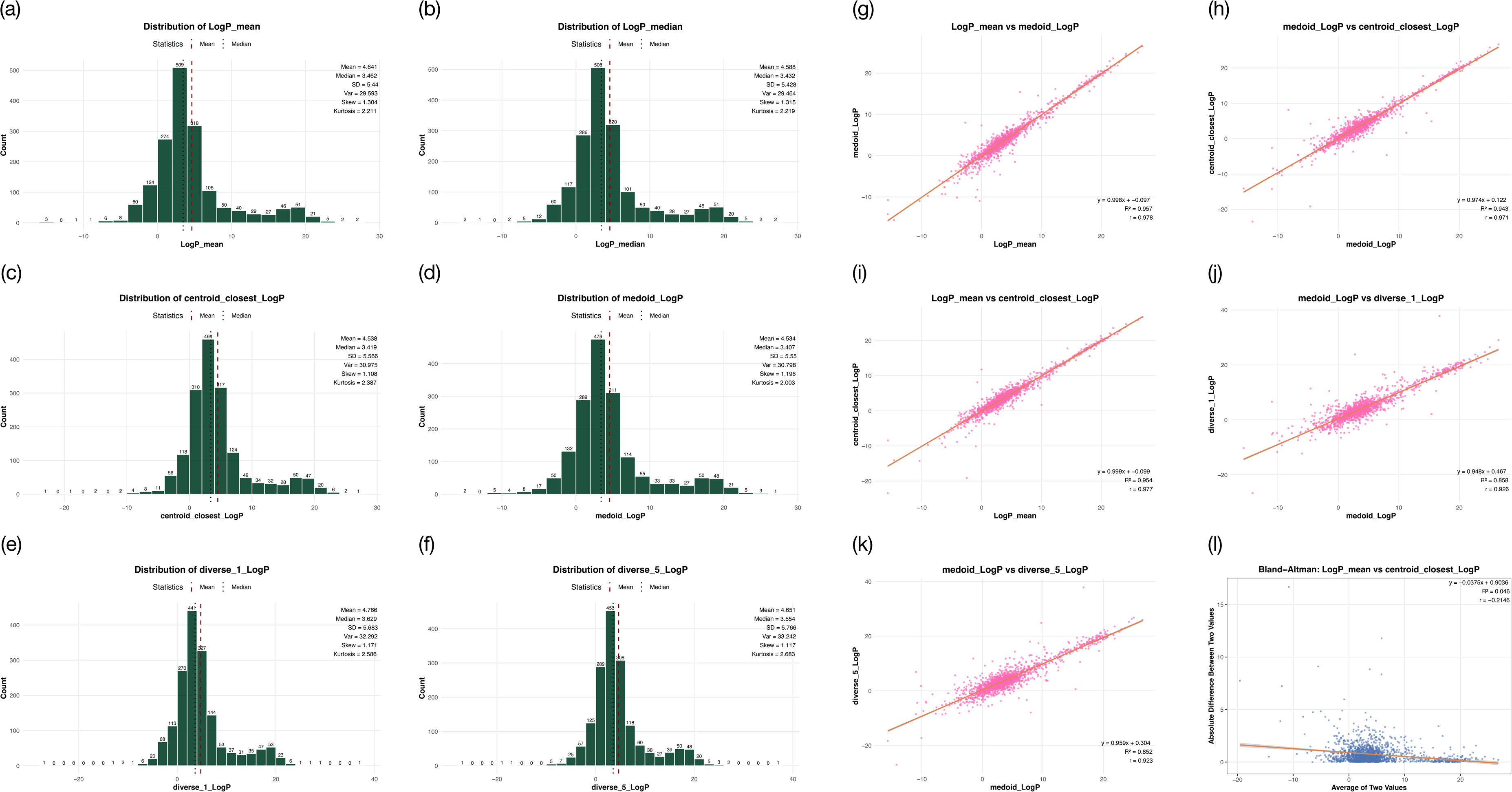
Chemical Profiling of Cluster Representatives Using Univariate and Bivariate Descriptor Analysis (LogP Example) **Distribution of Cluster Representative’s LogP values:** The histogram illustrates the distribution of LogP values across cluster representatives. The x-axis represents LogP values grouped into bins of width 2, while the y-axis indicates the frequency of cluster representatives within each bin. Bar annotations denote the number of cluster representatives in the corresponding bin. Summary statistics, including the mean, median, standard deviation, variance, skewness, and kurtosis, are displayed alongside the plot. The red dashed line indicates the mean LogP value, whereas the black dotted line indicates the median.

(a) Distribution of LogP mean values across Clusters
(b) Distribution of LogP median values across clusters
(c) Distribution of LogP values of centroid_closest representatives across Clusters
(d) Distribution of LogP values of medoid representatives across Clusters
(e) Distribution of LogP values of diverse_1 representatives across Clusters
(f) Distribution of LogP values of diverse_2 representatives across Clusters **Bivariate Analysis of Different Cluster Representatives:** The scatter plots illustrate the relationships between pairs of cluster representative descriptors, with LogP shown as an example. The x- and y-axes represent the descriptor values being compared, and each pink point corresponds to an individual cluster representative. A red regression line is overlaid to highlight the linear trend between the variables. The coefficient of determination (R²), Pearson correlation coefficient (*r*), and regression equation are annotated in the lower-right corner of each plot.

(g) Medoid LogP v/s mean LogP
(h) Centroid_closest LogP v/s medoid LogP
(i) Centroid_closest LogP v/s mean LogP
(j) diverse_1 LogP v/s medoid LogP
(k) diverse_5 LogP v/s medoid LogP **Bland-Altman Plot:** The x-axis illustrates the average of two values, and the y-axis indicates the mean absolute difference between the two values. A red regression line is overlaid to highlight the linear trend between the variables. The coefficient of determination (R²), Pearson correlation coefficient (*r*), and regression equation are annotated in the upper-right corner of each plot.

(l) Mean LogP v/s centroid_closest LogP

Correlation analyses demonstrated weak-to-moderate descriptor associations. Fcsp3 mean and logP mean showed a weak positive correlation (r = 0.271, R² = 0.073), indicating that increased saturation only mildly contributed to lipophilicity. Molecular weight and logP demonstrated similarly weak positive association (r = 0.257, R² = 0.066). In contrast, logP and TPSA exhibited a weak-to-moderate inverse relationship (r = -0.371, R² = 0.137), suggesting that higher lipophilicity was generally associated with lower polarity **(Supplementary Figure 61, Supplementary Table 51)**.

#### 3.8.2. Geometrical and Compactness Characterization

Mean core distance distributions indicated that most clusters remained geometrically compact on the UMAP manifold. Mean core distance was 0.283, with 83.6% of clusters occupying the 0.1–0.4 range. Post-noise assignment distances showed slightly broader dispersion (mean = 0.349), although 85.92% of clusters still remained within the 0.1–0.5 range, indicating limited geometric expansion after reassignment. Both distributions displayed strong positive skewness, reflecting the presence of a small number of highly diffuse clusters **(Supplementary Figures 62a, b; Supplementary Table 52)**.

Density gradient analysis demonstrated that most clusters retained stable density profiles following reassignment, with 1216 clusters (72.25%) exhibiting gradients between 0 and 0.1. Medoid-centroid distances were also low overall (mean = 0.086), suggesting strong geometric symmetry across clusters **(Supplementary Figures 62c, d; Supplementary Table 52)**. Only a small number of clusters demonstrated substantial asymmetry or density variation.

Strong positive correlations were observed between mean post-assignment distance and mean core distance (r = 0.923, R² = 0.852), medoid-centroid distance (r = 0.853, R² = 0.727), and post-distance standard deviation (r = 0.789, R² = 0.623). These findings indicate that increases in cluster spread were accompanied by increased asymmetry and internal geometric variability. Covariance volume showed only weak association with post-assignment distance (R² = 0.029) **(Supplementary Figure 64; Supplementary Table 52)**.

#### 3.8.3. Diversity, Separation and Cluster Confidence Analysis

Within-cluster Jaccard similarity demonstrated moderate internal structural similarity (mean = 0.354, median = 0.267), with most clusters concentrated within the 0.15–0.3 similarity range. Highly homogeneous clusters were comparatively rare, although a subset of clusters exhibited similarity values above 0.9. Separation ratio distributions indicated strong inter-cluster separation overall (mean = 3.328), with 1167 clusters (69.87%) exhibiting separation ratios greater than 2 **(Supplementary Figures 64a-b, Supplementary Table 53)**.

The separation ratio demonstrated a strong positive association with the dominant scaffold fraction (r = 0.745, R² = 0.555), suggesting that scaffold-dominated clusters tended to be more isolated in chemical space. In contrast, mean between-medoid similarity showed no association with within-cluster similarity **(Supplementary Figures 64c-e; Supplementary Table 51, 53)**.

Cluster confidence analysis revealed consistently high HDBSCAN assignment stability. Mean membership probability was high overall (mean = 0.926), with most clusters concentrated between 0.8 and 1.0. Variance in membership probability remained low (mean = 0.014), indicating limited uncertainty across clusters **(Supplementary Figures 65a-b, Supplementary Table 54)**. Variance probability and mean probability demonstrated a strong inverse relationship (r = -0.87, R² = 0.757), whereas dominant scaffold fraction showed negligible association with either confidence metric **(Supplementary Figures 65c-f; Supplementary Tables 51, 54)**.

### 3.9. Representative Molecule Analysis and Cluster Fidelity

#### 3.9.1. Cluster Fidelity Analysis

Core cluster size ranged from 100–531 molecules (mean = 143.96, median = 128), with most clusters (89.78%) concentrated between 100–200 molecules and only 24 clusters exceeding 300, indicating generally compact and consistent cluster cores **(Supplementary Figure 66a)**. Post-noise cluster size showed substantially greater dispersion, ranging from 100–2295 molecules (mean = 412.67, median = 339), with 75.1% of clusters between 100–500 molecules and only 4.21% exceeding 1000 molecules after reassignment, suggesting limited expansion for most clusters but substantial peripheral growth in a subset **(Supplementary Figure 66b)**. Core fraction ranged from 0.061–1.0 (mean = 0.442, median = 0.406), with 50.3% of clusters between 0.2–0.45, indicating moderate retention of dense cores following expansion **(Supplementary Figure 66c)**. HDBSCAN probability IQR remained low overall (mean = 0.124, median = 0.049), with 68.45% of clusters below 0.15 and only 4.1% above 0.5, suggesting consistently stable membership assignments **(Supplementary Figure 66, Supplementary Table 55)**.

Association analyses revealed moderate-to-weak relationships between stability metrics. Core and post-noise size demonstrated a moderate positive association (r = 0.504, R² = 0.254), indicating that larger cores generally expanded into larger clusters, although several clusters retained small cores while exhibiting substantial peripheral growth **(Supplementary Figure 67a, Supplementary Table 55)**. Core fraction showed weak negative association with density gradient (r = -0.29, R² = 0.084) and weak positive association with mean membership probability (r = 0.271, R² = 0.074), suggesting that clusters with larger stable cores tended to undergo lower density perturbation and slightly higher assignment confidence **(Supplementary Figure 67b-c; Supplementary Tables 52, 54-55)**. Moderate association with separation ratio (r = 0.476, R² = 0.227) and weak association with dominant scaffold fraction (r = 0.335, R² = 0.112) indicated that compact, scaffold-enriched clusters were generally more separated, although scaffold dominance alone poorly explained cluster stability **(Supplementary Figure 67d-e; Supplementary Tables 51, 53, 55)**.

#### 3.9.2. Representative Molecule Analysis

The mean-, median-, medoid-, and centroid-closest representatives exhibited highly similar descriptor distributions for MolWt, LogP, AromaticRingCount, FractionCSP3, TPSA, HBD, HBA, and RotatableBonds, indicating consistent preservation of the overall physicochemical landscape of the clustered chemical space. Central tendency measures remained closely aligned across these representative types, while comparable standard deviation and variance values suggested stable global descriptor distributions **(Figures 5a-d; Supplementary Figures 60-61, 68-73, Supplementary Table 56)**. Medoid and centroid-closest representatives consistently showed the smallest average absolute differences from cluster mean and median values across all descriptors, confirming that both strategies effectively capture the central physicochemical characteristics of the cluster population **(Figures 5e-f; Supplementary Figures 68-73, Supplementary Table 57)**. FractionCSP3 and AromaticRingCount displayed the most stable distributions overall, with relatively low dispersion and limited variation between representative strategies **(Supplementary Figures 68-73, Supplementary Tables 56-57)**.

In contrast, the diverse representatives produced broader and more heavy-tailed descriptor distributions across most physicochemical properties. MolWt, TPSA, HBD, HBA, and RotatableBonds showed markedly increased variance, skewness, and kurtosis among diverse representatives, indicating enrichment of structurally extreme, highly polar, hydrogen-bond-rich, and conformationally flexible compounds. Several diverse representatives exhibited substantially elevated kurtosis values, particularly for MolWt and TPSA, reflecting the presence of rare but highly extreme molecules within the sampled representative space **(Figures 5e-f; Supplementary Figures 74-83, Supplementary Table 56)**. Average absolute differences from cluster mean and median values were consistently larger for diverse representatives than for medoid or centroid representatives across all descriptors, while pairwise diverse differences further confirmed substantial physicochemical separation among diversity-selected compounds **(Supplementary Table 57)**. Overall, the results indicate that centroidal and medoid representatives preserve the dominant global physicochemical distribution of the clustered dataset, whereas diverse representatives expand descriptor-space coverage by preferentially sampling chemically extreme regions of the chemical space.

Across all molecular descriptors evaluated, central tendency-based representatives (mean, medoid, and centroid-closest) demonstrated consistently high concordance. Correlation coefficients remained near-unity for most descriptors (R ≈ 0.94-0.99, R² ≈ 0.88-0.98), with slopes close to 1.0, indicating strong linear preservation of descriptor structure across representation schemes **(Figures 5g-i; Supplementary Figures 84-86; Supplementary Table 58)**. Bland–Altman analysis further confirmed minimal systematic bias between these representatives, with narrow limits of agreement and low mean absolute differences **(Supplementary Figures 87-89; Supplementary Figure 59)**. Descriptors such as LogP, aromatic ring count, hydrogen bond donors/acceptors, and rotatable bonds exhibited particularly stable behavior, reflecting robust preservation of physicochemical properties regardless of representative selection strategy. The fractional sp³ character showed comparatively lower correlation but remained numerically stable due to its bounded distribution **(Supplementary Figures 84-89; Supplementary Tables 58-59)**.

In contrast, diverse representatives (diverse_1–diverse_5) exhibited consistently reduced agreement with central descriptors, with lower correlation (R ≈ 0.85–0.94), increased dispersion, and substantially higher RMSE and limits of agreement across all descriptors **(Figures 5j-l; Supplementary Figures 90-94, Supplementary Table 58)**. The effect was most pronounced for molecular weight and topological polar surface area, where variability increased several-fold relative to central representatives, indicating sensitivity of extreme or structurally diverse selections to cluster heterogeneity. Despite this dispersion, slope values generally remained near unity, suggesting that deviations are primarily variance-driven rather than systematic scaling distortions **(Supplementary Figures 90-99; Supplementary Tables 58-59)**. Overall, the results indicate a clear dichotomy between stability-preserving representatives (mean/medoid/centroid) and variability-enriching representatives (diverse selection), with the former reliably capturing cluster-level descriptor structure.

### 3.10. Representative Suitability for Drug Discovery Applications

Among the randomly selected clusters, five showed missing representative files: cluster 1147 (small), cluster 464 (medium), and clusters 1313, 964, and 1292 (large), indicating that missing entries were sparsely distributed across all size strata. The large clusters were retained despite these gaps, as they were primarily used for diversity assessment at the cluster level, where maintaining broad chemical space coverage was prioritized over complete representative-level completeness. For docking evaluation using 25 randomly selected molecules, five small clusters (993, 802, 591, 210, 598), five medium clusters (1308, 319, 952, 811, 971), and five large clusters (676, 831, 1292, 964, 1313) were assessed; while all small clusters had complete results, 3 medium clusters and 3 large clusters were complete, whereas clusters 811 and 971 (medium) and 1313 and 964 (large) had 2 missing molecules each, resulting in 24/25 results for 971, 811, and 964, and 23/25 for 1313.

Among the randomly selected molecules from the 15 analyzed clusters, four demonstrated consistently strong docking behavior with highly favorable mean affinities and relatively narrow score distributions, while four exhibited weak overall binding profiles indicative of low predicted activity. Three clusters showed intermediate docking performance, whereas four displayed broad affinity ranges exceeding approximately 3 kcal/mol, suggesting substantial internal heterogeneity and the coexistence of both strong and weak binders within the same chemical group. The close agreement between mean and median affinity values across most clusters further indicated relatively symmetric docking score distributions with minimal outlier influence. Overall, the observed variability confirms that the clustering framework effectively partitioned the natural product chemical space into regions with distinct binding characteristics toward the target protein **(Supplementary Figure 100-101, Supplementary Tables 60-61)**.

Representative-based docking analysis showed variability across selection strategies, with diverse_1 representatives exhibiting the widest docking score range (-11.7 to -3.8 kcal/mol). Centroid_closest and diverse_5 representatives displayed the broadest overall affinity distributions, while diverse_1, diverse_2, and diverse_3 showed comparatively narrower ranges. Mean docking scores ranged from -8.21 (diverse_1) to -8.75 kcal/mol (centroid_closest), and median scores ranged from -7.65 (diverse_2) to -8.8 kcal/mol (centroid_closest). Except for medoid and diverse_2 representatives, all representative types exhibited median scores greater than their corresponding means, indicating slight skewness toward lower-affinity outliers (Figure 6a, Supplementary Tables 62-63).

**Figure 6:**
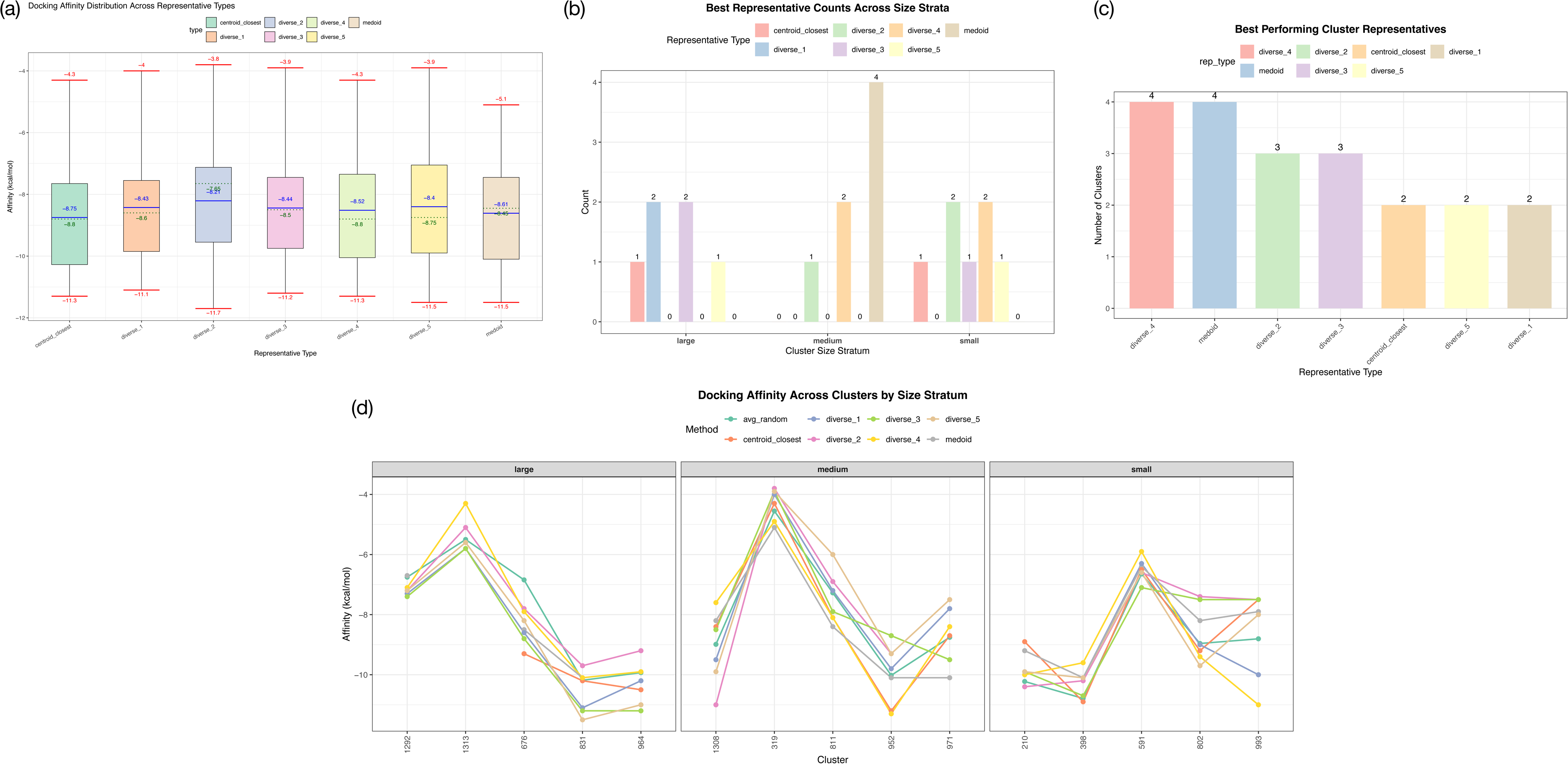
Docking Score-Based Analysis of Representative Suitability for Drug Discovery using KEAP1. **(a) Distribution of Docking Scores of Different Cluster Representatives:** The box plots summarize the distribution of docking scores across different cluster representative types. The x-axis represents the representative type, while the y-axis represents binding affinity (kcal/mol). For each representative type, the box denotes the interquartile range (Q1–Q3). The median affinity is indicated by a green dotted line, with the corresponding value annotated below the line, while the mean affinity is represented by a solid blue line, with its value annotated above the line. Red horizontal lines indicate the minimum and maximum observed affinities, with their corresponding values annotated below and above the respective lines. Different box colors distinguish the representative types. **(b) Best Representative Count across Size Strata:** The bar plot summarizes the number of clusters in which each representative type achieved the best docking score among the seven representative selection strategies. The x-axis represents the cluster size strata, while the y-axis indicates the number of clusters for which a given representative produced the most favorable docking score. Different bar colors correspond to different representative types. Numerical annotations above the bars denote the total number of clusters in which the respective representative achieved the best docking score. **(c) Global Analysis of Best Performing Representative:** The bar plot summarizes the number of clusters in which each representative type achieved the best docking score among the seven representative selection strategies. The x-axis represents the representative type, while the y-axis indicates the number of clusters for which a given representative produced the most favorable docking score. Different bar colors correspond to different representative types. Numerical annotations above the bars denote the total number of clusters in which the respective representative achieved the best docking score. **(d) Size-Stratified Comparison of Docking Scores of Different Representatives with Average of Random Molecules:** The faceted multi-line plot compares the docking scores of different cluster representative types against the average docking score of randomly selected molecules. Each facet corresponds to a cluster size stratum. The x-axis represents the cluster ID, while the y-axis represents binding affinity (kcal/mol). Line colors distinguish the representative types: centroid closest (orange), diverse 1 (blue), diverse 2 (pink), diverse 3 (green), diverse 4 (yellow), diverse 5 (camel), and medoid (grey). The dark green line represents the average docking score of 25 randomly selected molecules from each cluster, providing a baseline for comparison.

Diverse_4 and medoid representatives most frequently produced the best docking scores, each emerging as the top-performing representative in four clusters, followed by diverse_2 and diverse_3 with three clusters each, while centroid_closest, diverse_1, and diverse_5 were optimal in two clusters each. No consistent representative-selection pattern was observed across clusters overall **(Figure 6b, Supplementary Table 64a-b)**. Size-stratified analysis showed that large clusters favored diverse_1 and diverse_3 representatives, medium-sized clusters were predominantly represented by medoids, and small clusters favored diverse_2 and diverse_4 representatives, indicating that optimal representative selection varied with cluster size **(Figure 6c, Supplementary Table 64a, c)**.

Overall, the representative selection strategies were able to reasonably reproduce the average docking behavior of their parent clusters, as most representative affinities remained close to the corresponding average random docking score. Medoid and centroid-closest representatives generally exhibited the smallest deviations from cluster averages, indicating that central representatives effectively captured the typical affinity landscape of the cluster. In contrast, diversity-based representatives showed greater variability and frequently identified affinity extremes within the same cluster. Several clusters demonstrated substantially stronger affinities for specific diverse representatives relative to the average random score, including cluster 1308 (diverse_2,-11.0 kcal/mol vs. average random, -8.99 kcal/mol), cluster 831 (diverse_5, -11.5 kcal/mol vs. -10.18 kcal/mol), and cluster 993 (diverse_4, -11.0 kcal/mol vs. -8.80 kcal/mol) **(Supplementary Figure 102, Supplementary Table 65)**. These findings suggest that while central representatives are more suitable for approximating average cluster behavior, diversity-oriented representatives are more effective for exploring chemically distinct regions associated with enhanced docking performance.

Size-stratified analysis showed that small clusters generally exhibited the closest agreement between representative docking scores and average random cluster affinities, indicating relatively homogeneous chemical composition within these clusters. Clusters 210, 398, and 591 displayed only minor deviations among representative strategies, although cluster 993 represented an exception with a strongly favorable diverse_4 representative. Medium-sized clusters showed moderate variability, with occasional extreme-performing diverse representatives, as observed in cluster 1308. Large clusters exhibited the widest spread in representative docking scores, reflecting greater internal chemical heterogeneity and broader coverage of chemical space. For example, cluster 831 showed representative affinities ranging from -9.7 to -11.5 kcal/mol, while cluster 964 ranged from -9.2 to -11.2 kcal/mol **(Figure 6d, Supplementary Table 65)**. These observations indicate that cluster heterogeneity increases with cluster size, thereby increasing the importance of diversity-based representative selection for capturing high-affinity subregions within large clusters.

## 4. Discussion

Molecular clustering within the small-molecule chemical space remains challenging, primarily due to the scale, structural heterogeneity and high-dimensional nature of the data (Talevi & Bellera, 2024). These challenges grow to be highly pronounced in the natural product space, which is characterized by its high scaffold diversity and structural complexity (Chávez Hernández et al., 2020), caused due to evolutionary selection for biologically relevant interactions (Hong, 2011). Previous clustering studies are largely restricted to smaller datasets (Hernández-Hernández & Ballester, 2023), or subsets enriched for compounds with known drug discovery potential (Tao et al., 2015b). Many existing approaches also rely upon molecular fingerprints, and are often developed with existing and narrow drug discovery contexts. Hence, comprehensive and large-scale investigations integrating descriptor-based characterization, representative analysis, and broader exploration of the natural product chemical space are highly limited. The present study addresses these limitations, through large-scale clustering, combined with systematic evaluation of the descriptor space and extensive cluster characterization, and further demonstration of these approaches for downstream drug discovery applications.

Initial scaffold-based investigation of the dataset revealed the presence of extensive diversity, along with limited scaffold redundancy and broad structural dispersion. Strong right skew showed a “few-common, many rare” organization of the space, along with the prevalence of the benzene ring showing a widespread prevalence of aromatic frameworks. The sharp decline in scaffold frequencies beyond the most common scaffolds also indicates that no single chemotype is highly dominant. A high proportion of scaffolds are also seen to occur at low frequencies, with many singleton and low-occurrence scaffolds, which demonstrates a high amount of structural novelty and biosynthetic specialization. The observed scaffold landscape supports the need for diversity-aware representation. The overall results seen as consistent with scaffold diversity observed in a secondary metabolites of medicinal fungi (MeFSAT), as well as other chemical libraries including NPALTAS-Bacteria and NPATLAS-Fungi (Vivek-Ananth et al., 2023).

### 4.1 Correlation Analyses of the Descriptor Space and Feature Selection

Complete correlation analysis revealed substantial heterogeneity within the descriptor space, indicating that most descriptors encode relatively independent or weakly associated information. Although positive correlations were more prevalent overall, only a small fraction of descriptor pairs exhibited very high correlations (>|0.8|), suggesting limited large-scale redundancy across the feature set. Strong correlations were primarily localized within related descriptor families, including PMI, Chi indices, atom-count, size, and topological descriptors. The association between PMI and Chi descriptors is likely driven by shared dependence on molecular size and molecular weight, consistent with previous observations of size-driven spurious correlations in molecular descriptor analysis (Estrada, 2002). Collectively, the coexistence of broad heterogeneity and localized redundancy supports the use of feature reduction and threshold-based descriptor filtering strategies to minimize redundant information while preserving the overall variance structure of the dataset.

Analysis across correlation threshold ranges, descriptor counts, and selection strategies revealed distinct selection patterns. Moderate correlation ranges (Bands 1–3) showed substantial descriptor overlap across all methods, indicating the retention of a stable core descriptor set under moderate redundancy filtering. Relaxation of the lower correlation bound introduced additional descriptors while preserving this shared core structure, suggesting that many retained descriptors encode chemically robust information within the chemical space. In contrast, Band 4, representing stricter correlation thresholds, removed several strongly correlated descriptor groups observed in broader ranges. The reduced overlap and enrichment of Chi indices and topological descriptors indicate that stricter filtering preferentially retains complementary structural and topological information rather than highly co-varying physicochemical properties. Overall, the results suggest a transition from conserved descriptor retention under broader thresholds to increased descriptor diversity under stricter filtering, while consistently preserving a core set linked to molecular geometry and structural complexity.

Descriptor selection strategies exhibited distinct behaviors, reflecting different balances between descriptor diversity and redundancy. Greedy maximum coverage consistently retained the highest number of uniquely selected descriptors, as reflected by lower Jaccard similarity values, whereas the hierarchical clustering approaches showed substantial overlap, indicating a shared preference for structurally central descriptors and similar correlation-based grouping behavior. Among the hierarchical methods, the minimum average correlation-based approach demonstrated greater descriptor conservation across filtering conditions, suggesting higher selection stability. Across all methods, molecular complexity and geometry-related descriptors, including BertzCT and PMI ratio descriptors, were consistently enriched, indicating that topological complexity and three-dimensional shape are key features of the natural product chemical space. The recurrent enrichment of physicochemical, pharmacophoric, graph-based, and 3D descriptors further suggests that global structural and physicochemical properties are major contributors to dataset organization.

Descriptor count analysis revealed increased selection stability with increasing feature-space dimensionality. The 10- and 15-descriptor subsets showed inconsistent overlap patterns, suggesting that aggressive dimensionality reduction promotes competitive descriptor selection and greater method-dependent variability. In contrast, the 20- and 25-descriptor subsets exhibited substantial conservation across selection strategies and threshold ranges, indicating convergence toward a stable descriptor structure at higher dimensionality. Strong overlap between the 20- and 25-descriptor subsets further suggests that larger descriptor sets expand upon a conserved core rather than replacing previously retained descriptors. The enrichment of Chi indices, PMI ratios, and asphericity-related descriptors indicates the progressive inclusion of geometric and topological complexity alongside a stable core descriptor set.

### 4.2 Selection and Validation of Optimal Descriptor Selection Methods

Comparative analysis of descriptor selection strategies revealed substantial differences in redundancy minimization and descriptor-space coverage. Jaccard similarity analyses showed that the greedy maximum coverage approach produced lower cumulative similarity values than the hierarchical methods, indicating broader structural representation with reduced localized redundancy. Optimization of correlation threshold ranges and descriptor counts identified the 0.35–0.85 threshold range (Band 2) and the 20-descriptor configuration as the most balanced parameter combination, based on consistently low cumulative similarity values. Collectively, the results suggest that moderate redundancy filtering combined with intermediate descriptor dimensionality provides the most stable and informative representation of the descriptor space.

Pareto-based optimization further supported Band 2 and the 20-descriptor configuration as the optimal selection regime. Mean absolute correlation (MAC) analysis showed that Band 2 maintained the most stable redundancy profile across descriptor counts, whereas other bands exhibited progressive increases, decreases, or fluctuations in redundancy with increasing dimensionality. Although cumulative retained variance increased with descriptor count across all bands, redundancy also increased concurrently, particularly under stricter filtering conditions. Notably, the 20-descriptor configuration within Band 2 showed the smallest difference between retained variance and MAC, indicating the most balanced trade-off between descriptor-space coverage and redundancy minimization. Beyond 20 descriptors, gains in retained variance became marginal relative to the corresponding increase in redundancy, suggesting diminishing informational returns at higher dimensionality. Overall, the results indicate that descriptor optimization is governed by a balance between variance retention and redundancy control.

Redundancy-based refinement of the optimized descriptor subset produced a final 20-descriptor set enriched in physicochemical, graph-based, topological, and 3D descriptors, indicating preferential retention of features associated with molecular topology, connectivity, and geometry. The subset preserved descriptors representing both global structural properties and localized molecular characteristics. PCA-based evaluation showed strong information retention, with the reduced subset exceeding 85% cumulative variance at PC9 and reaching 100% variance retention by PC20, outperforming the full 82-descriptor set at comparable dimensionality. Perturbation-based analysis further demonstrated robust descriptor conservation across varying dataset splits and random seeds, with consistently high similarity between perturbed subsets and the final descriptor set. Collectively, these findings indicate that the framework generates a compact, low-redundancy, and stable feature space while preserving chemically relevant structural variance, supporting its suitability for downstream unsupervised and supervised machine learning applications.

### **4.3.** Performance of Algorithms and Full-Scale Clustering of the Dataset

Comparative evaluation of clustering algorithms on a 50,000-molecule subset revealed substantial differences in their ability to represent the diverse organization of the natural product chemical space. Although the Yeo–Johnson transformation effectively reduced descriptor skewness, conventional KMeans showed limited clustering performance across most configurations. The absence of a clear elbow point, coupled with steadily decreasing CH scores, suggested the lack of well-separated spherical cluster structures within the dataset. Low and stable silhouette scores across kmeans++ configurations, together with high DBI values, further indicated substantial cluster overlap and fragmentation of structurally related chemotypes. Incorporation of UMAP improved clustering performance, particularly in the UMAP–Bisecting KMeans framework, where lower minimum distance and moderate component values produced higher silhouette scores. The improved performance of cosine and Euclidean distance metrics further suggested that molecular similarity was better captured through continuous geometric and angular relationships. However, centroid-based approaches remained limited by the continuous and non-spherical nature of the chemical space, consistent with the low silhouette and high DBI values observed for hierarchical clustering (Hernández-Hernández & Ballester, 2023).

Among all approaches, UMAP–HDBSCAN showed the most chemically meaningful performance, consistently achieving higher silhouette scores and lower Davies–Bouldin indices than centroid-based methods, indicating better separation and local compactness. Higher silhouette values at lower cluster numbers suggest the presence of well-defined core chemotype regions, while the decline at higher cluster counts reflects subdivision of related molecular neighborhoods rather than formation of distinct clusters. This behavior aligns with the method’s ability to capture the irregular, non-spherical structure of natural product chemical space. Full-scale clustering further supported this, with strong initial silhouette scores indicating dense core molecular neighborhoods, followed by reduced performance after noise assignment due to boundary overlap. Overall, the results indicate that natural product chemical space behaves as a continuous structural manifold, where density-based clustering with nonlinear embedding provides a more appropriate representation than centroid-based approaches. Results also show consistency with previous UMAP studies on molecular clustering (Hernández-Hernández & Ballester, 2023).

### **4.4.** The Global Structure and Chemical Space Architecture of the Clustering Outcomes

A comprehensive evaluation of clustering outcomes was performed to assess global structural organization, incorporating cluster-level scaffold distribution, physicochemical trends, geometric descriptors, density patterns, and stability analyses.

Cluster-level scaffold organization indicates a globally heterogeneous yet locally structured chemical architecture. The generally low dominance of any single scaffold within most clusters suggests that multiple scaffold frameworks coexist within individual groups rather than being governed by a single core structure, reflecting strong structural mixing within local chemical neighborhoods. Only a limited number of clusters show pronounced scaffold dominance, indicating the presence of well-defined and isolated chemotype regions corresponding to structurally constrained natural product families. Overall, the distribution reflects a dual regime comprising a small number of scaffold-enriched clusters alongside a larger fraction of scaffold-diffuse clusters, supporting a chemical space that is globally continuous but locally structured.

Physicochemical trends across clusters indicate that most groups are composed of molecules with intermediate values of key properties such as hydrophobicity, polarity, and molecular size, rather than extremes. This suggests that clustering does not segregate molecules based on outlier physicochemical characteristics but instead groups chemically balanced compounds. Across clusters, properties such as lipophilicity, molecular weight, and polarity show weak inter-correlations, indicating largely independent variation and limited predictive dependence between descriptors. In addition, clusters are enriched in more saturated and structurally complex molecules compared to flat or purely aromatic systems, suggesting a stronger influence of three-dimensional structural features (Navarro-Vázquez et al., 2018). Overall, the results indicate that clustering is driven by a combination of multiple weakly correlated physicochemical and structural signals rather than any single dominant descriptor, consistent with the inherently multi-dimensional nature of natural product chemical space.

Geometric structure and stability analyses indicate that clusters are compact and internally consistent, embedded within a broader continuous chemical landscape. Most clusters form tight regions in embedding space, reflecting both spatial proximity and descriptor-level similarity among member molecules. Inclusion of noise leads to gradual cluster expansion rather than abrupt boundary shifts, suggesting smooth transitions between chemical neighborhoods and non-discrete cluster boundaries. Assignment confidence is highest within cluster cores and decreases toward boundary regions, indicating well-defined central regions with uncertainty concentrated at transition zones. Scaffold purity further supports this structure, with high-purity clusters appearing more isolated, while scaffold-diverse clusters act as bridges across the chemical manifold. Overall, the results support a continuous manifold structure for natural product space with dense local groupings rather than discrete, well-separated chemical classes.

### **4.5.** Representative Analysis and their Suitability for Drug Discovery Applications

Representative-based analysis distinguishes between stability-conserving (medoid and centroid-closest) and diversity-increasing selections, with central representatives closely matching cluster mean and median descriptor profiles and showing minimal deviation, thereby effectively capturing the typical chemical “core” of each cluster. In contrast, diversity-based representatives shift toward extreme descriptor values, increasing variability and exploring boundary regions of the cluster space. Despite this expansion toward extremes, underlying descriptor relationships and overall cluster structure remain stable, indicating that diversity sampling does not disrupt intrinsic patterns but instead extends coverage to underrepresented regions. Collectively, central representatives reflect average cluster identity, while diverse representatives complement them by capturing peripheral chemical variability within the same structured chemical landscape.

Docking results further reinforce the complementary roles of central and diverse representative selection. Central representatives (medoid and centroid-closest) consistently align with the average cluster binding behaviour, making them reliable proxies for estimating typical biological activity within each cluster. In contrast, diverse representatives exhibit a wider distribution of docking scores, capturing both weak and strong binders within the same cluster and thereby revealing hidden intra-cluster variability that is not reflected by central measures alone. Importantly, some diverse representatives outperform central representatives in binding affinity, highlighting their relevance for exploratory screening and early-stage hit discovery. Cluster size also plays a significant role, with larger clusters showing greater variability in docking outcomes across representatives, indicating increased internal chemical and biological heterogeneity. Overall, the results demonstrate that central representatives effectively summarize average activity, while diverse representatives are necessary to capture activity extremes and structural variability, together providing a more complete representation of cluster-level bioactivity.

### **4.6.** Limitations and Future Directions

Despite the comprehensive nature of the analysis, several limitations should be acknowledged in interpreting the results. The clustering performance is influenced by the presence of noise within the dataset, where increasing noise levels lead to a reduction in silhouette scores, reflecting the intrinsic difficulty of defining sharp boundaries in a continuous and highly overlapping chemical space. In addition, the descriptor space is limited to conventional physicochemical, topological, and 3D features, without incorporation of quantum chemical or ADMET-related descriptors, which constrains the depth of both chemical and pharmacokinetic interpretation. The study is also restricted to a single natural product dataset, which limits the generalizability of the findings across broader chemical libraries. Furthermore, clustering is performed in a reduced feature space, meaning that the resulting groupings represent structure in transformed descriptor space rather than explicit or experimentally validated chemotypes. Finally, the docking analysis is target-specific, and therefore does not capture broader biological activity across multiple protein systems. Overall, these limitations highlight the need for more comprehensive descriptor integration, multi-dataset validation, and multi-target evaluation in future studies.

Building on the present framework, several directions can be pursued to further strengthen and extend the analysis. A more detailed biological interpretation can be achieved by linking individual clusters to specific protein families or functional target classes, enabling a shift from purely structural grouping toward functional chemical space mapping. Expanding the analysis across multiple natural product and bioactive compound datasets would further help to assess the robustness and transferability of the observed clustering structure and representative selection behavior. From a methodological perspective, the use of graph-based learning and attention-driven models offers a promising alternative to descriptor-based representations, potentially capturing molecular structure in a more direct and flexible manner. In addition, extending the descriptor space to include quantum chemical properties and ADMET-related features would improve both the resolution of clustering and the biological relevance of the derived chemical organization. Collectively, these directions point toward a more integrated and biologically grounded framework for exploring complex molecular spaces.

## Conclusion

In conclusion, the present study provides a large-scale clustering-based analysis of the natural product chemical space represented within the COCONUT database, revealing a highly heterogeneous yet structurally organized molecular landscape. The results demonstrate that scaffold diversity within the dataset is not only consistent with previously described natural product spaces, but is even more pronounced, with coexistence of scaffold-dominant clusters, structurally diverse regions, and transitional clusters connecting neighboring chemotypes. Despite the complexity of the dataset, the reduced descriptor framework was able to preserve substantial chemical-space structure while enabling robust clustering and representative analysis. The identified clustering patterns and representative-selection strategies provide a foundation for downstream applications in drug discovery, biosynthetic investigations, and omics-driven molecular exploration. Overall, the study highlights the potential of integrated clustering frameworks for systematic organization and interpretation of large-scale natural product chemical spaces.

## Ethics

This study did not require approval from a human ethics committee or an animal welfare board.

## Data Accessibility

Scripts and supplementary data are available at: https://github.com/shrek-28/DescriptorClusteringNPSpace

## Author’s Contributions

PK: Conceptualization, Project administration, Supervision, Writing - review and editing; GS: Conceptualization, Data curation, Formal Analysis, Methodology, Investigation, Writing - Original draft, Visualization; Validation; AD - Data curation, Methodology; AN: Conceptualization, Project administration, Supervision, Writing - review and editing

## Declaration of Generative AI and AI-assisted Technologies in the Manuscript Preparation Process

The authors acknowledge the use of generative AI-assisted tools and technologies for the purpose of paraphrasing only.

## Acknowledgement

We would like to thank the Department of Biotechnology, Dayananda Sagar College of Engineering for infrastructural support. We would also like to thank J Varun Chowdary and Pranav Mehta, for technical assistance.

