## Supplementary figures for "Mapping Chemical Diversity: Descriptor-Guided Clustering of Natural Products in the COCONUT Database"

Figure 1

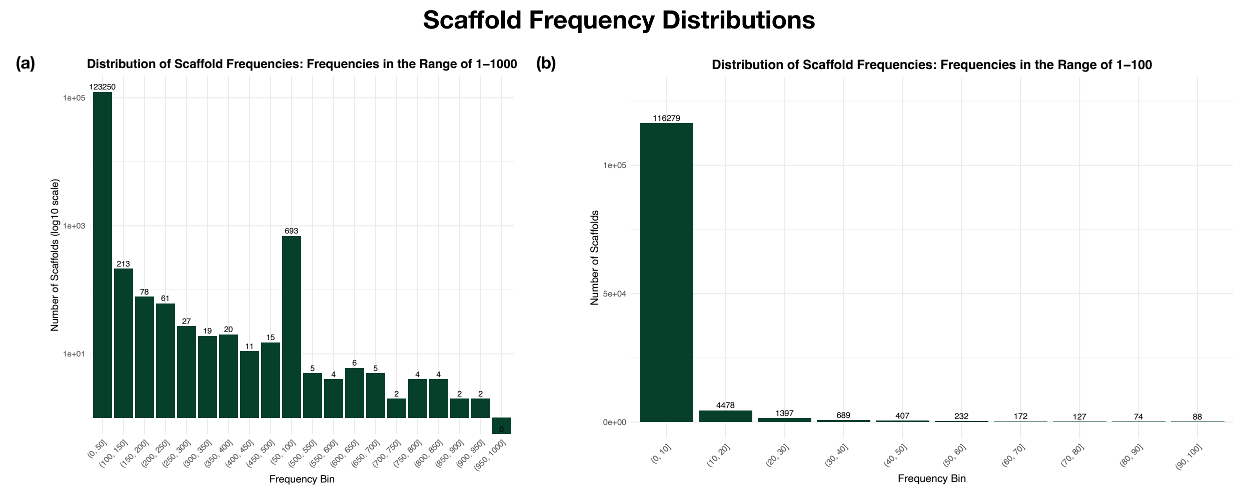


**Supplementary Figure 1. Scaffold frequency distributions across the dataset.**

**(a) Distribution of scaffold frequencies in the range 1–1000:** The x-axis represents scaffold frequency bins (bin width = 50), while the y-axis indicates the number of scaffolds on a $log_{10}$ scale. Numbers above the bars denote the scaffold count within each frequency bin.

**(b) Distribution of scaffold frequencies in the range 1–100**: The x-axis represents scaffold frequency bins (bin width = 10), while the y-axis indicates the number of scaffolds. Numbers above the bars denote the scaffold count within each frequency bin.

**Figure 2**

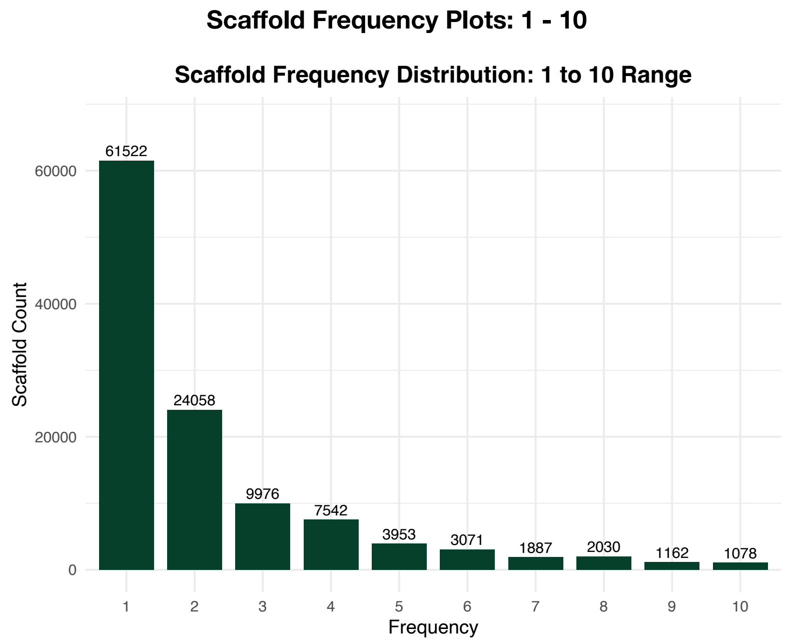


**Supplementary Figure 2: Scaffold frequency distributions across the dataset (Range of 1-10):** The x-axis represents the frequency of scaffold occurrence, while the y-axis indicates the number of scaffolds that occur at the given frequency. Numbers above the bards denote the scaffold count for the corresponding frequency.

**Figure 3**

#### Distribution of Pairwise Correlation Coefficients

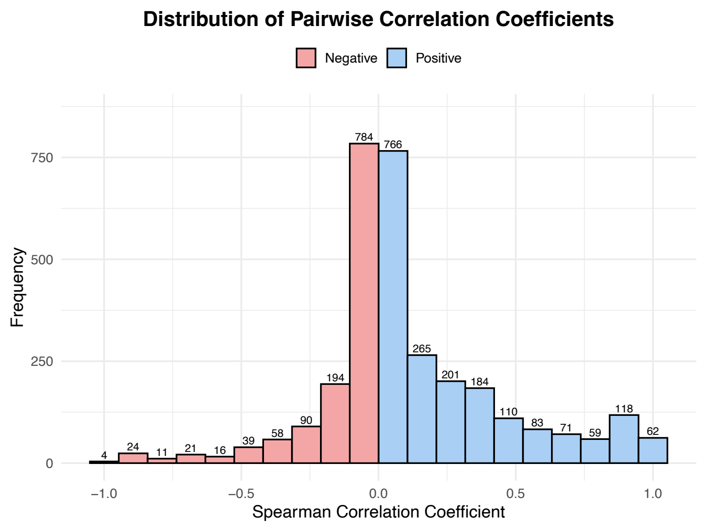


**Supplementary Figure 3: Distribution of Pairwise Correlation Coefficients:**
The x-axis represents ranges of Spearman correlation coefficient values, while the y-axis indicates the frequency of coefficients within each range. Numbers above the bars denote the number of coefficients in the corresponding range. Red bars represent negative correlation coefficients, whereas blue bars represent positive correlation coefficients.

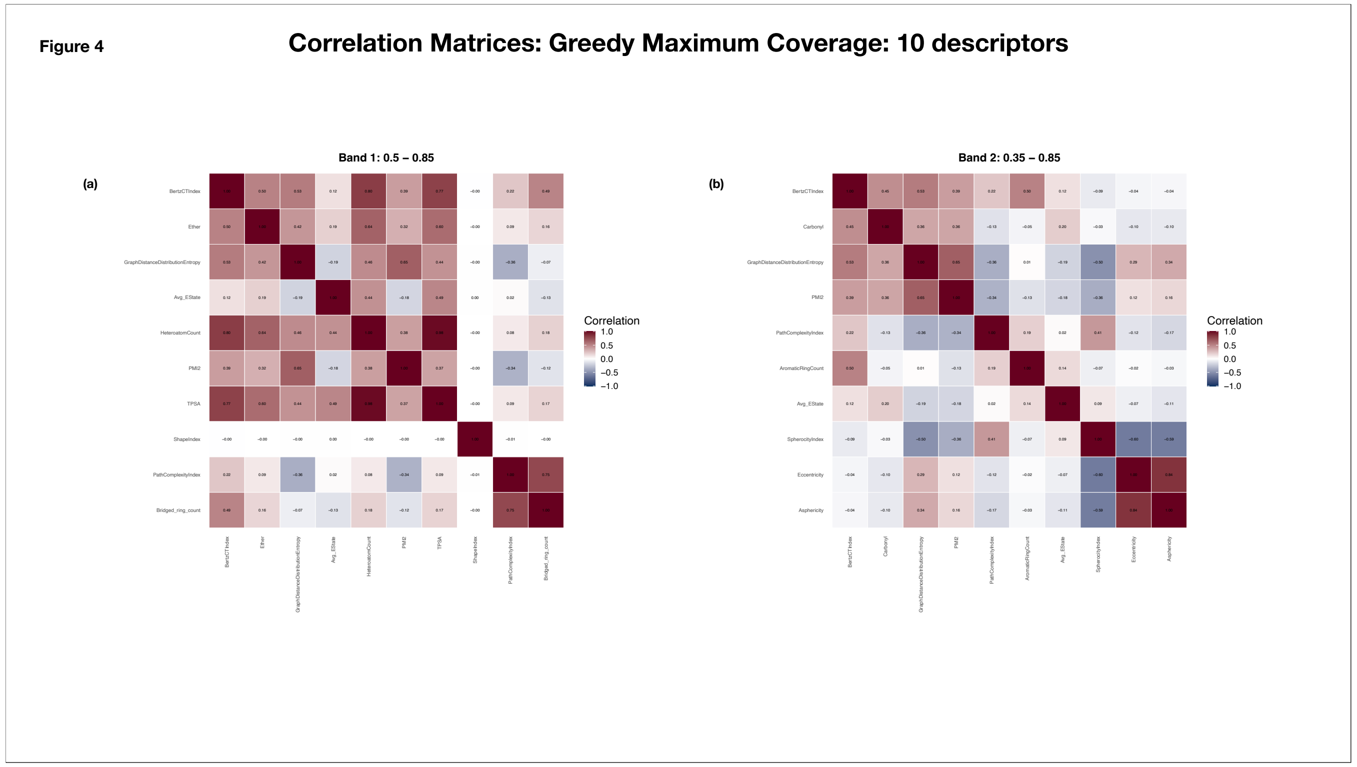


**Supplementary Figure 4: Correlation matrices generated using the Greedy Maximum Coverage method with 10 selected descriptors**

The x-axis and y-axis represent the selected molecular descriptors. Each cell denotes the pairwise Spearman correlation coefficient between descriptor pairs, with coefficient values annotated within the cells. Color intensity ranges from dark blue (−1, strong negative correlation) to dark red (+1, strong positive correlation), with white indicating no correlation (0).

(a) Band 1 (0.5 - 0.85)

(b) Band 2 (0.35 - 0.85)

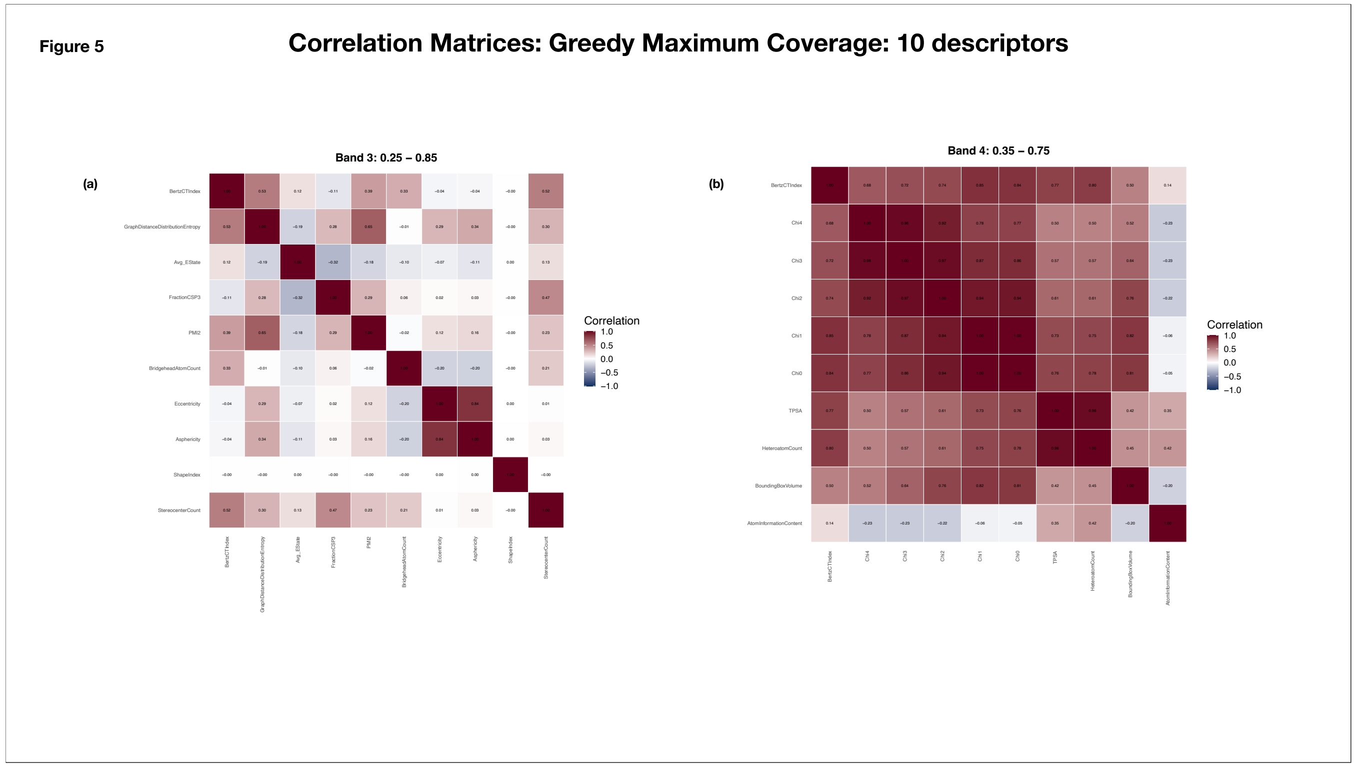


**Supplementary Figure 5: Correlation matrices generated using the Greedy Maximum Coverage method with 10 selected descriptors**

The x-axis and y-axis represent the selected molecular descriptors. Each cell denotes the pairwise Spearman correlation coefficient between descriptor pairs, with coefficient values annotated within the cells. Color intensity ranges from dark blue (−1, strong negative correlation) to dark red (+1, strong positive correlation), with white indicating no correlation (0).

(a) Band 3 (0.25 - 0.85)

(b) Band 4 (0.35 - 0.75)

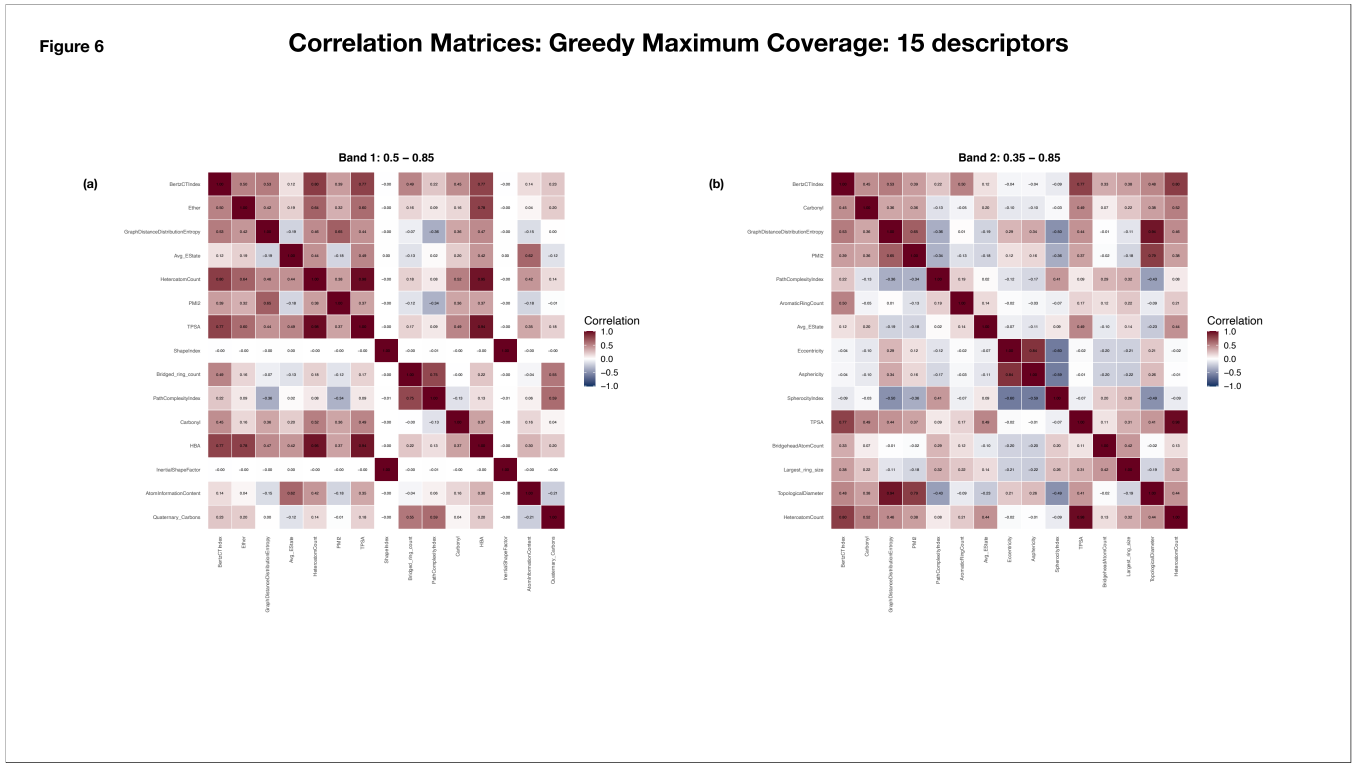


**Supplementary Figure 6: Correlation matrices generated using the Greedy Maximum Coverage method with 15 selected descriptors**

The x-axis and y-axis represent the selected molecular descriptors. Each cell denotes the pairwise Spearman correlation coefficient between descriptor pairs, with coefficient values annotated within the cells. Color intensity ranges from dark blue (−1, strong negative correlation) to dark red (+1, strong positive correlation), with white indicating no correlation (0).

(a) Band 1 (0.5 - 0.85)

(b) Band 2 (0.35 - 0.85)

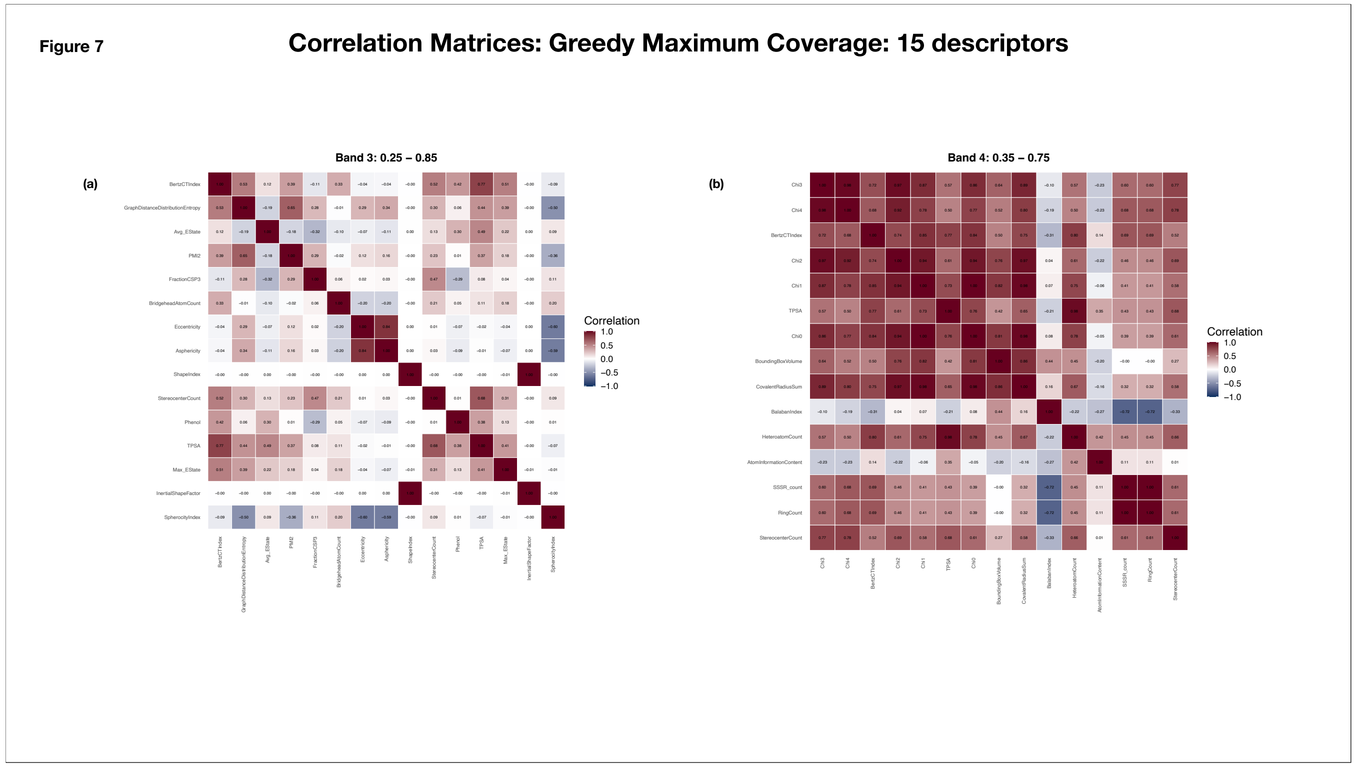


**Supplementary Figure 7: Correlation matrices generated using the Greedy Maximum Coverage method with 10 selected descriptors**

The x-axis and y-axis represent the selected molecular descriptors. Each cell denotes the pairwise Spearman correlation coefficient between descriptor pairs, with coefficient values annotated within the cells. Color intensity ranges from dark blue (−1, strong negative correlation) to dark red (+1, strong positive correlation), with white indicating no correlation (0).

(a) Band 3 (0.25 - 0.85)

(b) Band 4 (0.35 - 0.75)

Figure 8

#### Correlation Matrices: Greedy Maximum Coverage: 20 descriptors

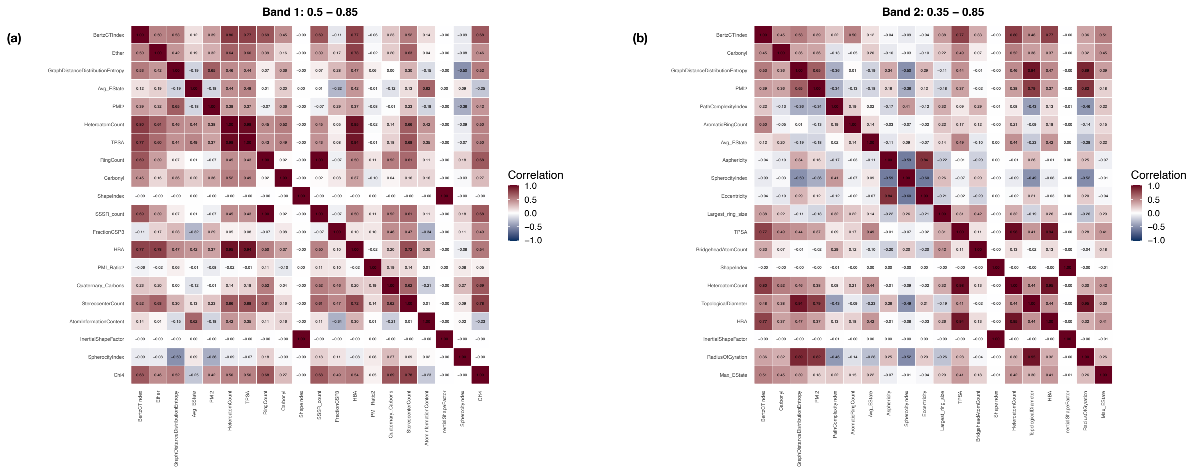

**Supplementary Figure 8: Correlation matrices generated using the Greedy Maximum Coverage method with 20 selected descriptors**
The x-axis and y-axis represent the selected molecular descriptors. Each cell denotes the pairwise Spearman correlation coefficient between descriptor pairs, with coefficient values annotated within the cells. Color intensity ranges from dark blue (−1, strong negative correlation) to dark red (+1, strong positive correlation), with white indicating no correlation (0).
(a) Band 1 (0.5 - 0.85)
(b) Band 2 (0.35 - 0.85)

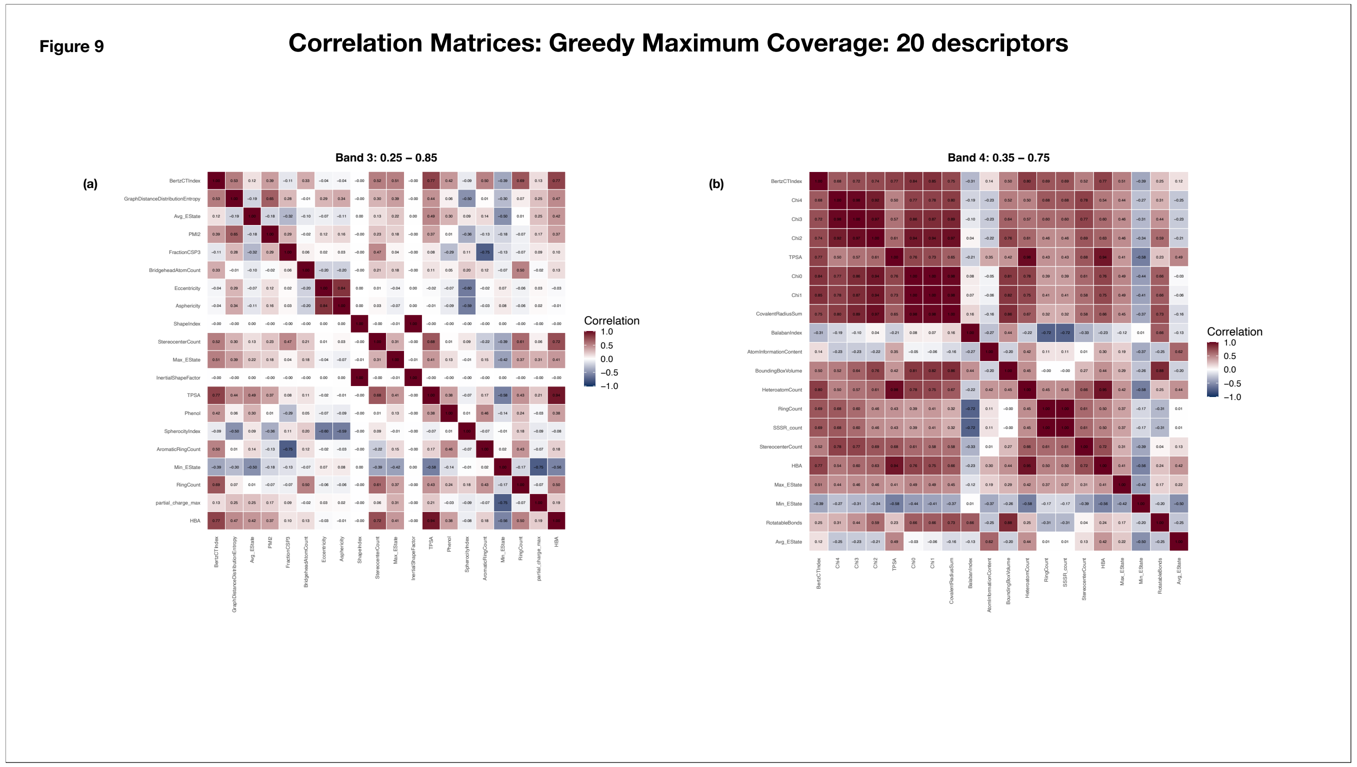


**Supplementary Figure 9: Correlation matrices generated using the Greedy Maximum Coverage method with 20 selected descriptors**

The x-axis and y-axis represent the selected molecular descriptors. Each cell denotes the pairwise Spearman correlation coefficient between descriptor pairs, with coefficient values annotated within the cells. Color intensity ranges from dark blue (−1, strong negative correlation) to dark red (+1, strong positive correlation), with white indicating no correlation (0).

(a) Band 3 (0.25 - 0.85)

(b) Band 4 (0.35 - 0.75)

Figure 10

#### Correlation Matrices: Greedy Maximum Coverage: 25 descriptors

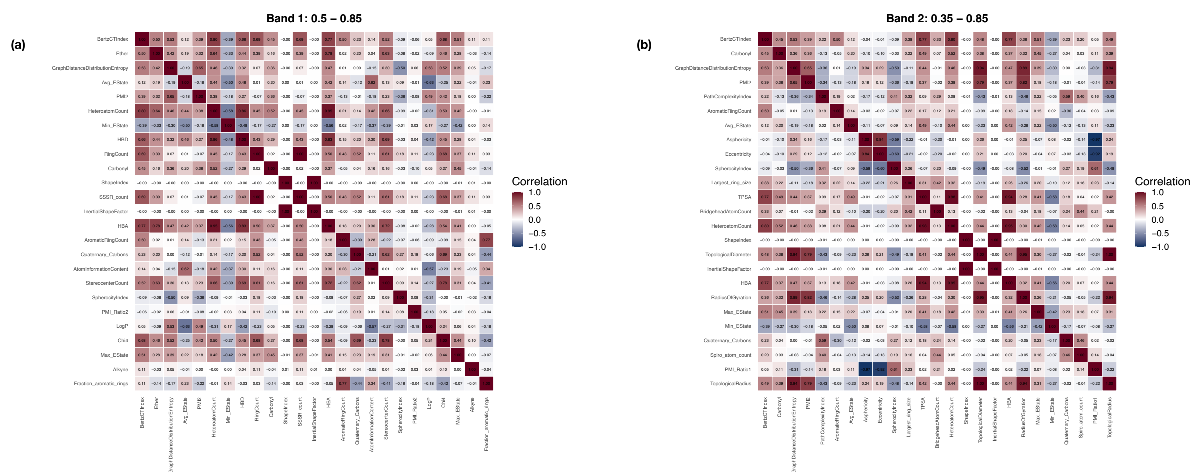

**Supplementary Figure 10: Correlation matrices generated using the Greedy Maximum Coverage method with 25 selected descriptors**

The x-axis and y-axis represent the selected molecular descriptors. Each cell denotes the pairwise Spearman correlation coefficient between descriptor pairs, with coefficient values annotated within the cells. Color intensity ranges from dark blue (−1, strong negative correlation) to dark red (+1, strong positive correlation), with white indicating no correlation (0).

(a) Band 1 (0.5 - 0.85)

(b) Band 2 (0.35 - 0.85)

Figure 11

#### Correlation Matrices: Greedy Maximum Coverage: 25 descriptors

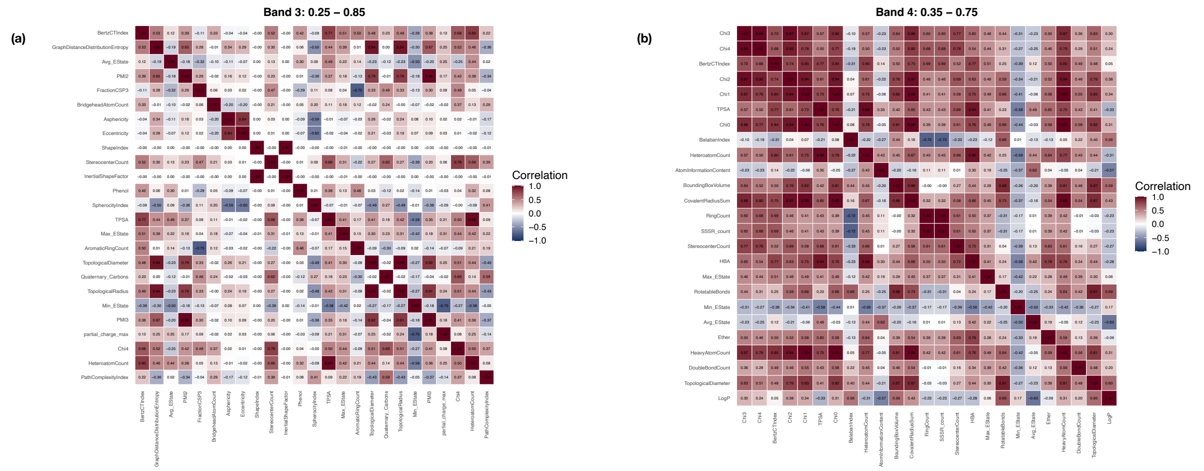


**Supplementary Figure 11: Correlation matrices generated using the Greedy Maximum Coverage method with 25 selected descriptors**

The x-axis and y-axis represent the selected molecular descriptors. Each cell denotes the pairwise Spearman correlation coefficient between descriptor pairs, with coefficient values annotated within the cells. Color intensity ranges from dark blue (−1, strong negative correlation) to dark red (+1, strong positive correlation), with white indicating no correlation (0).

(a) Band 3 (0.25 - 0.85)

(b) Band 4 (0.35 - 0.75)

**Figure 12**

#### Correlation Matrices: Hierarchical Clustering - Maximum Average Correlation: 10 descriptors

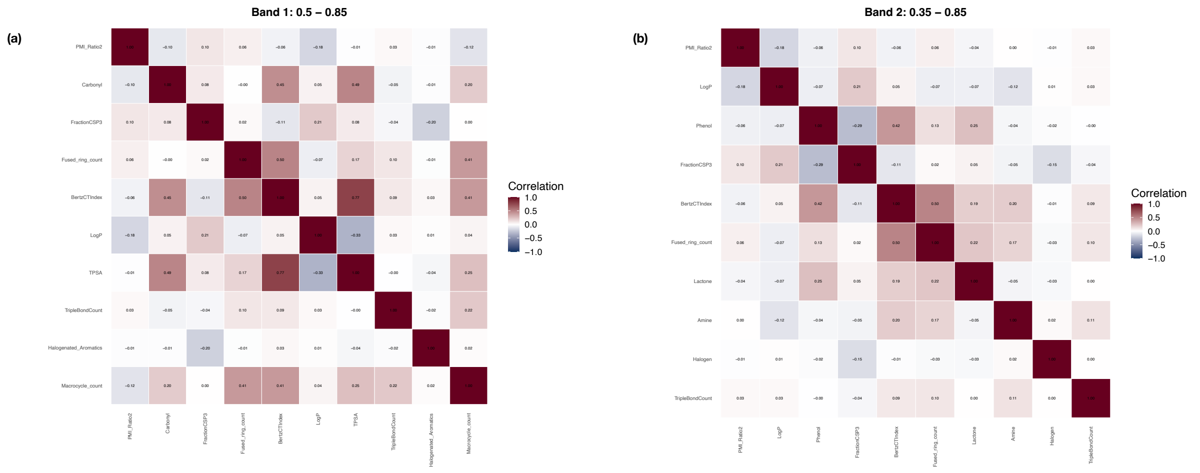


**Supplementary Figure 12: Correlation matrices generated using the Hierarchical Clustering (Maximum Average Correlation) method with 10 selected descriptors**

The x-axis and y-axis represent the selected molecular descriptors. Each cell denotes the pairwise Spearman correlation coefficient between descriptor pairs, with coefficient values annotated within the cells. Color intensity ranges from dark blue (−1, strong negative correlation) to dark red (+1, strong positive correlation), with white indicating no correlation (0).

(a) Band 1 (0.5 - 0.85)

(b) Band 2 (0.35 - 0.85)

**Figure 13** **Correlation Matrices: Hierarchical Clustering - Maximum Average Correlation: 10 descriptors**

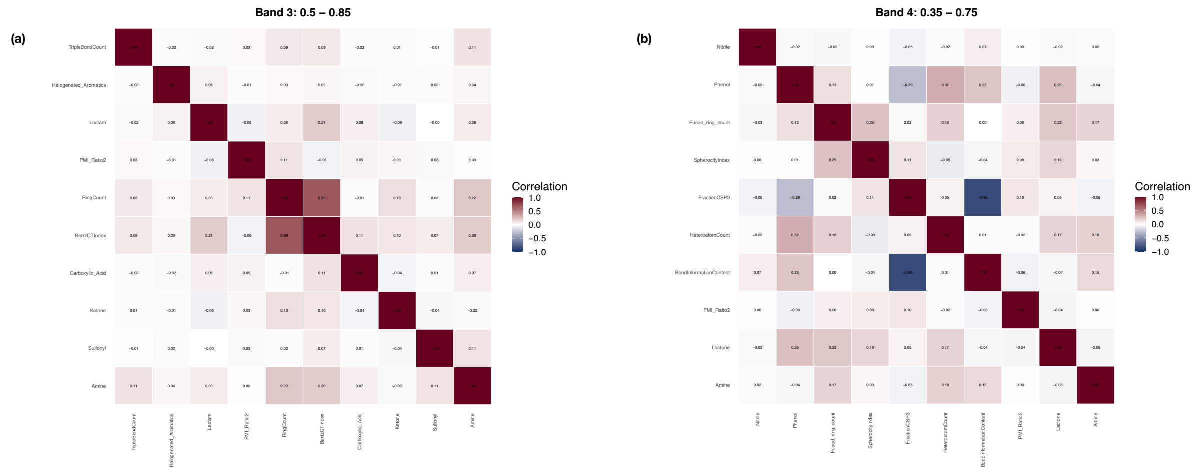


**Supplementary Figure 13: Correlation matrices generated using the Hierarchical Clustering (Maximum Average Correlation) method with 10 selected descriptors**

The x-axis and y-axis represent the selected molecular descriptors. Each cell denotes the pairwise Spearman correlation coefficient between descriptor pairs, with coefficient values annotated within the cells. Color intensity ranges from dark blue (−1, strong negative correlation) to dark red (+1, strong positive correlation), with white indicating no correlation (0).

(a) Band 3 (0.25 - 0.85)

(b) Band 4 (0.35 - 0.75)

Figure 14

#### Correlation Matrices: Hierarchical Clustering - Maximum Average Correlation: 15 descriptors

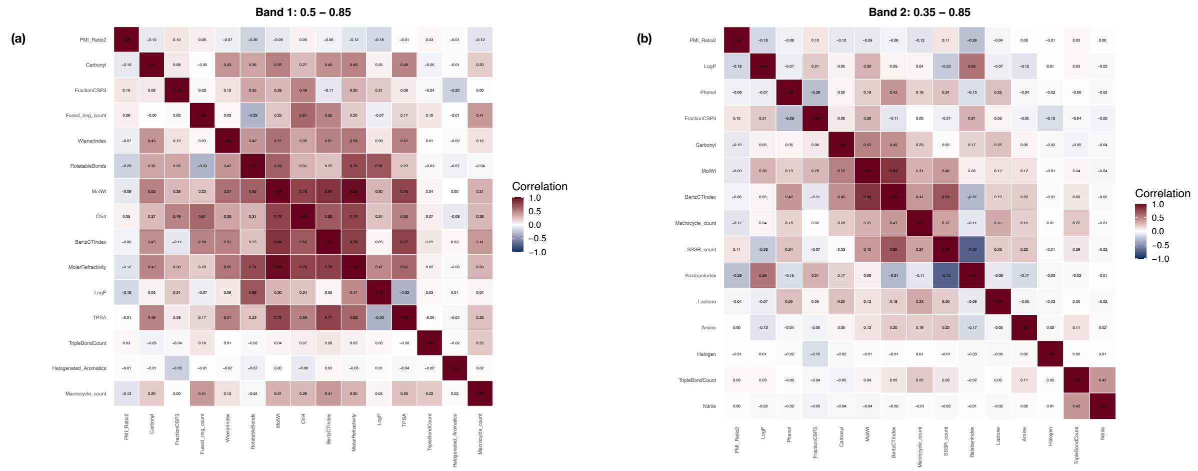

**Supplementary Figure 14: Correlation matrices generated using the Hierarchical Clustering (Maximum Average Correlation) method with 15 selected descriptors**
The x-axis and y-axis represent the selected molecular descriptors. Each cell denotes the pairwise Spearman correlation coefficient between descriptor pairs, with coefficient values annotated within the cells. Color intensity ranges from dark blue (−1, strong negative correlation) to dark red (+1, strong positive correlation), with white indicating no correlation (0).
(a) Band 1 (0.5 - 0.85)
(b) Band 2 (0.35 - 0.85)

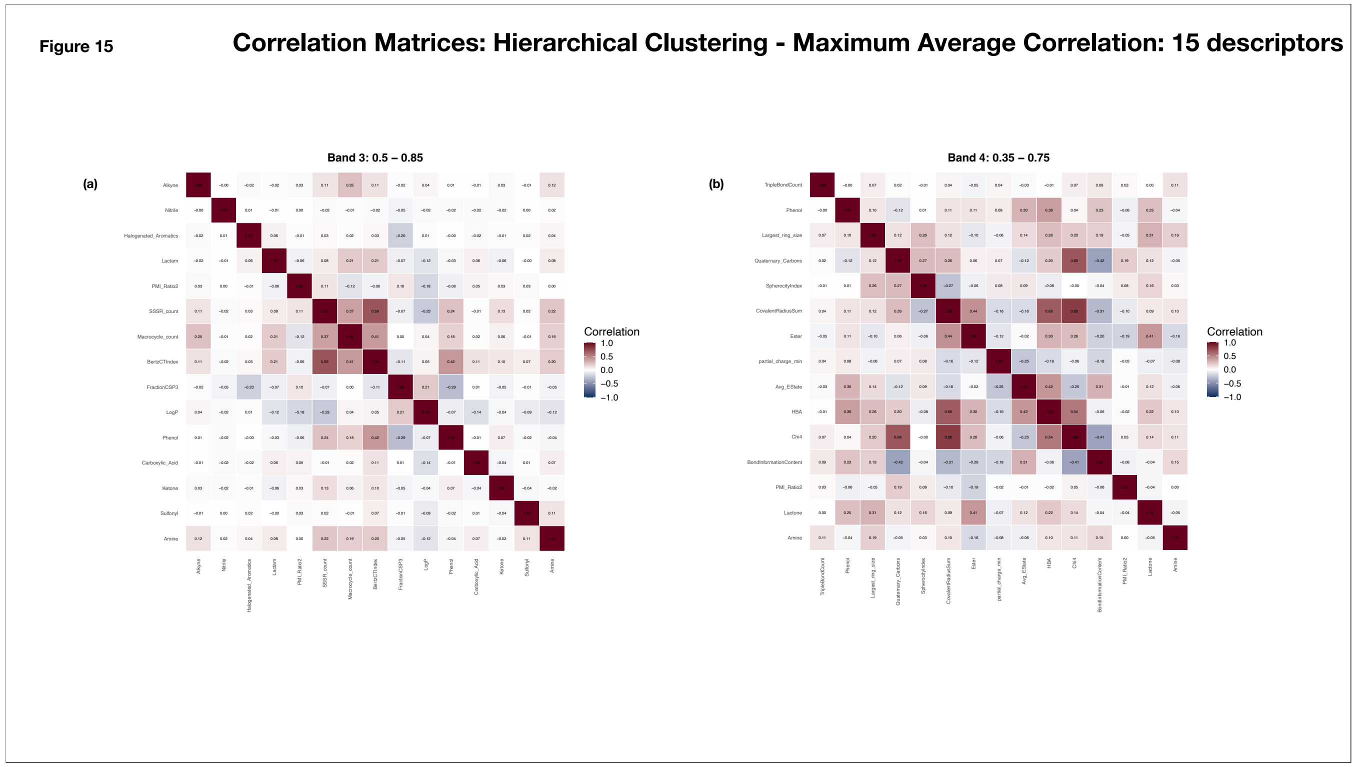


**Supplementary Figure 15: Correlation matrices generated using the Hierarchical Clustering (Maximum Average Correlation) method with 15 selected descriptors**

The x-axis and y-axis represent the selected molecular descriptors. Each cell denotes the pairwise Spearman correlation coefficient between descriptor pairs, with coefficient values annotated within the cells. Color intensity ranges from dark blue (−1, strong negative correlation) to dark red (+1, strong positive correlation), with white indicating no correlation (0).

(a) Band 3 (0.25 - 0.85)

(b) Band 4 (0.35 - 0.75)

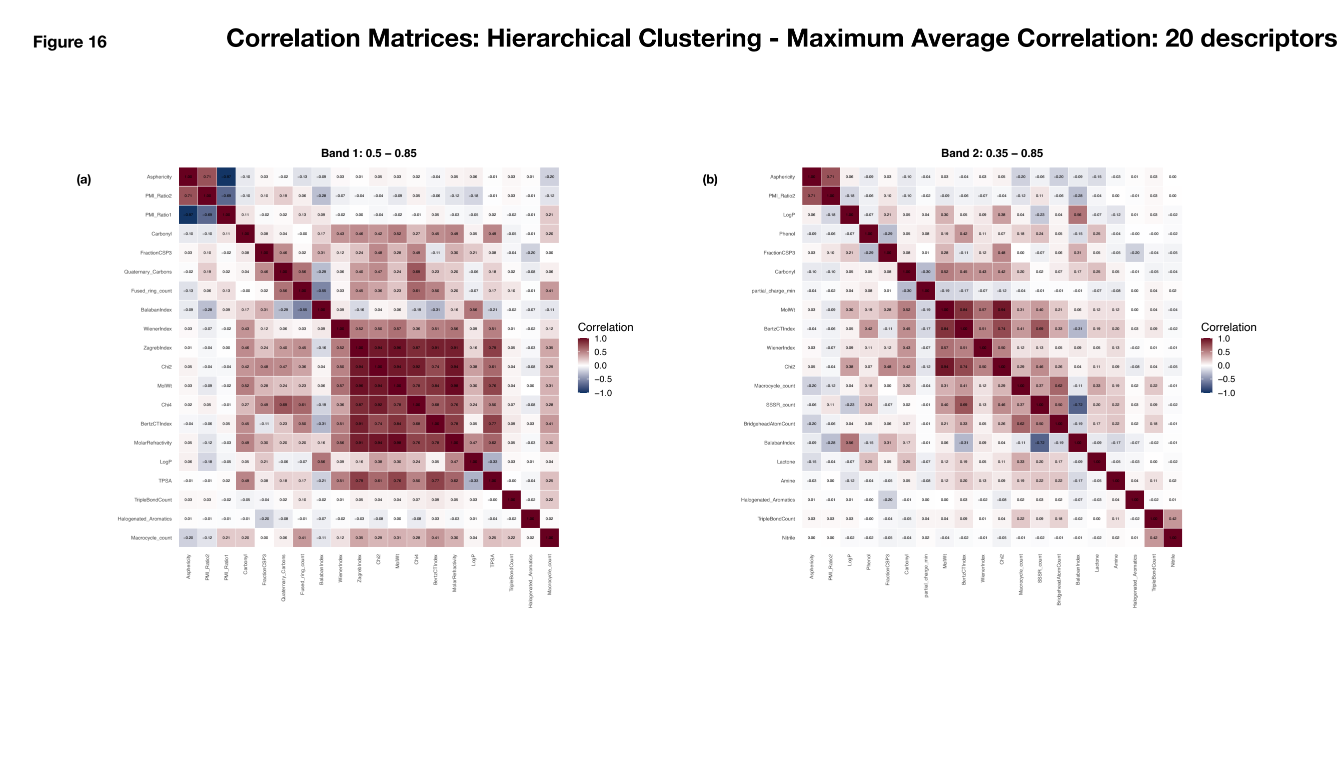


**Supplementary Figure 16: Correlation matrices generated using the Hierarchical Clustering (Maximum Average Correlation) method with 15 selected descriptors**

The x-axis and y-axis represent the selected molecular descriptors. Each cell denotes the pairwise Spearman correlation coefficient between descriptor pairs, with coefficient values annotated within the cells. Color intensity ranges from dark blue (−1, strong negative correlation) to dark red (+1, strong positive correlation), with white indicating no correlation (0).

(a) Band 1 (0.5 - 0.85)

(b) Band 2 (0.35 - 0.85)

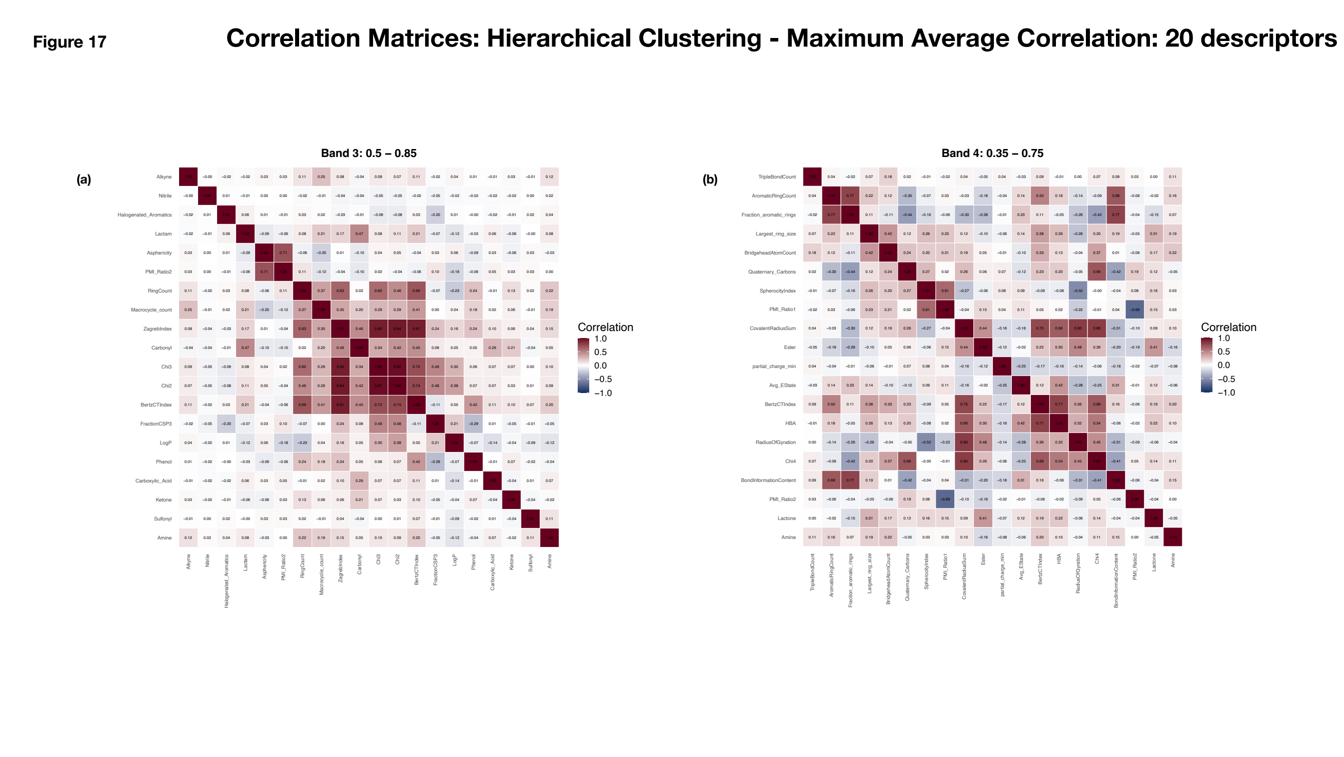


**Supplementary Figure 17: Correlation matrices generated using the Hierarchical Clustering (Maximum Average Correlation) method with 20 selected descriptors**

The x-axis and y-axis represent the selected molecular descriptors. Each cell denotes the pairwise Spearman correlation coefficient between descriptor pairs, with coefficient values annotated within the cells. Color intensity ranges from dark blue (−1, strong negative correlation) to dark red (+1, strong positive correlation), with white indicating no correlation (0).

(a) Band 3 (0.25 - 0.85)

(b) Band 4 (0.35 - 0.75)

Figure 18

#### Correlation Matrices: Hierarchical Clustering - Maximum Average Correlation: 25 descriptors

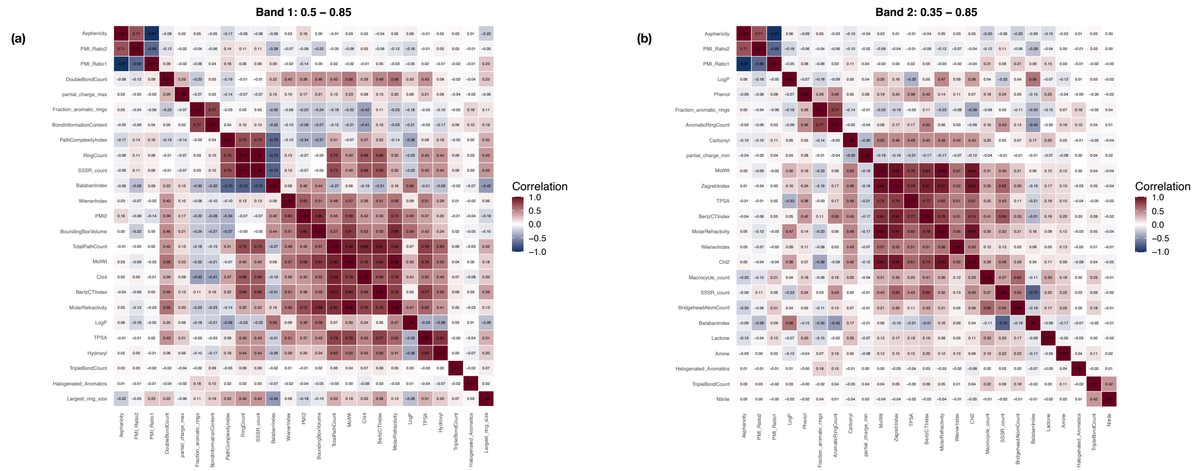

**Supplementary Figure 18: Correlation matrices generated using the Hierarchical Clustering (Maximum Average Correlation) method with 25 selected descriptors**

The x-axis and y-axis represent the selected molecular descriptors. Each cell denotes the pairwise Spearman correlation coefficient between descriptor pairs, with coefficient values annotated within the cells. Color intensity ranges from dark blue (–1, strong negative correlation) to dark red (+1, strong positive correlation), with white indicating no correlation (0).

(a) Band 1 (0.5 - 0.85)

(b) Band 2 (0.35 - 0.85)

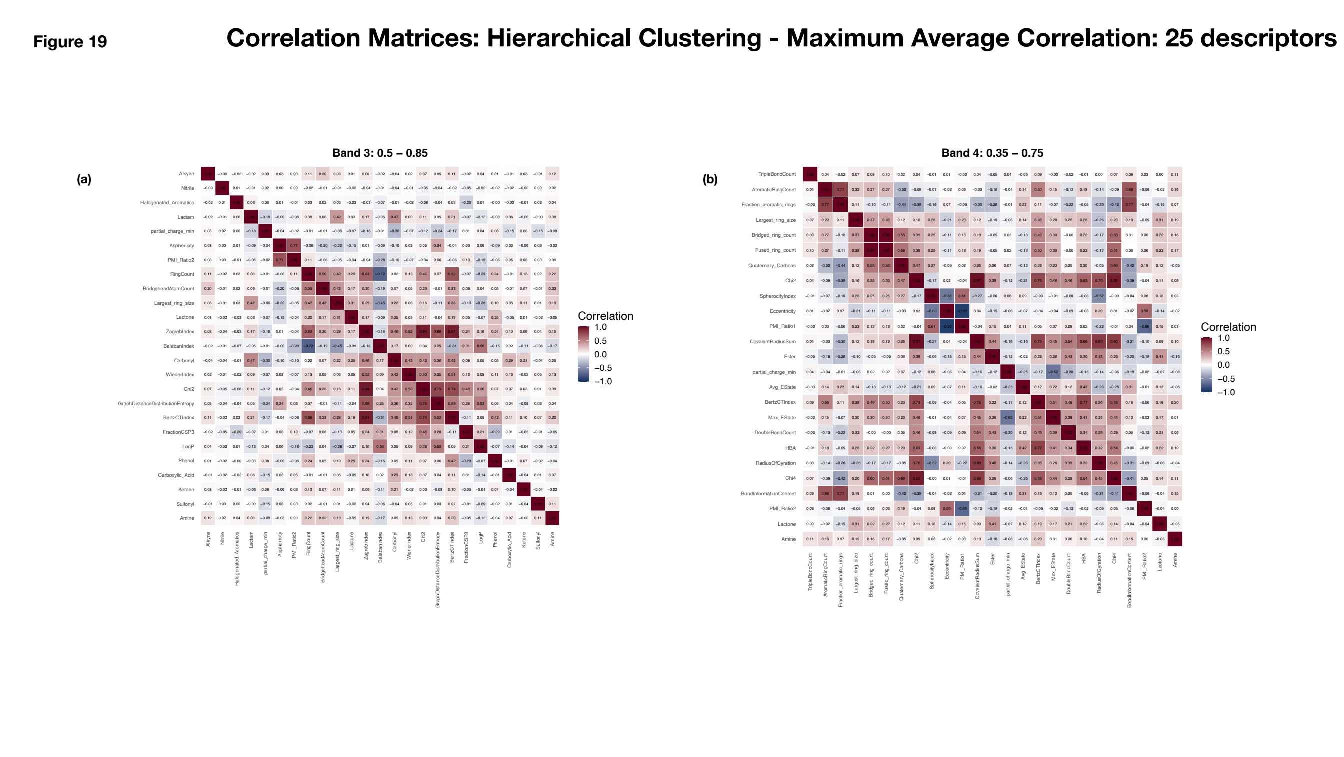


**Supplementary Figure 19: Correlation matrices generated using the Hierarchical Clustering (Maximum Average Correlation) method with 25 selected descriptors**

The x-axis and y-axis represent the selected molecular descriptors. Each cell denotes the pairwise Spearman correlation coefficient between descriptor pairs, with coefficient values annotated within the cells. Color intensity ranges from dark blue (−1, strong negative correlation) to dark red (+1, strong positive correlation), with white indicating no correlation (0).

(a) Band 3 (0.25 - 0.85)

(b) Band 4 (0.35 - 0.75)

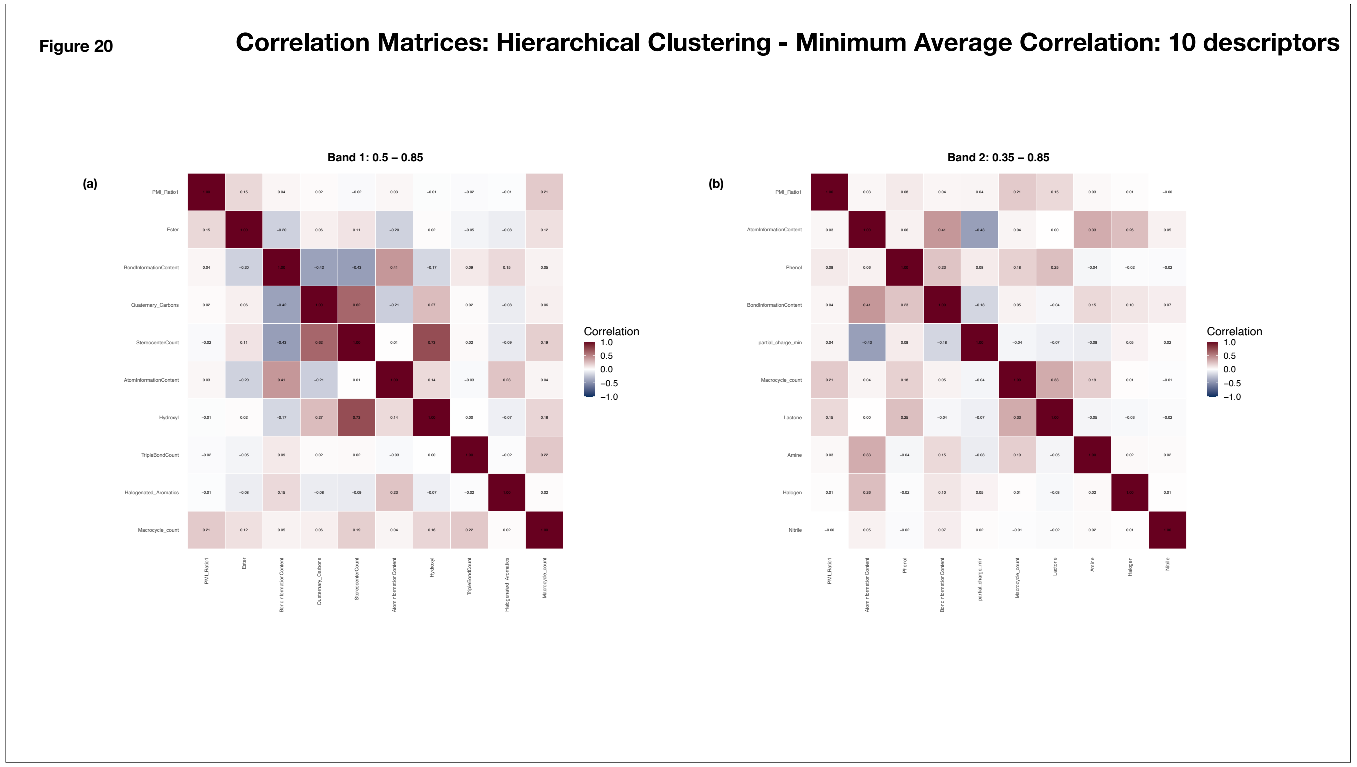

**Supplementary Figure 20: Correlation matrices generated using the Hierarchical Clustering (Minimum Average Correlation) method with 10 selected descriptors**
The x-axis and y-axis represent the selected molecular descriptors. Each cell denotes the pairwise Spearman correlation coefficient between descriptor pairs, with coefficient values annotated within the cells. Color intensity ranges from dark blue (–1, strong negative correlation) to dark red (+1, strong positive correlation), with white indicating no correlation (0).
(a) Band 1 (0.5 - 0.85)
(b) Band 2 (0.35 - 0.85)

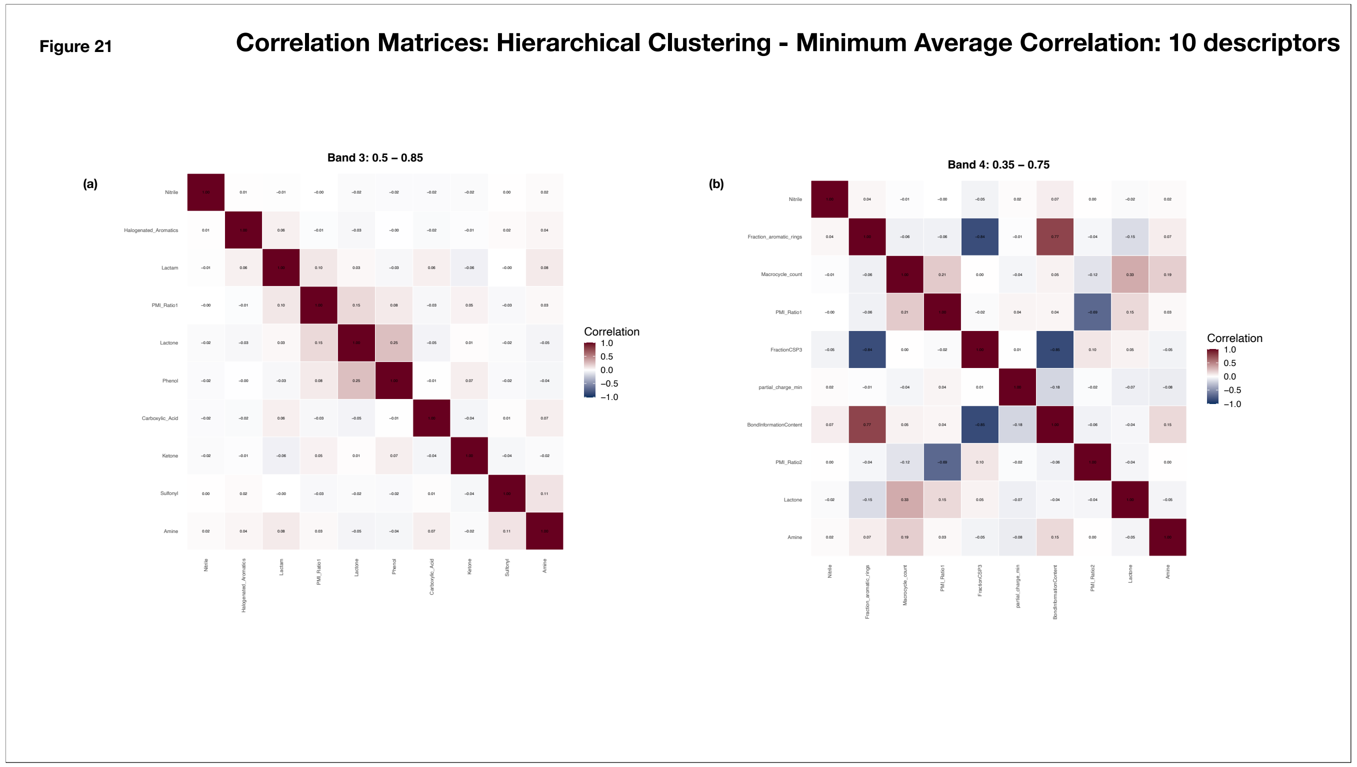


**Supplementary Figure 21: Correlation matrices generated using the Hierarchical Clustering (Minimum Average Correlation) method with 10 selected descriptors**

The x-axis and y-axis represent the selected molecular descriptors. Each cell denotes the pairwise Spearman correlation coefficient between descriptor pairs, with coefficient values annotated within the cells. Color intensity ranges from dark blue (−1, strong negative correlation) to dark red (+1, strong positive correlation), with white indicating no correlation (0).

(a) Band 3 (0.25 - 0.85)

(b) Band 4 (0.35 - 0.75)

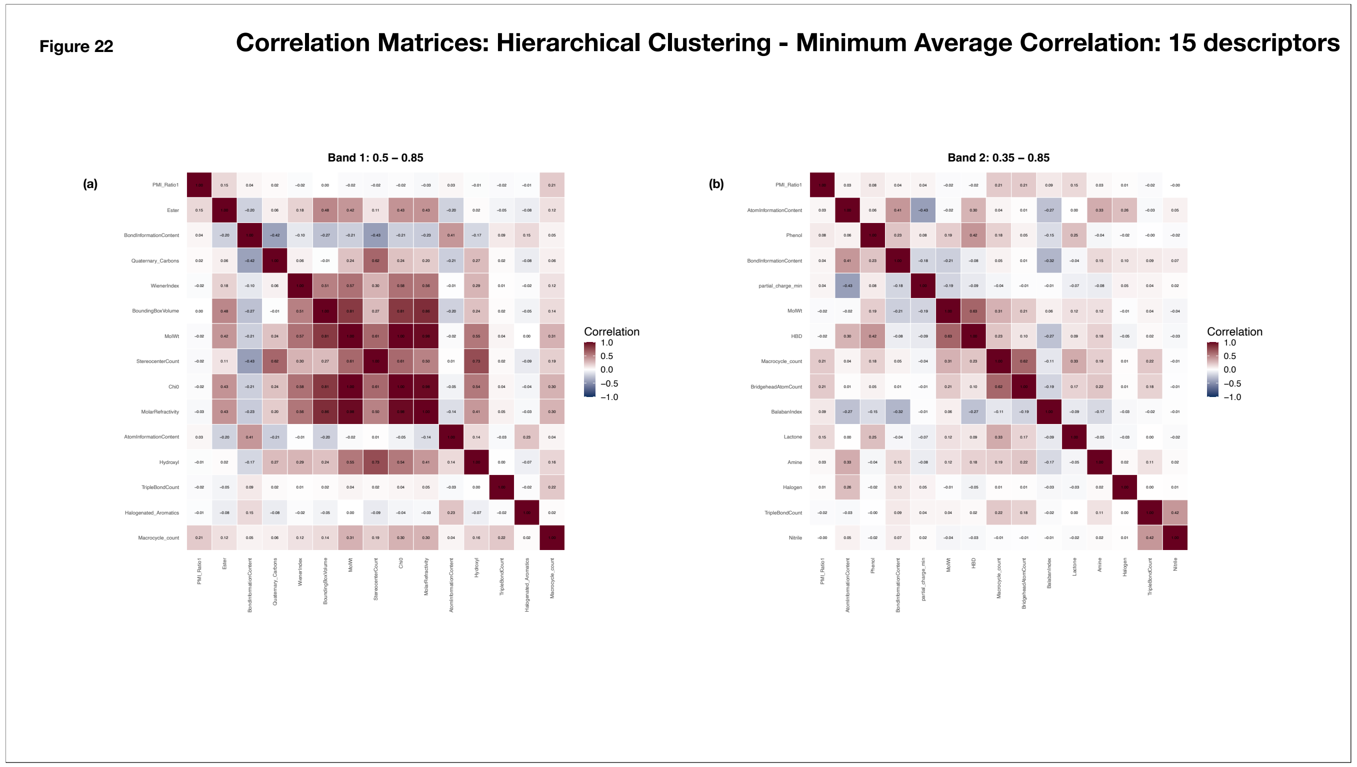


**Supplementary Figure 22: Correlation matrices generated using the Hierarchical Clustering (Minimum Average Correlation) method with 15 selected descriptors**

The x-axis and y-axis represent the selected molecular descriptors. Each cell denotes the pairwise Spearman correlation coefficient between descriptor pairs, with coefficient values annotated within the cells. Color intensity ranges from dark blue (−1, strong negative correlation) to dark red (+1, strong positive correlation), with white indicating no correlation (0).

(a) Band 1 (0.5 - 0.85)

(b) Band 2 (0.35 - 0.85)

**Figure 23** **Correlation Matrices: Hierarchical Clustering - Minimum Average Correlation: 15 descriptors**

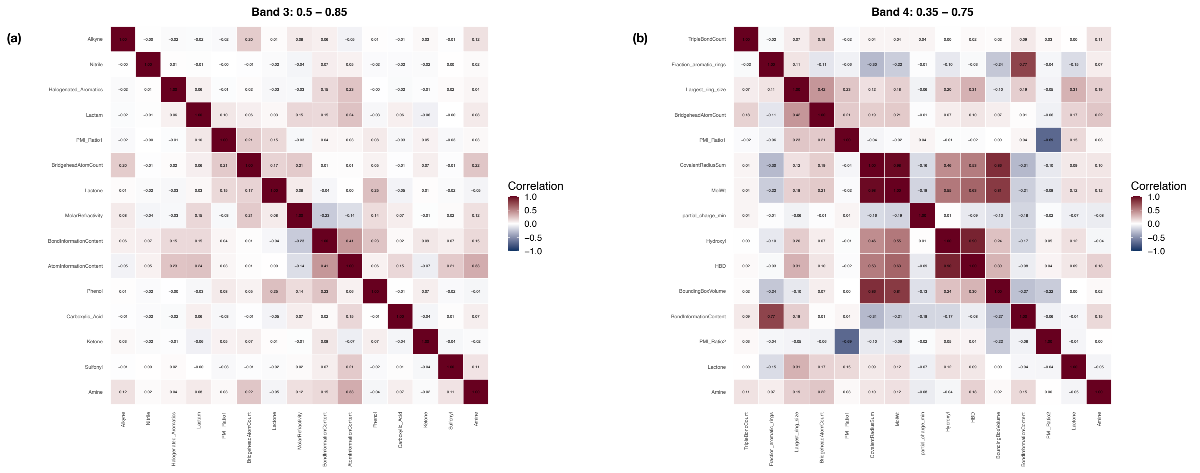


**Supplementary Figure 23: Correlation matrices generated using the Hierarchical Clustering (Minimum Average Correlation) method with 15 selected descriptors**

The x-axis and y-axis represent the selected molecular descriptors. Each cell denotes the pairwise Spearman correlation coefficient between descriptor pairs, with coefficient values annotated within the cells. Color intensity ranges from dark blue (−1, strong negative correlation) to dark red (+1, strong positive correlation), with white indicating no correlation (0).

(a) Band 3 (0.25 - 0.85)

(b) Band 4 (0.35 - 0.75)

Figure 24

#### Correlation Matrices: Hierarchical Clustering - Minimum Average Correlation: 20 descriptors

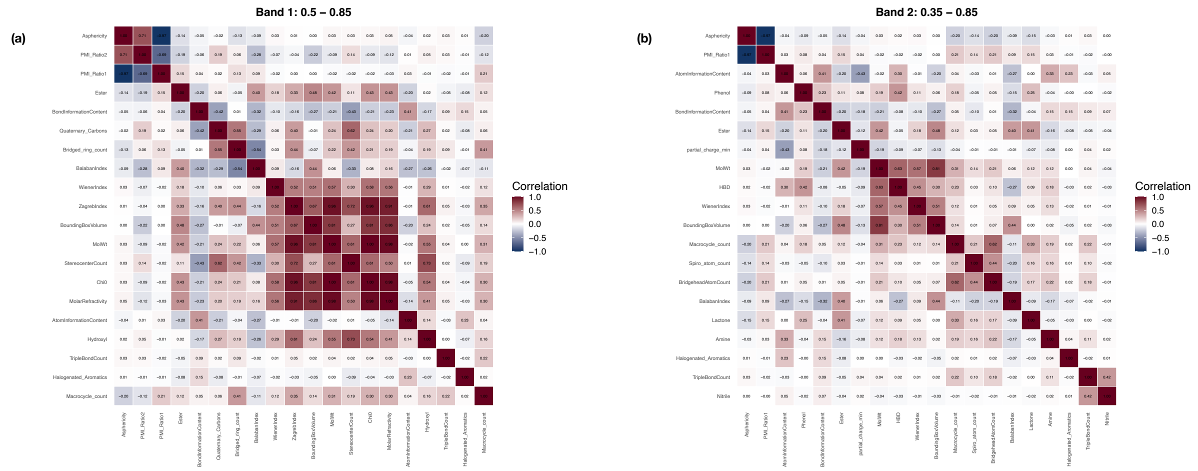


**Supplementary Figure 24: Correlation matrices generated using the Hierarchical Clustering (Minimum Average Correlation) method with 20 selected descriptors**

The x-axis and y-axis represent the selected molecular descriptors. Each cell denotes the pairwise Spearman correlation coefficient between descriptor pairs, with coefficient values annotated within the cells. Color intensity ranges from dark blue (−1, strong negative correlation) to dark red (+1, strong positive correlation), with white indicating no correlation (0).

(a) Band 1 (0.5 - 0.85)

(b) Band 2 (0.35 - 0.85)

Figure 25

#### Correlation Matrices: Hierarchical Clustering - Minimum Average Correlation: 20 descriptors

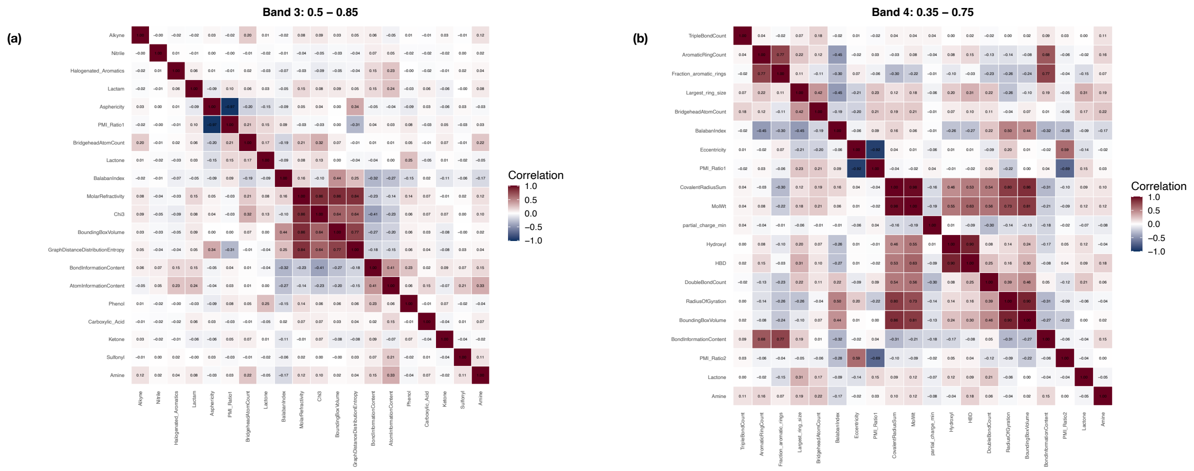


**Supplementary Figure 25: Correlation matrices generated using the Hierarchical Clustering (Minimum Average Correlation) method with 20 selected descriptors**

The x-axis and y-axis represent the selected molecular descriptors. Each cell denotes the pairwise Spearman correlation coefficient between descriptor pairs, with coefficient values annotated within the cells. Color intensity ranges from dark blue (−1, strong negative correlation) to dark red (+1, strong positive correlation), with white indicating no correlation (0).

(a) Band 3 (0.25 - 0.85)

(b) Band 4 (0.35 - 0.75)

**Figure 26** **Correlation Matrices: Hierarchical Clustering - Minimum Average Correlation: 25 descriptors**

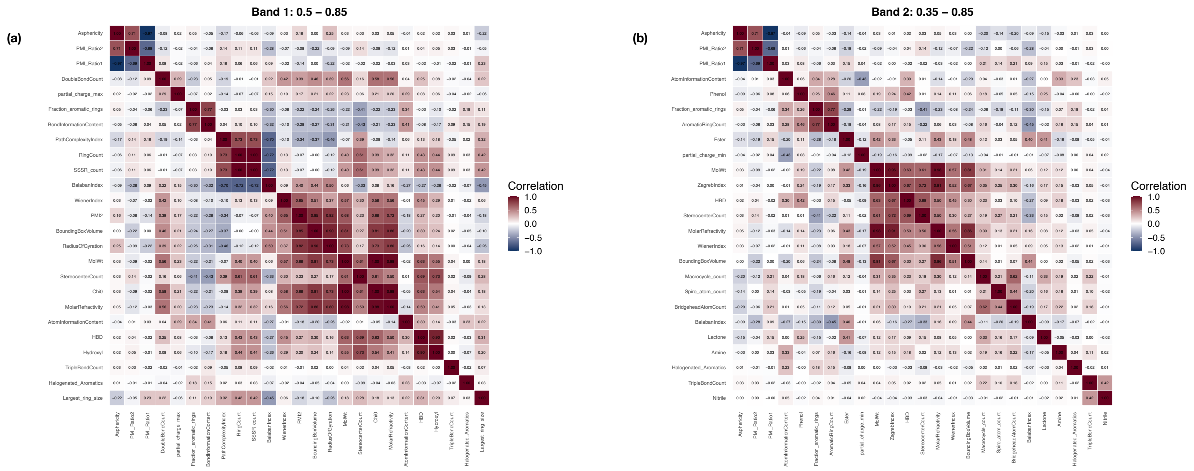


**Supplementary Figure 26: Correlation matrices generated using the Hierarchical Clustering (Minimum Average Correlation) method with 25 selected descriptors**

The x-axis and y-axis represent the selected molecular descriptors. Each cell denotes the pairwise Spearman correlation coefficient between descriptor pairs, with coefficient values annotated within the cells. Color intensity ranges from dark blue (−1, strong negative correlation) to dark red (+1, strong positive correlation), with white indicating no correlation (0).

(a) Band 1 (0.5 - 0.85)

(b) Band 2 (0.35 - 0.85)

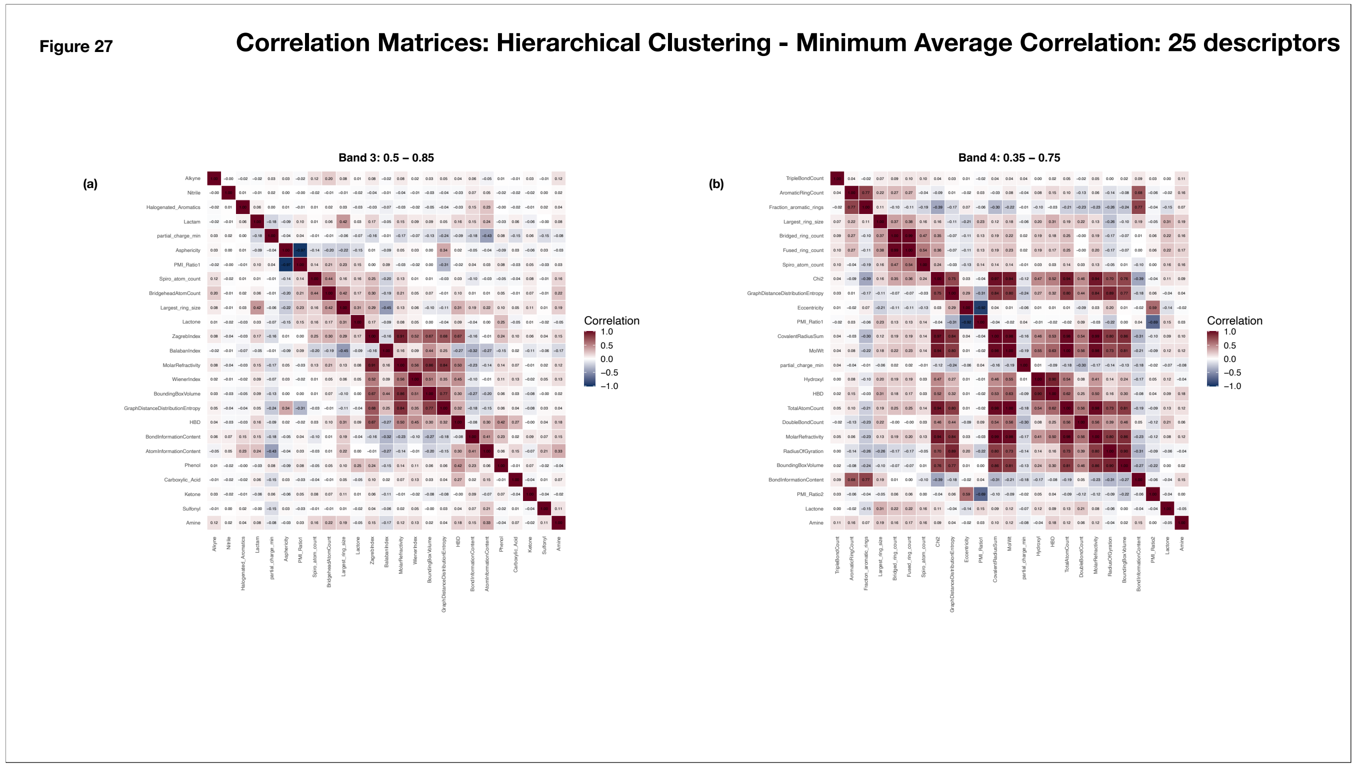


**Supplementary Figure 27: Correlation matrices generated using the Hierarchical Clustering (Minimum Average Correlation) method with 25 selected descriptors**

The x-axis and y-axis represent the selected molecular descriptors. Each cell denotes the pairwise Spearman correlation coefficient between descriptor pairs, with coefficient values annotated within the cells. Color intensity ranges from dark blue (−1, strong negative correlation) to dark red (+1, strong positive correlation), with white indicating no correlation (0).

(a) Band 3 (0.25 - 0.85)

(b) Band 4 (0.35 - 0.75)

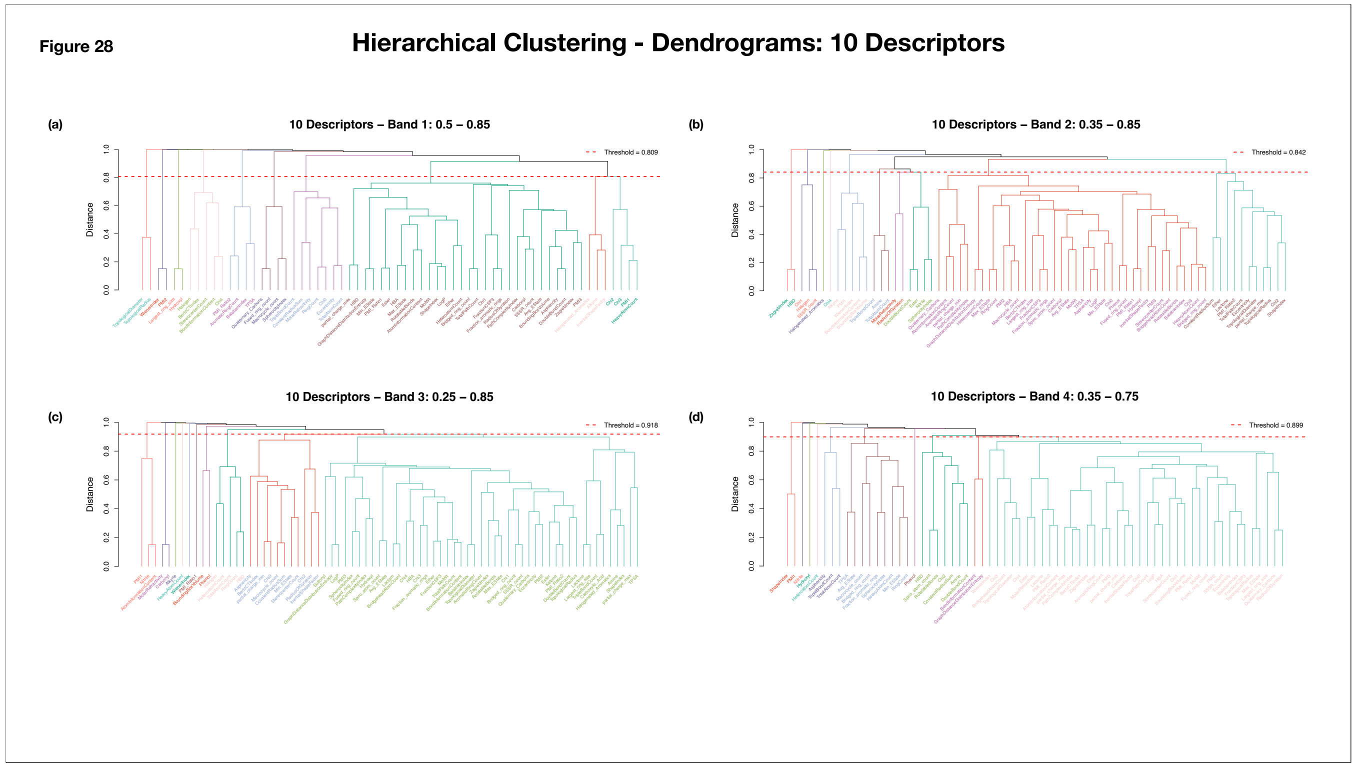


**Supplementary Figure 28: Dendrograms Representing Clusters formed through Hierarchical Clustering (10 Descriptors)**
The dendrograms depict descriptor clustering obtained through hierarchical clustering. The x-axis represents the molecular descriptors, and the y-axis represents the clustering distance. The red dashed line indicates the clustering threshold, with its value annotated in the upper-right corner. Descriptors belonging to the same cluster are shown in the same color.

**(a)** Band 1: 0.5 - 0.85
**(b)** Band 2: 0.35 - 0.85
**(c)** Band 3: 0.25 - 0.85
**(d)** Band 4: 0.35 - 0.75

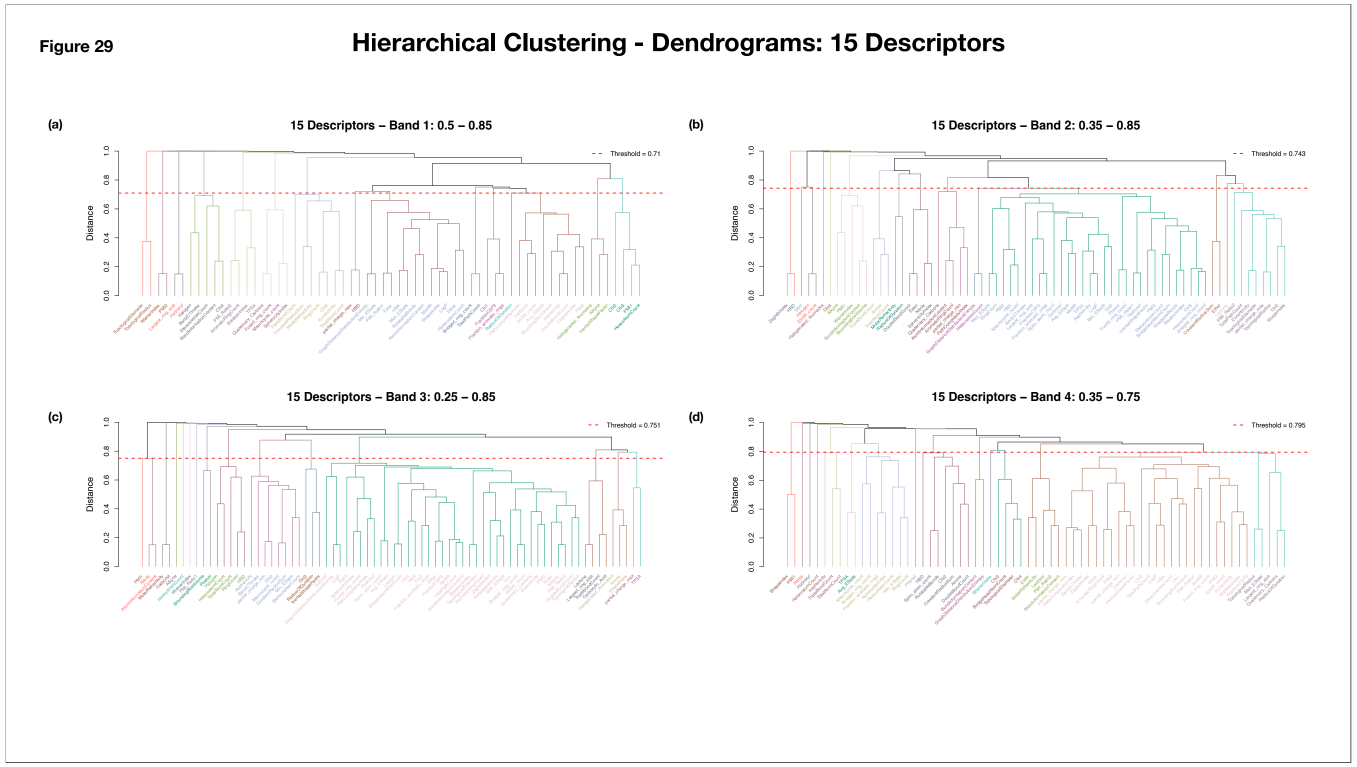


**Supplementary Figure 29: Dendrograms Representing Clusters formed through Hierarchical Clustering (15 Descriptors)**
The dendrograms depict descriptor clustering obtained through hierarchical clustering. The x-axis represents the molecular descriptors, and the y-axis represents the clustering distance. The red dashed line indicates the clustering threshold, with its value annotated in the upper-right corner. Descriptors belonging to the same cluster are shown in the same color.
**(a)** Band 1: 0.5 - 0.85
**(b)** Band 2: 0.35 - 0.85
**(c)** Band 3: 0.25 - 0.85
**(d)** Band 4: 0.35 - 0.75

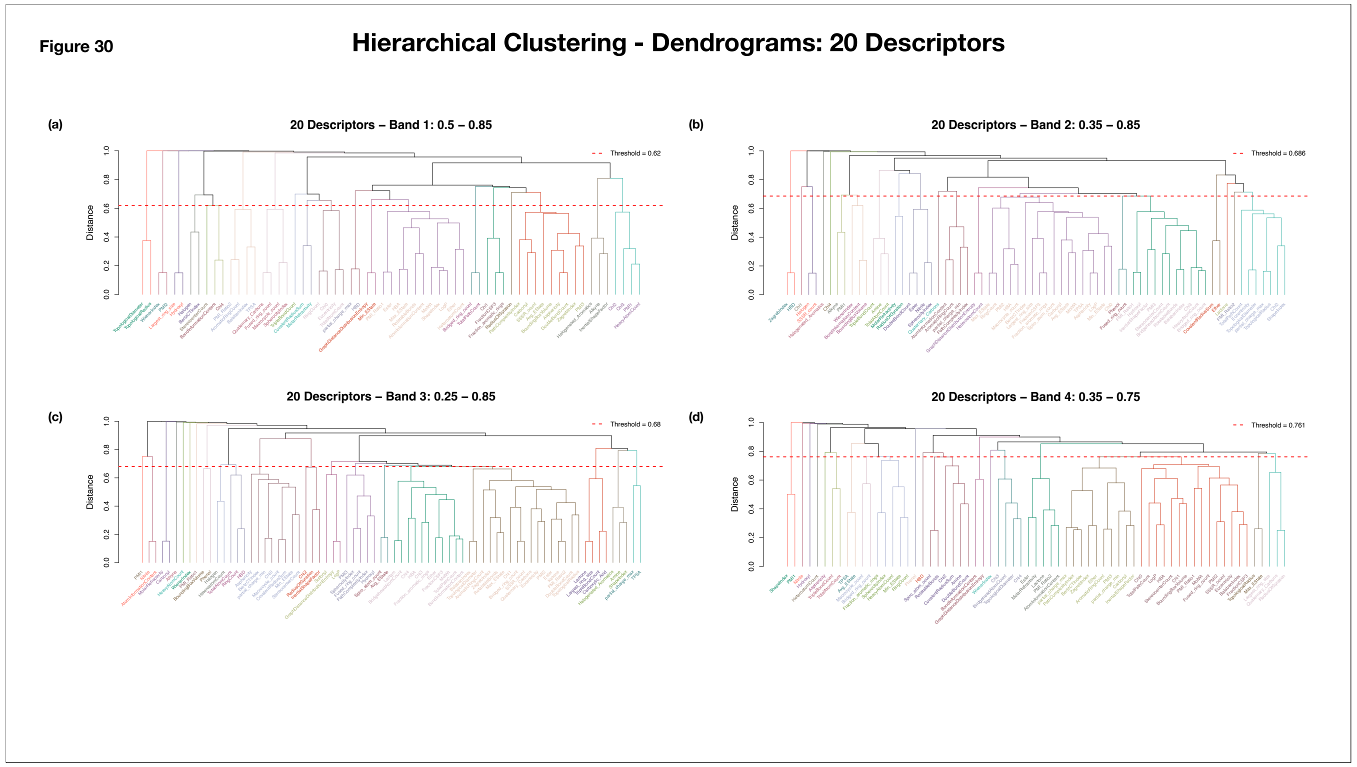


**Supplementary Figure 30: Dendrograms Representing Clusters formed through Hierarchical Clustering (20 Descriptors)**
The dendrograms depict descriptor clustering obtained through hierarchical clustering. The x-axis represents the molecular descriptors, and the y-axis represents the clustering distance. The red dashed line indicates the clustering threshold, with its value annotated in the upper-right corner. Descriptors belonging to the same cluster are shown in the same color.
**(a)** Band 1: 0.5 - 0.85
**(b)** Band 2: 0.35 - 0.85
**(c)** Band 3: 0.25 - 0.85
**(d)** Band 4: 0.35 - 0.75

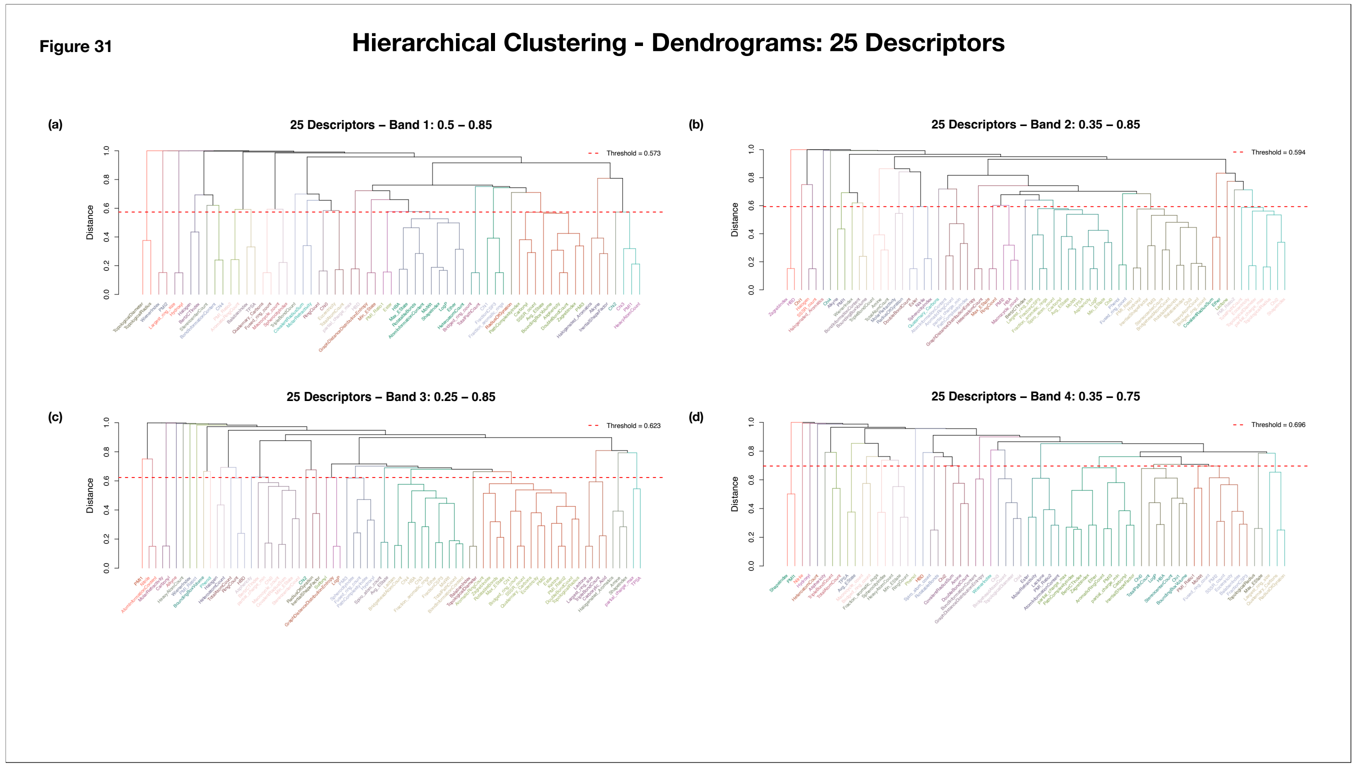


**Supplementary Figure 31: Dendrograms Representing Clusters formed through Hierarchical Clustering (25 Descriptors)**
The dendrograms depict descriptor clustering obtained through hierarchical clustering. The x-axis represents the molecular descriptors, and the y-axis represents the clustering distance. The red dashed line indicates the clustering threshold, with its value annotated in the upper-right corner. Descriptors belonging to the same cluster are shown in the same color.
**(a)** Band 1: 0.5 - 0.85
**(b)** Band 2: 0.35 - 0.85
**(c)** Band 3: 0.25 - 0.85
**(d)** Band 4: 0.35 - 0.75

**Figure 32**

#### Descriptor Distribution across the 48 configurations

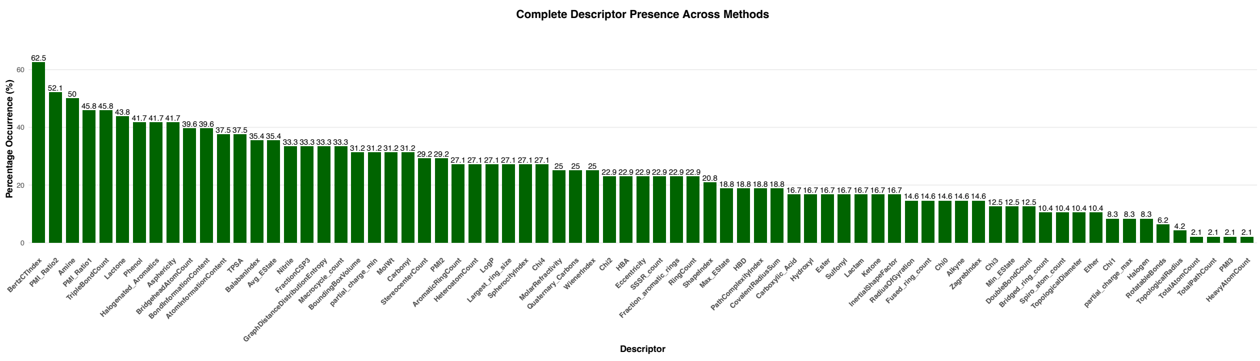


**Supplementary Figure 32. Descriptor Presence Across Selection Strategies, Descriptor Counts, and Correlation Threshold Ranges**
The bar plot summarizes the frequency of descriptor occurrence across all combinations of descriptor selection strategies, descriptor counts, and correlation threshold ranges. The x-axis represents the descriptor names, while the y-axis indicates the percentage of combinations in which each descriptor was selected. Percentage occurrence values are annotated above the corresponding bars.

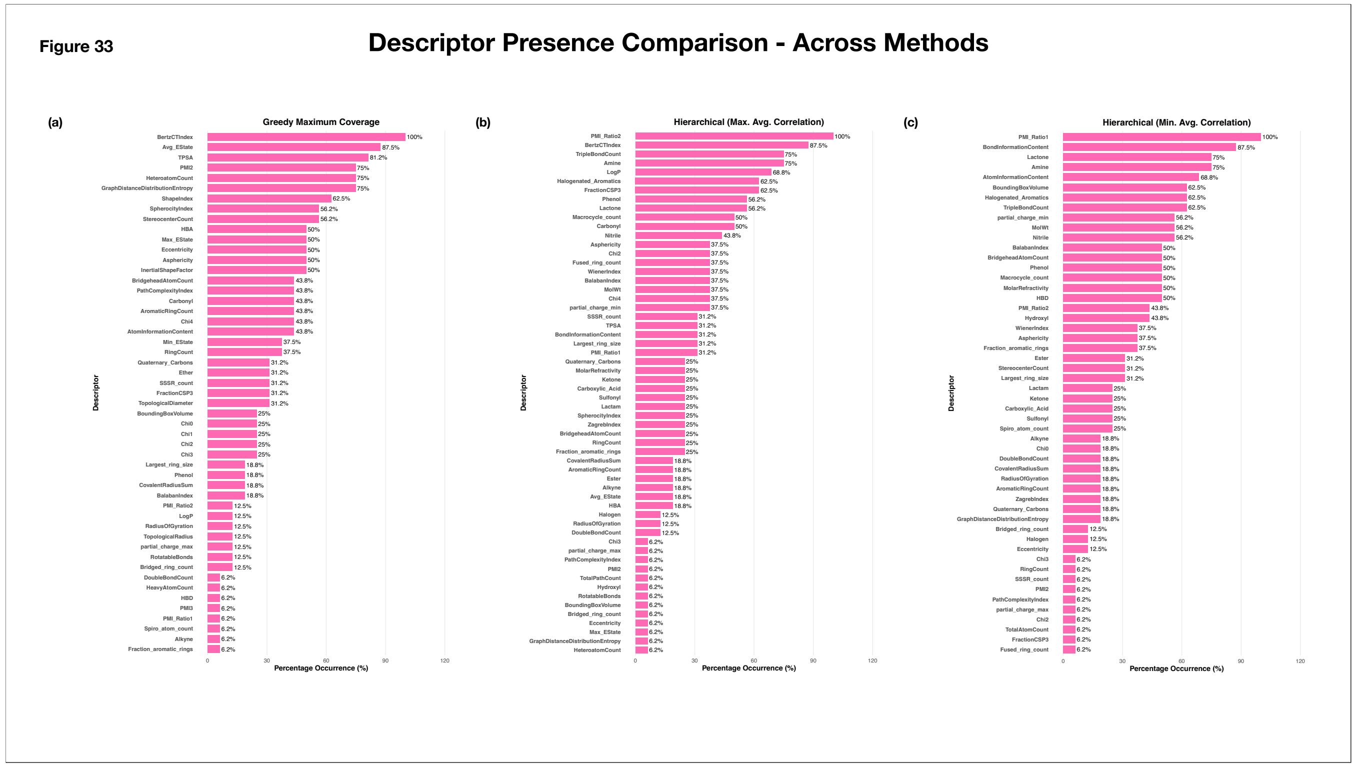


**Supplementary Figure 33. Descriptor Presence Across Selection Strategies**

The bar plot summarizes the frequency of descriptor occurrence across all combinations of descriptor counts and correlation threshold ranges for individual selection strategies. The x-axis represents the percentage of combinations in which each descriptor was selected while the y-axis indicates the descriptor names. Percentage occurrence values are annotated towards the right of the corresponding bars.

(a) Greedy maximum coverage

(b) Hierarchical clustering (maximum average correlation)

(c) Hierarchical clustering (minimum average correlation)

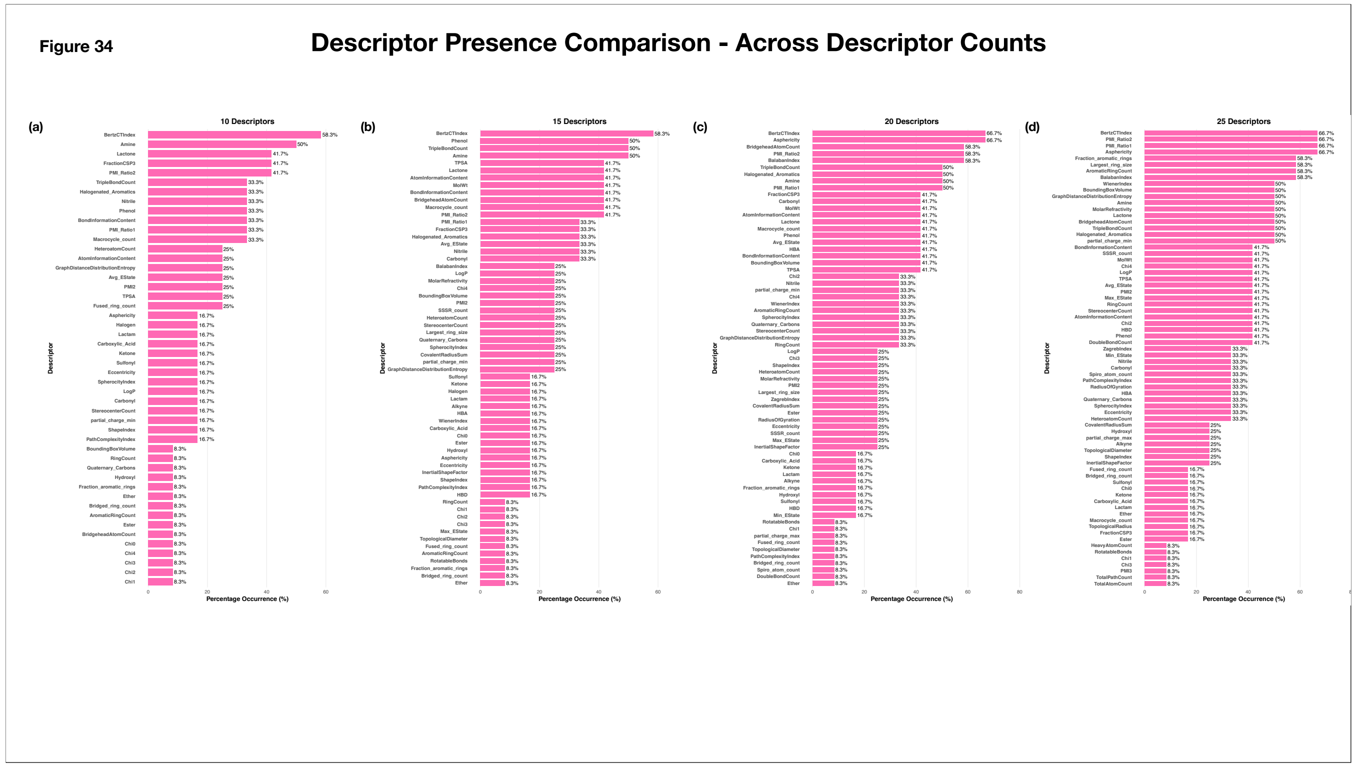


**Supplementary Figure 34. Descriptor Presence Across Descriptor Counts**

The bar plot summarizes the frequency of descriptor occurrence across all combinations of descriptor selection strategies and correlation threshold ranges for individual descriptor counts. The x-axis represents the percentage of combinations in which each descriptor was selected while the y-axis indicates the descriptor names. Percentage occurrence values are annotated towards the right of the corresponding bars.

(a) 10 descriptors
(b) 15 descriptors
(c) 20 descriptors
(d) 25 descriptors

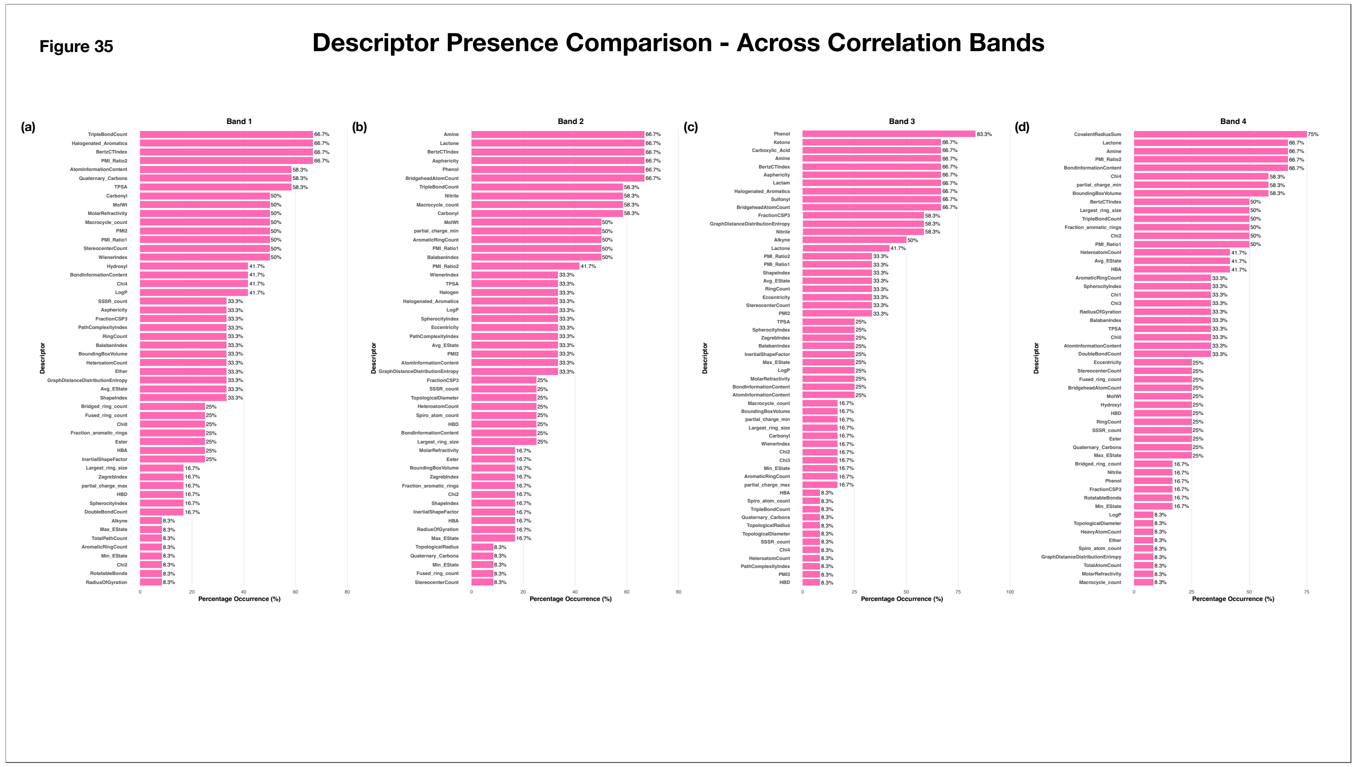


**Supplementary Figure 35. Descriptor Presence Across Correlation Threshold Ranges**

The bar plot summarizes the frequency of descriptor occurrence across all combinations of descriptor selection strategies and descriptor count for individual correlation threshold ranges. The x-axis represents the percentage of combinations in which each descriptor was selected while the y-axis indicates the descriptor names. Percentage occurrence values are annotated towards the right of the corresponding bars.

(a) Band 1 (0.5 - 0.85)
(b) Band 2 (0.35 - 0.85)
(c) Band 3 (0.25 - 0.85)
(d) Band 4 (0.35 - 0.75)

**Figure 36**

#### Descriptor Type Presence - Global Analysis

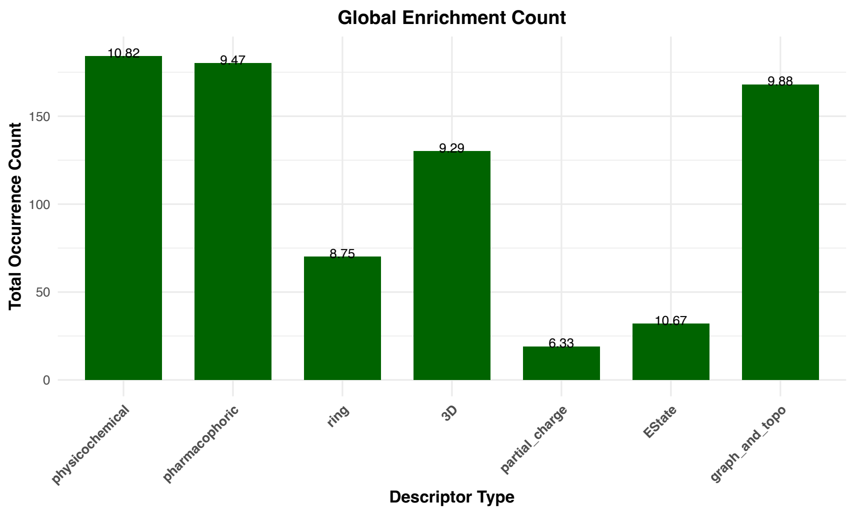


**Supplementary Figure 36. Global Descriptor Type Enrichment Across Descriptor Selection Strategies, Correlation Threshold Ranges, and Descriptor Counts**
The bar plot summarizes the normalized occurrence of descriptor types across all combinations of descriptor selection strategies, correlation threshold ranges, and descriptor counts. The x-axis represents the descriptor type, while the y-axis denotes the normalized occurrence frequency. Values annotated above each bar indicate the corresponding occurrence of each descriptor type.

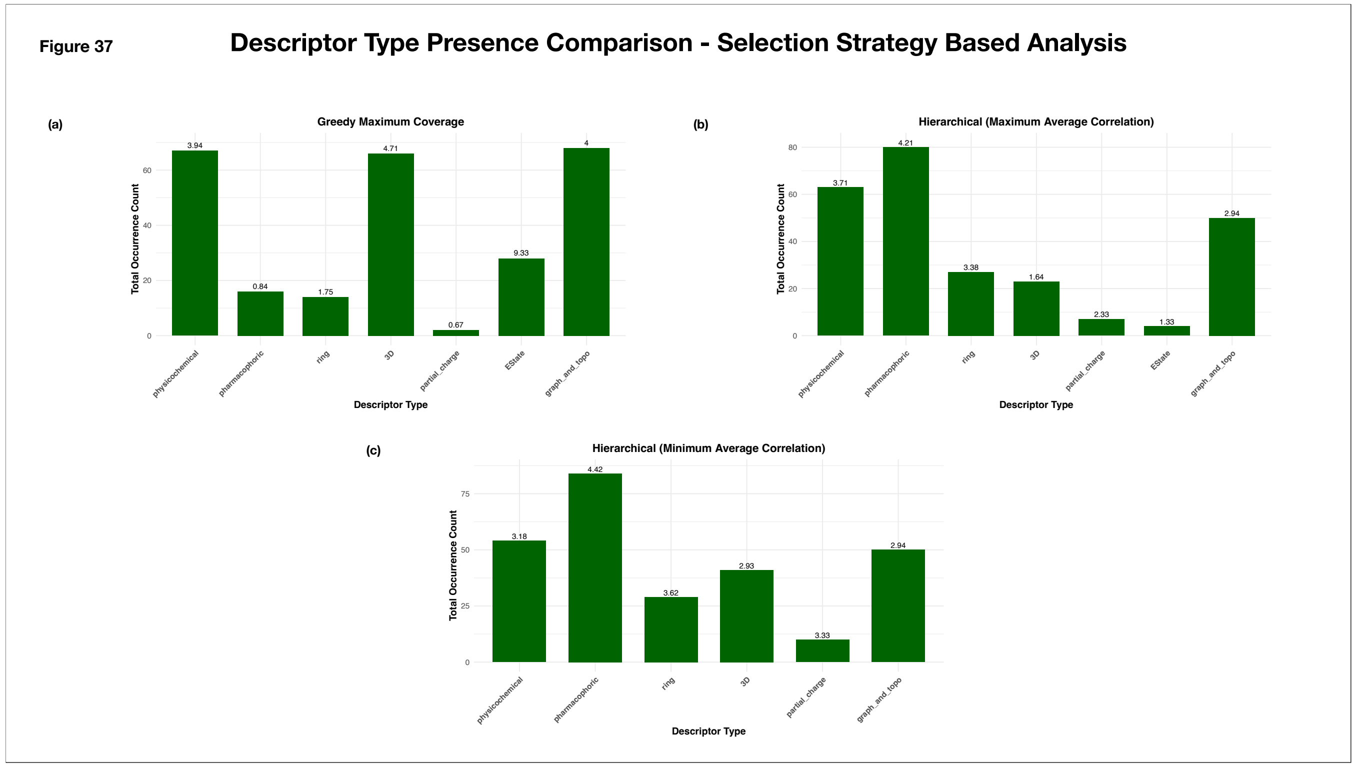


**Supplementary Figure 37. Global Descriptor Type Enrichment Across Descriptor Selection Strategies**

The bar plots summarize the normalized occurrence of descriptor types across all combinations of correlation threshold ranges and descriptor counts for the individual selection strategies. The x-axis represents the descriptor type, while the y-axis denotes the normalized occurrence frequency. Values annotated above each bar indicate the corresponding occurrence of each descriptor type.

(a) Greedy Maximum Coverage

(b) Hierarchical Clustering (Maximum Average Correlation)

(c) Hierarchical Clustering (Minimum Average Correlation)

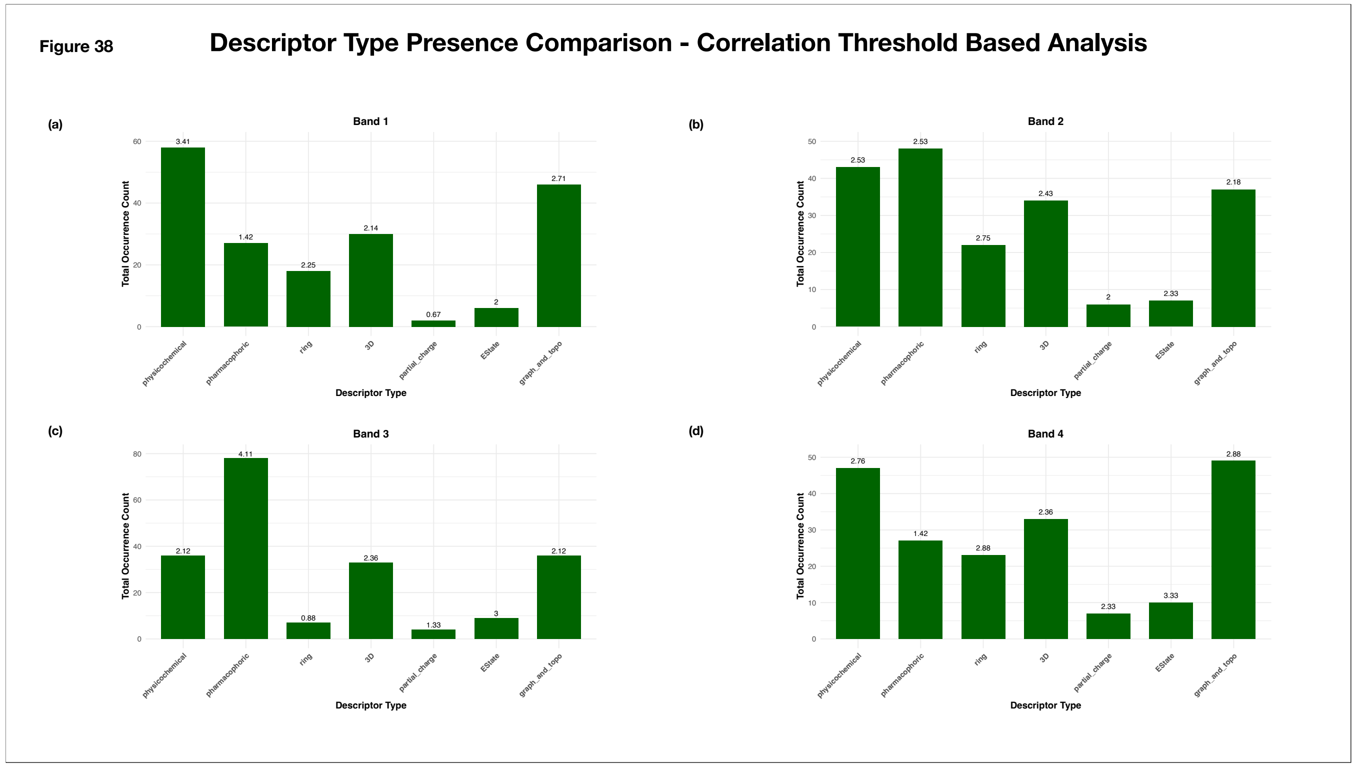


**Supplementary Figure 38. Global Descriptor Type Enrichment Across Correlation Threshold Ranges**

The bar plots summarize the normalized occurrence of descriptor types across all combinations of descriptor selection strategies and descriptor counts for the individual correlation threshold ranges. The x-axis represents the descriptor type, while the y-axis denotes the normalized occurrence frequency. Values annotated above each bar indicate the corresponding occurrence of each descriptor type.

(a) Band 1 (0.5 - 0.85)
(b) Band 2 (0.35 - 0.85)
(c) Band 3 (0.25 - 0.85)
(d) Band 4 (0.35 - 0.75)

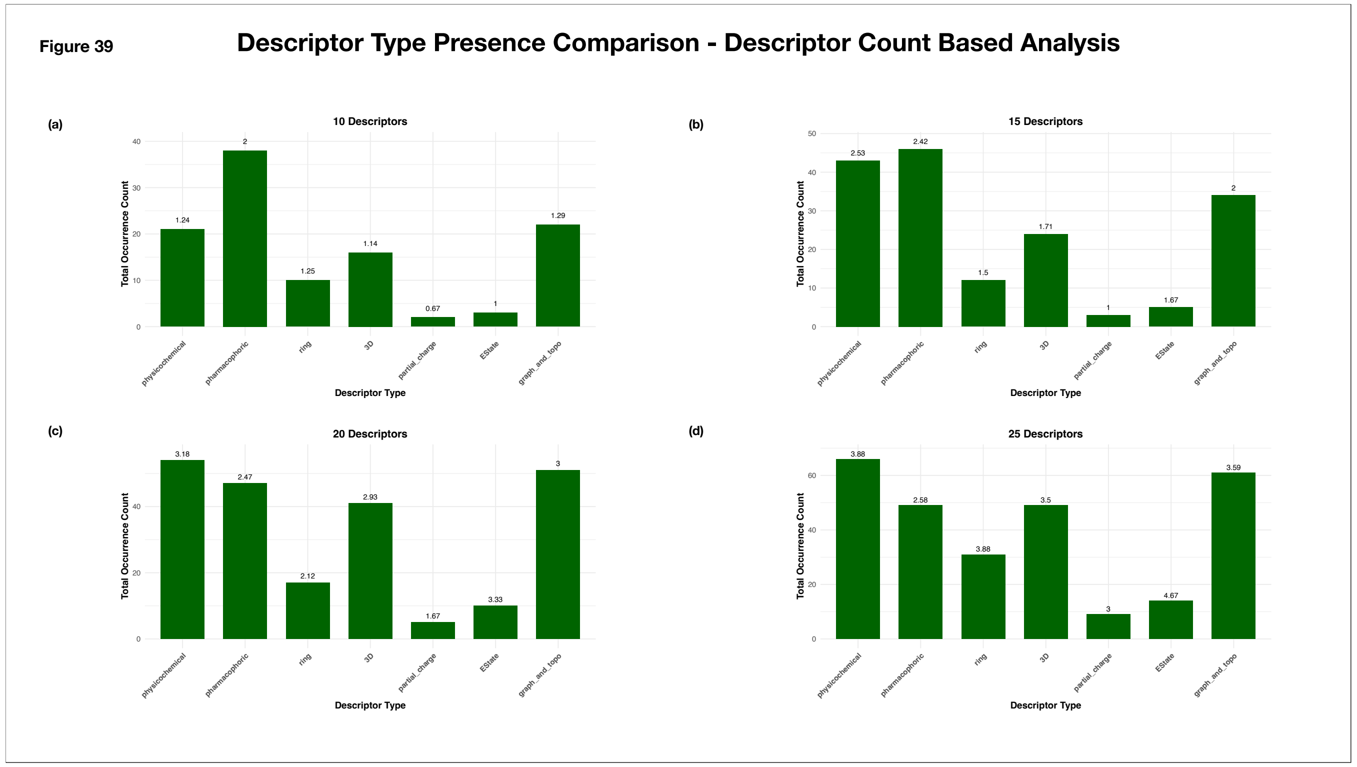


**Supplementary Figure 39. Global Descriptor Type Enrichment Across Descriptor Types**

The bar plots summarize the normalized occurrence of descriptor types across all combinations of descriptor selection strategies and correlation threshold ranges for the individual descriptor counts. The x-axis represents the descriptor type, while the y-axis denotes the normalized occurrence frequency. Values annotated above each bar indicate the corresponding occurrence of each descriptor type.

(a) 10 descriptors
(b) 15 descriptors
(c) 20 descriptors
(d) 25 descriptors

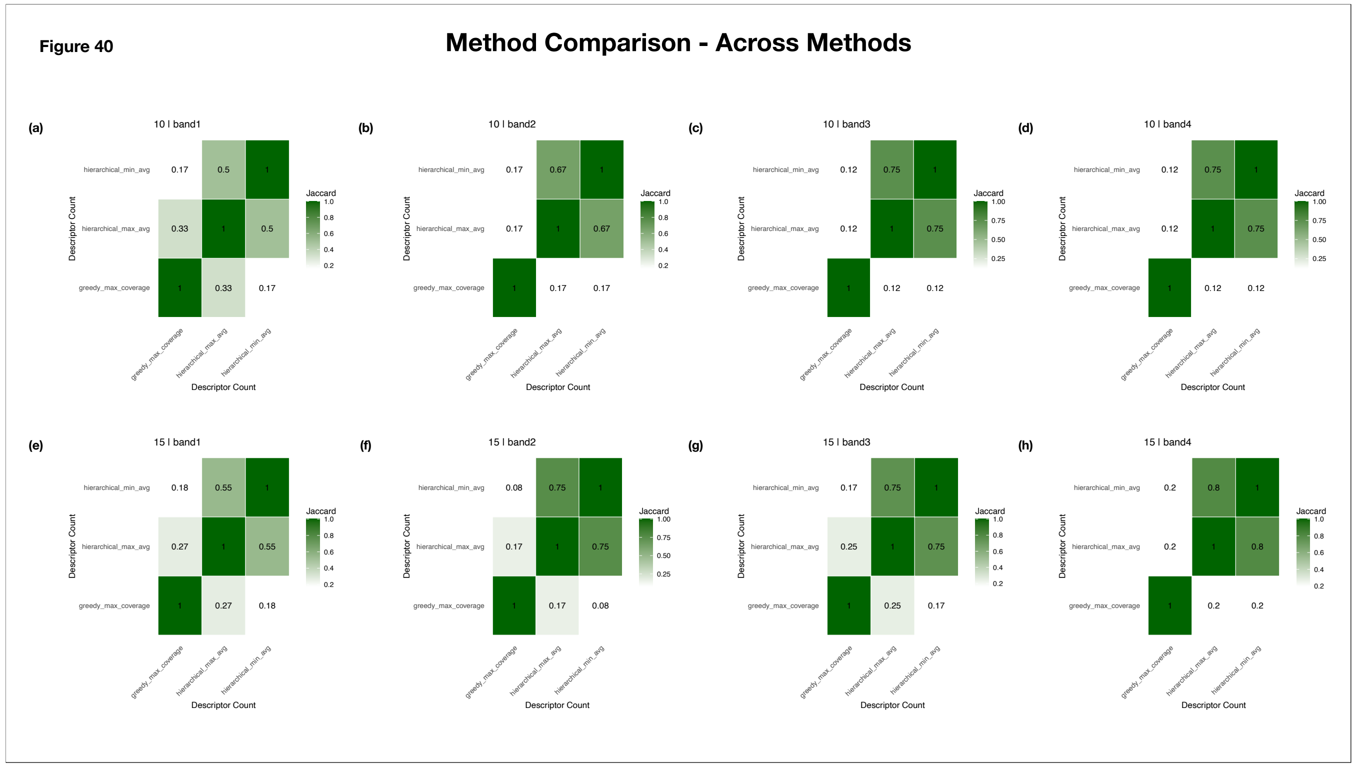


**Supplementary Figure 40: Jaccard Similarity Heatmaps for Evaluation of Descriptor Selection Strategies through Descriptor Overlap**

The heatmaps summarize descriptor overlap through Jaccard Similarity across different combinations of descriptor counts and correlation threshold ranges. Both axes represent descriptor selection strategy, and cell annotations indicate the Jaccard similarity coefficient. Darker shades of green correspond to greater descriptor overlap.

(a) 10 descriptors | Band 1 (0.5 - 0.85)
(b) 10 descriptors | Band 2 (0.35 - 0.85)
(c) 10 descriptors | Band 3 (0.25 - 0.85)
(d) 10 descriptors | Band 4 (0.35 - 0.75)
(e) 15 descriptors | Band 1 (0.5 - 0.85)
(f) 15 descriptors | Band 2 (0.35 - 0.85)
(g) 15 descriptors | Band 3 (0.25 - 0.85)
(h) 15 descriptors | Band 4 (0.35 - 0.75)

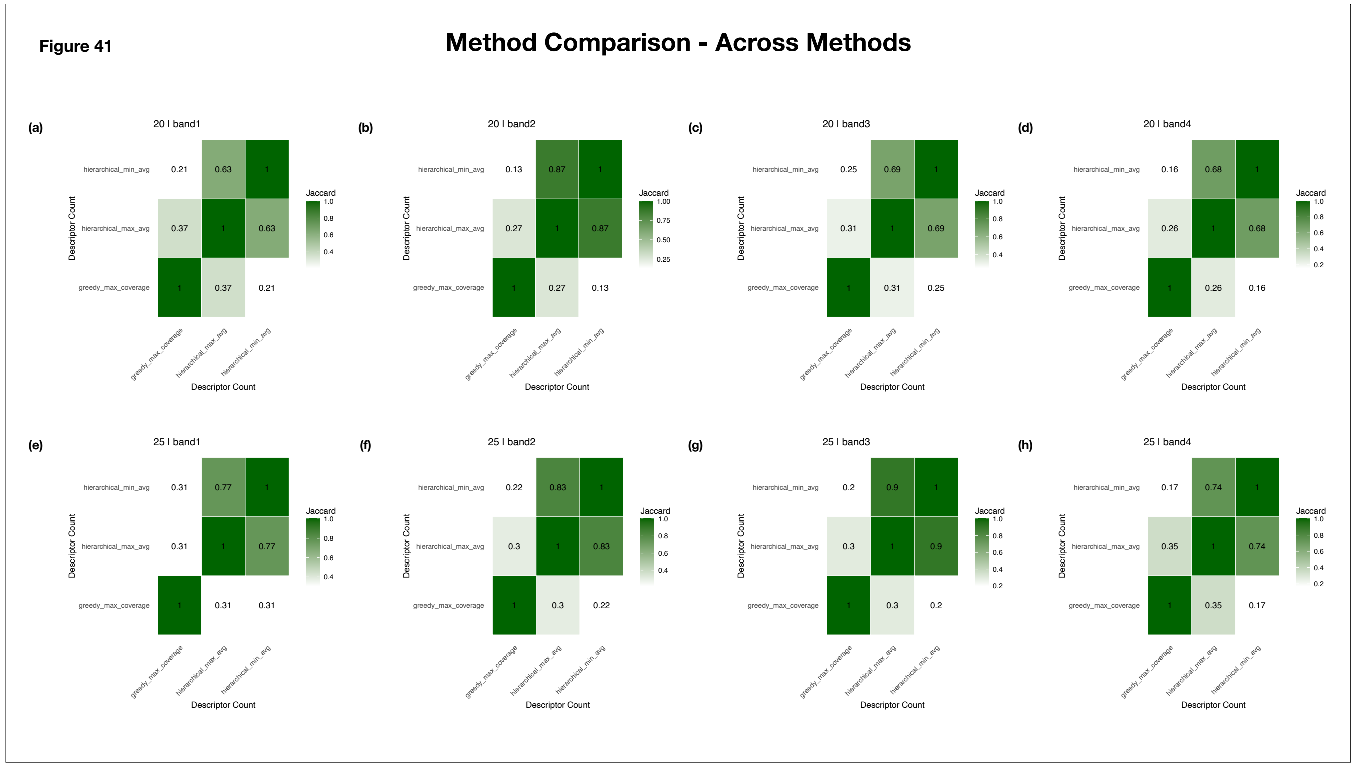


**Supplementary Figure 41: Jaccard Similarity Heatmaps for Evaluation of Descriptor Selection Strategies through Descriptor Overlap**
The heatmaps summarize descriptor overlap through Jaccard Similarity across different combinations of descriptor counts and correlation threshold ranges. Both axes represent descriptor selection strategy, and cell annotations indicate the Jaccard similarity coefficient. Darker shades of green correspond to greater descriptor overlap.
(a) 20 descriptors | Band 1 (0.5 - 0.85)
(b) 20 descriptors | Band 2 (0.35 - 0.85)
(c) 20 descriptors | Band 3 (0.25 - 0.85)
(d) 20 descriptors | Band 4 (0.35 - 0.75)
(e) 25 descriptors | Band 1 (0.5 - 0.85)
(f) 25 descriptors | Band 2 (0.35 - 0.85)
(g) 25 descriptors | Band 3 (0.25 - 0.85)
(h) 25 descriptors | Band 4 (0.35 - 0.75)

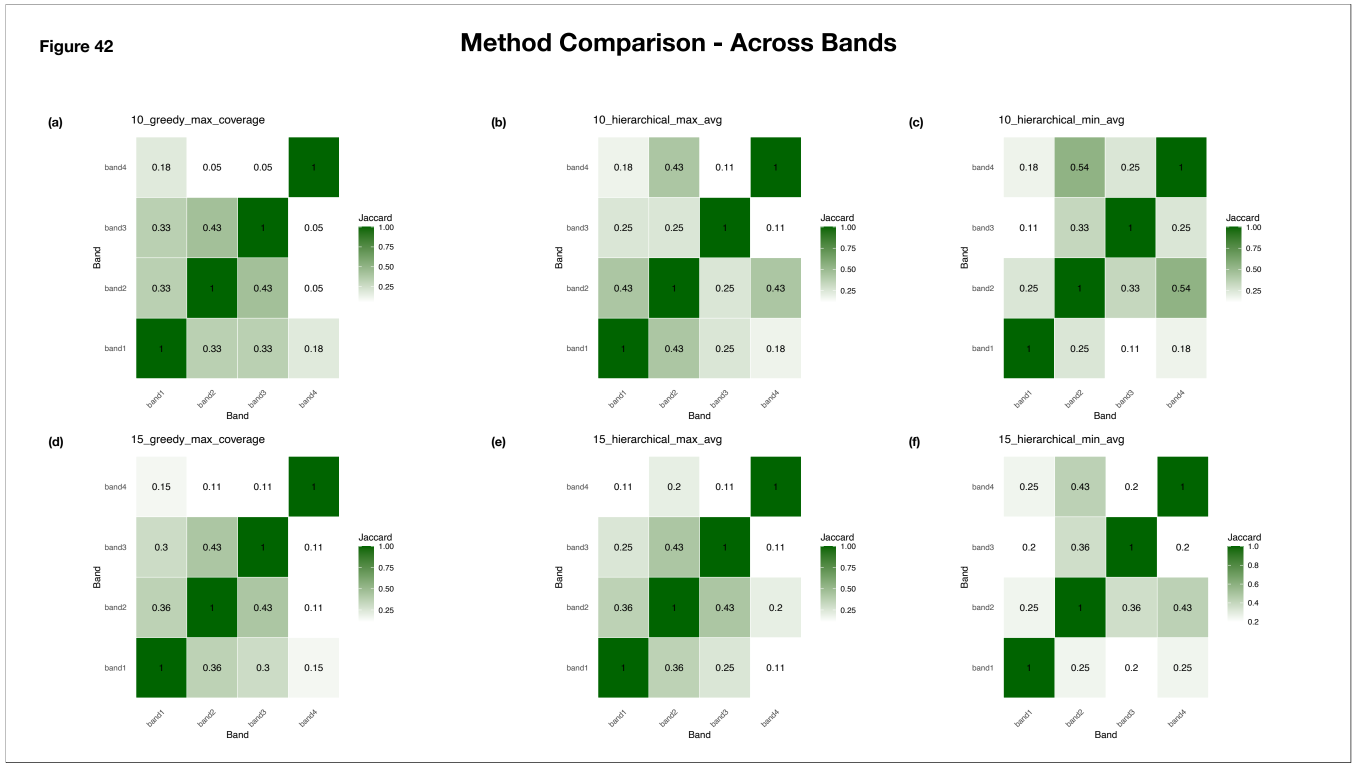


**Supplementary Figure 42: Jaccard Similarity Heatmaps for Evaluation of Correlation Threshold Ranges through Descriptor Overlap**

The heatmaps summarize descriptor overlap through Jaccard Similarity across different combinations of descriptor counts and descriptor selection strategies. Both axes represent correlation threshold ranges, and cell annotations indicate the Jaccard similarity coefficient. Darker shades of green correspond to greater descriptor overlap.

(a) 10 descriptors | Greedy Maximum Coverage

(b) 10 descriptors | Hierarchical Clustering (Maximum Average Correlation)

(c) 10 descriptors | Hierarchical Clustering (Minimum Average Correlation)

(d) 15 descriptors | Greedy Maximum Coverage

(e) 15 descriptors | Hierarchical Clustering (Maximum Average Correlation)

(f) 15 descriptors | Hierarchical Clustering (Minimum Average Correlation)

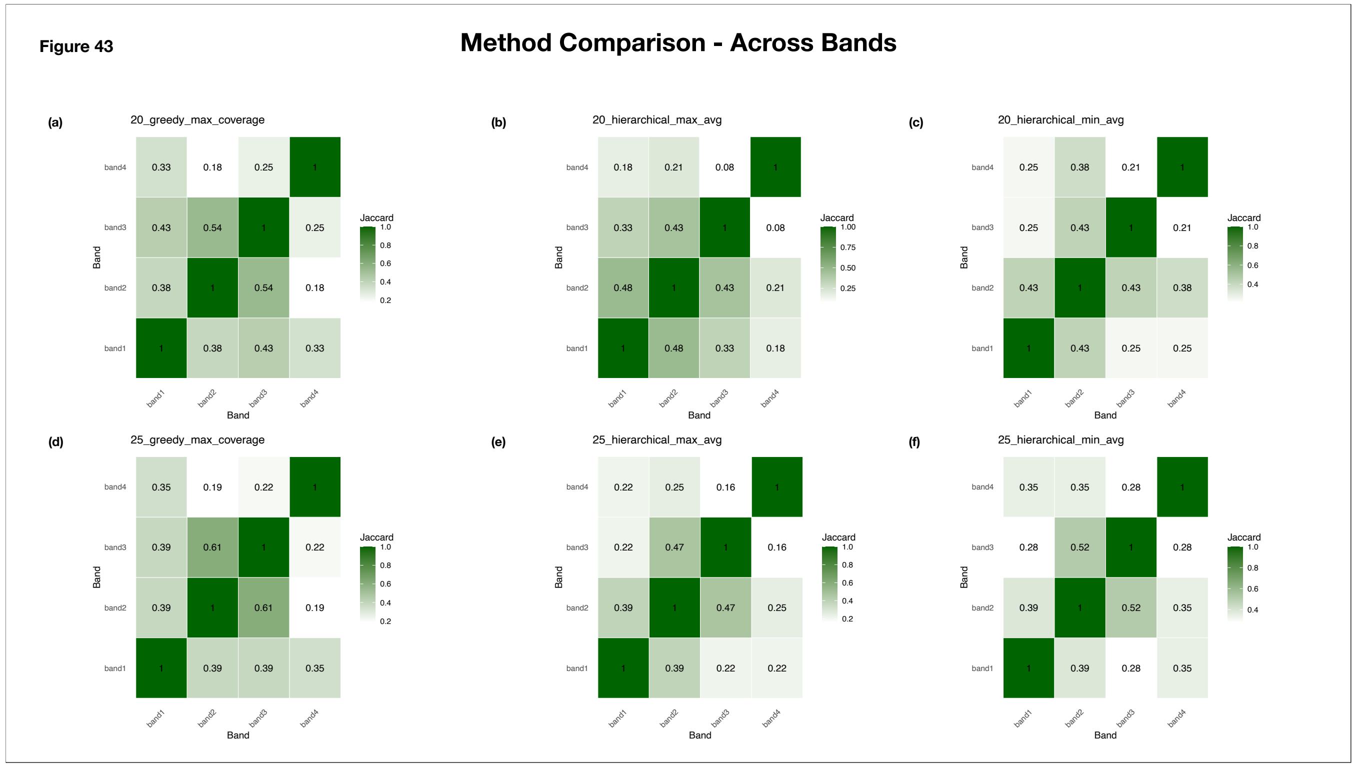


**Supplementary Figure 43: Jaccard Similarity Heatmaps for Evaluation of Correlation Threshold Ranges through Descriptor Overlap**
The heatmaps summarize descriptor overlap through Jaccard Similarity across different combinations of descriptor counts and descriptor selection strategies. Both axes represent correlation threshold ranges, and cell annotations indicate the Jaccard similarity coefficient. Darker shades of green correspond to greater descriptor overlap.
(a) 20 descriptors | Greedy Maximum Coverage
(b) 20 descriptors | Hierarchical Clustering (Maximum Average Correlation)
(c) 20 descriptors | Hierarchical Clustering (Minimum Average Correlation)
(d) 25 descriptors | Greedy Maximum Coverage
(e) 25 descriptors | Hierarchical Clustering (Maximum Average Correlation)
(f) 25 descriptors | Hierarchical Clustering (Minimum Average Correlation)

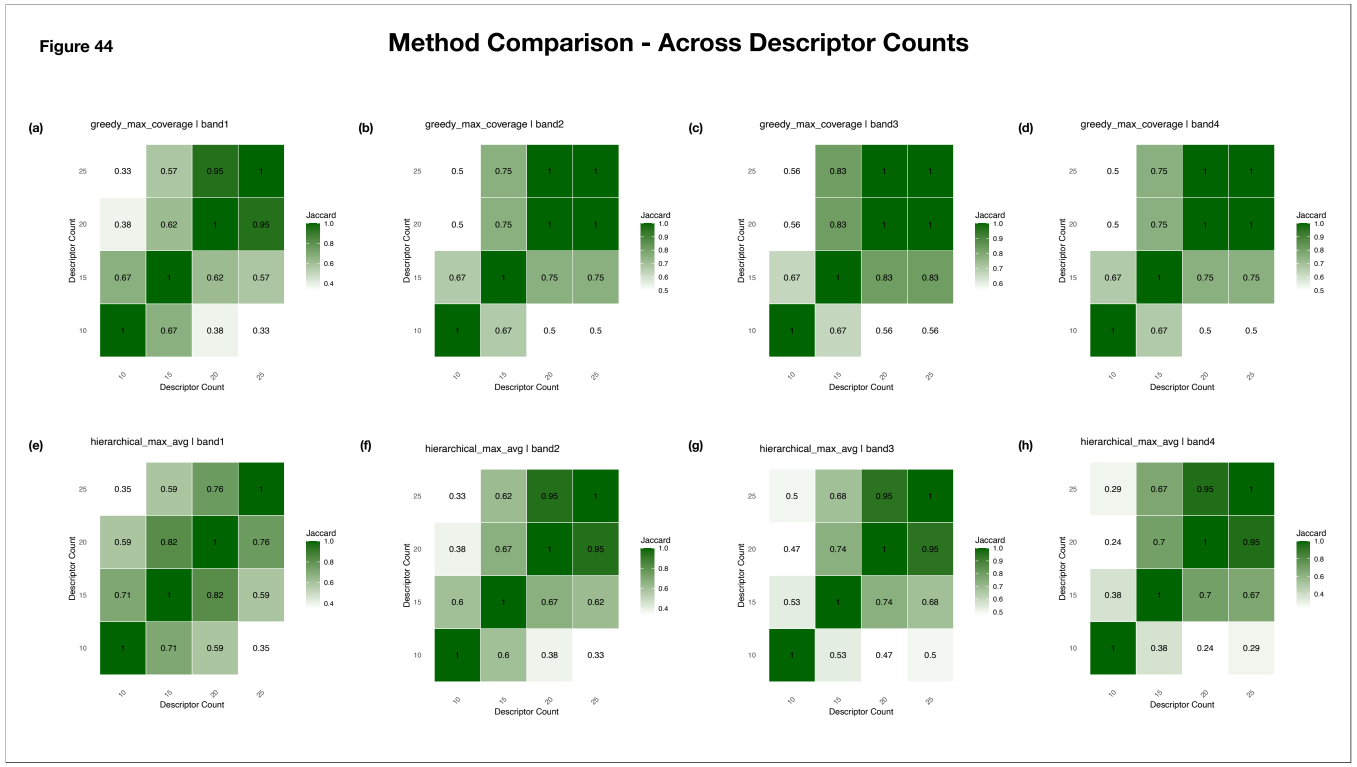


**Supplementary Figure 44: Jaccard Similarity Heatmaps for Evaluation of Descriptor Counts through Descriptor Overlap**
The heatmaps summarize descriptor overlap through Jaccard Similarity across different combinations of correlation threshold ranges and descriptor selection strategies. Both axes represent , and cell annotations indicate the Jaccard similarity coefficient. Darker shades of green correspond to greater descriptor overlap.

(a) Band 1 (0.5 - 0.85) | Greedy Maximum Coverage
(b) Band 2 (0.35 - 0.85) | Greedy Maximum Coverage
(c) Band 3 (0.25 - 0.85) | Greedy Maximum Coverage
(d) Band 4 (0.35 - 0.75) | Greedy Maximum Coverage
(e) Band 1 (0.5 - 0.85) | Hierarchical Clustering (Maximum Average Correlation)
(f) Band 2 (0.35 - 0.85) | Hierarchical Clustering (Maximum Average Correlation)
(g) Band 3 (0.25 - 0.85) | Hierarchical Clustering (Maximum Average Correlation)
(h) Band 4 (0.35 - 0.75) |Hierarchical Clustering (Maximum Average Correlation)

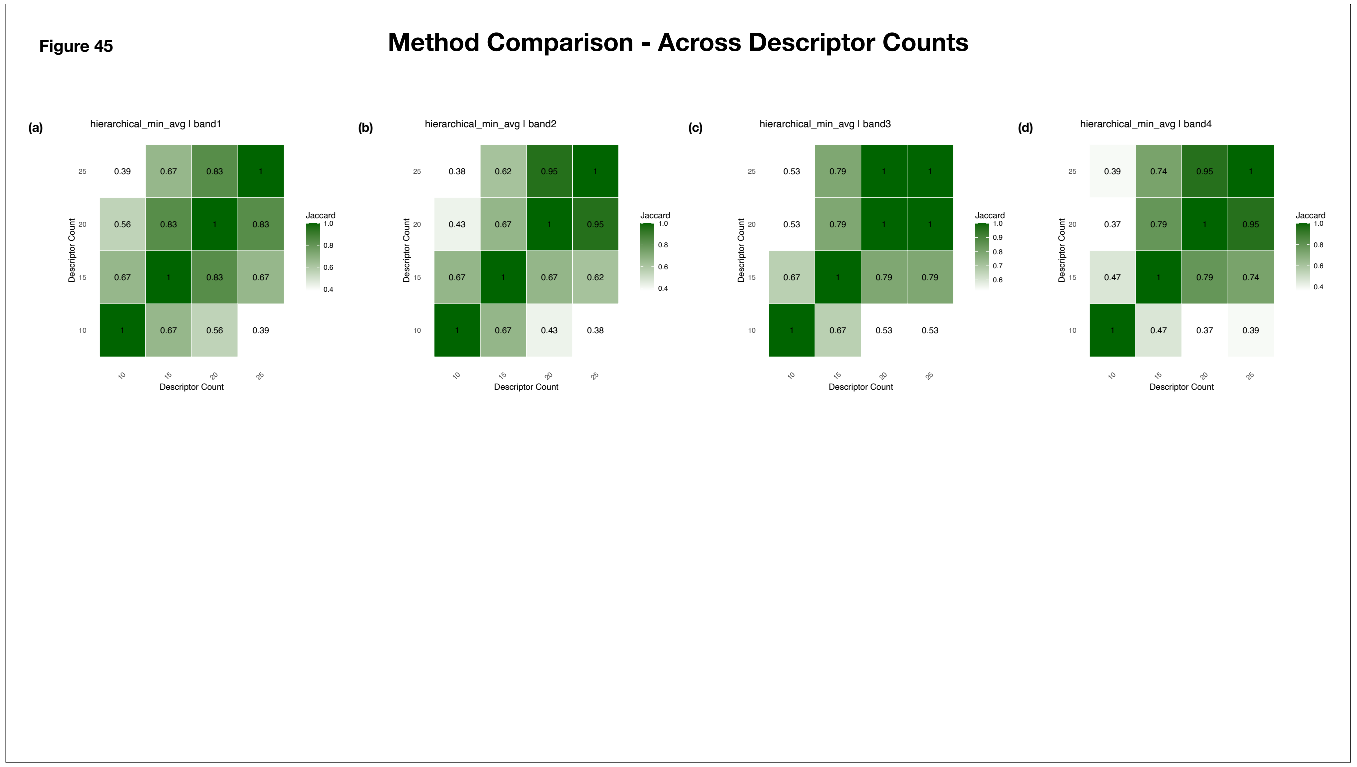


**Supplementary Figure 45: Jaccard Similarity Heatmaps for Evaluation of Descriptor Counts through Descriptor Overlap**
The heatmaps summarize descriptor overlap through Jaccard Similarity across different combinations of correlation threshold ranges and descriptor selection strategies. Both axes represent , and cell annotations indicate the Jaccard similarity coefficient. Darker shades of green correspond to greater descriptor overlap.
(a) Band 1 (0.5 - 0.85) | Hierarchical Clustering (Minimum Average Correlation)
(b) Band 2 (0.35 - 0.85) | Hierarchical Clustering (Minimum Average Correlation)
(c) Band 3 (0.25 - 0.85) | Hierarchical Clustering (Minimum Average Correlation)
(d) Band 4 (0.35 - 0.75) |Hierarchical Clustering (Minimum Average Correlation)

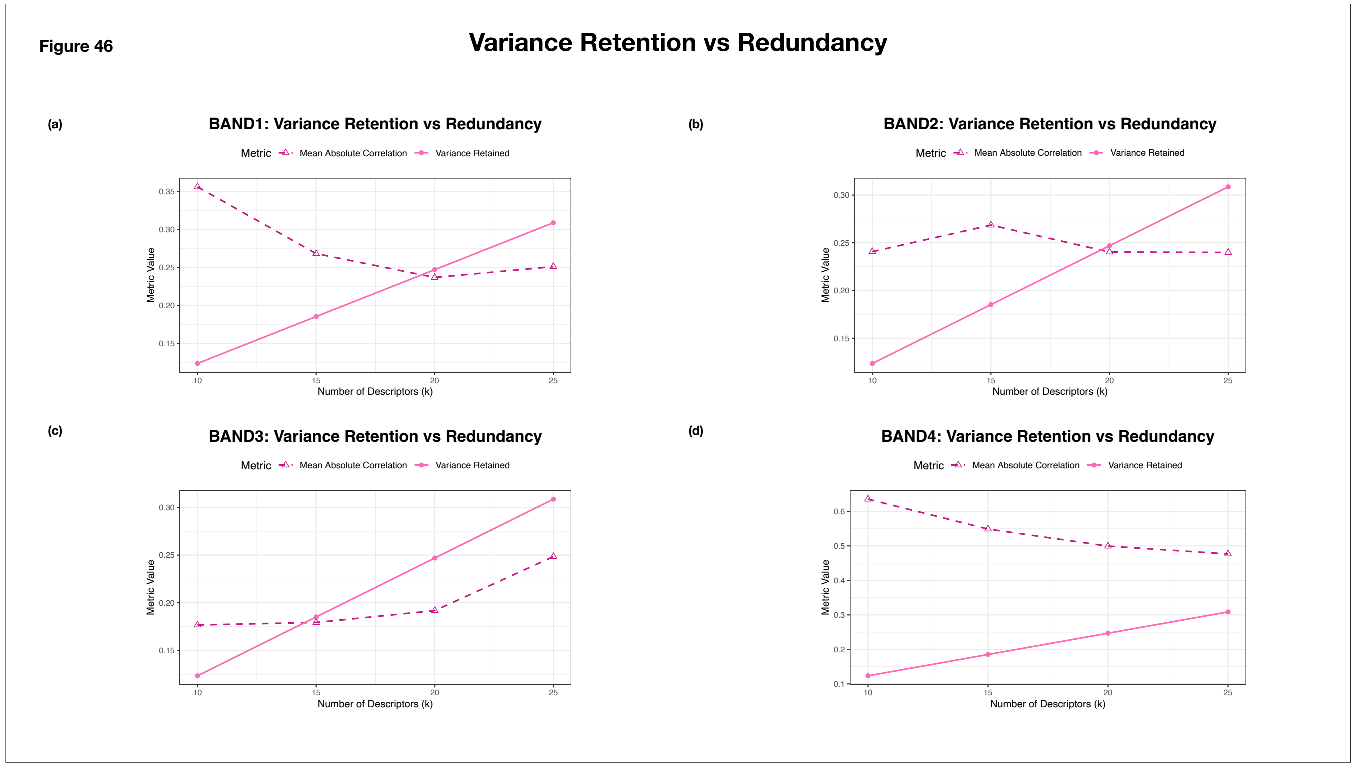


**Supplementary Figure 46: Variance Retention vs Redundancy**

The line plots summarize the trade-off between retained variance and descriptor redundancy for the selected correlation threshold range. The x-axis represents descriptor count, and the y-axis represents the corresponding metric value. The solid pink line denotes the proportion of variance retained, while the dotted pink line with triangular markers represents the mean absolute correlation.

(a) Band 1 (0.5 - 0.85)
(b) Band 2 (0.35 - 0.85)
(c) Band 3 (0.25 - 0.85)
(d) Band 4 (0.35 - 0.75)

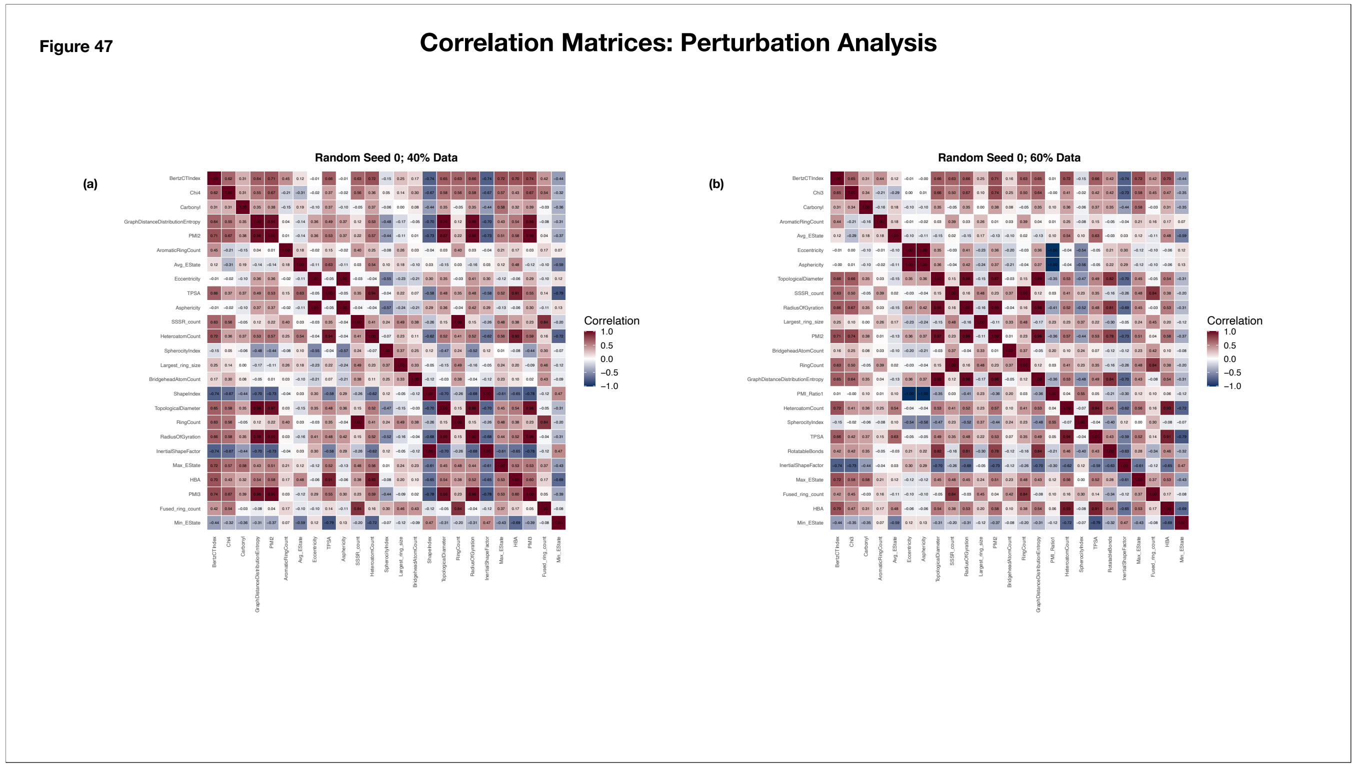


**Supplementary Figure 47. Correlation Matrix of Descriptors Selected by the Greedy Maximum Coverage Method (Correlation Threshold Range: 0.35–0.85; 25 Descriptors) During Perturbation Analysis (Random Seed: 0 )**

The heat map displays pairwise Spearman correlation coefficients among the 25 descriptors selected by the Greedy Maximum Coverage method from a perturbed dataset split. Both axes represent the selected descriptors, and annotated cells indicate the corresponding correlation coefficients. Colors range from dark blue (−1) to dark red (+1), with white indicating no correlation (0).

(a) 40% of the dataset

(b) 60% of the dataset

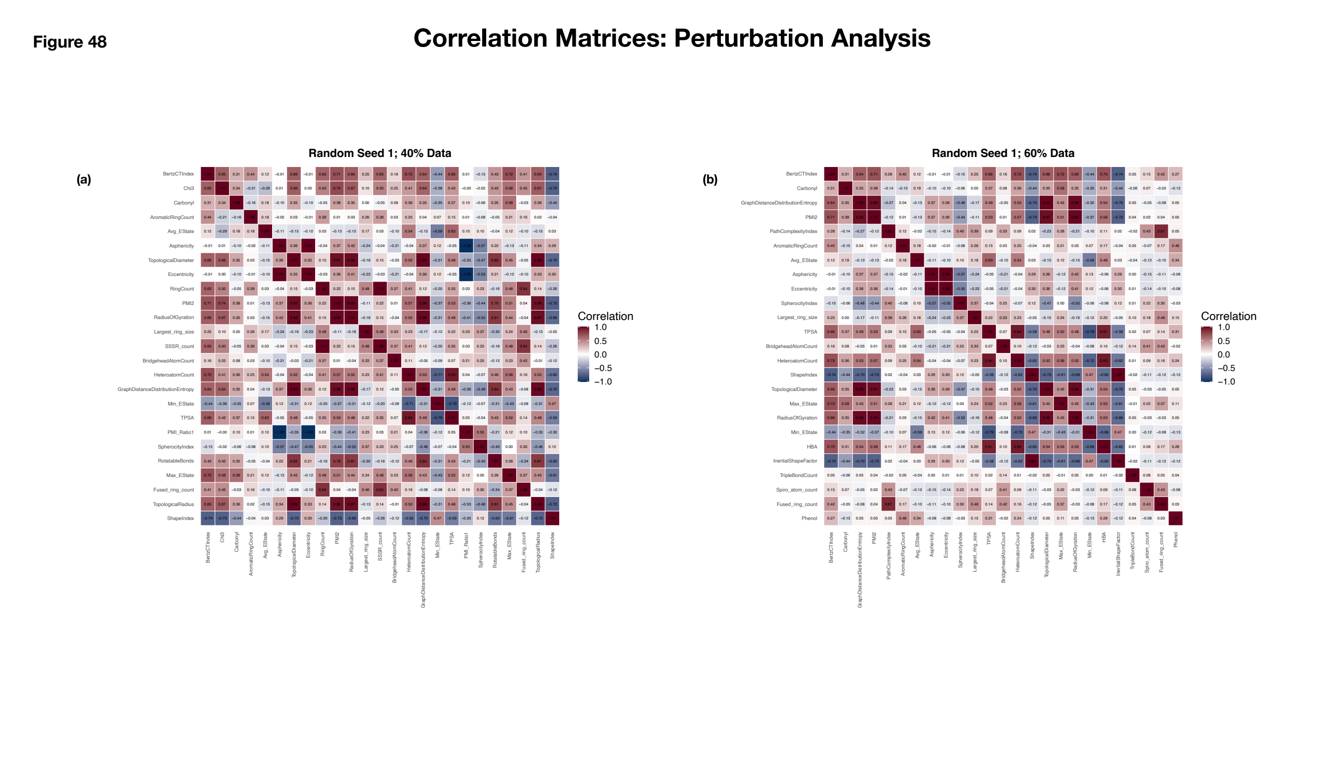


**Supplementary Figure 48. Correlation Matrix of Descriptors Selected by the Greedy Maximum Coverage Method (Correlation Threshold Range: 0.35–0.85; 25 Descriptors) During Perturbation Analysis (Random Seed: 1)**
The heat map displays pairwise Spearman correlation coefficients among the 25 descriptors selected by the Greedy Maximum Coverage method from a perturbed dataset split. Both axes represent the selected descriptors, and annotated cells indicate the corresponding correlation coefficients. Colors range from dark blue (−1) to dark red (+1), with white indicating no correlation (0).
(a) 40% of the dataset
(b) 60% of the dataset

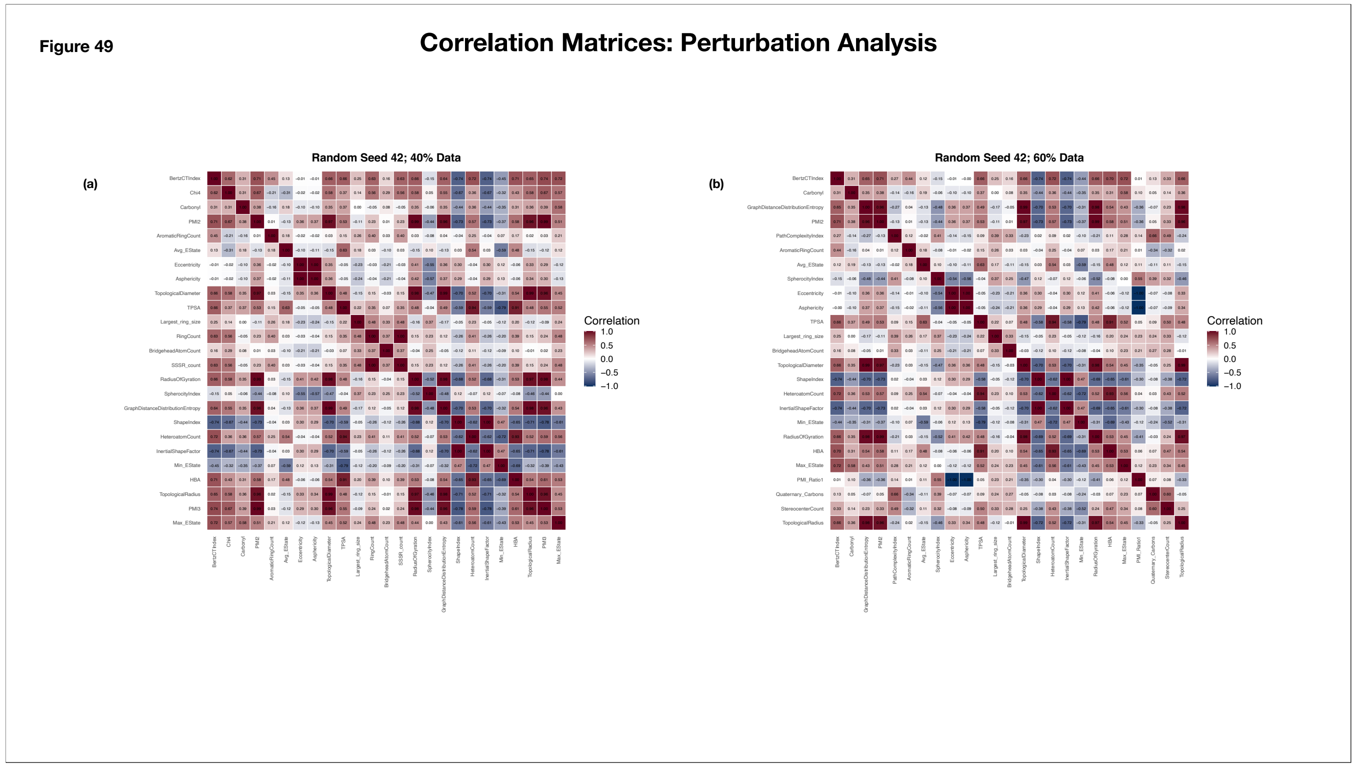


**Supplementary Figure 49. Correlation Matrix of Descriptors Selected by the Greedy Maximum Coverage Method (Correlation Threshold Range: 0.35–0.85; 25 Descriptors) During Perturbation Analysis (Random Seed: 42)**

The heat map displays pairwise Spearman correlation coefficients among the 25 descriptors selected by the Greedy Maximum Coverage method from a perturbed dataset split. Both axes represent the selected descriptors, and annotated cells indicate the corresponding correlation coefficients. Colors range from dark blue (−1) to dark red (+1), with white indicating no correlation (0).

(a) 40% of the dataset

(b) 60% of the dataset

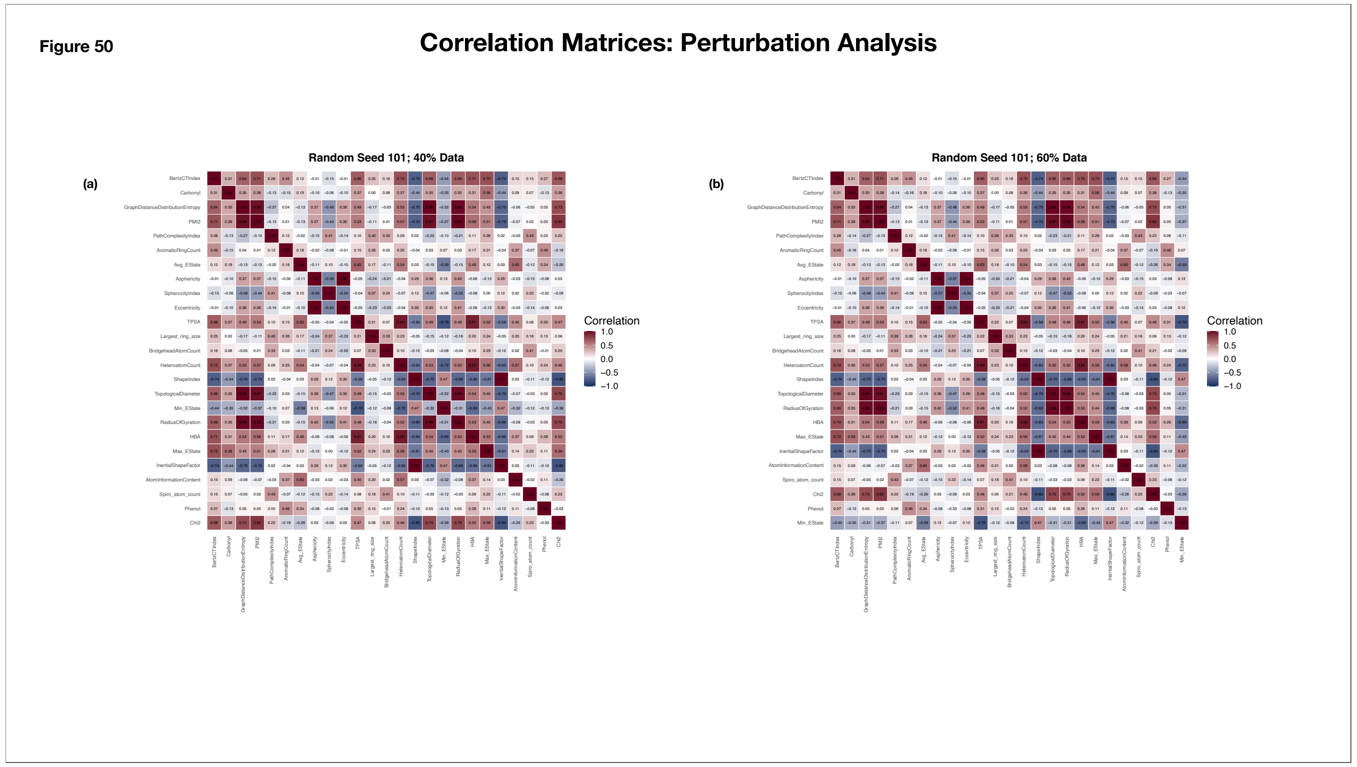

**Supplementary Figure 50. Correlation Matrix of Descriptors Selected by the Greedy Maximum Coverage Method (Correlation Threshold Range: 0.35–0.85; 25 Descriptors) During Perturbation Analysis (Random Seed: 101)**
The heat map displays pairwise Spearman correlation coefficients among the 25 descriptors selected by the Greedy Maximum Coverage method from a perturbed dataset split. Both axes represent the selected descriptors, and annotated cells indicate the corresponding correlation coefficients. Colors range from dark blue (−1) to dark red (+1), with white indicating no correlation (0).
(a) 40% of the dataset
(b) 60% of the dataset

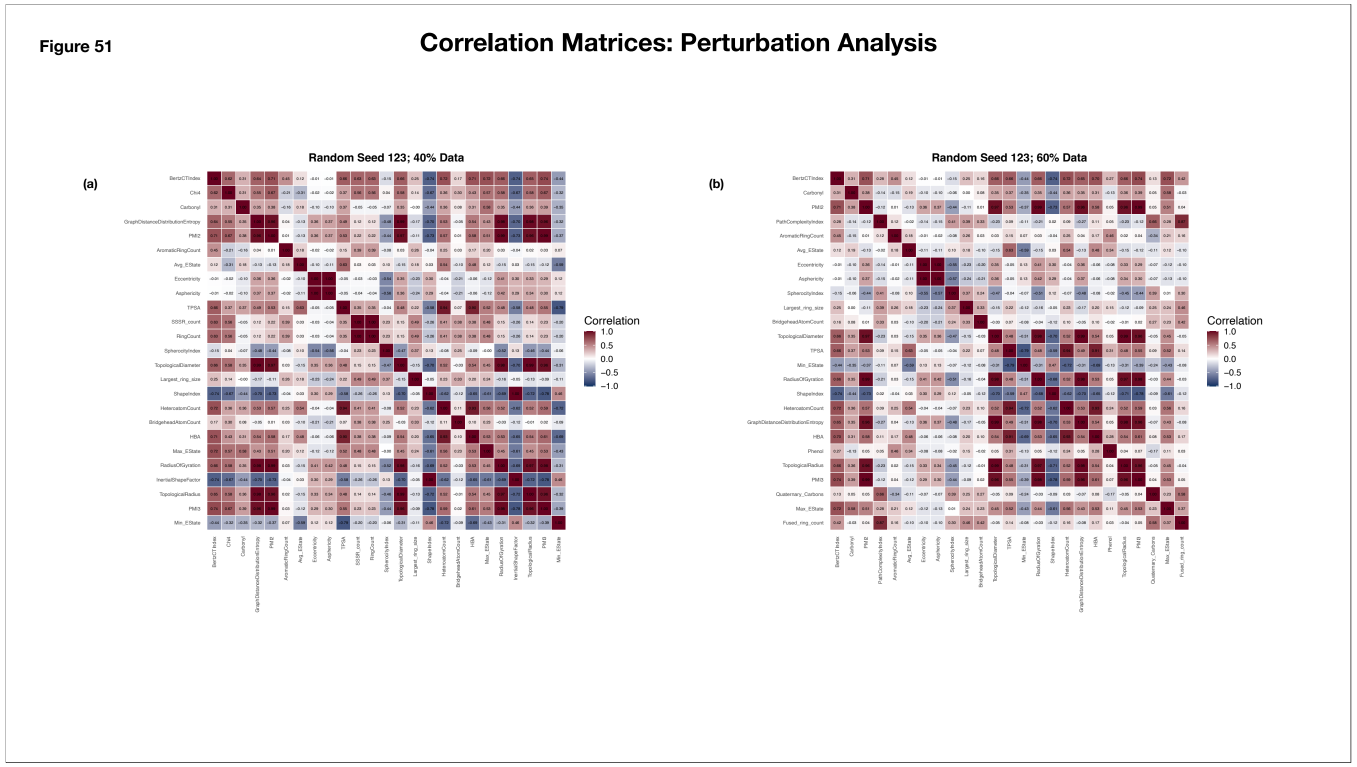


**Supplementary Figure 51. Correlation Matrix of Descriptors Selected by the Greedy Maximum Coverage Method (Correlation Threshold Range: 0.35–0.85; 25 Descriptors) During Perturbation Analysis (Random Seed: 123)**

The heat map displays pairwise Spearman correlation coefficients among the 25 descriptors selected by the Greedy Maximum Coverage method from a perturbed dataset split. Both axes represent the selected descriptors, and annotated cells indicate the corresponding correlation coefficients. Colors range from dark blue (−1) to dark red (+1), with white indicating no correlation (0).

(a) 40% of the dataset

(b) 60% of the dataset

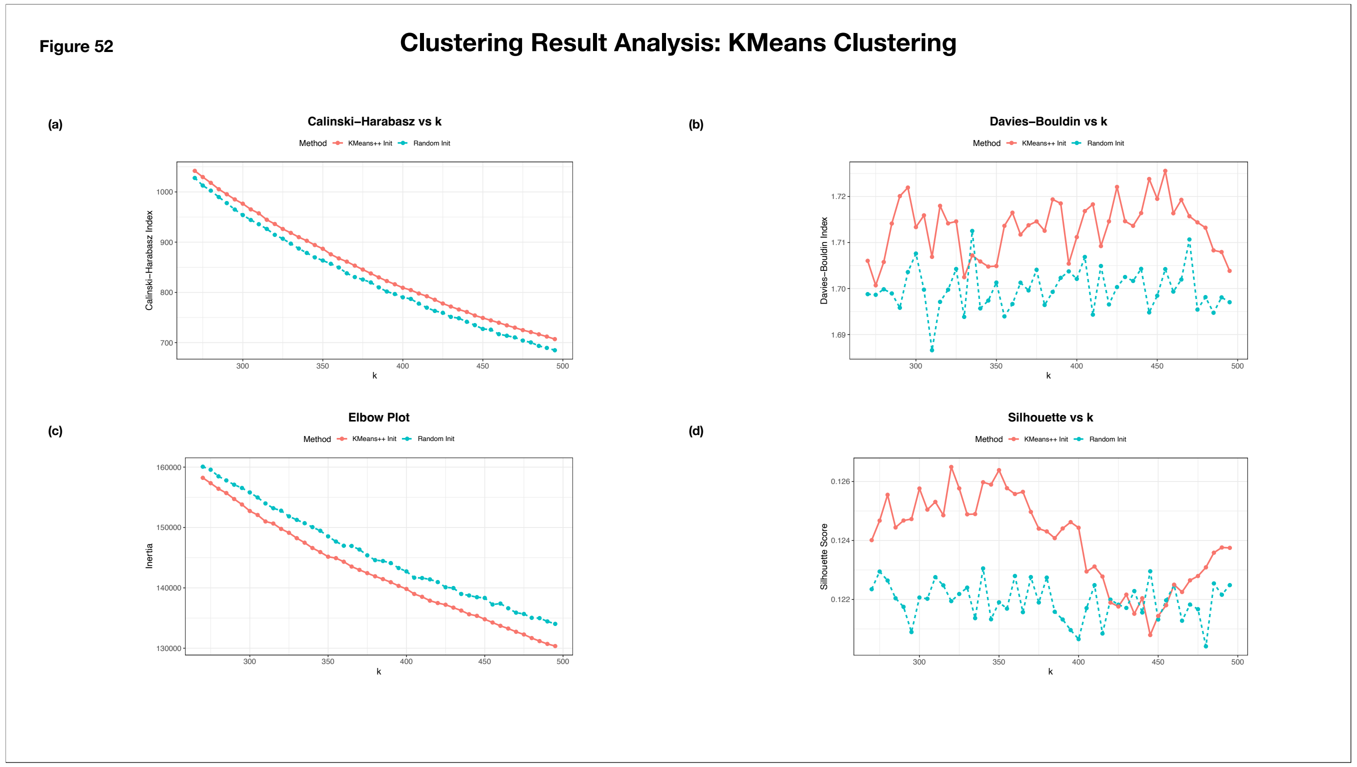


**Supplementary Figure 52: Clustering Metrics for K-Means Clustering using the Sample Dataset**

The line plots summarize the performance of different clustering metrics across varying numbers of clusters generated using K-Means clustering. The x-axis represents the number of clusters, while the y-axis represents the corresponding clustering metric value. Red solid lines denote results obtained using k-means++ initialization, whereas green dotted lines represent results obtained using random initialization.

(a) Calinski-Harabasz Index
(b) Davies Bouldin Index
(c) Inertia
(d) Silhouette Score

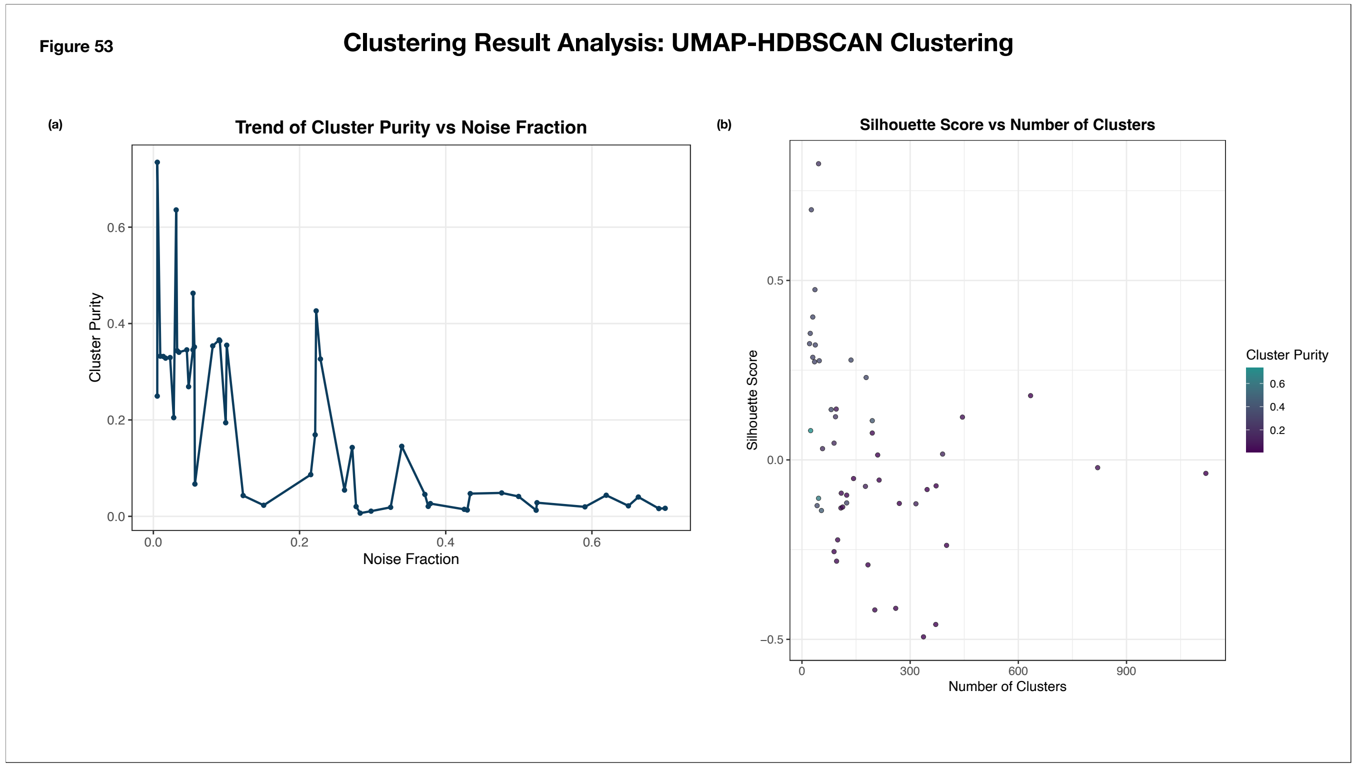


**Supplementary Figure 53: Clustering Metrics for UMAP-HDBSCAN Clustering using the Sample Dataset**

(a) **Trend of Cluster Purity versus Noise Fraction**: The line plot illustrates the relationship between cluster purity and noise fraction for clusters generated using UMAP–HDBSCAN. The x-axis represents the noise fraction, while the y-axis represents cluster purity. Each highlighted point corresponds to a unique UMAP–HDBSCAN hyperparameter combination.

(b) **Trend of Silhouette Score versus Number of Clusters**: The scatter plot summarizes the relationship between the number of clusters and silhouette score across different UMAP–HDBSCAN hyperparameter combinations. The x-axis represents the number of clusters, while the y-axis represents the silhouette score. Each point corresponds to a unique hyperparameter combination, with point color indicating cluster purity. Colors range from purple (low cluster purity) to green (high cluster purity).

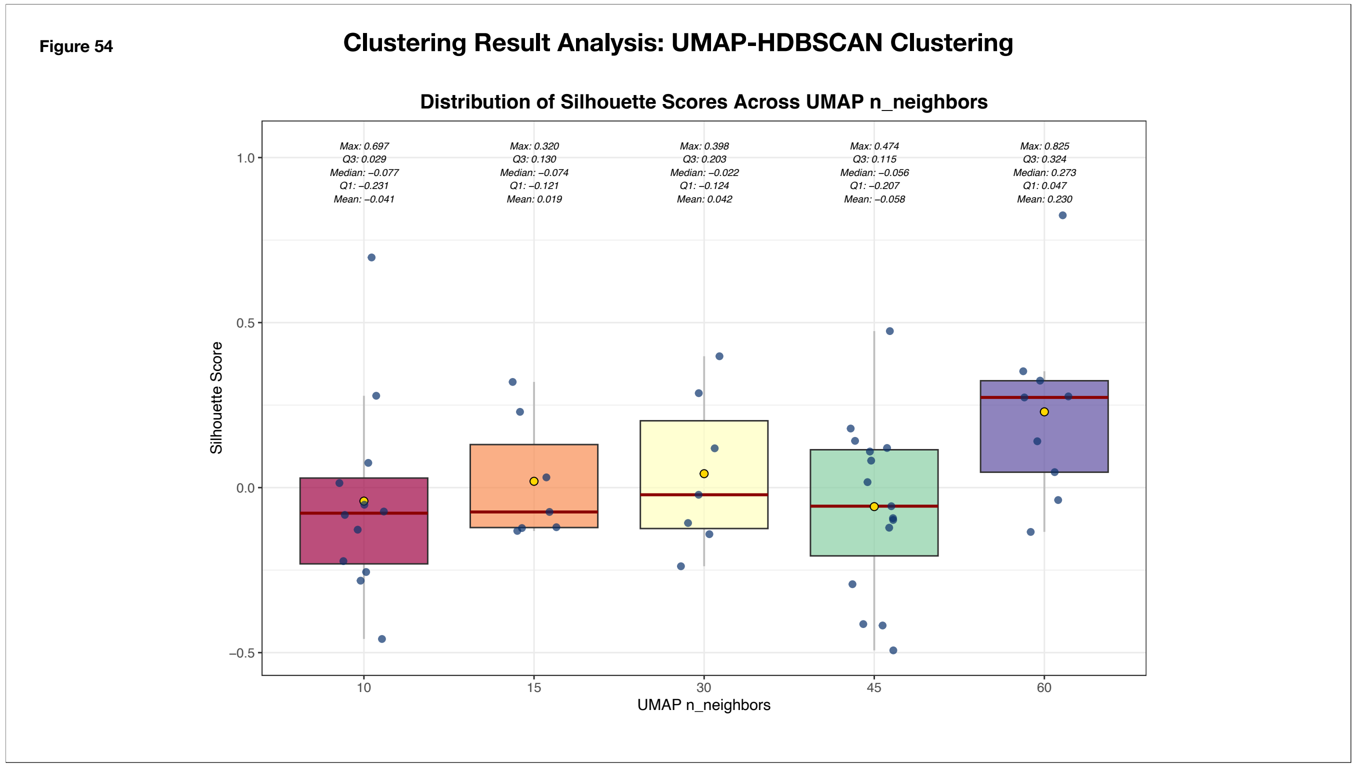


**Supplementary Figure 54: Clustering Metrics for UMAP-HDBSCAN Clustering using the Sample Dataset - Distribution of Silhouette Scores across different values of UMAP n_neighbours**

The box plots summarize the distribution of silhouette scores across different values of the UMAP n_neighbors hyperparameter. The x-axis represents individual n_neighbors values, while the y-axis represents the silhouette score. Each box denotes the interquartile range (Q1–Q3), with the median indicated by a red horizontal line and the mean represented by a yellow point. Individual points correspond to distinct hyperparameter combinations. Whiskers extend to the minimum and maximum observed silhouette scores. The mean, minimum, maximum, Q1, and Q3 values are annotated above each box.

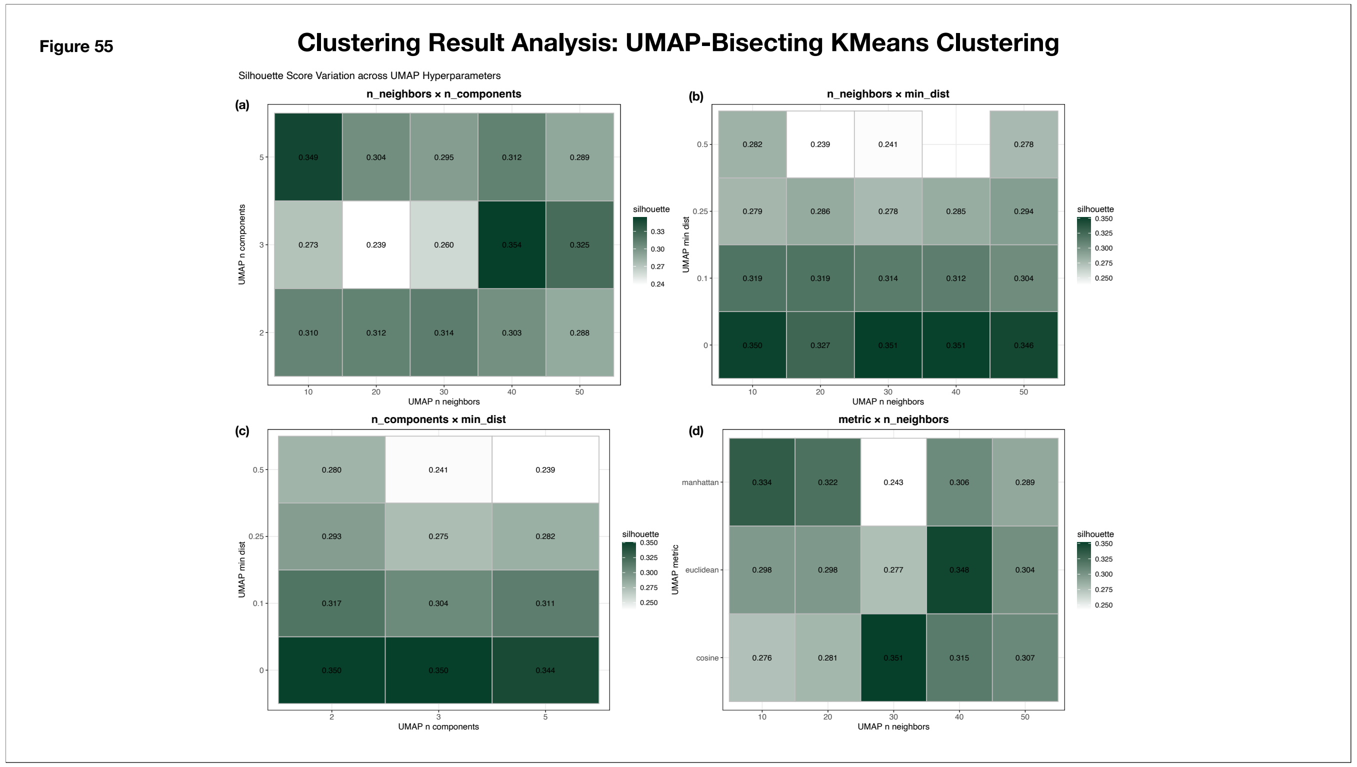


**Supplementary Figure 55: Clustering Result Analysis for UMAP-Bisecting KMeans: Silhouette Score Variations across Different UMAP Hyperparameters**

The heat maps summarize clustering performance, measured by silhouette score, across different UMAP hyperparameter combinations used for UMAP–Bisecting K-Means clustering. The x- and y-axes represent the corresponding UMAP hyperparameters being compared. Darker shades indicate higher silhouette scores, and each cell is annotated with the corresponding score.

(a) **n_components vs n_neighbors**: Comparison of silhouette scores across combinations of UMAP n_components and n_neighbors.

(b) **min_dist vs n_neighbors:** Comparison of silhouette scores across combinations of UMAP min_dist and n_neighbors.

(c) **min_dist vs n_components**: Comparison of silhouette scores across combinations of UMAP min_dist and n_components.

(d) **Metric vs n_neighbors**: Comparison of silhouette scores across combinations of UMAP distance metrics and n_neighbors values.

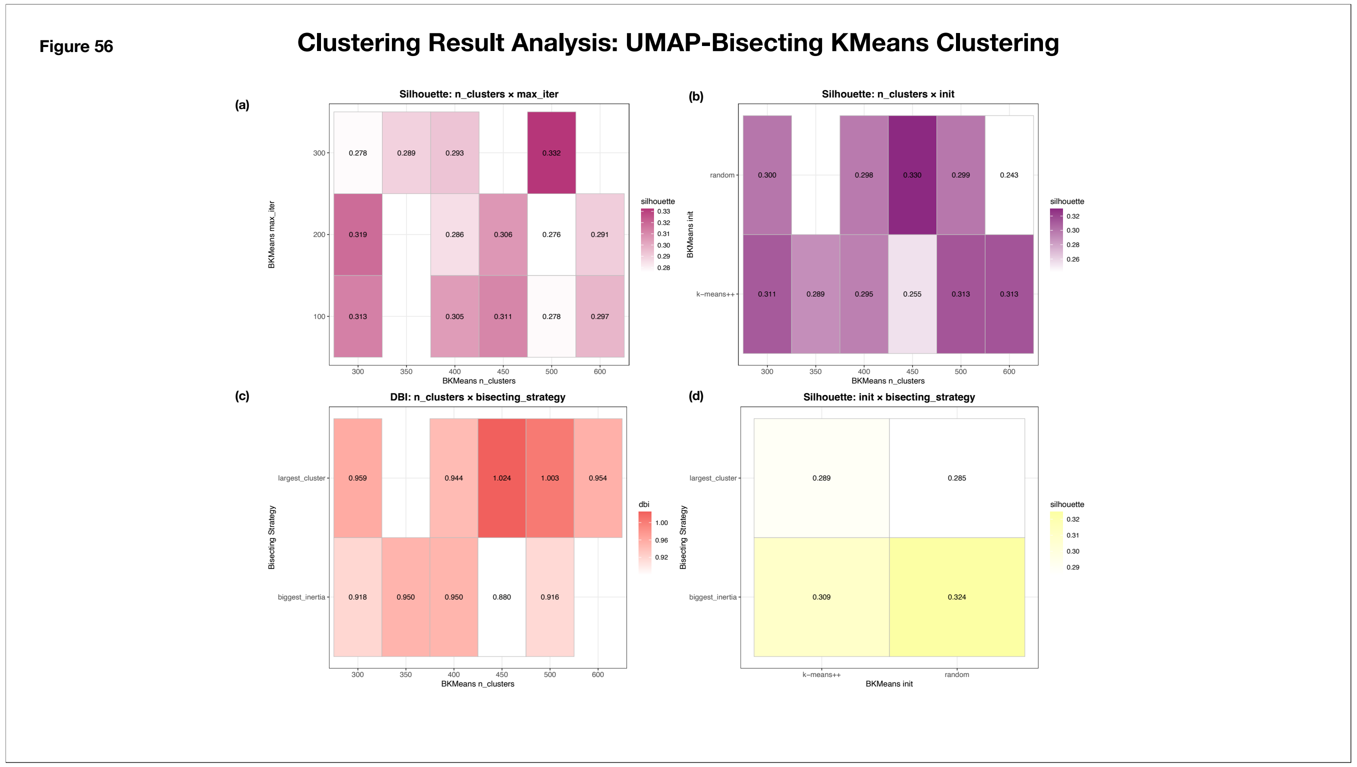


**Supplementary Figure 56: Clustering Result Analysis for UMAP-Bisecting KMeans: Silhouette Score and DBI Variations across Different UMAP Hyperparameters**

The heat maps summarize clustering performance, measured by silhouette score and DBI, across different UMAP hyperparameter combinations used for UMAP–Bisecting K-Means clustering. The x- and y-axes represent the corresponding Bisecting KMeans hyperparameters being compared. Darker shades indicate higher silhouette/DBI scores, and each cell is annotated with the corresponding score.

**(a) max_iter v/s n_neighbours:** comparison of Silhouette scores across combinations of max_iter and n_clusters

**(b) Init v/s n_clusters**: comparison of Silhouette scores across combinations of init and n_clusters

**(c) bisecting_strategy v/s n_clusters**: Davies-Bouldin indices across combinations of bisecting strategy and n_clusters

**(d) bisecting_strategy v/s init**: Silhouette scores across combinations of bisecting strategy and initialization technique

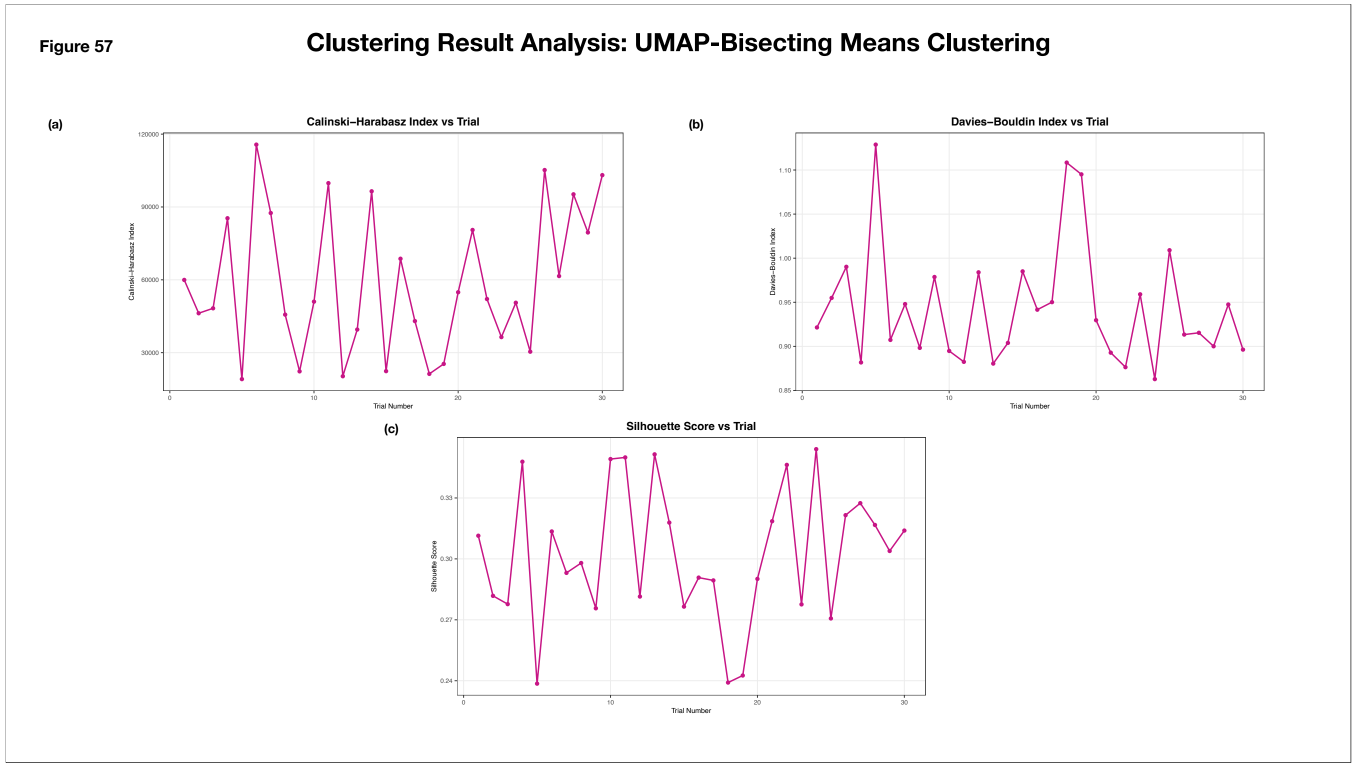


**Supplementary Figure 57: Trial-Wise Results of UMAP-Bisecting KMeans Clustering on the Sample Dataset**

The line plots summarize clustering performance across different UMAP–Bisecting K-Means hyperparameter combinations. The x-axis represents the trial number, while the y-axis represents the value of the corresponding clustering metric. Each marked point corresponds to an individual hyperparameter trial.

(a) **Calinski–Harabasz Index per Trial:** Distribution of Calinski–Harabasz index values across all hyperparameter trials

(b) **Davies–Bouldin Index per Trial**: Distribution of Davies–Bouldin index values across all hyperparameter trials.

(c) **Silhouette Score per Trial**: Distribution of silhouette score values across all hyperparameter trials.

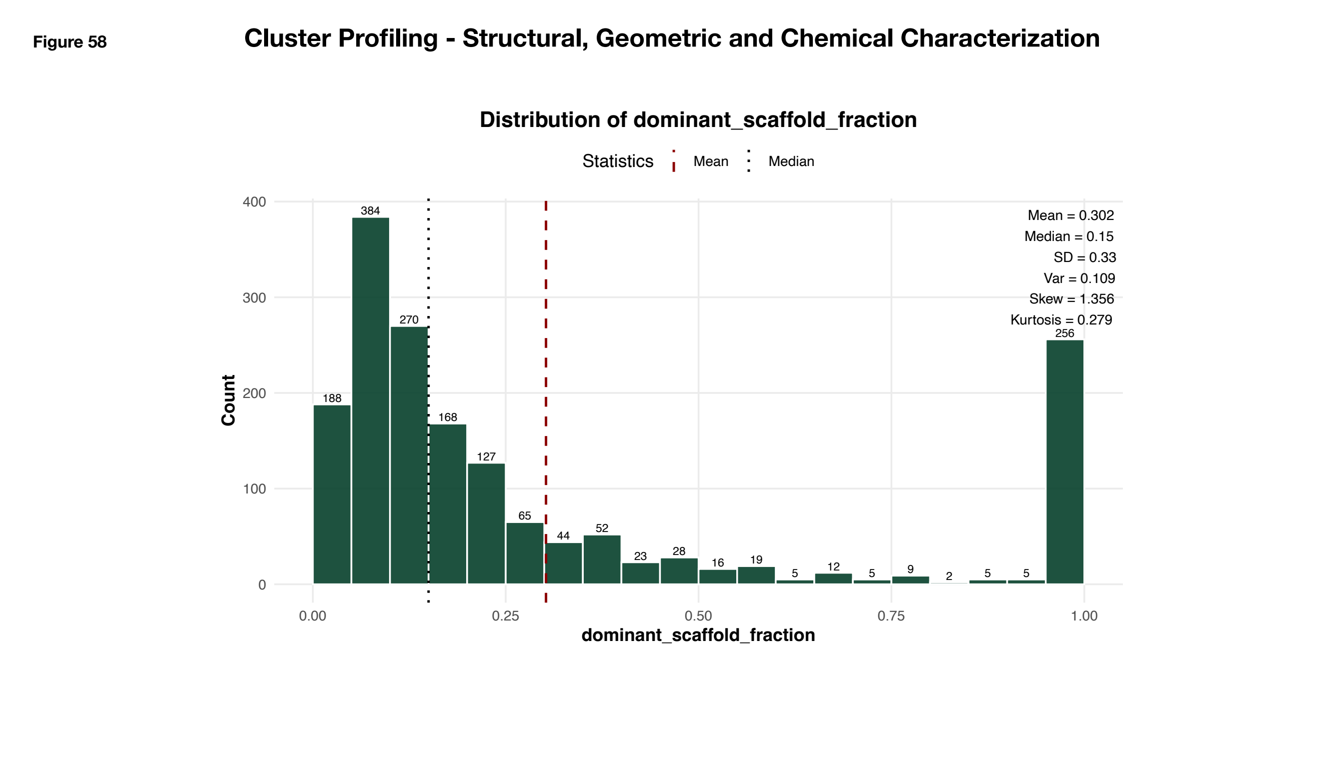


**Supplementary Figure 58: Distribution of Dominant Scaffold Fraction**

The histogram illustrates the distribution of dominant scaffold fraction (DSF) values across clusters. The x-axis represents DSF values grouped into predefined bins, while the y-axis indicates the number of clusters within each bin. Bar annotations denote the corresponding cluster counts. Summary statistics, including the mean, median, standard deviation, variance, skewness, and kurtosis, are displayed alongside the plot. The red dashed line marks the mean DSF value, while the black dotted line indicates the median, providing a visual summary of the distribution's central tendency and shape.

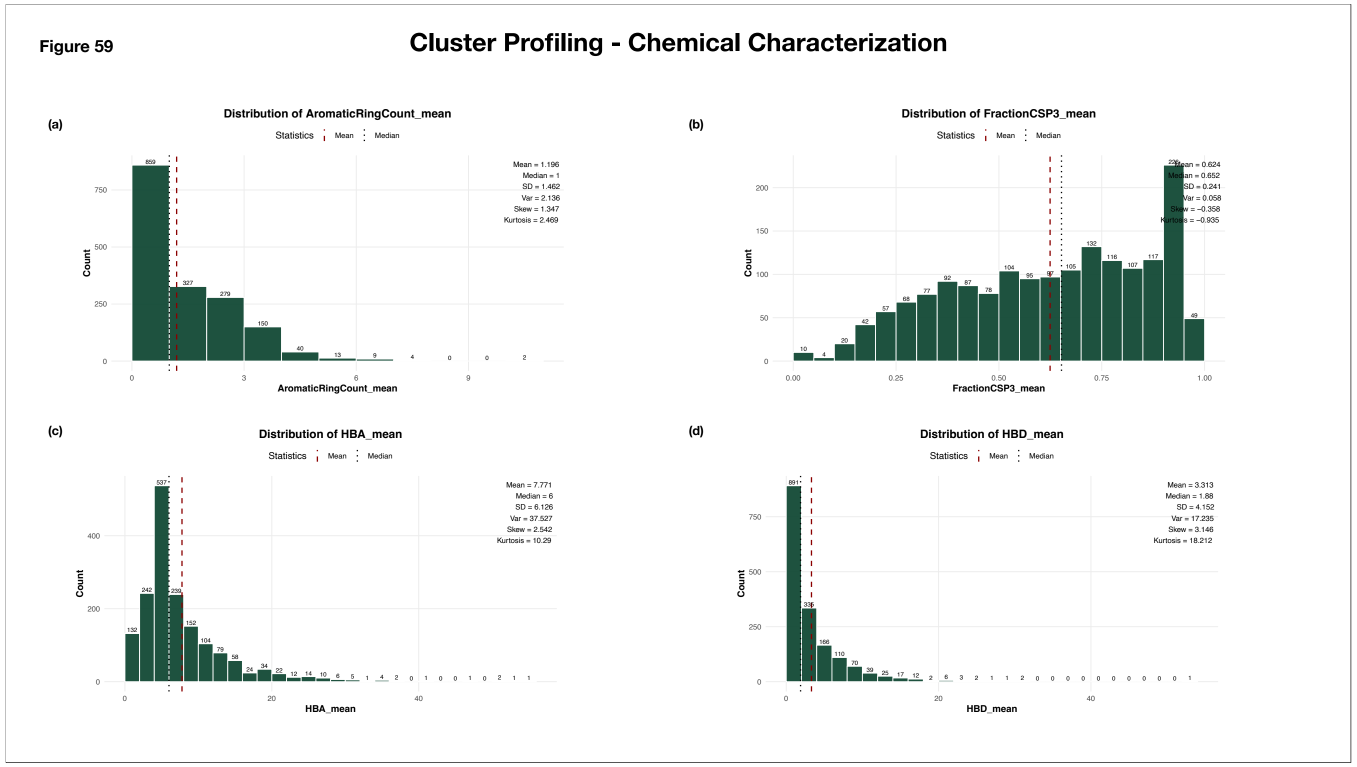


**Supplementary Figure 59: Chemical Characterization of Clusters - Distribution of mean descriptor values of Cluster Molecules**

The histogram illustrates the distribution of mean descriptor values across clusters. The x-axis represents the mean descriptor values grouped into predefined bins, while the y-axis indicates the number of clusters within each bin. Bar annotations denote the corresponding cluster counts. Summary statistics, including the mean, median, standard deviation, variance, skewness, and kurtosis, are displayed alongside the plot. The red dashed line marks the mean value, while the black dotted line indicates the median, providing a visual summary of the distribution's central tendency and variability.

(a) Mean Aromatic Ring Count

(b) Mean Fraction CSP3

(c) Mean HBA (Hydrogen Bond Acceptor) Count

(d) Mean HBD (Hydrogen Bond Donor) Count

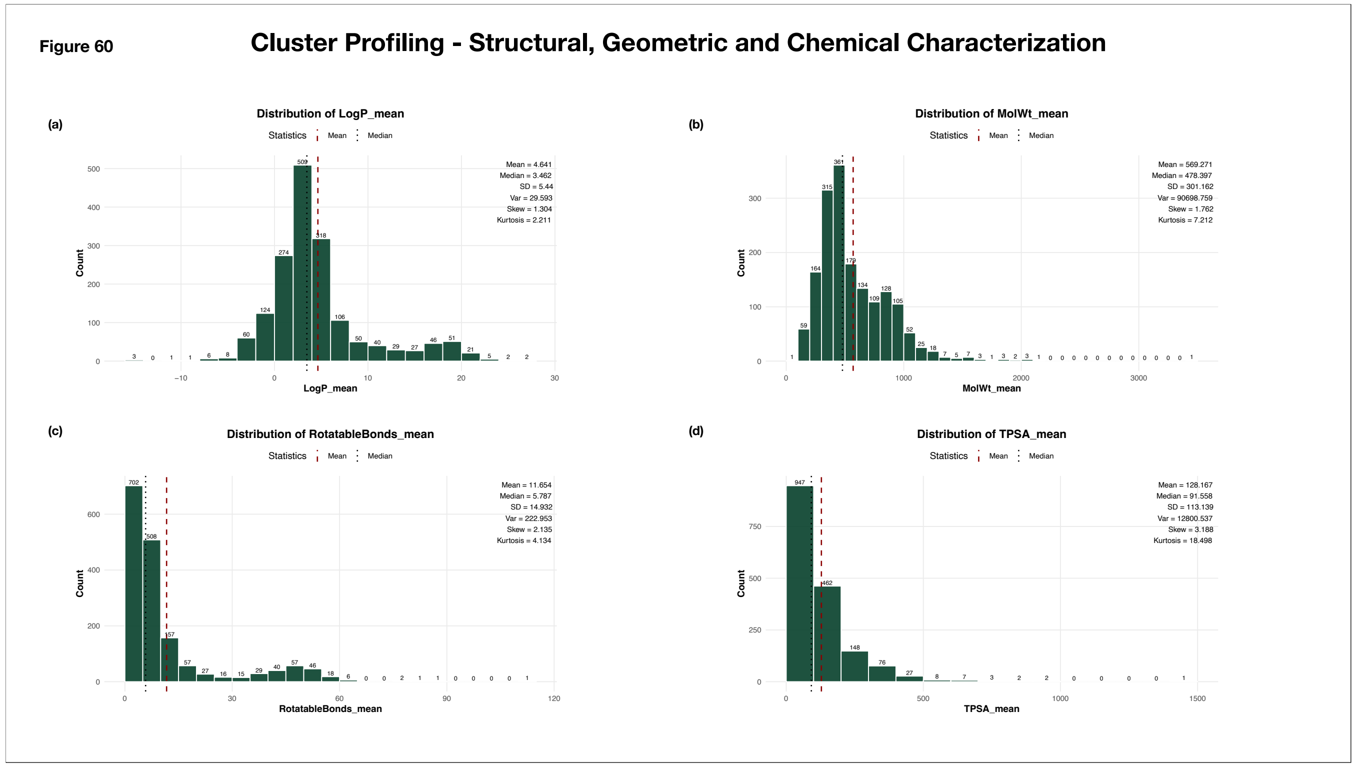

**Supplementary Figure 60: Chemical Characterization of Clusters - Distribution of mean descriptor values of Cluster Molecules**
The histogram illustrates the distribution of mean descriptor values across clusters. The x-axis represents the mean descriptor values grouped into predefined bins, while the y-axis indicates the number of clusters within each bin. Bar annotations denote the corresponding cluster counts. Summary statistics, including the mean, median, standard deviation, variance, skewness, and kurtosis, are displayed alongside the plot. The red dashed line marks the mean value, while the black dotted line indicates the median, providing a visual summary of the distribution's central tendency and variability.
(a) Mean LogP
(b) Mean Molecular Weight
(c) Mean Rotatable Bonds
(d) Mean TPSA (Total Polar Surface Area)

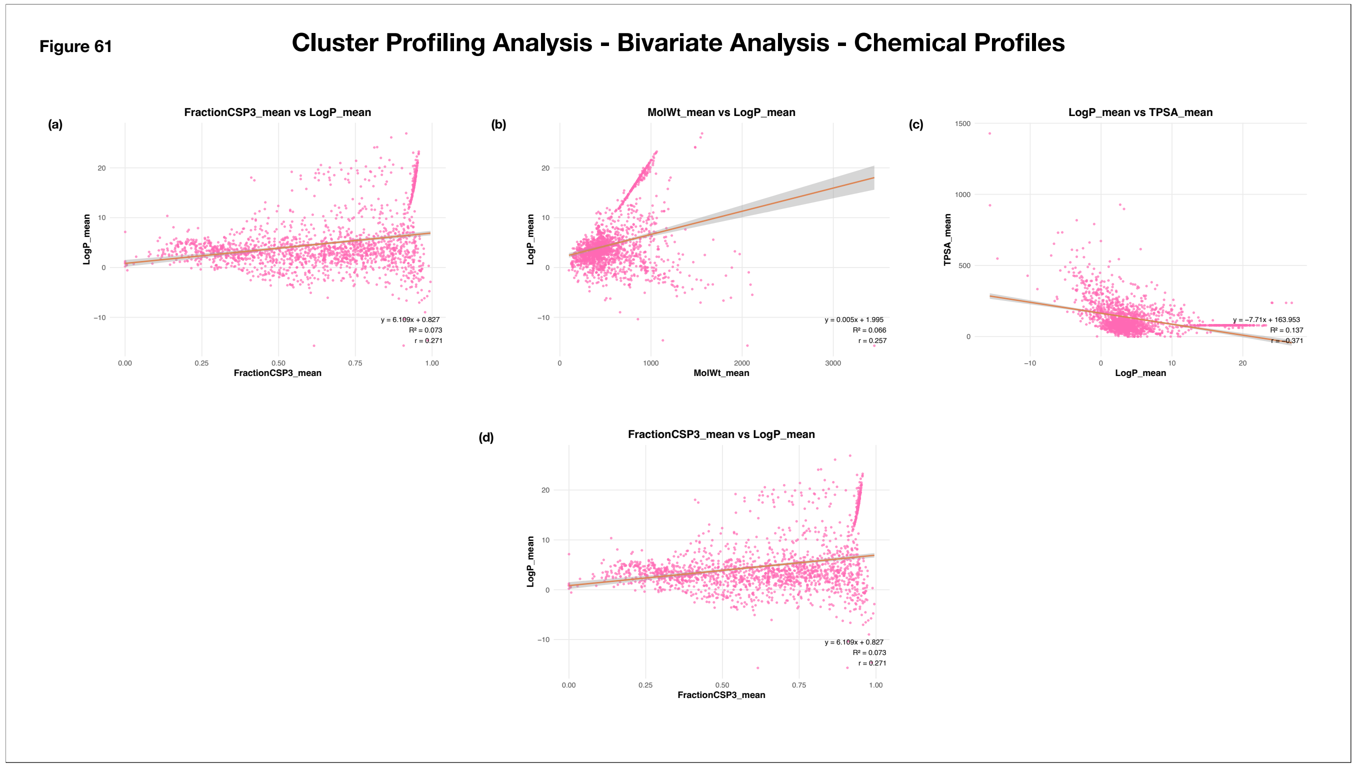


**Supplementary Figure 61: Bivariate Analysis of Mean Descriptor Values across Clusters**

The scatter plots illustrate the relationships between pairs of mean descriptor values across clusters. The x- and y-axes represent the descriptor values being compared, and each pink point corresponds to an individual cluster. A red regression line is overlaid to highlight the linear trend between the variables. The coefficient of determination ($R^2$), Pearson correlation coefficient (r), and regression equation are annotated in the lower-right corner of each plot.

(a) Mean LogP v/s Mean Fraction CSP3

(b) Mean LogP v/s Mean LogP

(c) Mean TPSA v/s Mean LogP

(d) Mean LogP v/s Mean Fraction CSP3

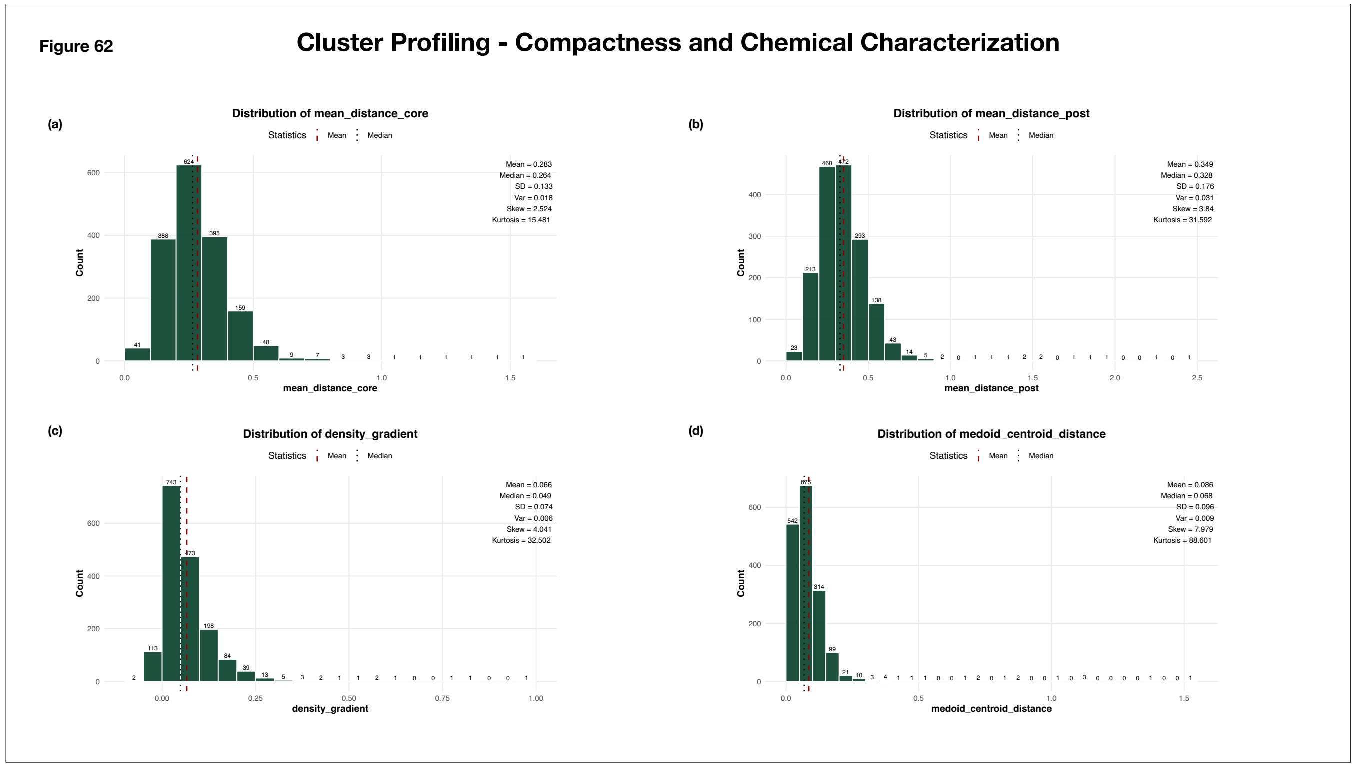


**Supplementary Figure 62: Distribution of Cluster Compactness Metrics**
The histograms illustrate the distributions of cluster compactness and characterization metrics across all clusters. The x-axis represents the observed values grouped into predefined bins, while the y-axis indicates the number of clusters within each bin. Bar annotations denote the corresponding cluster counts. Summary statistics, including the mean, median, standard deviation, variance, skewness, and kurtosis, are displayed alongside each plot. The red dashed line marks the mean value, while the black dotted line indicates the median, providing a visual summary of the central tendency, variability, and overall distribution of each metric.
(a) Distribution of mean distance (core clusters)
(b) Distribution of mean distance (post noise-reassignment)
(c) Distribution of density gradient
(d) Distribution of medoid-centroid distance

Figure 63

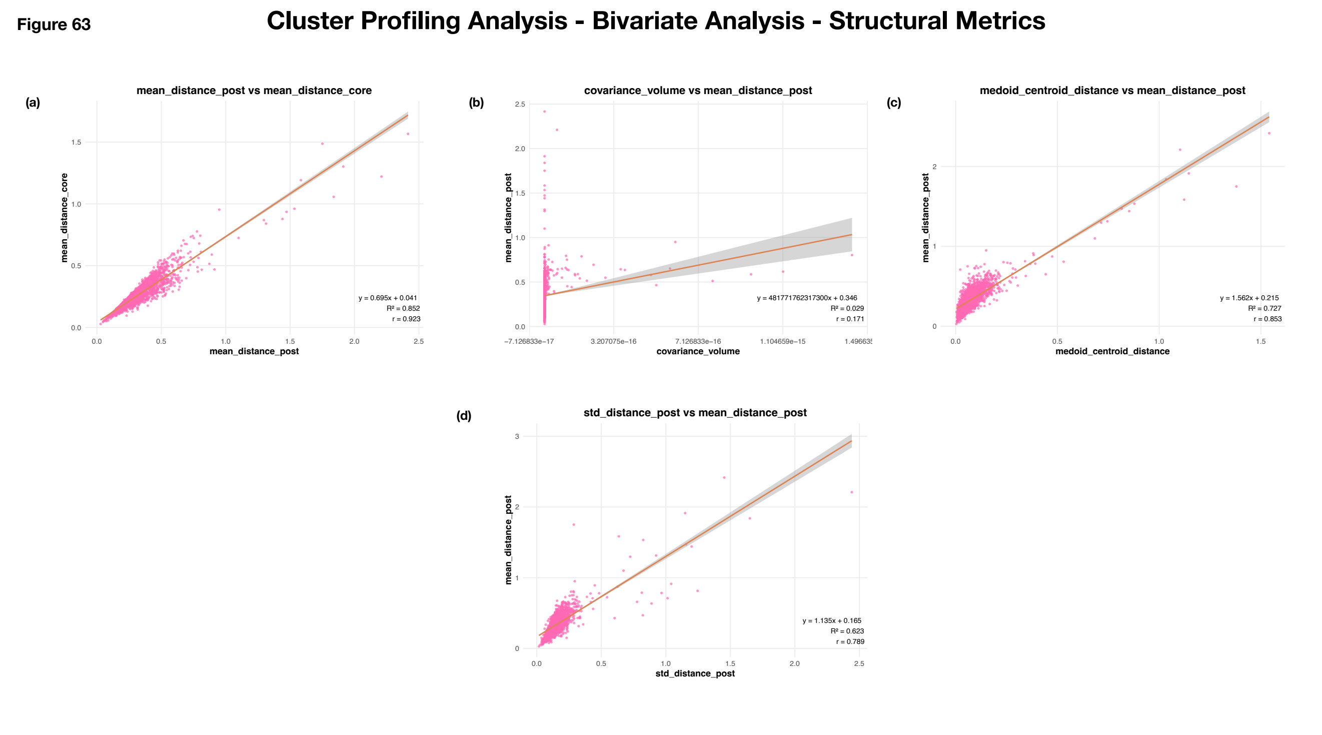


**Supplementary Figure 63: Bivariate Analysis of Structural Metrics across Clusters**

The scatter plots illustrate the relationships between pairs of structural metrics across clusters. The x- and y-axes represent the metrics being compared, and each pink point corresponds to an individual cluster. A red regression line is overlaid to highlight the linear trend between the variables. The coefficient of determination ($R^2$), Pearson correlation coefficient (r), and regression equation are annotated in the lower-right corner of each plot.

(a) Mean distance (core clusters) v/s mean distance (post-noise reassignment)
(b) Mean distance (post-noise reassignment) vs Covariance volume
(c) Mean distance (post-noise reassignment) vs Medoid-centroid distance
(d) Mean distance (post-noise reassignment) v/s Standard deviation of distance (post-noise reassignment)

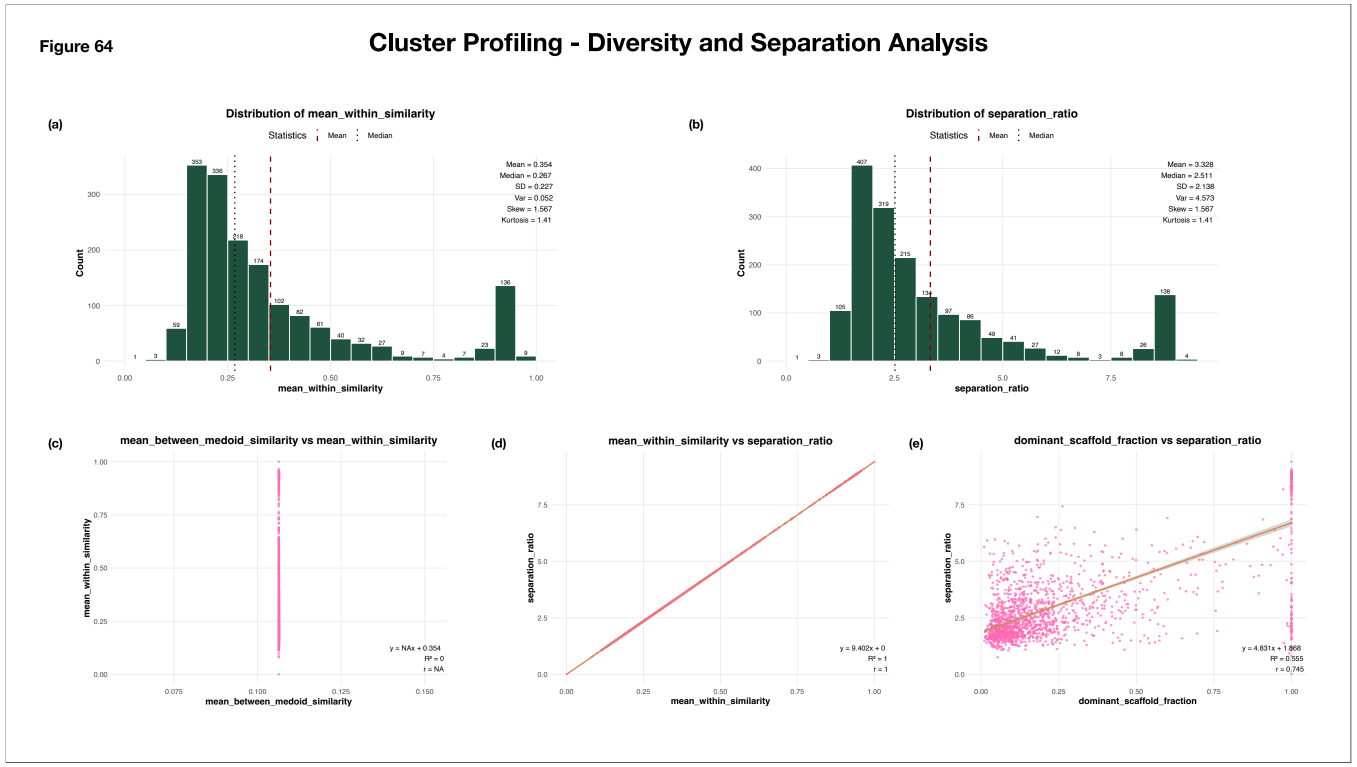


**Supplementary Figure 64: Distribution and Correlations of Diversity and Separation Metrics**

The histograms illustrate the distributions of diversity and separation metrics across all clusters. The x-axis represents the observed values grouped into predefined bins, while the y-axis indicates the number of clusters within each bin. Bar annotations denote the corresponding cluster counts. Summary statistics, including the mean, median, standard deviation, variance, skewness, and kurtosis, are displayed alongside each plot. The red dashed line marks the mean value, while the black dotted line indicates the median, providing a visual summary of the central tendency, variability, and overall distribution of each metric.

(a) Distribution of Mean Within Cluster Jaccard Similarity

(b) Distribution of Separation Ratio

The scatter plots illustrate the relationships between pairs of diversity, separation and chemical characteristic metrics across clusters. The x- and y-axes represent the metrics being compared, and each pink point corresponds to an individual cluster. A red regression line is overlaid to highlight the linear trend between the variables. The coefficient of determination ($R^2$), Pearson correlation coefficient (r), and regression equation are annotated in the lower-right corner of each plot.

(c) Mean-Within Cluster Jaccard Similarity v/s Mean-Between-Medoid Similarity

(d) Separation Ratio v/s Mean-Within Cluster Jaccard Similarity

(e) Separation Ratio v/s Dominant Scaffold Fraction

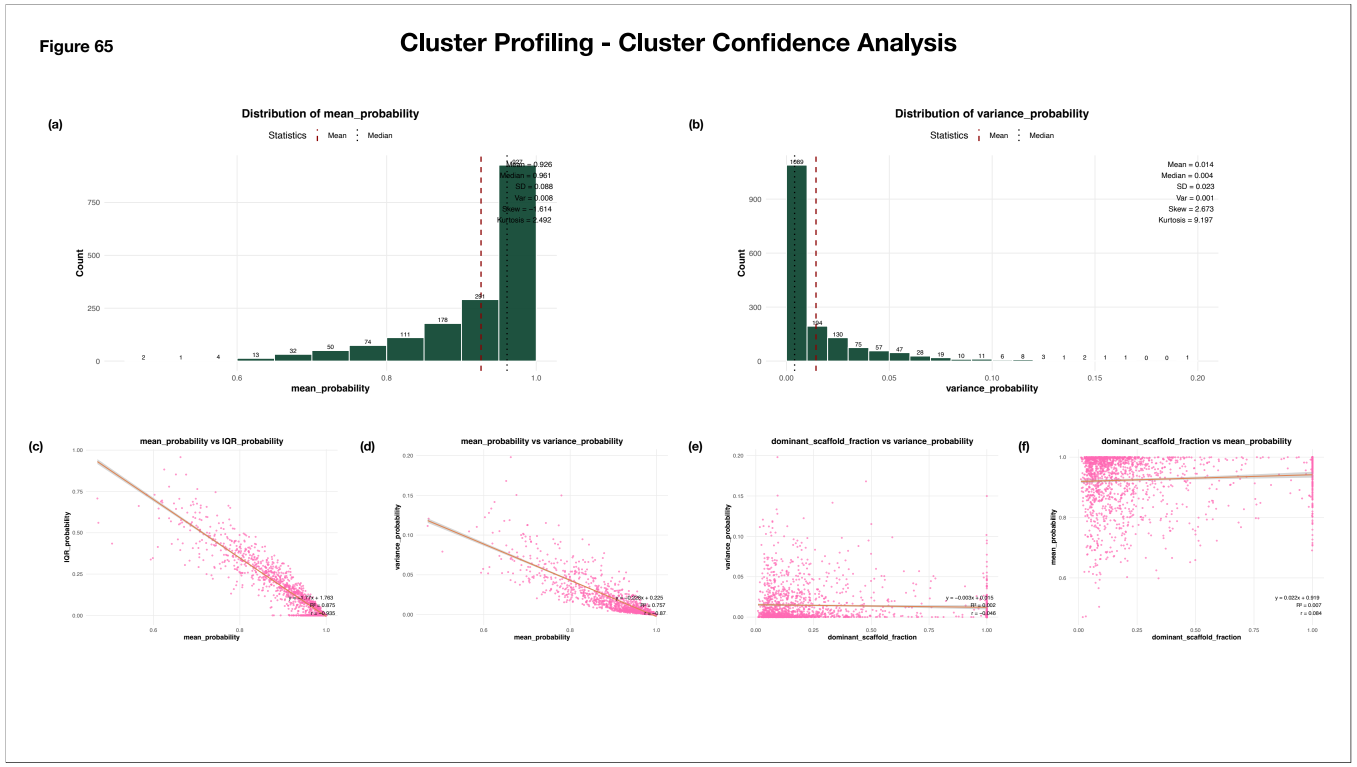


**Supplementary Figure 65: Distribution and Correlations of Cluster Confidence Metrics**

The histograms illustrate the distributions of cluster confidence metrics across all clusters. The x-axis represents the observed values grouped into predefined bins, while the y-axis indicates the number of clusters within each bin. Bar annotations denote the corresponding cluster counts. Summary statistics, including the mean, median, standard deviation, variance, skewness, and kurtosis, are displayed alongside each plot. The red dashed line marks the mean value, while the black dotted line indicates the median, providing a visual summary of the central tendency, variability, and overall distribution of each metric.

(a) Distribution of Mean HDBSCAN Assignment Probability

(b) Distribution of Variance of HDBSCAN Assignment Probability

The scatter plots illustrate the relationships between pairs of assignment confidence and chemical characteristic metrics across clusters. The x- and y-axes represent the metrics being compared, and each pink point corresponds to an individual cluster. A red regression line is overlaid to highlight the linear trend between the variables. The coefficient of determination ($R^2$), Pearson correlation coefficient (r), and regression equation are annotated in the lower-right corner of each plot.

(c) IQR (HDBSCAN probability assignment) v/s Mean (HDBSCAN probability assignment)

(d) Variance (HDBSCAN probability assignment) v/s Mean (HDBSCAN probability assignment)

(e) Variance (HDBSCAN probability assignment) v/s Dominant Scaffold Fraction

(f) Mean (HDBSCAN probability assignment) v/s Dominant Scaffold Fraction

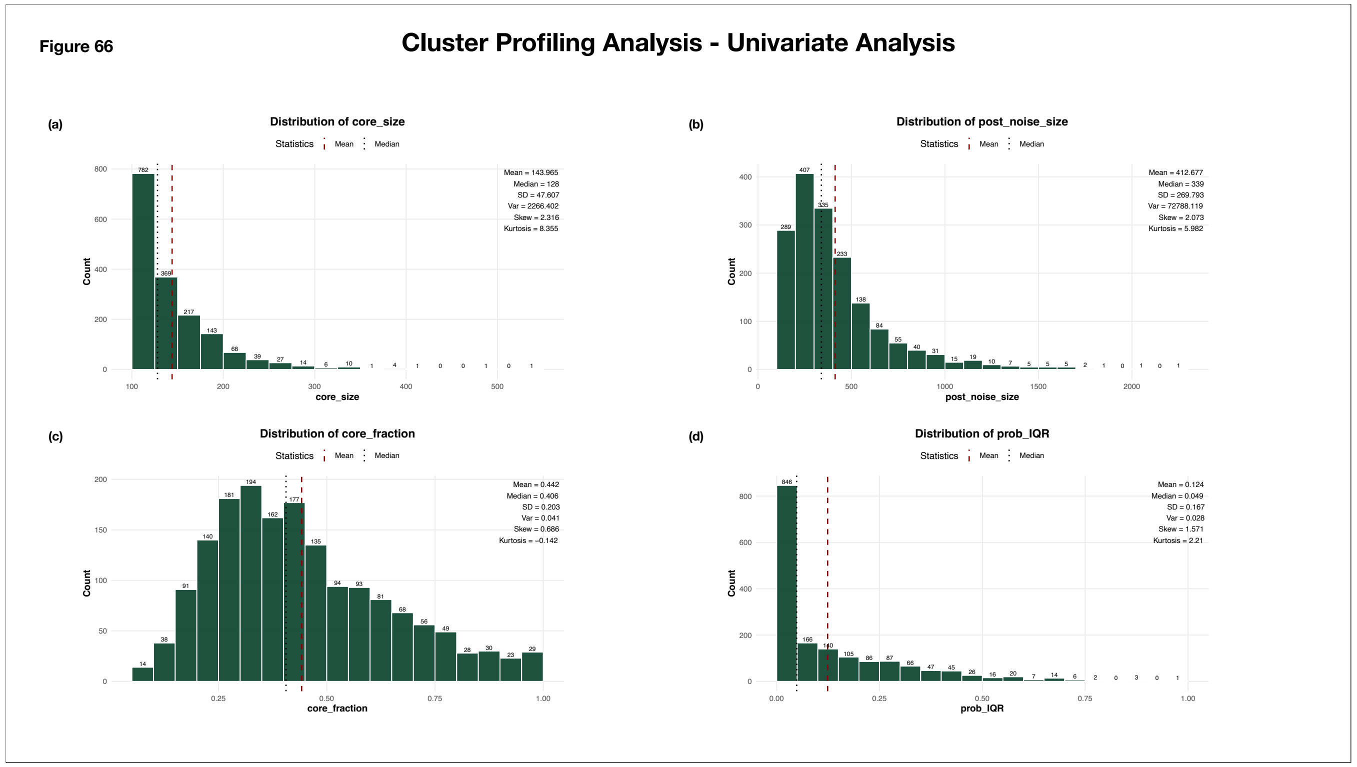


**Supplementary Figure 66: Distribution of Cluster Fidelity Metrics**

The histograms illustrate the distributions of cluster fidelity metrics across all clusters. The x-axis represents the observed values grouped into predefined bins, while the y-axis indicates the number of clusters within each bin. Bar annotations denote the corresponding cluster counts. Summary statistics, including the mean, median, standard deviation, variance, skewness, and kurtosis, are displayed alongside each plot. The red dashed line marks the mean value, while the black dotted line indicates the median, providing a visual summary of the central tendency, variability, and overall distribution of each metric.

(a) Distribution of cluster core size
(b) Distribution of post-noise assignment cluster size
(c) Distribution of core fraction
(d) Distribution of IQR of HDBSCAN probability assignments

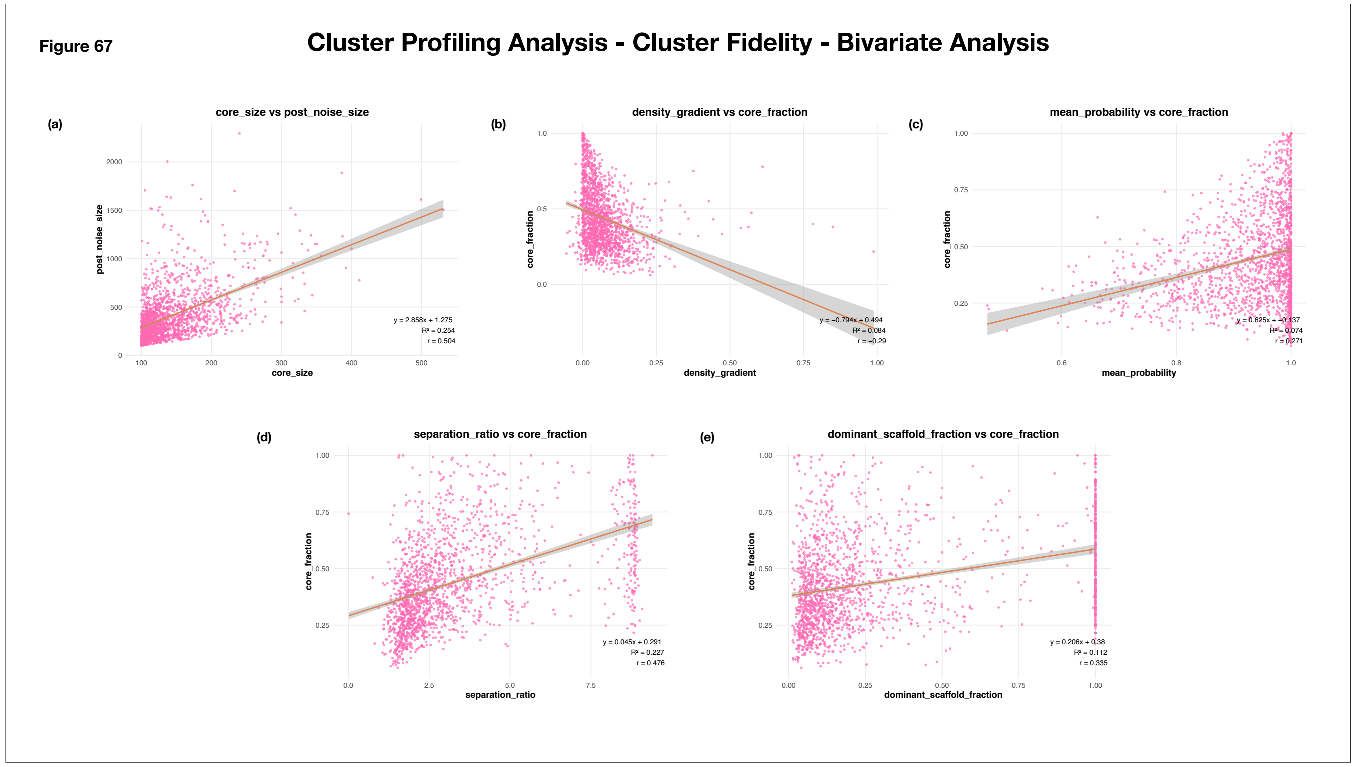


**Supplementary Figure 67: Bivariate Analysis of Fidelity, Separation, Assignment Probability and Chemical Characteristics across Clusters**

The scatter plots illustrate the relationships between pairs of fidelity, separation, assignment probability and chemical characterization metrics across clusters. The x- and y-axes represent the metrics being compared, and each pink point corresponds to an individual cluster. A red regression line is overlaid to highlight the linear trend between the variables. The coefficient of determination ($R^2$), Pearson correlation coefficient (r), and regression equation are annotated in the lower-right corner of each plot.

(a) Cluster size (post-noise assignment) v/s Cluster size (core cluster)
(b) Core fraction v/s Density Gradient
(c) Core fraction v/s Mean HDBSCAN Membership Probability
(d) Core fraction v/s Separation Ratio
(e) Core fraction v/s Dominant Scaffold Fraction

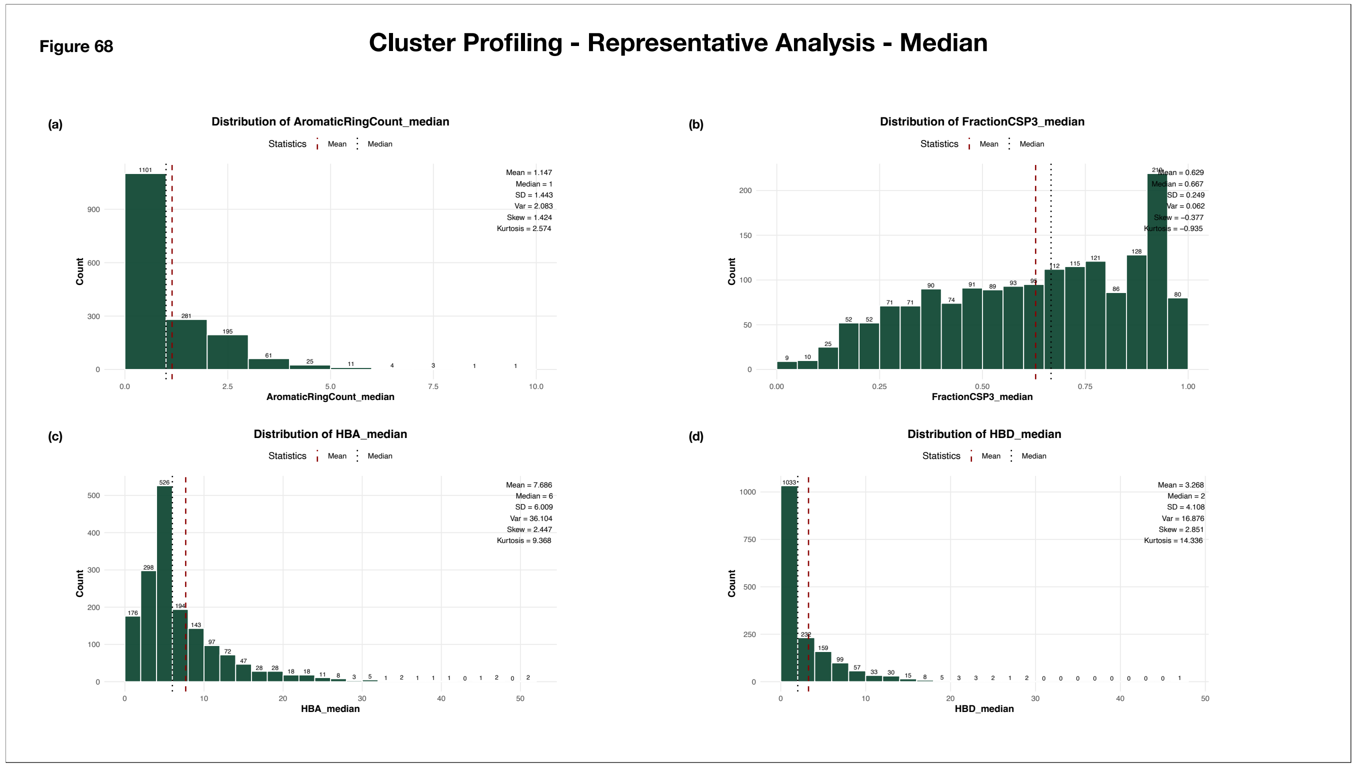


**Supplementary Figure 68: Distribution of Median Descriptor Values of Cluster Molecules**

The histograms illustrate the distributions of median descriptor values of cluster molecules across all clusters. The x-axis represents the observed values grouped into predefined bins, while the y-axis indicates the number of clusters within each bin. Bar annotations denote the corresponding cluster counts. Summary statistics, including the mean, median, standard deviation, variance, skewness, and kurtosis, are displayed alongside each plot. The red dashed line marks the mean value, while the black dotted line indicates the median, providing a visual summary of the central tendency, variability, and overall distribution of each metric.

(a) Distribution of Median Aromatic Ring Count
(b) Distribution of Median Fraction CSP3
(c) Distribution of Median HBA
(d) Distribution of Median HBD

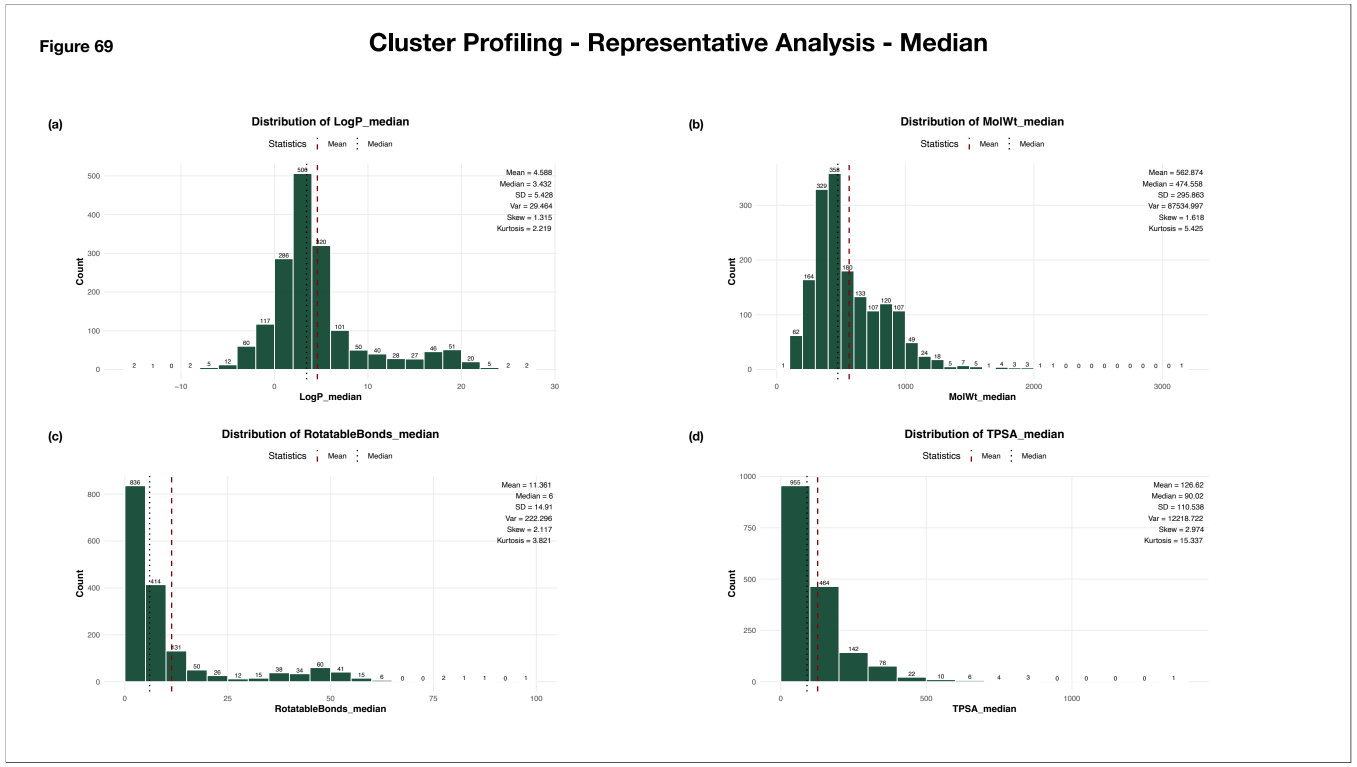


**Supplementary Figure 69: Distribution of Median Descriptor Values of Cluster Molecules**

The histograms illustrate the distributions of median descriptor values of cluster molecules across all clusters. The x-axis represents the observed values grouped into predefined bins, while the y-axis indicates the number of clusters within each bin. Bar annotations denote the corresponding cluster counts. Summary statistics, including the mean, median, standard deviation, variance, skewness, and kurtosis, are displayed alongside each plot. The red dashed line marks the mean value, while the black dotted line indicates the median, providing a visual summary of the central tendency, variability, and overall distribution of each metric.

(a) Distribution of Median LogP
(b) Distribution of Median Molecular Weight
(c) Distribution of Median Rotatable Bonds
(d) Distribution of Median TPSA

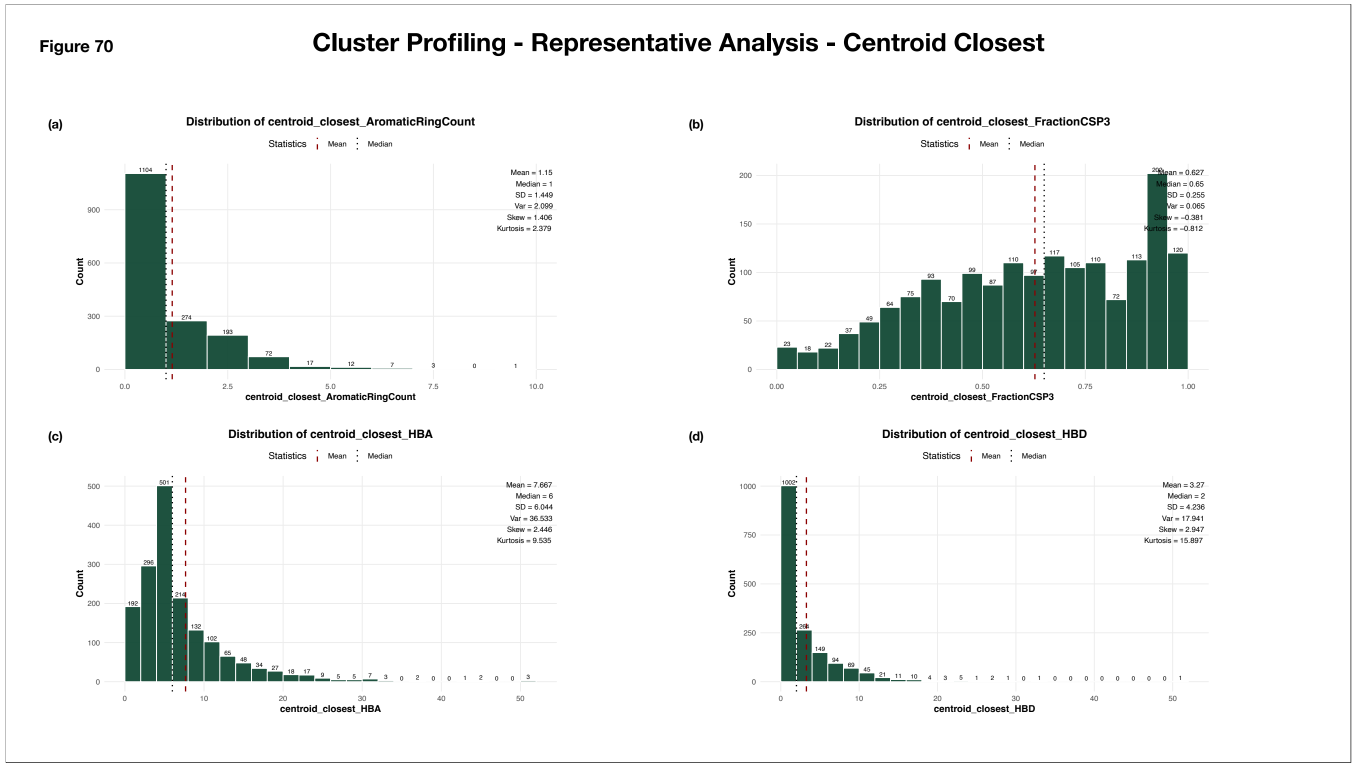


**Supplementary Figure 70: Distribution of Descriptor-Based Properties of Centroid-Closest Representatives Across All Clusters**

The histograms illustrate the distributions of centroid-closest representative values across all clusters. The x-axis represents the observed values grouped into predefined bins, while the y-axis indicates the number of clusters within each bin. Bar annotations denote the corresponding cluster counts. Summary statistics, including the mean, median, standard deviation, variance, skewness, and kurtosis, are displayed alongside each plot. The red dashed line marks the mean value, while the black dotted line indicates the median, providing a visual summary of the central tendency, variability, and overall distribution of each metric.

(a) Distribution of Centroid_Closest Aromatic Ring Count
(b) Distribution of Centroid_Closest Fraction CSP3
(c) Distribution of Centroid_Closest HBA
(d) Distribution of Centroid_Closest HBD

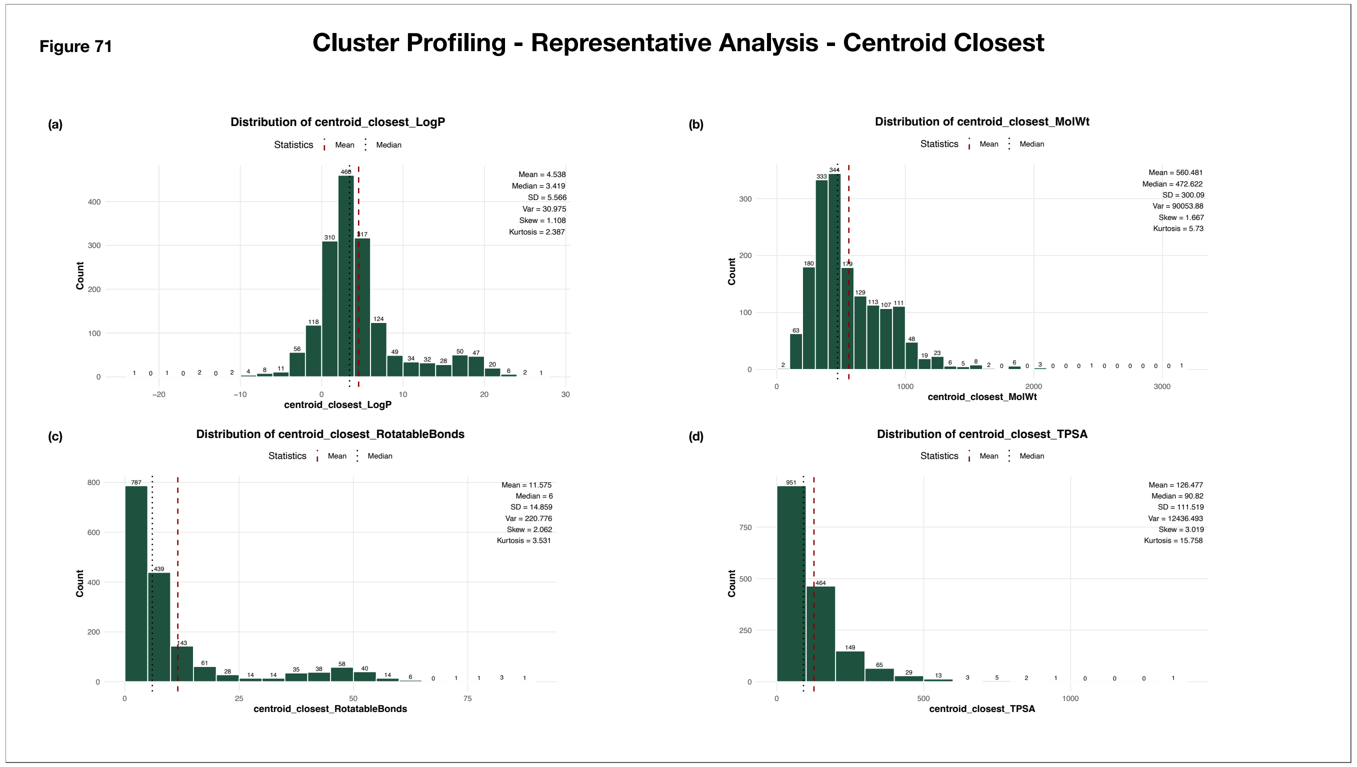


**Supplementary Figure 71: Distribution of Descriptor-Based Properties of Centroid-Closest Representatives Across All Clusters**

The histograms illustrate the distributions of centroid-closest representative values across all clusters. The x-axis represents the observed values grouped into predefined bins, while the y-axis indicates the number of clusters within each bin. Bar annotations denote the corresponding cluster counts. Summary statistics, including the mean, median, standard deviation, variance, skewness, and kurtosis, are displayed alongside each plot. The red dashed line marks the mean value, while the black dotted line indicates the median, providing a visual summary of the central tendency, variability, and overall distribution of each metric.

(a) Distribution of Centroid_Closest LogP
(b) Distribution of Centroid_Closest Molecular Weight
(c) Distribution of Centroid_Closest Rotatable Bonds
(d) Distribution of Centroid_Closest TPSA

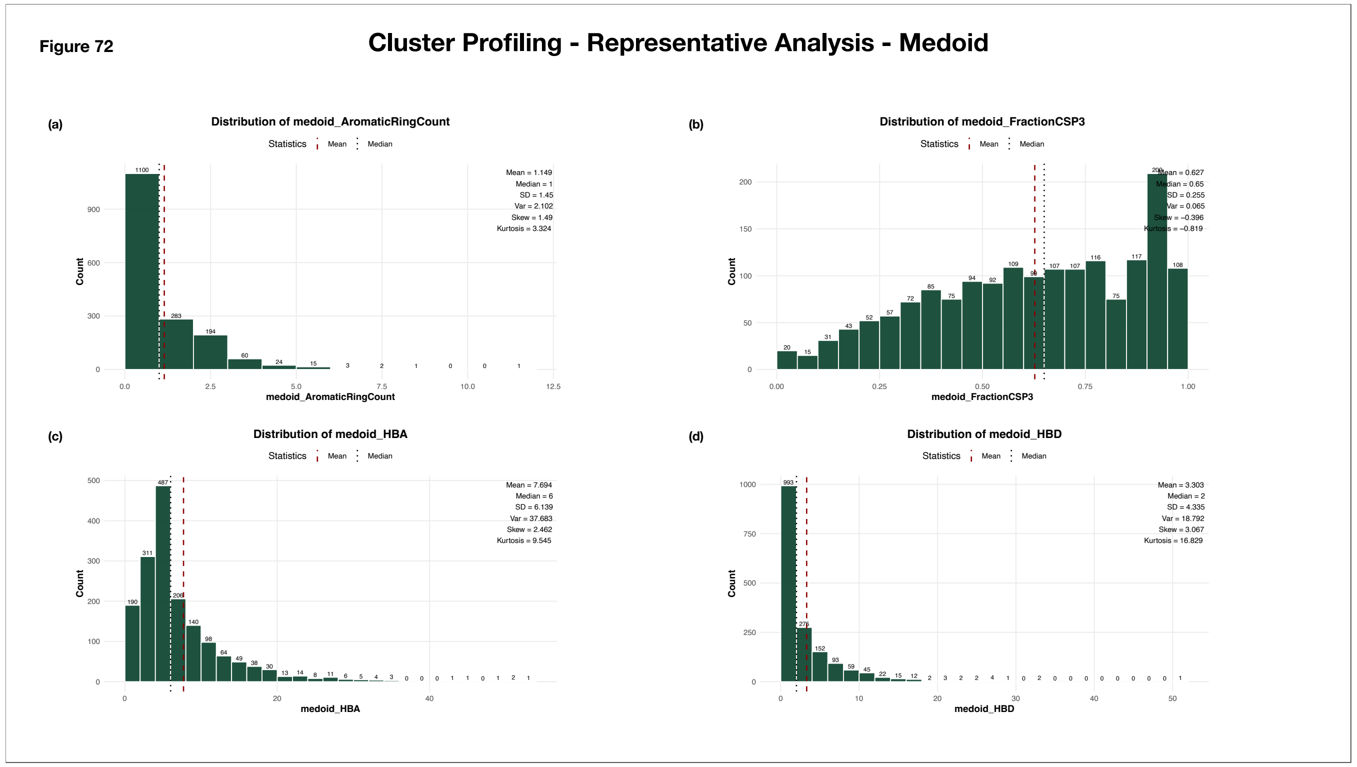


**Supplementary Figure 72: Distribution of Descriptor-Based Properties of Medoid Representatives Across All Clusters**

The histograms illustrate the distributions of medoid representative values across all clusters. The x-axis represents the observed values grouped into predefined bins, while the y-axis indicates the number of clusters within each bin. Bar annotations denote the corresponding cluster counts. Summary statistics, including the mean, median, standard deviation, variance, skewness, and kurtosis, are displayed alongside each plot. The red dashed line marks the mean value, while the black dotted line indicates the median, providing a visual summary of the central tendency, variability, and overall distribution of each metric.

(a) Distribution of Medoid Aromatic Ring Count
(b) Distribution of Medoid Fraction CSP3
(c) Distribution of Media HBA
(d) Distribution of Medoid HBD

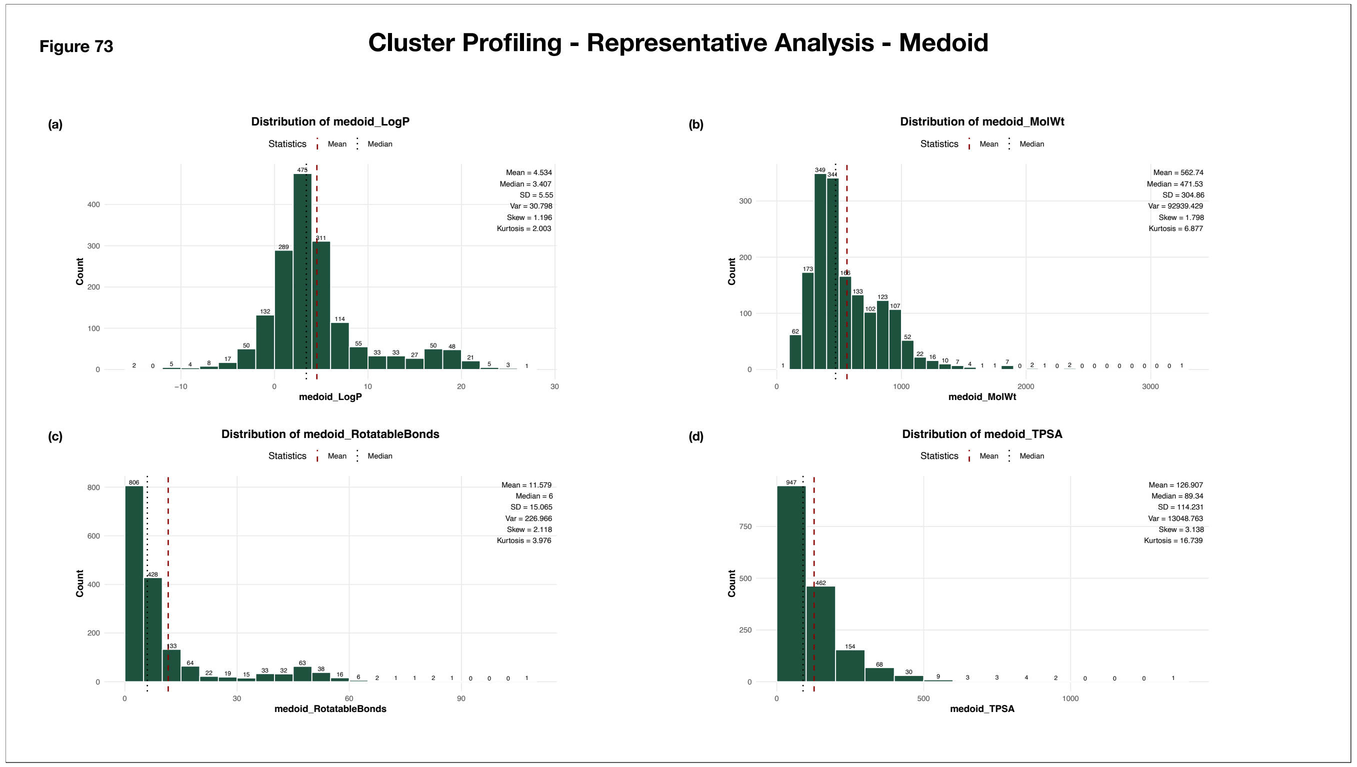


**Supplementary Figure 73: Distribution of Descriptor-Based Properties of Medoid Representatives Across All Clusters**

The histograms illustrate the distributions of medoid representative values across all clusters. The x-axis represents the observed values grouped into predefined bins, while the y-axis indicates the number of clusters within each bin. Bar annotations denote the corresponding cluster counts. Summary statistics, including the mean, median, standard deviation, variance, skewness, and kurtosis, are displayed alongside each plot. The red dashed line marks the mean value, while the black dotted line indicates the median, providing a visual summary of the central tendency, variability, and overall distribution of each metric.

(a) Distribution of Medoid LogP
(b) Distribution of Medoid Molecular Weight
(c) Distribution of Medoid Rotatable Bonds
(d) Distribution of Medoid TPSA

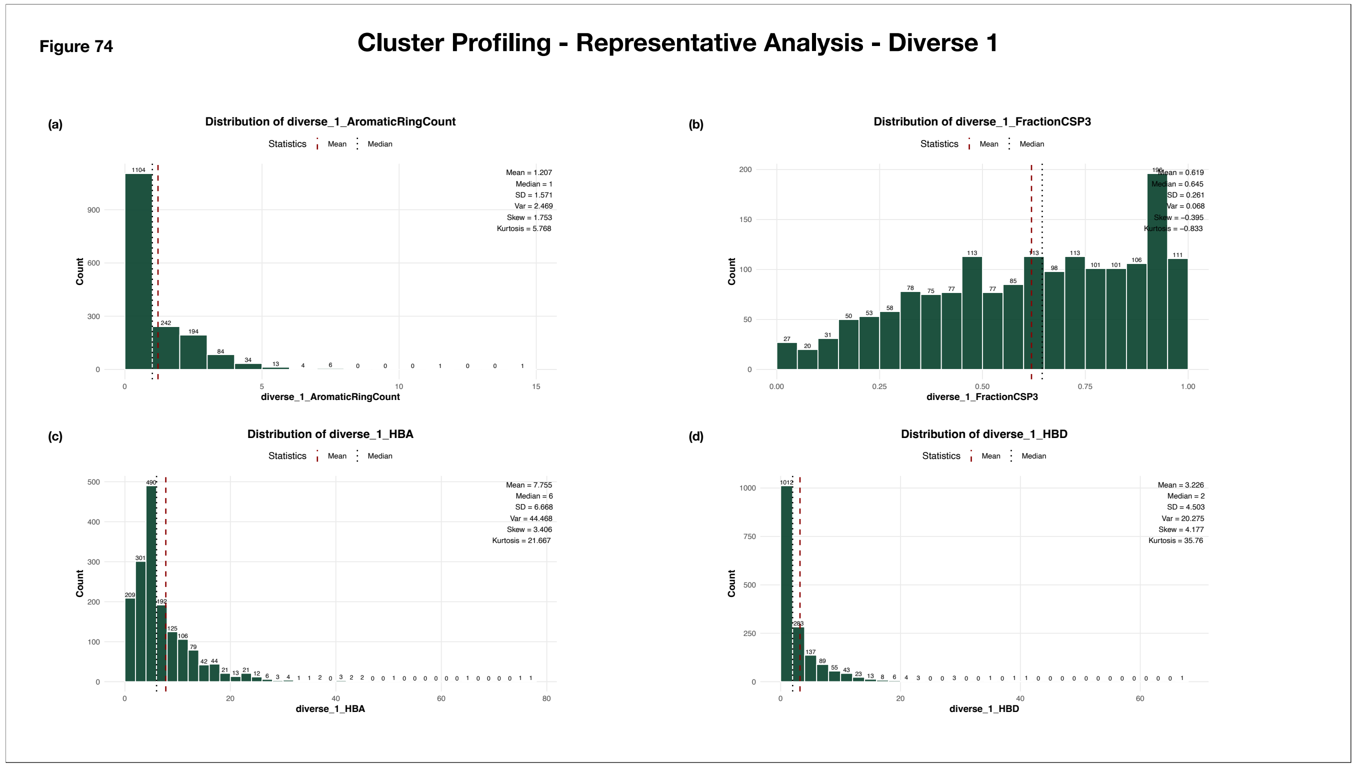


**Supplementary Figure 74: Distribution of Descriptor-Based Properties of Diverse_1 Representatives Across All Clusters**

The histograms illustrate the distributions of diverse_1 representative values across all clusters. The x-axis represents the observed values grouped into predefined bins, while the y-axis indicates the number of clusters within each bin. Bar annotations denote the corresponding cluster counts. Summary statistics, including the mean, median, standard deviation, variance, skewness, and kurtosis, are displayed alongside each plot. The red dashed line marks the mean value, while the black dotted line indicates the median, providing a visual summary of the central tendency, variability, and overall distribution of each metric.

(a) Distribution of Diverse_1 Aromatic Ring Count
(b) Distribution of Diverse_1 Fraction CSP3
(c) Distribution of Diverse_1 HBA
(d) Distribution of Diverse_1 HBD

Figure 75

### Cluster Profiling - Representative Analysis - Diverse 1

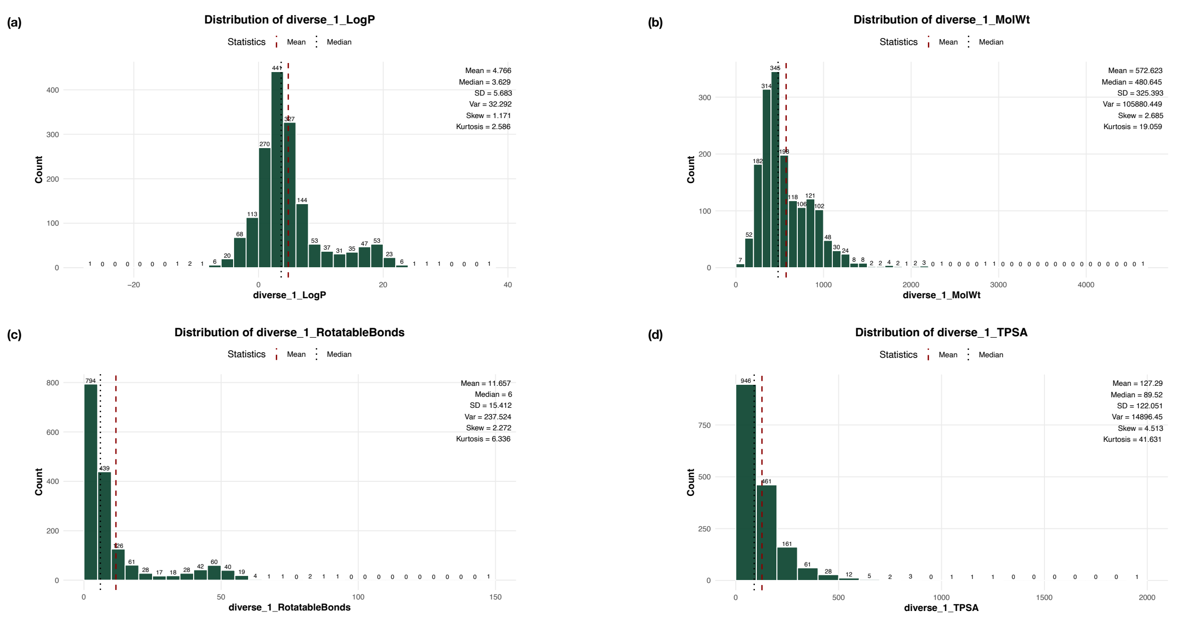


**Supplementary Figure 75: Distribution of Descriptor-Based Properties of Diverse_1 Representatives Across All Clusters**

The histograms illustrate the distributions of diverse_1 representative values across all clusters. The x-axis represents the observed values grouped into predefined bins, while the y-axis indicates the number of clusters within each bin. Bar annotations denote the corresponding cluster counts. Summary statistics, including the mean, median, standard deviation, variance, skewness, and kurtosis, are displayed alongside each plot. The red dashed line marks the mean value, while the black dotted line indicates the median, providing a visual summary of the central tendency, variability, and overall distribution of each metric.

(a) Distribution of Diverse_1 LogP
(b) Distribution of Diverse_1 Molecular Weight
(c) Distribution of Diverse_1 Rotatable Bonds
(d) Distribution of Diverse_1 TPSA

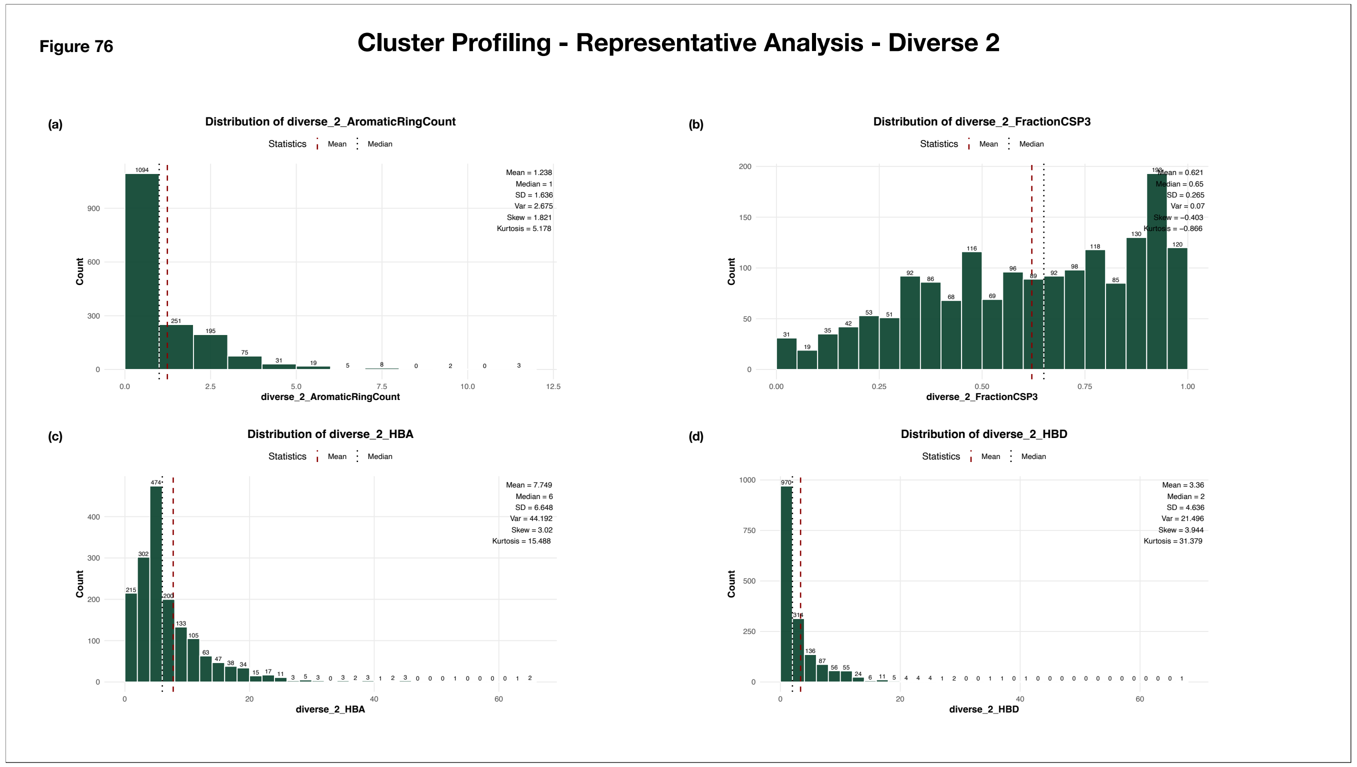


**Supplementary Figure 76: Distribution of Descriptor-Based Properties of Diverse_2 Representatives Across All Clusters**

The histograms illustrate the distributions of diverse_2 representative values across all clusters. The x-axis represents the observed values grouped into predefined bins, while the y-axis indicates the number of clusters within each bin. Bar annotations denote the corresponding cluster counts. Summary statistics, including the mean, median, standard deviation, variance, skewness, and kurtosis, are displayed alongside each plot. The red dashed line marks the mean value, while the black dotted line indicates the median, providing a visual summary of the central tendency, variability, and overall distribution of each metric.

(a) Distribution of Diverse_2 Aromatic Ring Count
(b) Distribution of Diverse_2 Fraction CSP3
(c) Distribution of Diverse_2 HBA
(d) Distribution of Diverse_2 HBD

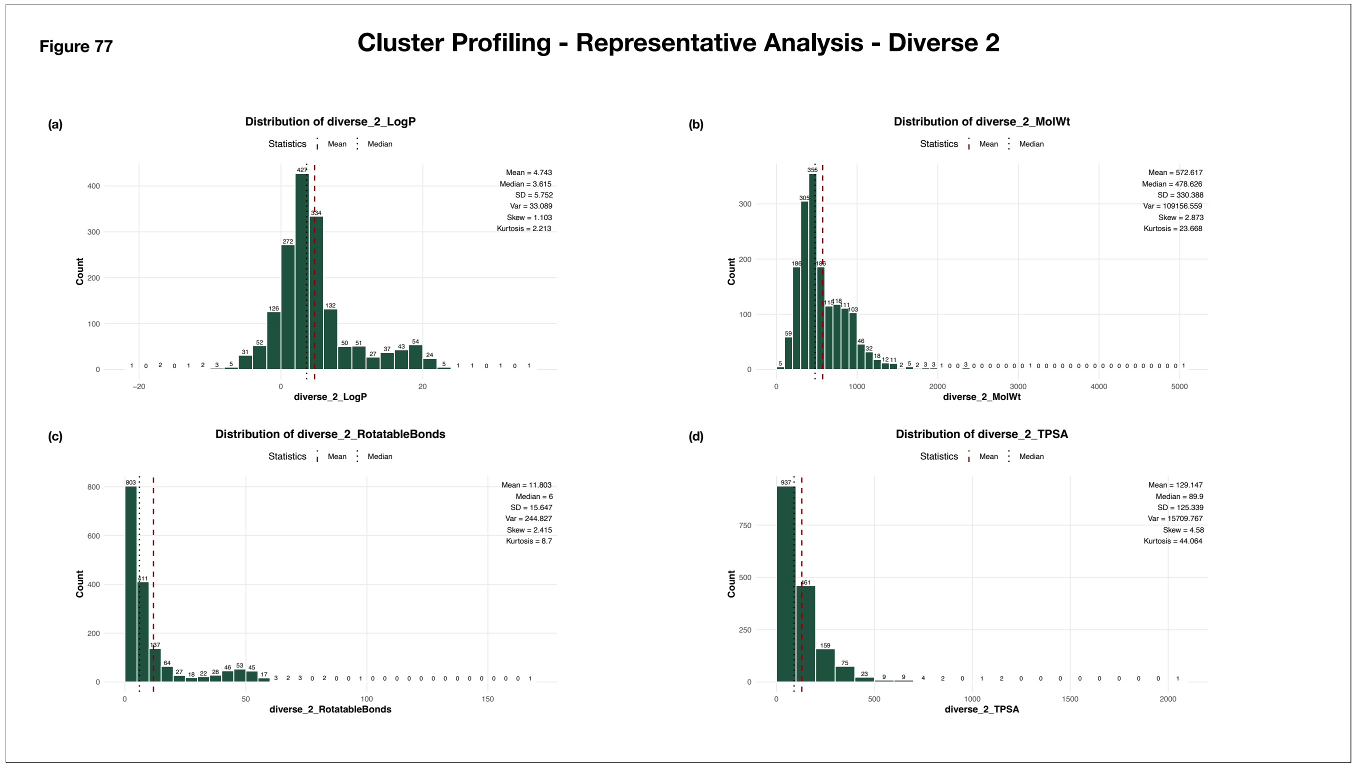


**Supplementary Figure 77: Distribution of Descriptor-Based Properties of Diverse_2 Representatives Across All Clusters**

The histograms illustrate the distributions of diverse_2 representative values across all clusters. The x-axis represents the observed values grouped into predefined bins, while the y-axis indicates the number of clusters within each bin. Bar annotations denote the corresponding cluster counts. Summary statistics, including the mean, median, standard deviation, variance, skewness, and kurtosis, are displayed alongside each plot. The red dashed line marks the mean value, while the black dotted line indicates the median, providing a visual summary of the central tendency, variability, and overall distribution of each metric.

(a) Distribution of Diverse_2 LogP
(b) Distribution of Diverse_2 Molecular Weight
(c) Distribution of Diverse_2 Rotatable Bonds
(d) Distribution of Diverse_2 TPSA

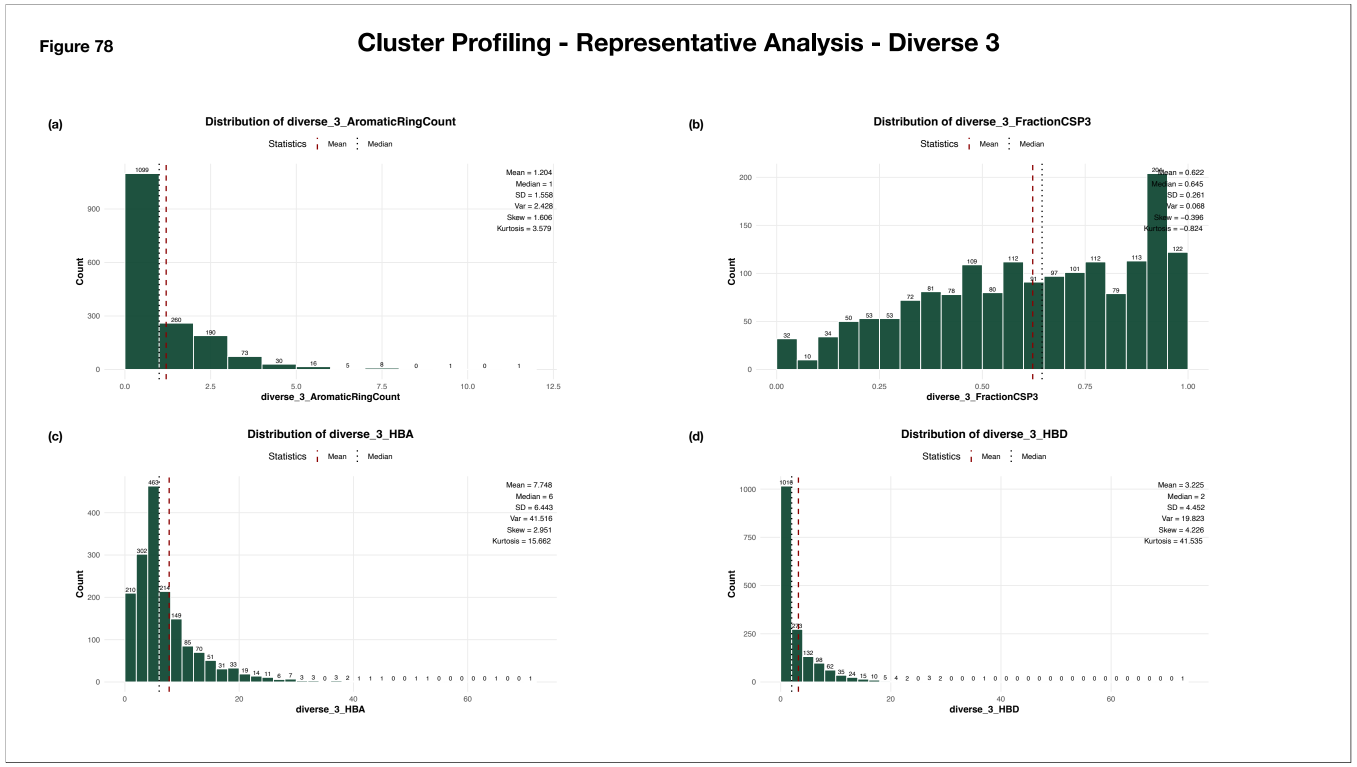


**Supplementary Figure 78: Distribution of Descriptor-Based Properties of Diverse_3 Representatives Across All Clusters**

The histograms illustrate the distributions of diverse_3 representative values across all clusters. The x-axis represents the observed values grouped into predefined bins, while the y-axis indicates the number of clusters within each bin. Bar annotations denote the corresponding cluster counts. Summary statistics, including the mean, median, standard deviation, variance, skewness, and kurtosis, are displayed alongside each plot. The red dashed line marks the mean value, while the black dotted line indicates the median, providing a visual summary of the central tendency, variability, and overall distribution of each metric.

(a) Distribution of Diverse_3 Aromatic Ring Count
(b) Distribution of Diverse_3 Fraction CSP3
(c) Distribution of Diverse_3 HBA
(d) Distribution of Diverse_3 HBD

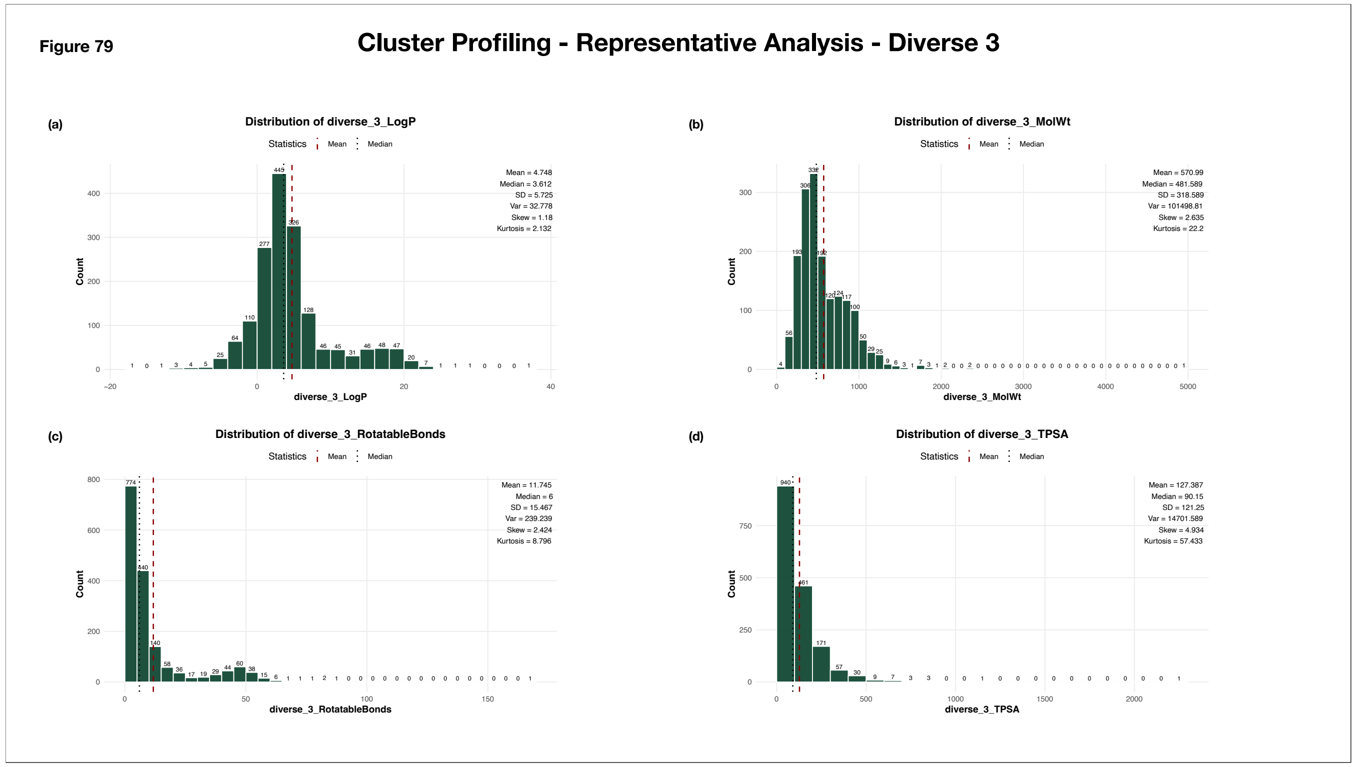


**Supplementary Figure 79: Distribution of Descriptor-Based Properties of Diverse_3 Representatives Across All Clusters**

The histograms illustrate the distributions of diverse_3 representative values across all clusters. The x-axis represents the observed values grouped into predefined bins, while the y-axis indicates the number of clusters within each bin. Bar annotations denote the corresponding cluster counts. Summary statistics, including the mean, median, standard deviation, variance, skewness, and kurtosis, are displayed alongside each plot. The red dashed line marks the mean value, while the black dotted line indicates the median, providing a visual summary of the central tendency, variability, and overall distribution of each metric.

(a) Distribution of Diverse_3 LogP
(b) Distribution of Diverse_3 Molecular Weight
(c) Distribution of Diverse_3 Rotatable Bonds
(d) Distribution of Diverse_3 TPSA

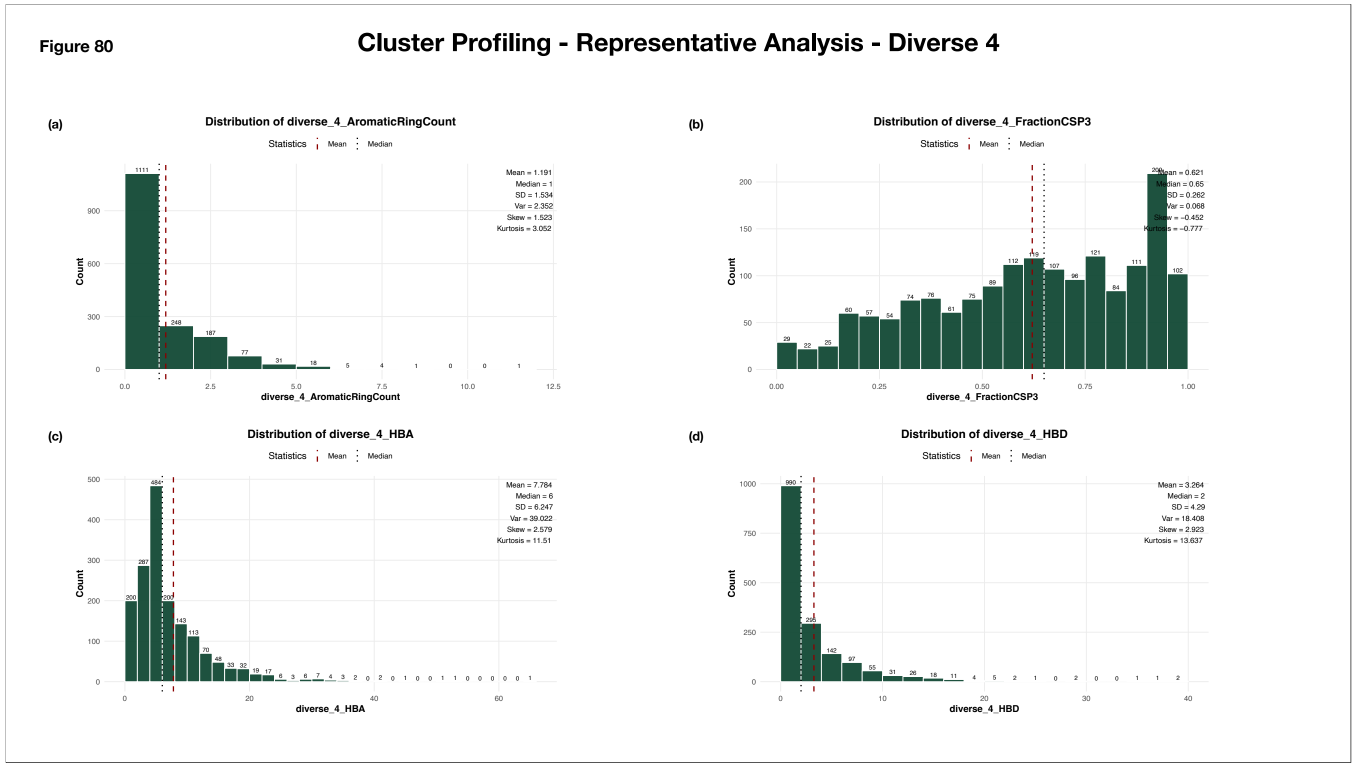


**Supplementary Figure 80: Distribution of Descriptor-Based Properties of Diverse_4 Representatives Across All Clusters**

The histograms illustrate the distributions of diverse_4 representative values across all clusters. The x-axis represents the observed values grouped into predefined bins, while the y-axis indicates the number of clusters within each bin. Bar annotations denote the corresponding cluster counts. Summary statistics, including the mean, median, standard deviation, variance, skewness, and kurtosis, are displayed alongside each plot. The red dashed line marks the mean value, while the black dotted line indicates the median, providing a visual summary of the central tendency, variability, and overall distribution of each metric.

(a) Distribution of Diverse_4 Aromatic Ring Count
(b) Distribution of Diverse_4 Fraction CSP3
(c) Distribution of Diverse_4 HBA
(d) Distribution of Diverse_4 HBD

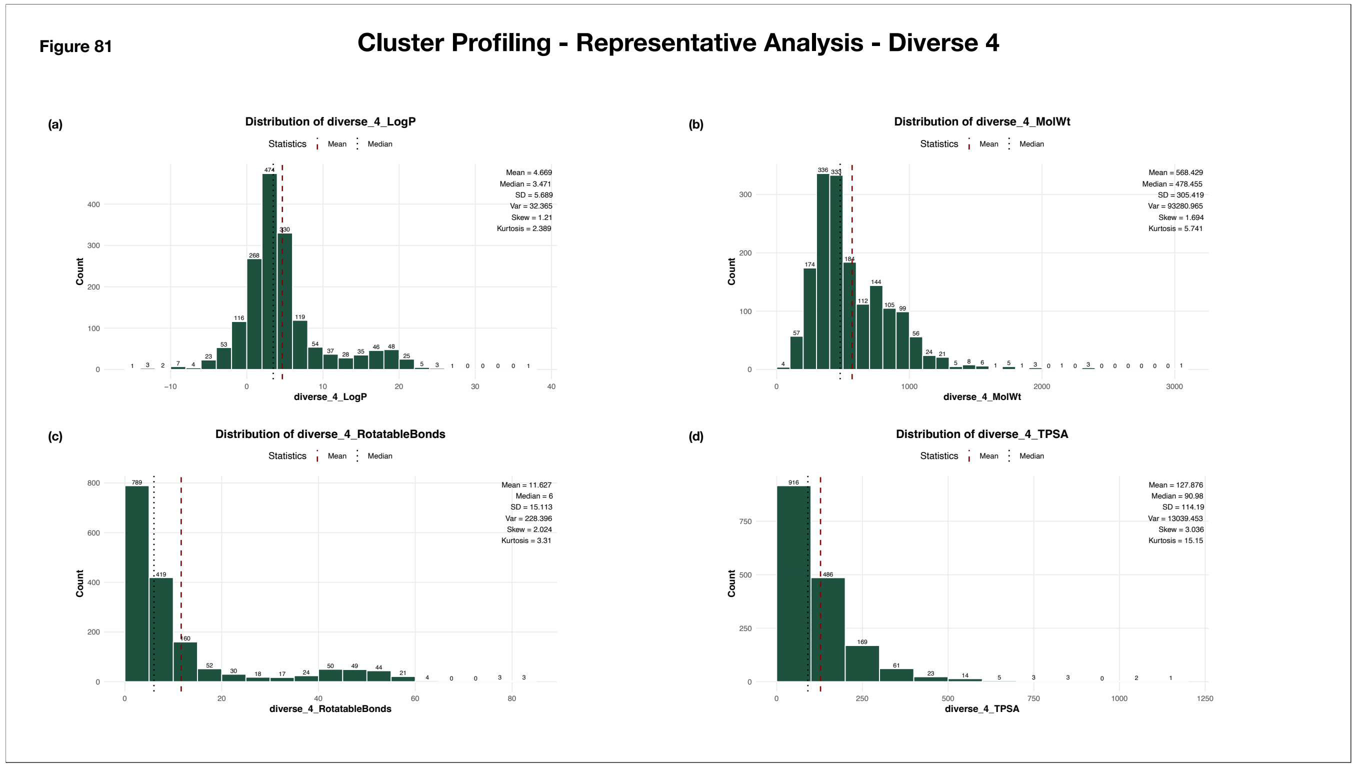


**Supplementary Figure 81: Distribution of Descriptor-Based Properties of Diverse_4 Representatives Across All Clusters**

The histograms illustrate the distributions of diverse_4 representative values across all clusters. The x-axis represents the observed values grouped into predefined bins, while the y-axis indicates the number of clusters within each bin. Bar annotations denote the corresponding cluster counts. Summary statistics, including the mean, median, standard deviation, variance, skewness, and kurtosis, are displayed alongside each plot. The red dashed line marks the mean value, while the black dotted line indicates the median, providing a visual summary of the central tendency, variability, and overall distribution of each metric.

(a) Distribution of Diverse_4 LogP
(b) Distribution of Diverse_4 Molecular Weight
(c) Distribution of Diverse_4 Rotatable Bonds
(d) Distribution of Diverse_4 TPSA

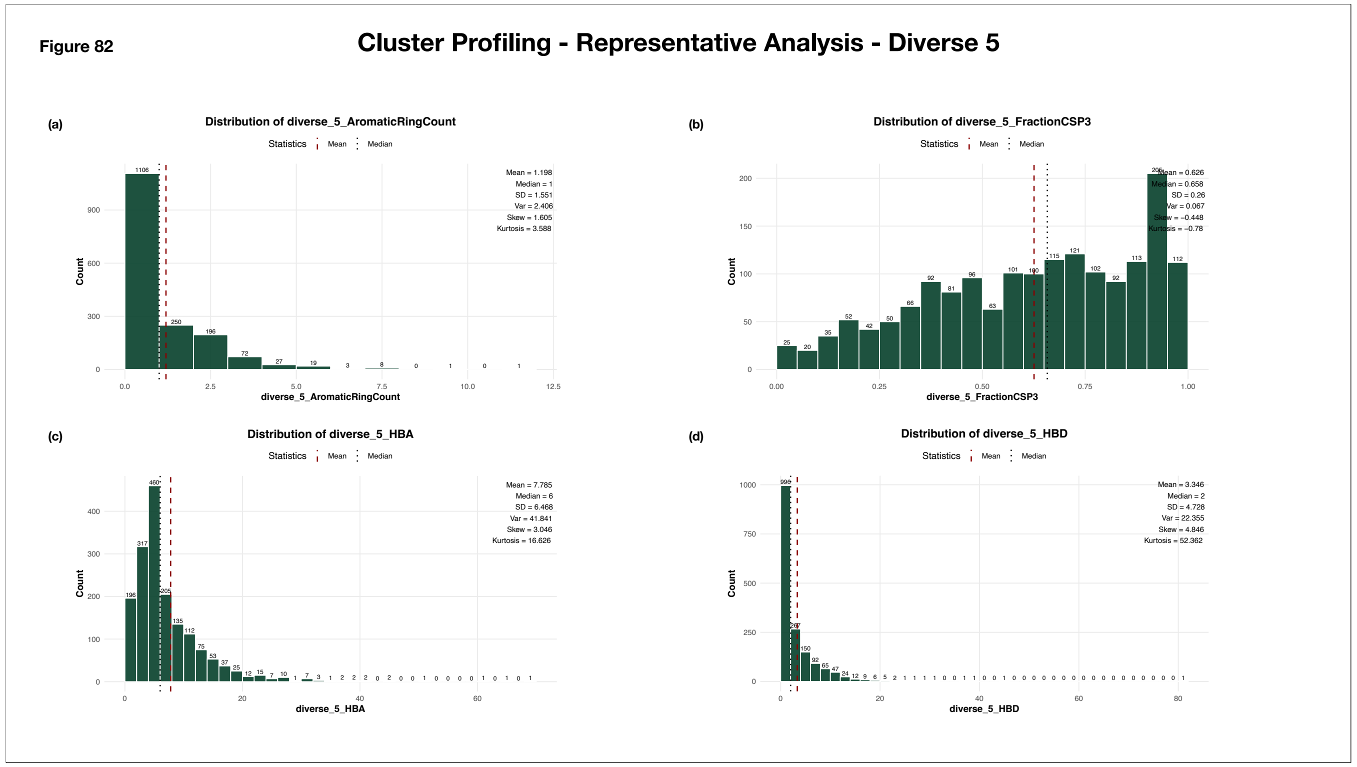


**Supplementary Figure 82: Distribution of Descriptor-Based Properties of Diverse_5 Representatives Across All Clusters**

The histograms illustrate the distributions of diverse_5 representative values across all clusters. The x-axis represents the observed values grouped into predefined bins, while the y-axis indicates the number of clusters within each bin. Bar annotations denote the corresponding cluster counts. Summary statistics, including the mean, median, standard deviation, variance, skewness, and kurtosis, are displayed alongside each plot. The red dashed line marks the mean value, while the black dotted line indicates the median, providing a visual summary of the central tendency, variability, and overall distribution of each metric.

(a) Distribution of Diverse_5 Aromatic Ring Count
(b) Distribution of Diverse_5 Fraction CSP3
(c) Distribution of Diverse_5 HBA
(d) Distribution of Diverse_5 HBD

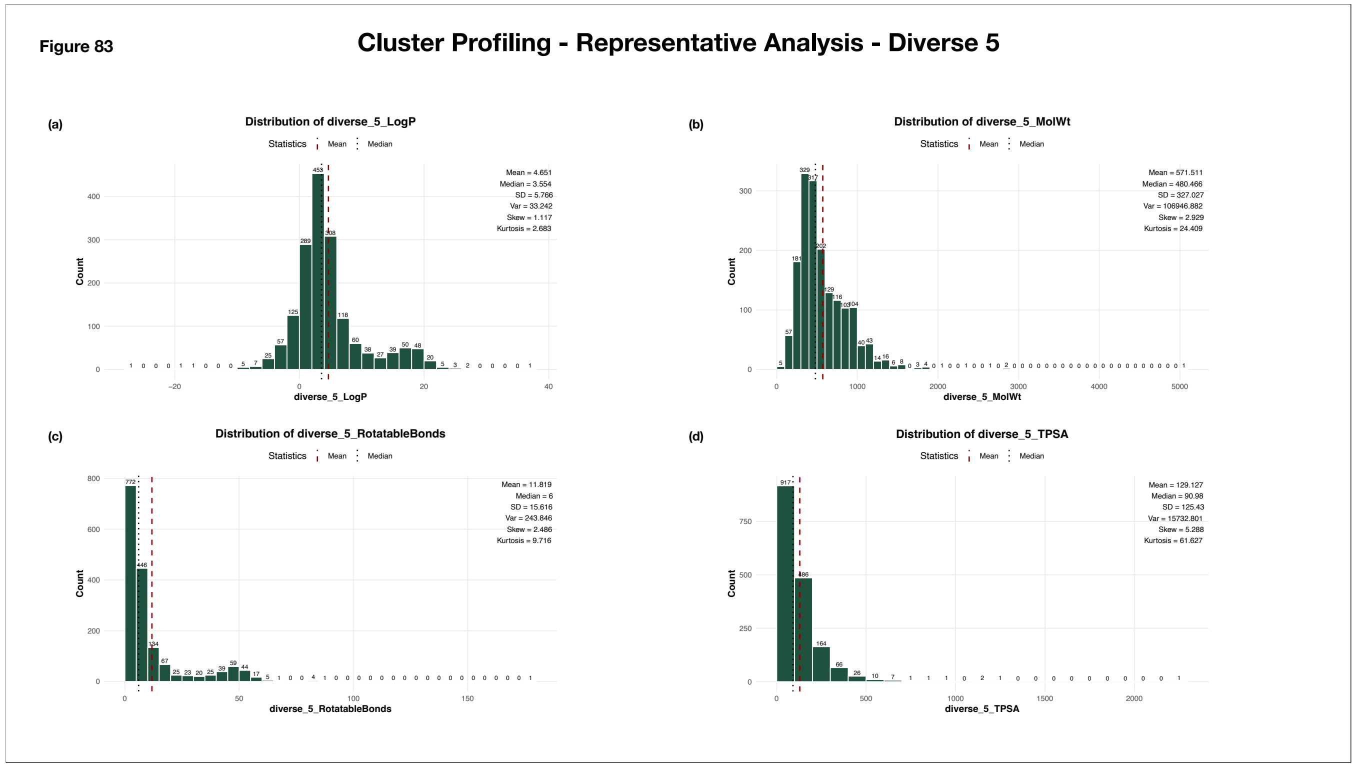


**Supplementary Figure 83: Distribution of Descriptor-Based Properties of Diverse_5 Representatives Across All Clusters**

The histograms illustrate the distributions of diverse_5 representative values across all clusters. The x-axis represents the observed values grouped into predefined bins, while the y-axis indicates the number of clusters within each bin. Bar annotations denote the corresponding cluster counts. Summary statistics, including the mean, median, standard deviation, variance, skewness, and kurtosis, are displayed alongside each plot. The red dashed line marks the mean value, while the black dotted line indicates the median, providing a visual summary of the central tendency, variability, and overall distribution of each metric.

(a) Distribution of Diverse_5 LogP
(b) Distribution of Diverse_5 Molecular Weight
(c) Distribution of Diverse_5 Rotatable Bonds
(d) Distribution of Diverse_5 TPSA

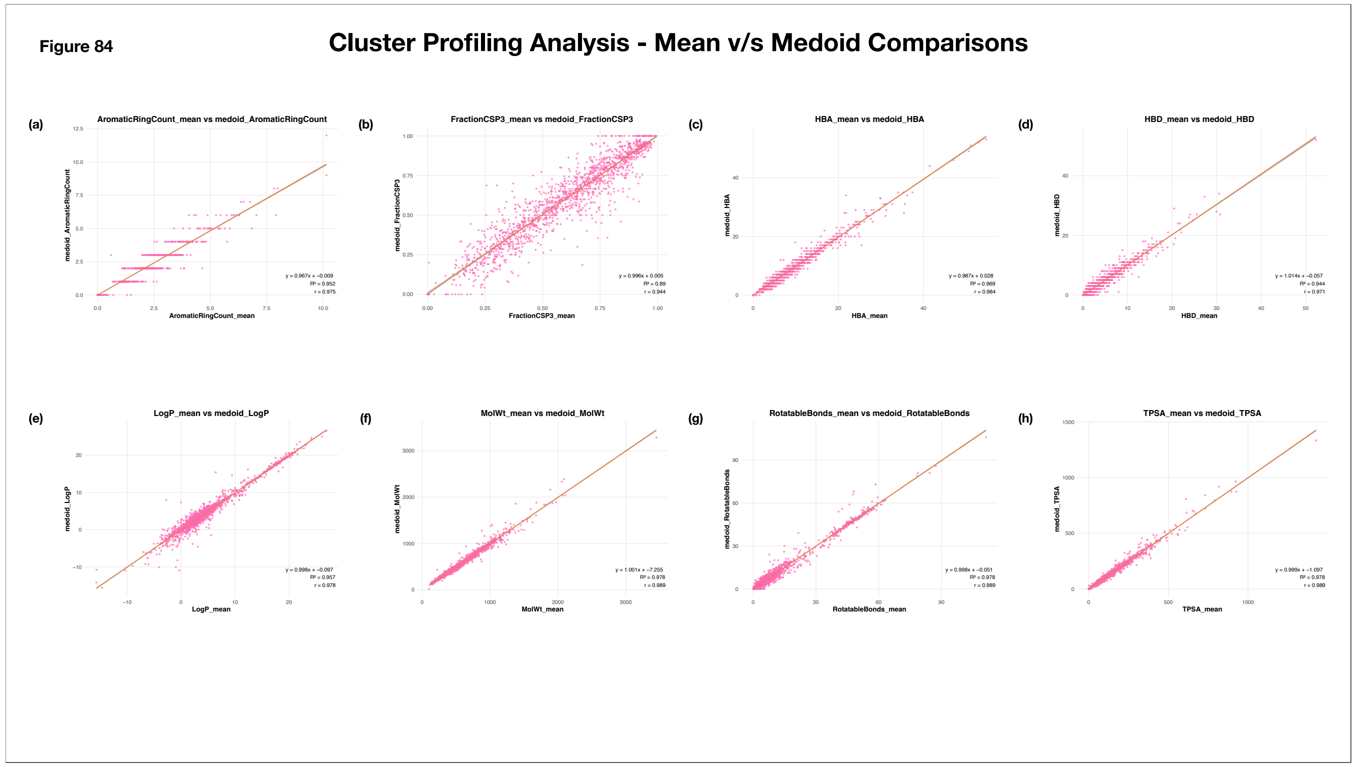


**Supplementary Figure 84: Bivariate Analysis of Cluster Representative Associations**

The scatterplots illustrate the relationships between descriptor values of cluster medoids and mean descriptor value of cluster molecules. The x-axis represents the mean descriptor value for the clusters, and y-axis represents the descriptor value for the medoid of the cluster. Each pink point corresponds to an individual cluster. A red regression line is overlaid to highlight the linear trend between the variables. The coefficient of determination ($R^2$), Pearson correlation coefficient (r), and regression equation are annotated in the lower-right corner of each plot.

(a) Medoid v/s Mean (Aromatic Ring Count)
(b) Medoid v/s Mean (Fraction CSP3)
(c) Medoid v/s Mean (HBA)
(d) Medoid v/s Mean (HBD)
(e) Medoid v/s Mean (LogP)
(f) Medoid v/s Mean (Molecular Weight)
(g) Medoid v/s Mean (Rotatable Bonds)
(h) Medoid v/s Mean (TPSA)

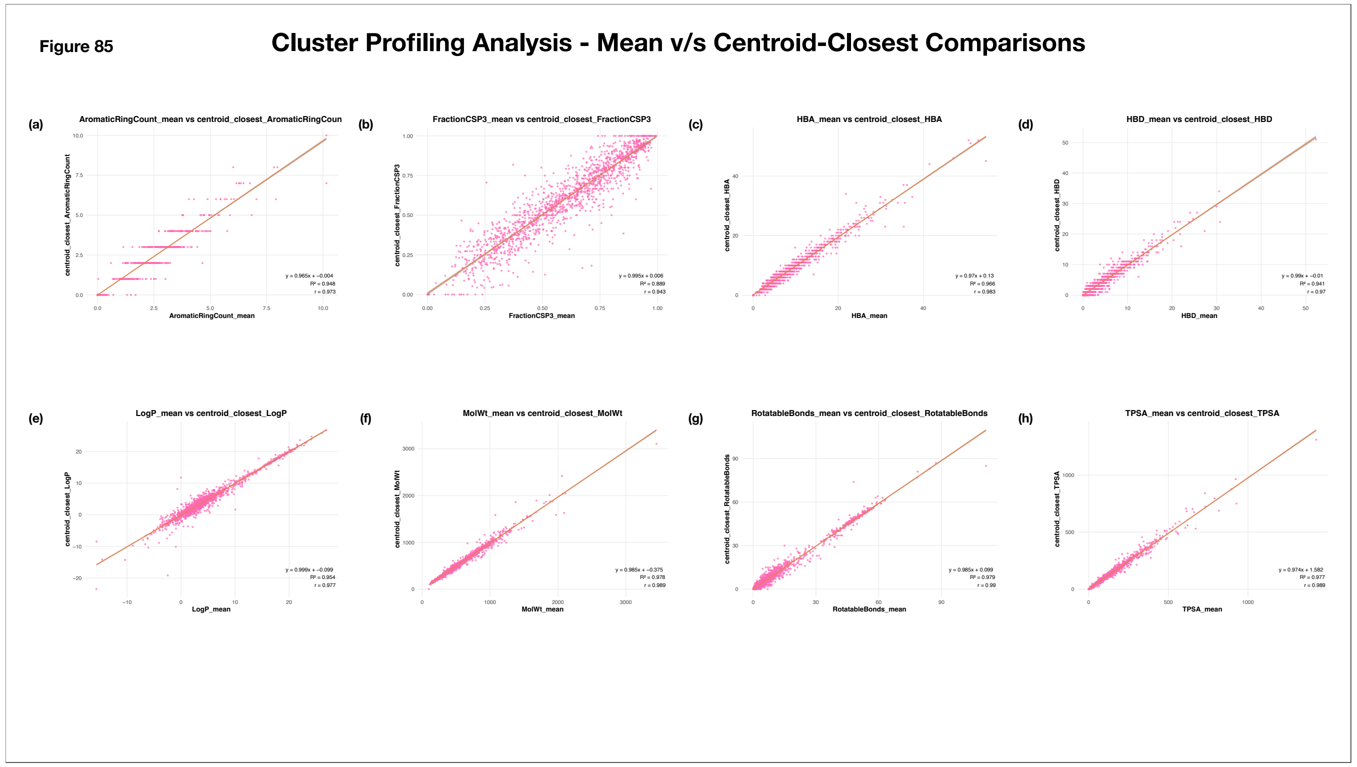


**Supplementary Figure 85: Bivariate Analysis of Cluster Representative Associations**

The scatterplots illustrate the relationships between descriptor values of cluster's centroid-closest representatives and mean descriptor value of cluster molecules. The x-axis represents the mean descriptor value for the clusters, and y-axis represents the descriptor value for the centroid-closest representative of the cluster. Each pink point corresponds to an individual cluster. A red regression line is overlaid to highlight the linear trend between the variables. The coefficient of determination ($R^2$), Pearson correlation coefficient (r), and regression equation are annotated in the lower-right corner of each plot.

(a) Centroid-Closest v/s Mean (Aromatic Ring Count)
(b) Centroid-Closest v/s Mean (Fraction CSP3)
(c) Centroid-Closest v/s Mean (HBA)
(d) Centroid-Closest v/s Mean (HBD)
(e) Centroid-Closest v/s Mean (LogP)
(f) Centroid-Closest v/s Mean (Molecular Weight)
(g) Centroid-Closest v/s Mean (Rotatable Bonds)
(h) Centroid-Closest v/s Mean (TPSA)

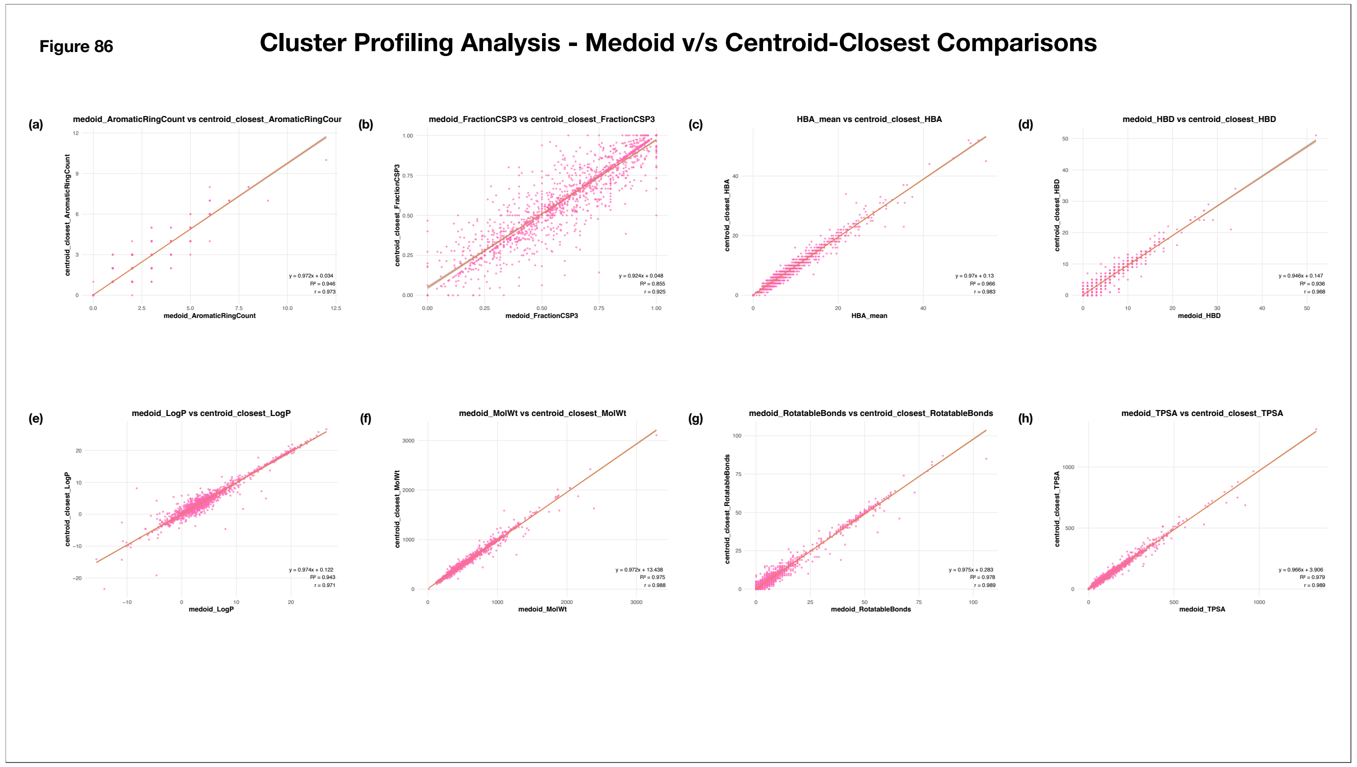


**Supplementary Figure 86: Bivariate Analysis of Cluster Representative Associations**

The scatterplots illustrate the relationships between descriptor values of cluster's centroid-closest and medoid representative. The x- and y-axes represents the descriptor values for the medoid and centroid-closest representative of the clusters respectively. Each pink point corresponds to an individual cluster. A red regression line is overlaid to highlight the linear trend between the variables. The coefficient of determination ($R^2$), Pearson correlation coefficient (r), and regression equation are annotated in the lower-right corner of each plot.

(a) Centroid-Closest v/s Medoid (Aromatic Ring Count)
(b) Centroid-Closest v/s Medoid (Fraction CSP3)
(c) Centroid-Closest v/s Medoid (HBA)
(d) Centroid-Closest v/s Medoid (HBD)
(e) Centroid-Closest v/s Medoid (LogP)
(f) Centroid-Closest v/s Medoid (Molecular Weight)
(g) Centroid-Closest v/s Medoid (Rotatable Bonds)
(h) Centroid-Closest v/s Medoid (TPSA)

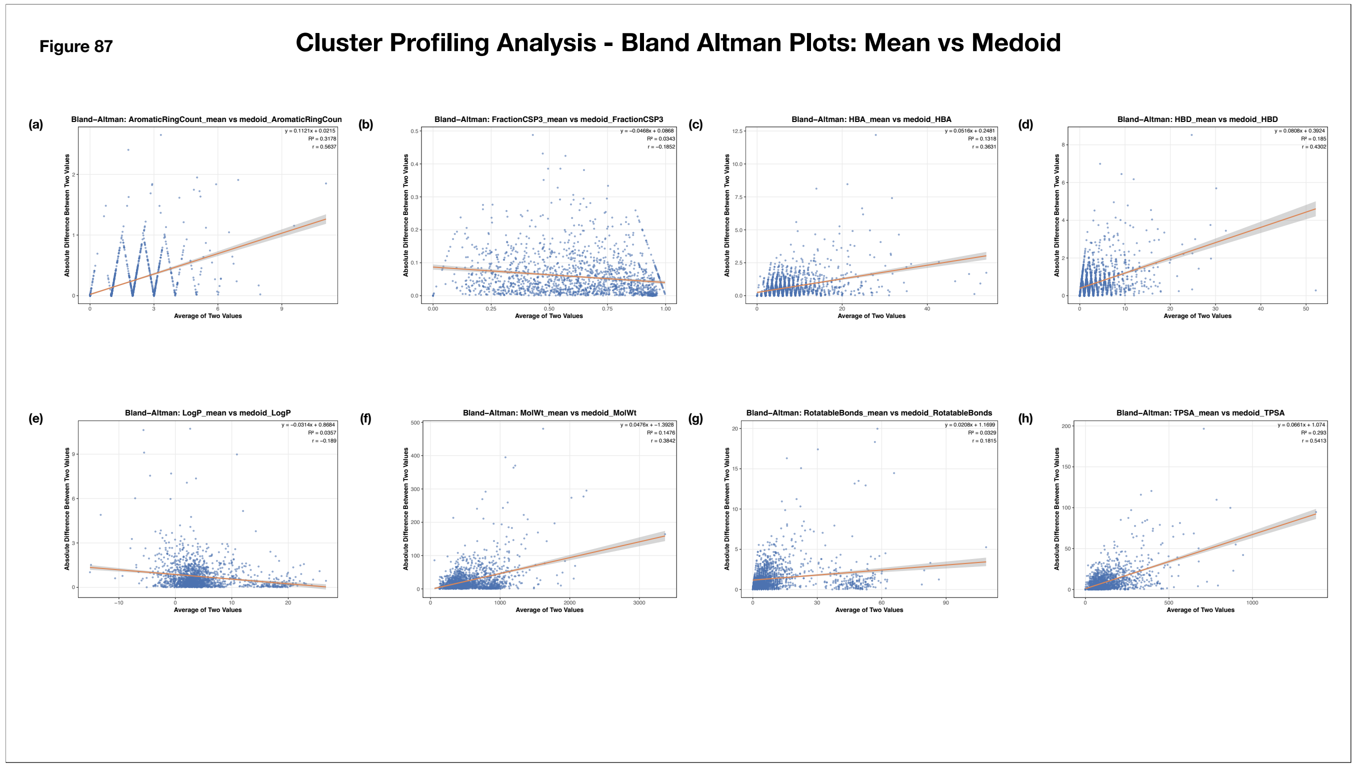


**Supplementary Figure 87: Bland Altman Analyses of Cluster Representatives**

The scatterplots represent Bland-Altman analyses of the relationships between mean descriptor values and the descriptor values for the medoid representative of the clusters. The x-axis represents the average of the mean descriptor value and the medoid's descriptor value, whereas the y-axis represents the absolute difference between the two. Each point represents an individual cluster. A red regression line is overlaid to highlight the linear trend between the variables. The coefficient of determination ($R^2$), Pearson correlation coefficient (r), and regression equation are annotated in the upper-right corner of each plot.

(a) Aromatic Ring Count
(b) Fraction CSP3
(c) HBA
(d) HBD
(e) LogP
(f) Molecular Weight
(g) Rotatable Bonds
(h) TPSA

Figure 88

#### Cluster Profiling Analysis - Bland Altman Plots: Mean v/s Centroid-Closest

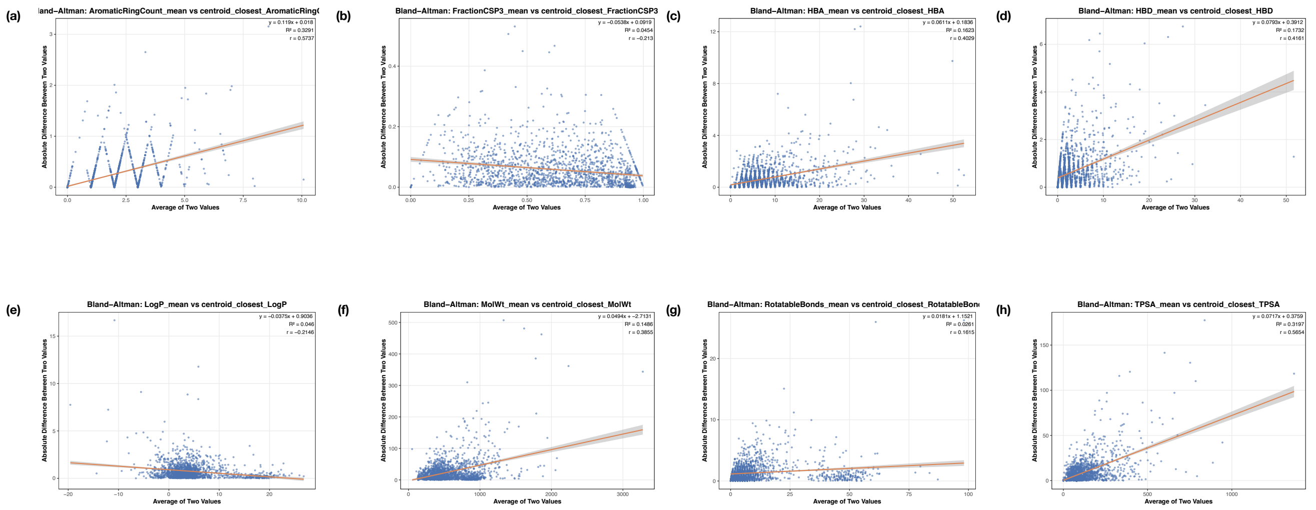


**Supplementary Figure 88: Bland Altman Analyses of Cluster Representatives**

The scatterplots represent Bland-Altman analyses of the relationships between mean descriptor values and the descriptor values for the centroid-closest representative of the clusters. The x-axis represents the average of the mean descriptor value and the centroid-closest's descriptor value, whereas the y-axis represents the absolute difference between the two. Each point represents an individual cluster. A red regression line is overlaid to highlight the linear trend between the variables. The coefficient of determination ($R^2$), Pearson correlation coefficient (r), and regression equation are annotated in the upper-right corner of each plot.

(a) Aromatic Ring Count
(b) Fraction CSP3
(c) HBA
(d) HBD
(e) LogP
(f) Molecular Weight
(g) Rotatable Bonds
(h) TPSA

Figure 89

### Cluster Profiling Analysis - Bland Altman Plots: Medoid vs Centroid-Closest

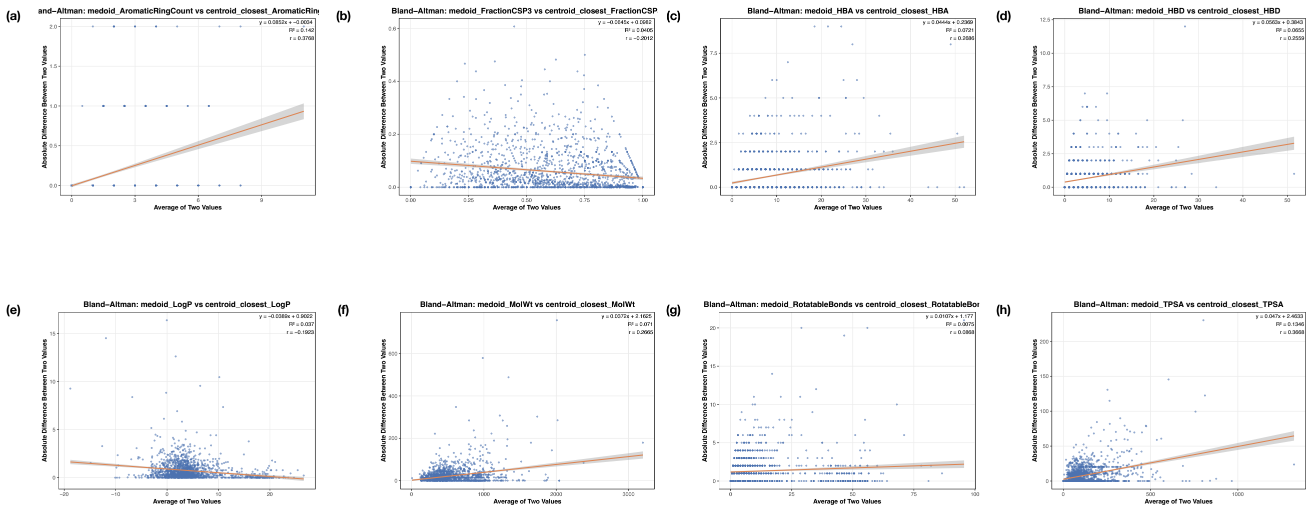

**Supplementary Figure 89: Bland Altman Analyses of Cluster Representatives**

The scatterplots represent Bland-Altman analyses of the relationships between the descriptor values for the medoid and centroid-closest representatives of the clusters. The x-axis represents the average of the medoid and centroid-closest's descriptor values, whereas the y-axis represents the absolute difference between the two. Each point represents an individual cluster. A red regression line is overlaid to highlight the linear trend between the variables. The coefficient of determination ($R^2$), Pearson correlation coefficient (r), and regression equation are annotated in the upper-right corner of each plot.

(a) Aromatic Ring Count
(b) Fraction CSP3
(c) HBA
(d) HBD
(e) LogP
(f) Molecular Weight
(g) Rotatable Bonds
(h) TPSA

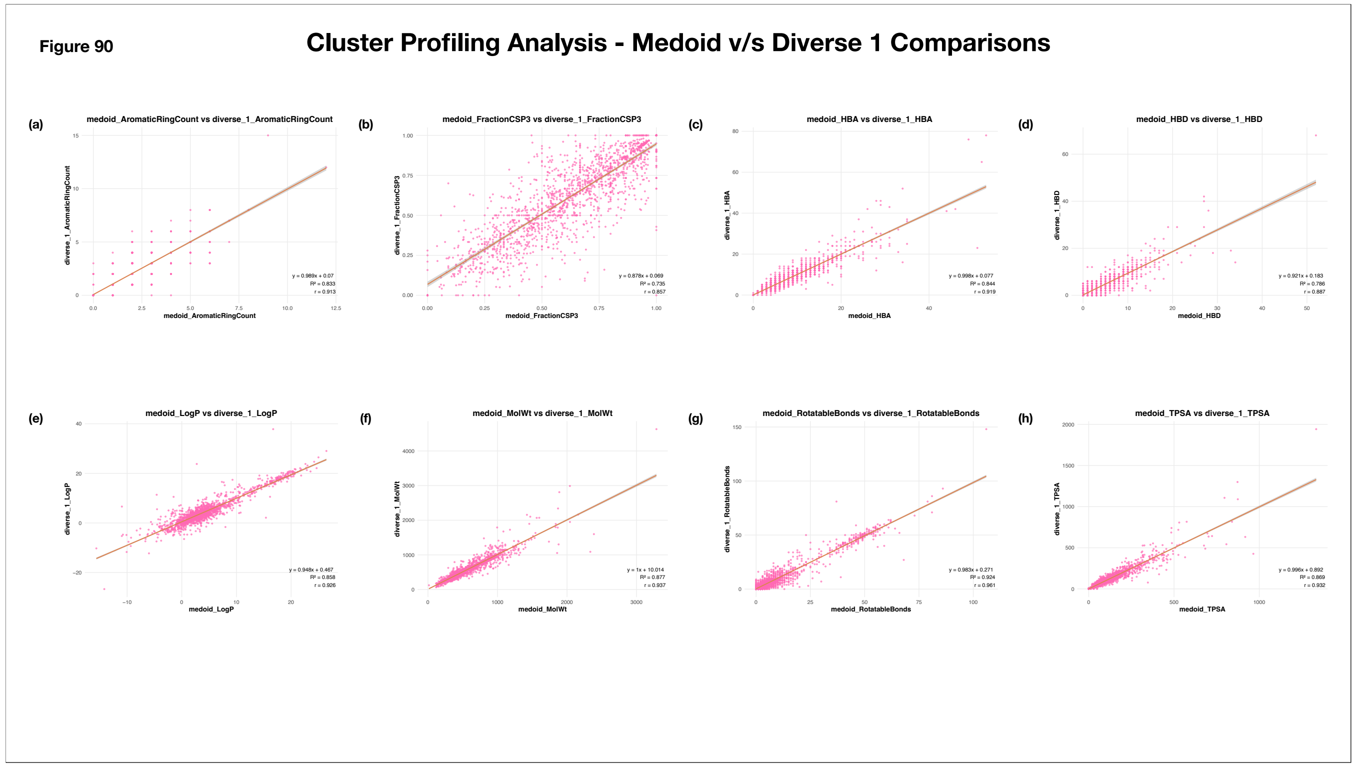


**Supplementary Figure 90: Bivariate Analysis of Cluster Representative Associations**

The scatterplots illustrate the relationships between descriptor values of cluster's diverse_1 and medoid representatives. The x- and y-axes represents the descriptor values for the medoid and diverse_1 representative of the clusters respectively. Each pink point corresponds to an individual cluster. A red regression line is overlaid to highlight the linear trend between the variables. The coefficient of determination ($R^2$), Pearson correlation coefficient (r), and regression equation are annotated in the lower-right corner of each plot.

(a) Diverse_1 v/s Medoid (Aromatic Ring Count)
(b) Diverse_1 v/s Medoid (Fraction CSP3)
(c) Diverse_1 v/s Medoid (HBA)
(d) Diverse_1 v/s Medoid (HBD)
(e) Diverse_1 v/s Medoid (LogP)
(f) Diverse_1 v/s Medoid (Molecular Weight)
(g) Diverse_1 v/s Medoid (Rotatable Bonds)
(h) Diverse_1 v/s Medoid (TPSA)

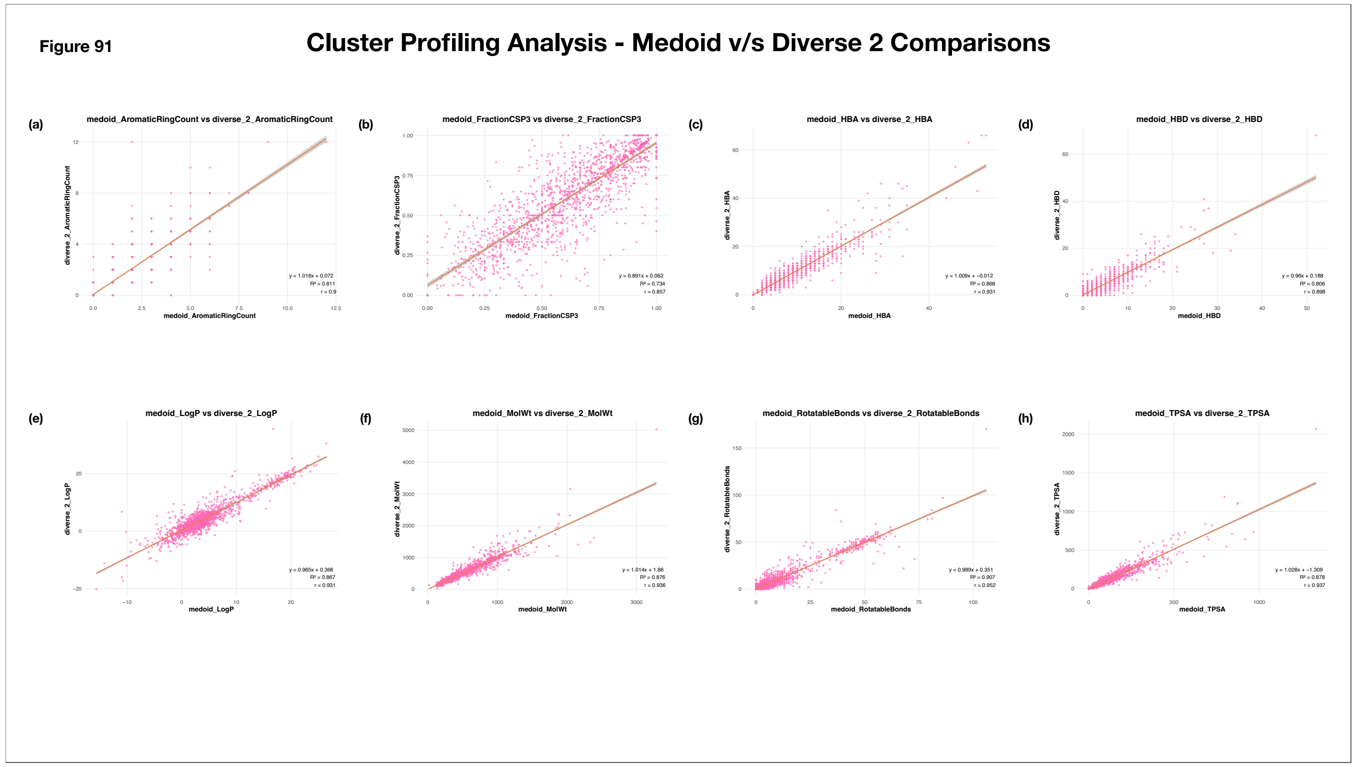


**Supplementary Figure 91: Bivariate Analysis of Cluster Representative Associations**

The scatterplots illustrate the relationships between descriptor values of cluster's diverse_2 and medoid representatives. The x- and y-axes represents the descriptor values for the medoid and diverse_2 representative of the clusters respectively. Each pink point corresponds to an individual cluster. A red regression line is overlaid to highlight the linear trend between the variables. The coefficient of determination ($R^2$), Pearson correlation coefficient (r), and regression equation are annotated in the lower-right corner of each plot.

(a) Diverse_2 v/s Medoid (Aromatic Ring Count)
(b) Diverse_2 v/s Medoid (Fraction CSP3)
(c) Diverse_2 v/s Medoid (HBA)
(d) Diverse_2 v/s Medoid (HBD)
(e) Diverse_2 v/s Medoid (LogP)
(f) Diverse_2 v/s Medoid (Molecular Weight)
(g) Diverse_2 v/s Medoid (Rotatable Bonds)
(h) Diverse_2 v/s Medoid (TPSA)

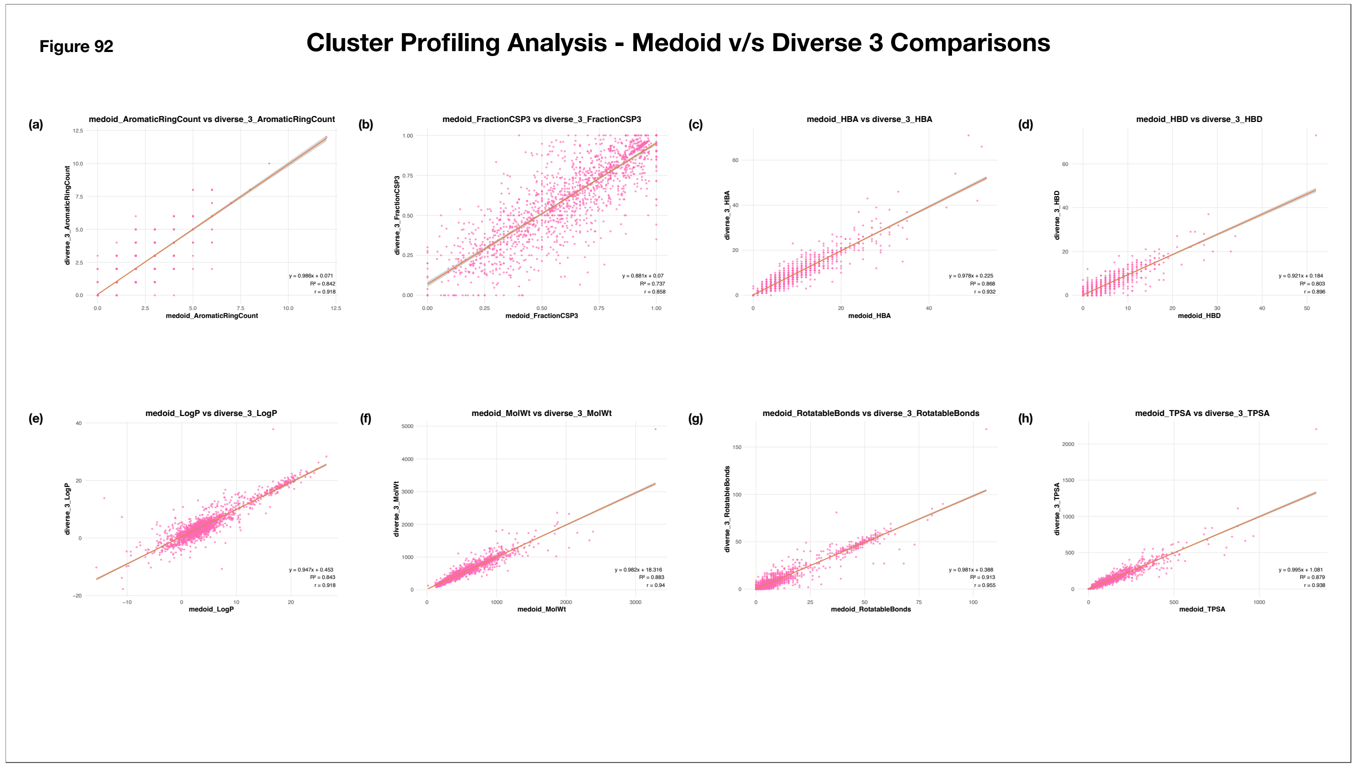


**Supplementary Figure 92: Bivariate Analysis of Cluster Representative Associations**

The scatterplots illustrate the relationships between descriptor values of cluster's diverse_3 and medoid representatives. The x- and y-axes represents the descriptor values for the medoid and diverse_3 representative of the clusters respectively. Each pink point corresponds to an individual cluster. A red regression line is overlaid to highlight the linear trend between the variables. The coefficient of determination ($R^2$), Pearson correlation coefficient (r), and regression equation are annotated in the lower-right corner of each plot.

(a) Diverse_3 v/s Medoid (Aromatic Ring Count)
(b) Diverse_3 v/s Medoid (Fraction CSP3)
(c) Diverse_3 v/s Medoid (HBA)
(d) Diverse_3 v/s Medoid (HBD)
(e) Diverse_3 v/s Medoid (LogP)
(f) Diverse_3 v/s Medoid (Molecular Weight)
(g) Diverse_3 v/s Medoid (Rotatable Bonds)
(h) Diverse_3 v/s Medoid (TPSA)

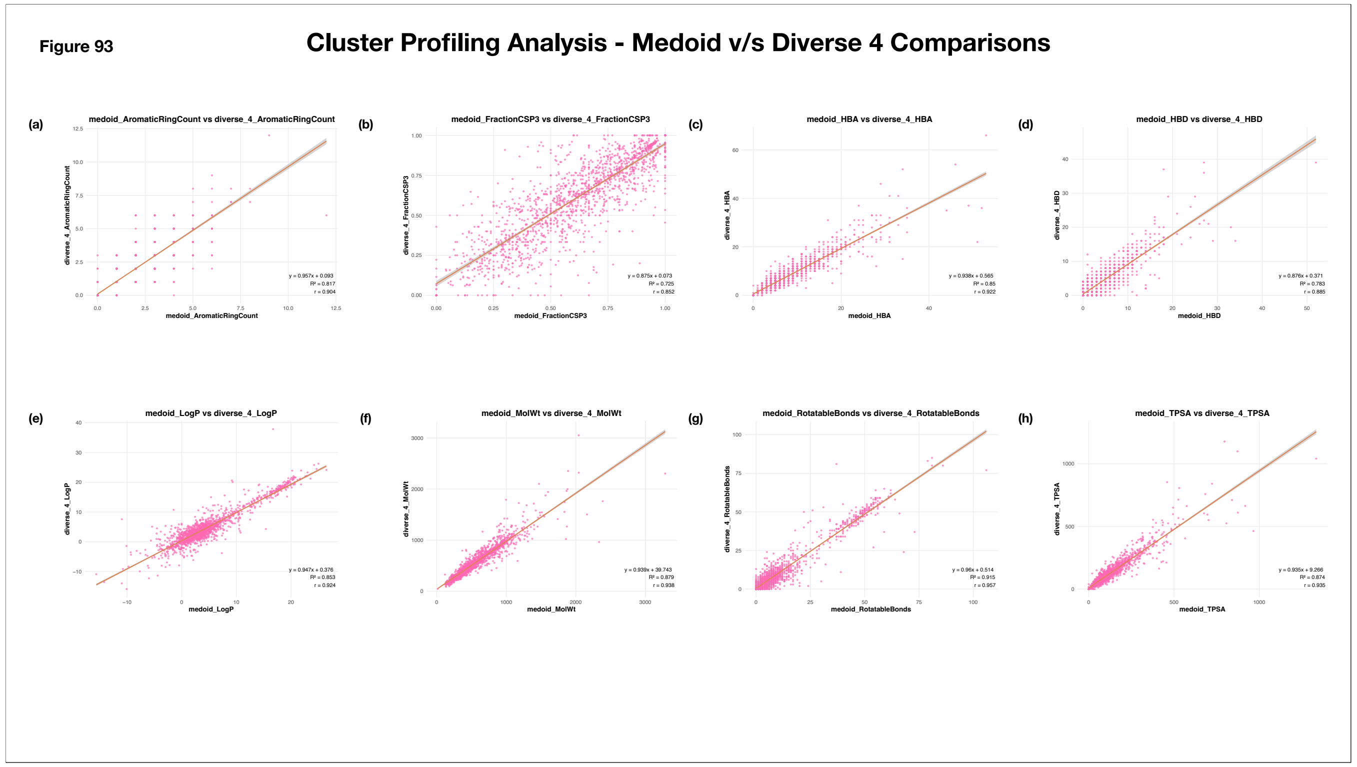


**Supplementary Figure 93: Bivariate Analysis of Cluster Representative Associations**

The scatterplots illustrate the relationships between descriptor values of cluster's diverse_4 and medoid representatives. The x- and y-axes represents the descriptor values for the medoid and diverse_4 representative of the clusters respectively. Each pink point corresponds to an individual cluster. A red regression line is overlaid to highlight the linear trend between the variables. The coefficient of determination ($R^2$), Pearson correlation coefficient (r), and regression equation are annotated in the lower-right corner of each plot.

(a) Diverse_4 v/s Medoid (Aromatic Ring Count)
(b) Diverse_4 v/s Medoid (Fraction CSP3)
(c) Diverse_4 v/s Medoid (HBA)
(d) Diverse_4 v/s Medoid (HBD)
(e) Diverse_4 v/s Medoid (LogP)
(f) Diverse_4 v/s Medoid (Molecular Weight)
(g) Diverse_4 v/s Medoid (Rotatable Bonds)
(h) Diverse_4 v/s Medoid (TPSA)

**Figure 94**

#### Cluster Profiling Analysis - Medoid v/s Diverse 5 Comparisons

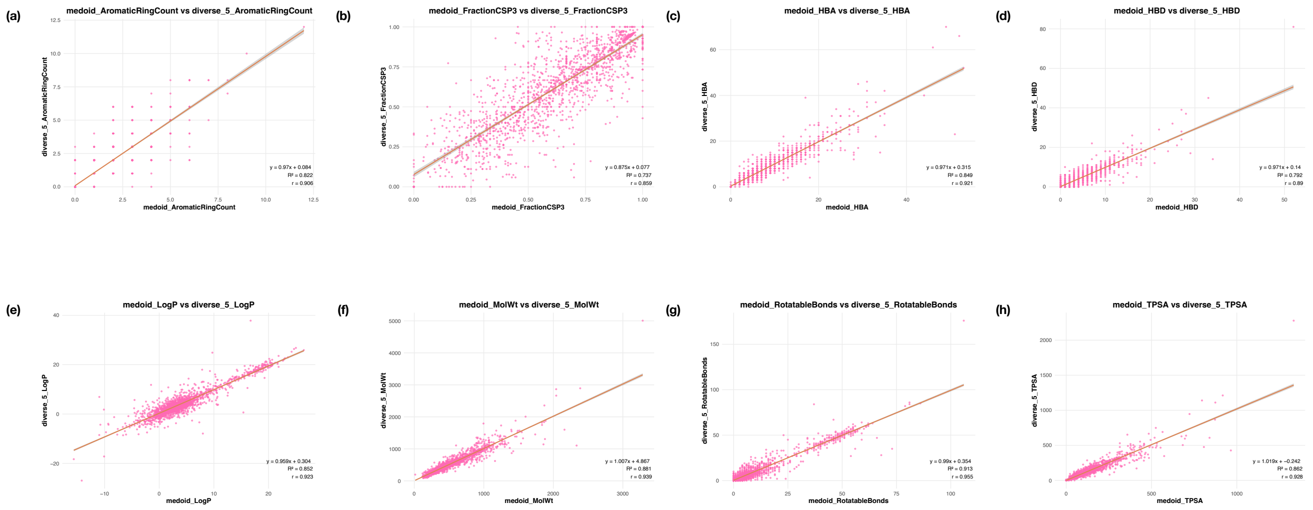


**Supplementary Figure 94: Bivariate Analysis of Cluster Representative Associations**

The scatterplots illustrate the relationships between descriptor values of cluster's diverse_5 and medoid representatives. The x- and y-axes represents the descriptor values for the medoid and diverse_5 representative of the clusters respectively. Each pink point corresponds to an individual cluster. A red regression line is overlaid to highlight the linear trend between the variables. The coefficient of determination ($R^2$), Pearson correlation coefficient (r), and regression equation are annotated in the lower-right corner of each plot.

(a) Diverse_5 v/s Medoid (Aromatic Ring Count)
(b) Diverse_5 v/s Medoid (Fraction CSP3)
(c) Diverse_5 v/s Medoid (HBA)
(d) Diverse_5 v/s Medoid (HBD)
(e) Diverse_5 v/s Medoid (LogP)
(f) Diverse_5 v/s Medoid (Molecular Weight)
(g) Diverse_5 v/s Medoid (Rotatable Bonds)
(h) Diverse_5 v/s Medoid (TPSA)

Figure 95

#### Cluster Profiling Analysis - Bland-Atlman Analyses: Medoid v/s Diverse 1

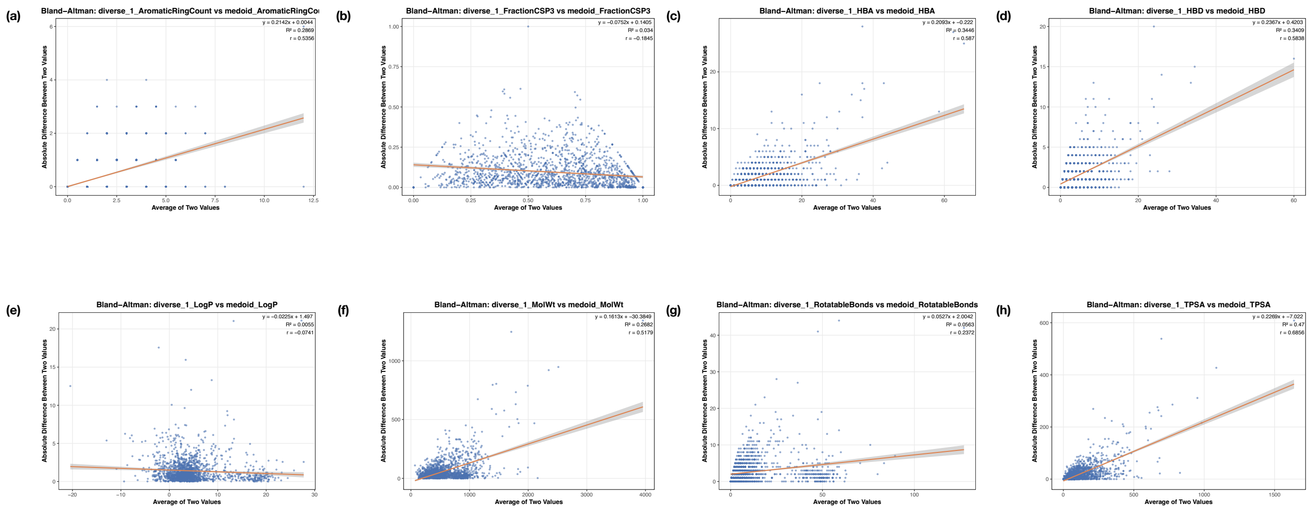


**Supplementary Figure 95: Bland Altman Analyses of Cluster Representatives**

The scatterplots represent Bland-Altman analyses of the relationships between the descriptor values for the medoid and diverse_1 representatives of the clusters. The x-axis represents the average of the medoid and diverse_1's descriptor values, whereas the y-axis represents the absolute difference between the two. Each point represents an individual cluster. A red regression line is overlaid to highlight the linear trend between the variables. The coefficient of determination ($R^2$), Pearson correlation coefficient (r), and regression equation are annotated in the upper-right corner of each plot.

(a) Aromatic Ring Count
(b) Fraction CSP3
(c) HBA
(d) HBD
(e) LogP
(f) Molecular Weight
(g) Rotatable Bonds
(h) TPSA

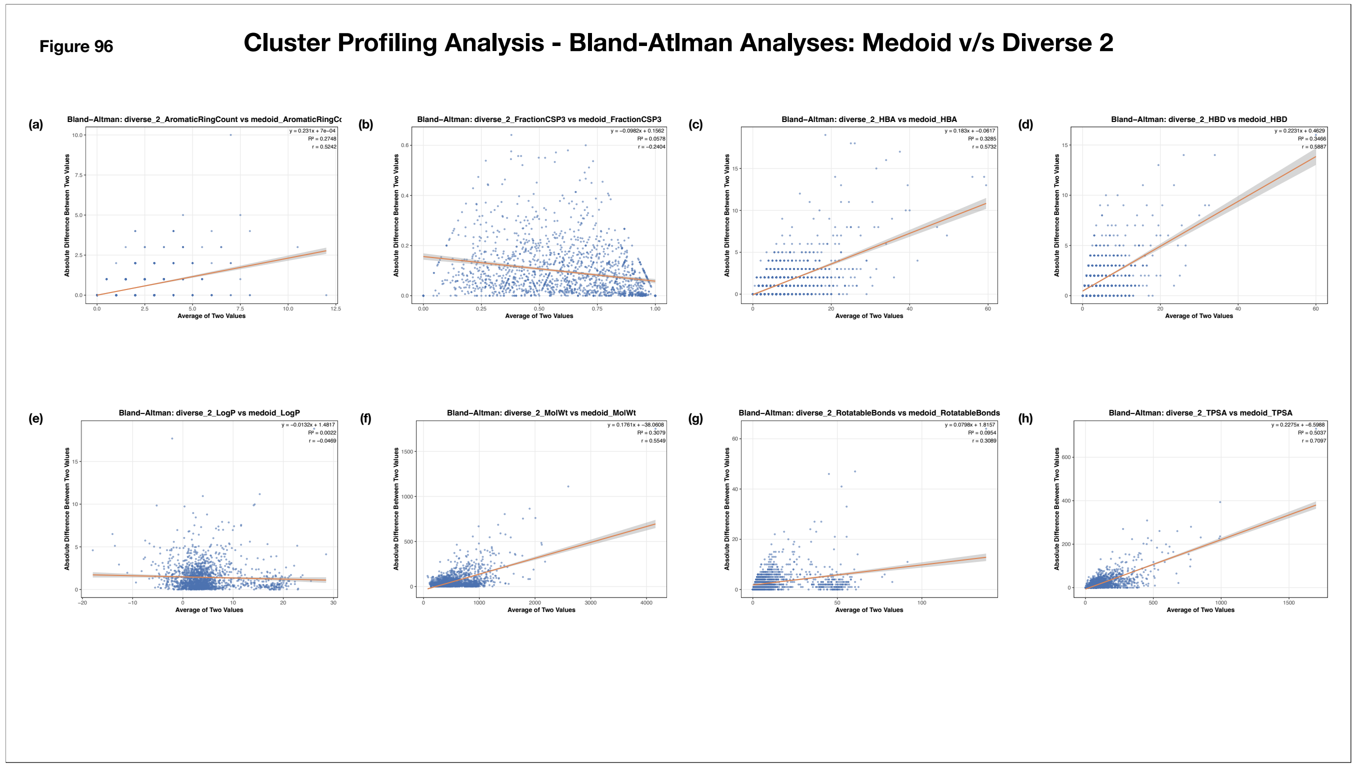


**Supplementary Figure 96: Bland Altman Analyses of Cluster Representatives**

The scatterplots represent Bland-Altman analyses of the relationships between the descriptor values for the medoid and diverse_2 representatives of the clusters. The x-axis represents the average of the medoid and diverse_2's descriptor values, whereas the y-axis represents the absolute difference between the two. Each point represents an individual cluster. A red regression line is overlaid to highlight the linear trend between the variables. The coefficient of determination ($R^2$), Pearson correlation coefficient (r), and regression equation are annotated in the upper-right corner of each plot.

(a) Aromatic Ring Count
(b) Fraction CSP3
(c) HBA
(d) HBD
(e) LogP
(f) Molecular Weight
(g) Rotatable Bonds
(h) TPSA

**Figure 97**

#### Cluster Profiling Analysis - Bland-Atlman Analyses: Medoid v/s Diverse 3

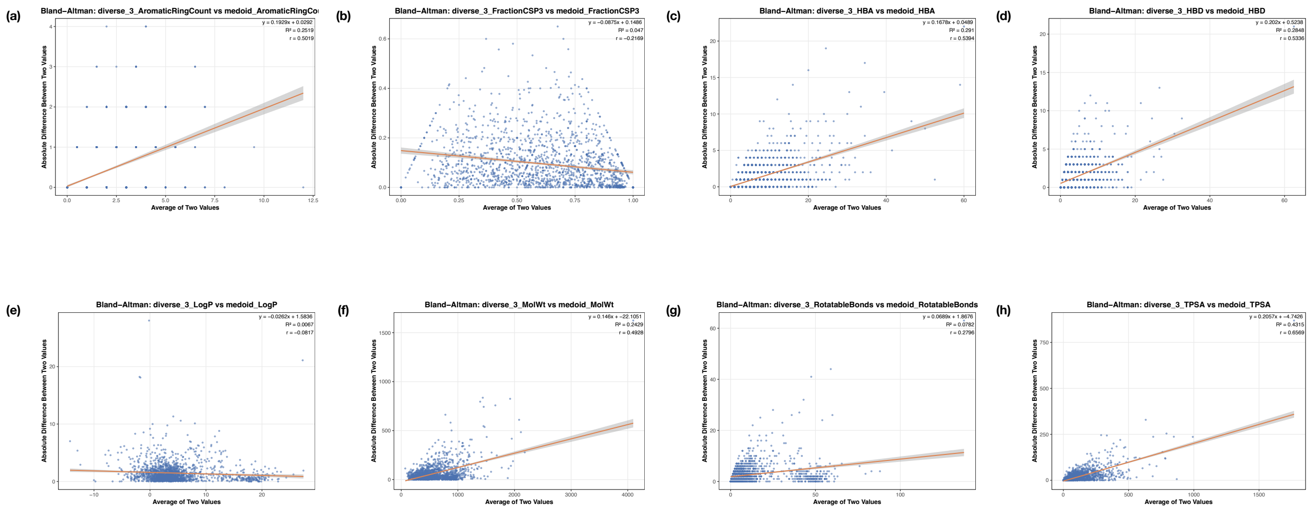


**Supplementary Figure 97: Bland Altman Analyses of Cluster Representatives**

The scatterplots represent Bland-Altman analyses of the relationships between the descriptor values for the medoid and diverse_3 representatives of the clusters. The x-axis represents the average of the medoid and diverse_3's descriptor values, whereas the y-axis represents the absolute difference between the two. Each point represents an individual cluster. A red regression line is overlaid to highlight the linear trend between the variables. The coefficient of determination ($R^2$), Pearson correlation coefficient (r), and regression equation are annotated in the upper-right corner of each plot.

(a) Aromatic Ring Count
(b) Fraction CSP3
(c) HBA
(d) HBD
(e) LogP
(f) Molecular Weight
(g) Rotatable Bonds
(h) TPSA

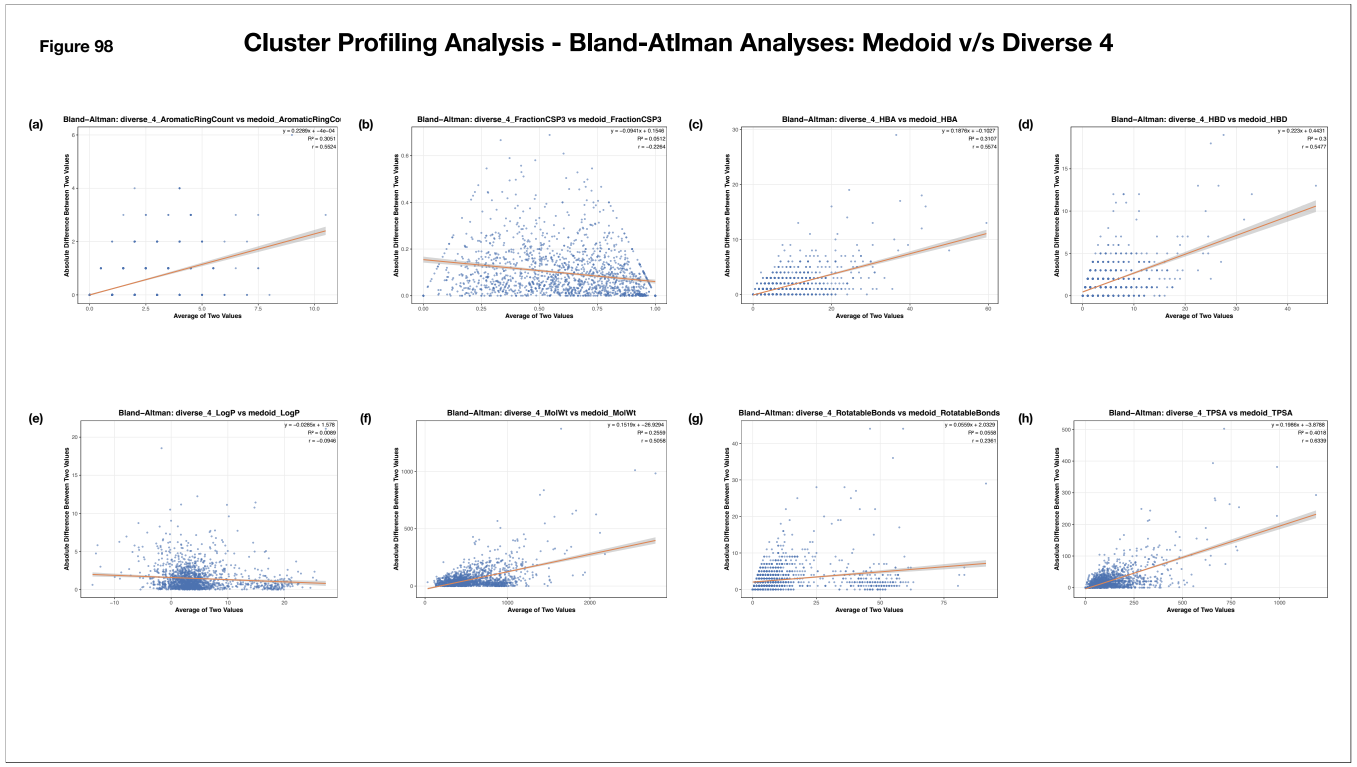


**Supplementary Figure 98: Bland Altman Analyses of Cluster Representatives**

The scatterplots represent Bland-Altman analyses of the relationships between the descriptor values for the medoid and diverse_4 representatives of the clusters. The x-axis represents the average of the medoid and diverse_4's descriptor values, whereas the y-axis represents the absolute difference between the two. Each point represents an individual cluster. A red regression line is overlaid to highlight the linear trend between the variables. The coefficient of determination ($R^2$), Pearson correlation coefficient (r), and regression equation are annotated in the upper-right corner of each plot.

(a) Aromatic Ring Count
(b) Fraction CSP3
(c) HBA
(d) HBD
(e) LogP
(f) Molecular Weight
(g) Rotatable Bonds
(h) TPSA

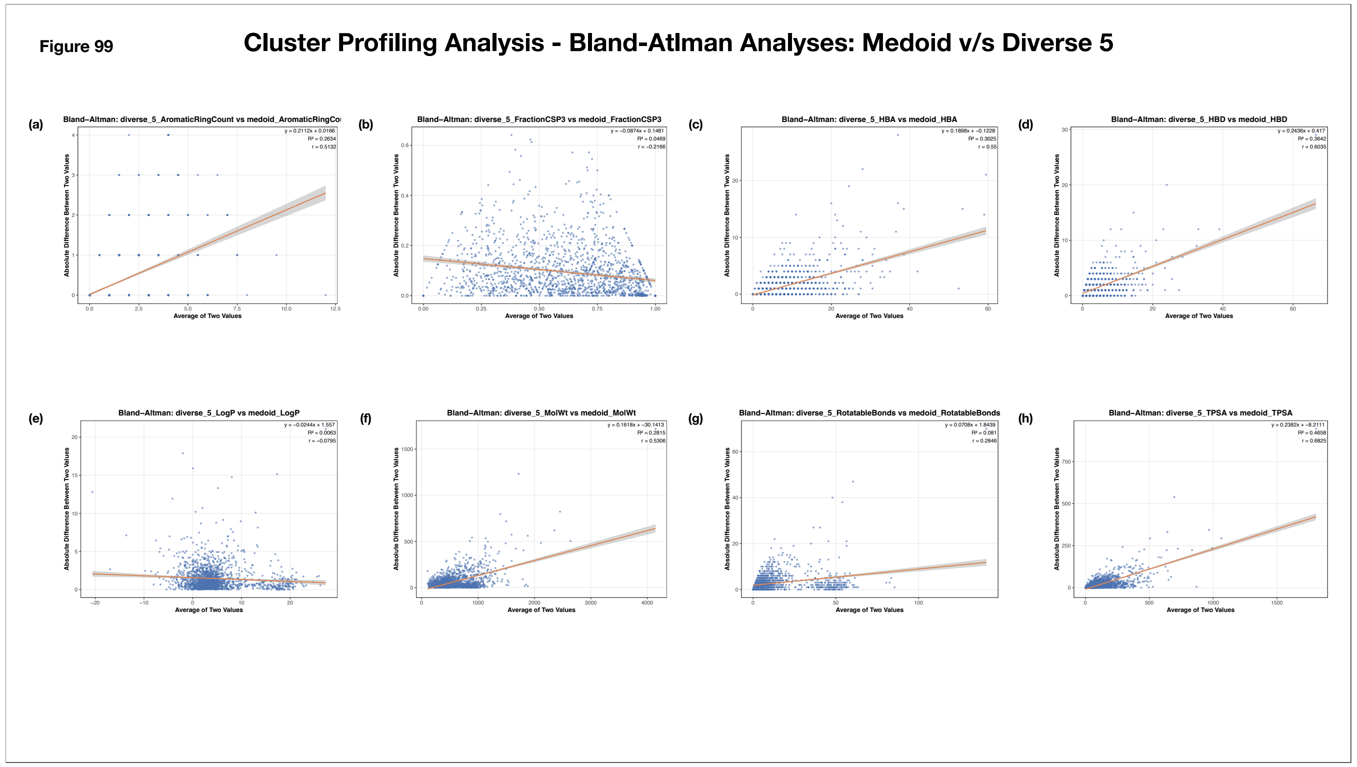


**Supplementary Figure 99: Bland Altman Analyses of Cluster Representatives**

The scatterplots represent Bland-Altman analyses of the relationships between the descriptor values for the medoid and diverse_5 representatives of the clusters. The x-axis represents the average of the medoid and diverse_5's descriptor values, whereas the y-axis represents the absolute difference between the two. Each point represents an individual cluster. A red regression line is overlaid to highlight the linear trend between the variables. The coefficient of determination ($R^2$), Pearson correlation coefficient (r), and regression equation are annotated in the upper-right corner of each plot.

(a) Aromatic Ring Count
(b) Fraction CSP3
(c) HBA
(d) HBD
(e) LogP
(f) Molecular Weight
(g) Rotatable Bonds
(h) TPSA

**Figure 100**

#### Docking Score Distribution of Representatives

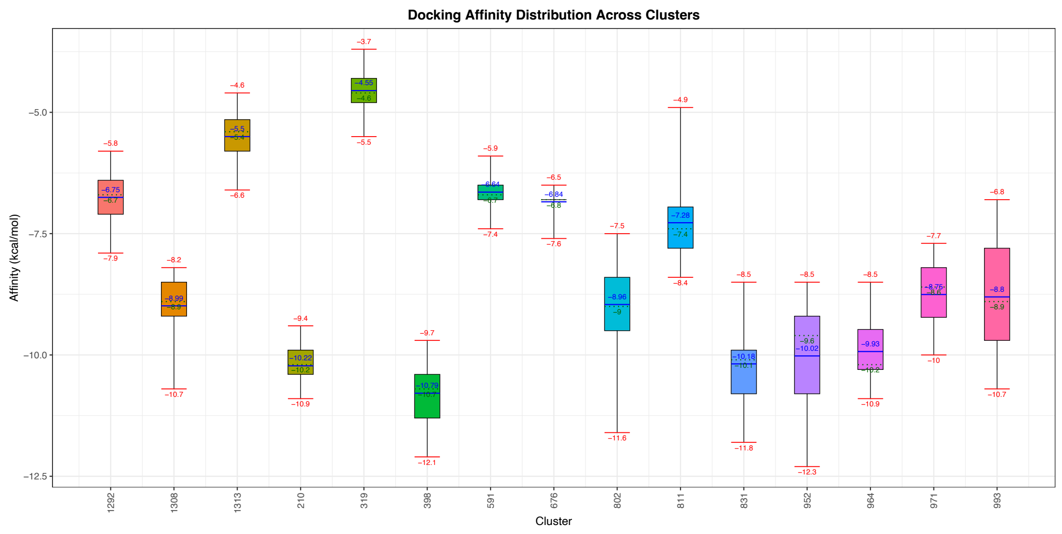


**Supplementary Figure 100: Docking Affinity Distribution of Randomly Selected Molecules from the Representative Clusters**

The box plots summarize the distribution of docking scores across the randomly selected molecules from each cluster. The x-axis represents the cluster ID, while the y-axis represents binding affinity (kcal/mol). For each representative type, the box denotes the interquartile range (Q1–Q3). The median affinity is indicated by a green dotted line, with the corresponding value annotated below the line, while the mean affinity is represented by a solid blue line, with its value annotated above the line. Red horizontal lines indicate the minimum and maximum observed affinities within the cluster, with their corresponding values annotated below and above the respective lines. Different box colors distinguish the different clusters.

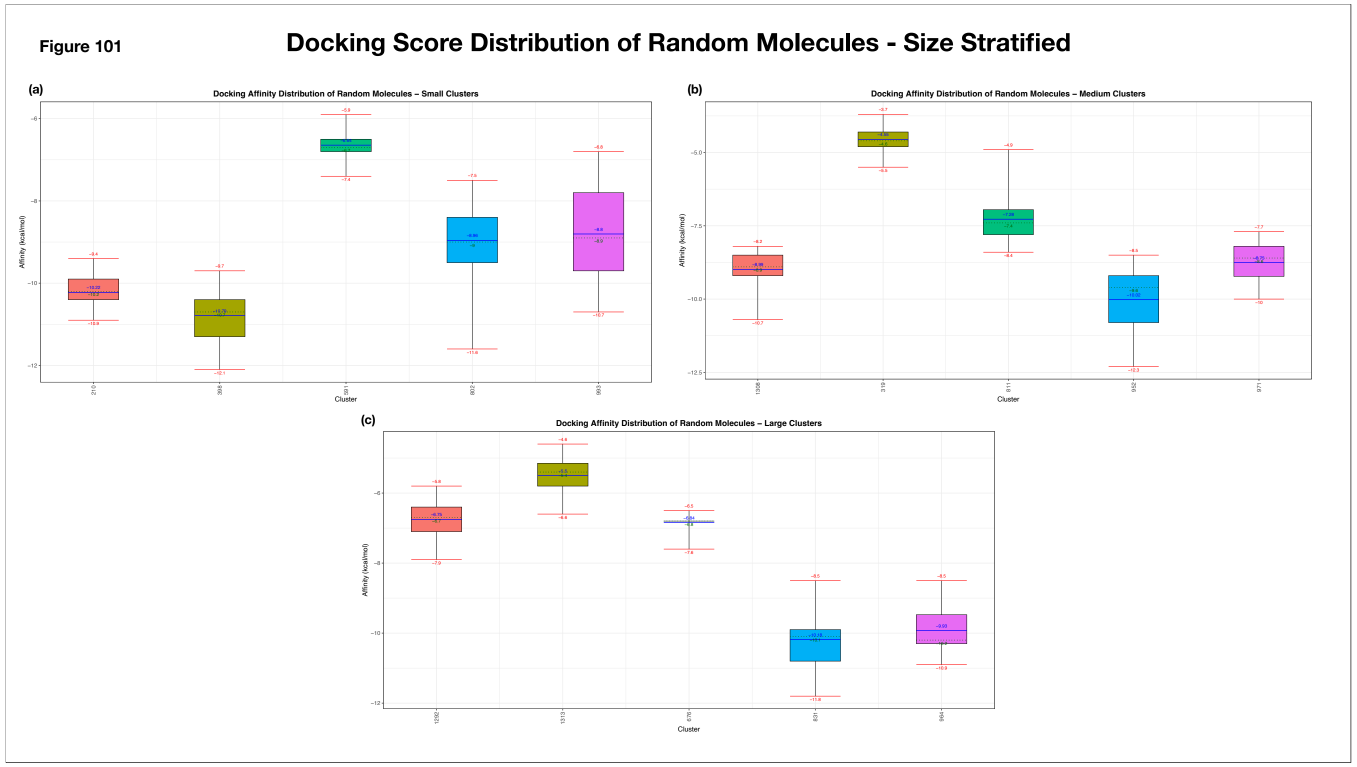


**Supplementary Figure 101: Distribution of Docking Scores of Randomly Selected Molecules across Different Size Strata**

The box plots summarize the distribution of docking scores across the randomly selected molecules from each cluster. The x-axis represents the cluster ID, while the y-axis represents binding affinity (kcal/mol). For each representative type, the box denotes the interquartile range (Q1–Q3). The median affinity is indicated by a green dotted line, with the corresponding value annotated below the line, while the mean affinity is represented by a solid blue line, with its value annotated above the line. Red horizontal lines indicate the minimum and maximum observed affinities within the cluster, with their corresponding values annotated below and above the respective lines. Different box colors distinguish the different clusters.

(a) Distribution of Docking Affinities of Randomly Selected Molecules - Small-Sized Clusters

(b) Distribution of Docking Affinities of Randomly Selected Molecules - Medium-Sized Clusters

(c) Distribution of Docking Affinities of Randomly Selected Molecules - Large-Sized Clusters

**Figure 102**

#### Representative Suitability for Docking - Global Analysis

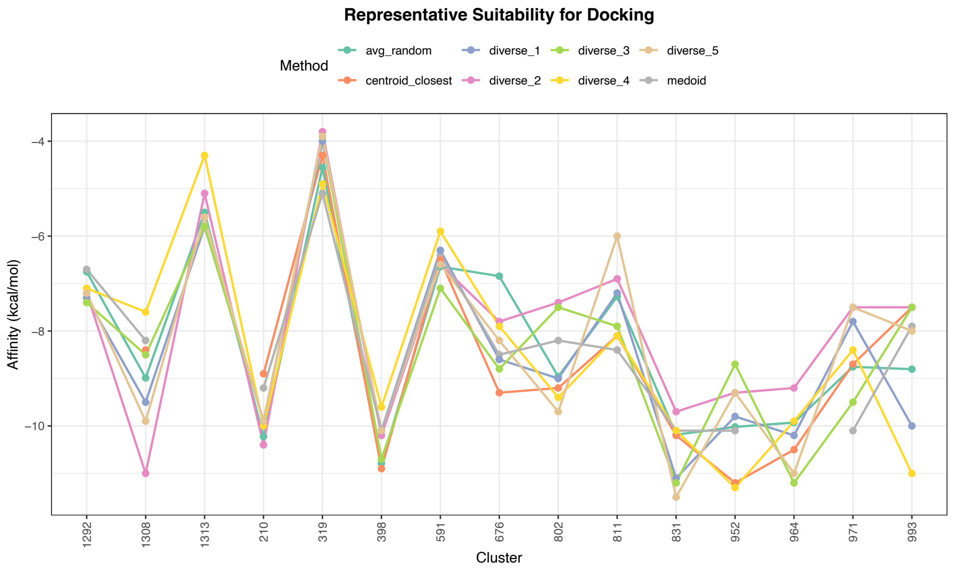


**Supplementary Figure 102: Suitability of Representatives for Docking (Global Analysis)**
The multi-line plot compares the docking scores of different cluster representative types against the average docking score of randomly selected molecules. The x-axis represents the cluster ID, while the y-axis represents binding affinity (kcal/mol). Line colors distinguish the representative types: centroid closest (orange), diverse 1 (blue), diverse 2 (pink), diverse 3 (green), diverse 4 (yellow), diverse 5 (camel), and medoid (grey). The dark green line represents the average docking score of 25 randomly selected molecules from each cluster, providing a baseline for comparison.
